# An Arm-to-Disarm Strategy to Overcome Phenotypic AMR in *Mycobacterium tuberculosis*

**DOI:** 10.1101/2023.03.23.533925

**Authors:** T. Anand Kumar, Shalini Birua, M. Sharath Chandra, Piyali Mukherjee, Samsher Singh, Grace Kaul, Abdul Akhir, Sidharth Chopra, Jennifer Hirschi, Amit Singh, Harinath Chakrapani

**Affiliations:** Department of Chemistry, Indian Institute of Science Education and Research (IISER), Pune, India; Division of Microbiology and Cell Biology, Indian Institute of Science, Bangalore, India; Department of Chemistry, Binghamton University, New York, USA; Division of Molecular Microbiology and Immunology, CSIR-Central Drug Research Institute, Janakipuram Extension, Sitapur Road, Lucknow-226031, Uttar Pradesh, India; Academy of Scientific and Innovative Research (AcSIR), Ghaziabad, India

**Keywords:** Antimicrobial resistance, tuberculosis, persisters, mycobacteria, fluoroquinolone, permeability, drug accumulation, phenotypic antimicrobial resistance

## Abstract

Most front-line tuberculosis drugs are ineffective against hypoxic non-replicating drug-tolerant *Mycobacterium tuberculosis* (*Mtb*) contributing to phenotypic antimicrobial resistance (AMR). This is largely due to the poor permeability in the thick and waxy cell wall of persister cells, leading to diminished drug accumulation and reduced drug-target engagement. Here, using an “arm-to-disarm” prodrug approach, we demonstrate that non-replicating *Mtb* persisters can be sensitized to Moxifloxacin (MXF), a front-line TB drug. We design and develop a series of nitroheteroaryl MXF prodrugs that are substrates for bacterial nitroreductases (NTR), a class of enzymes that are over-expressed in hypoxic *Mtb*. Enzymatic activation involves electron-transfer to the nitroheteroaryl compound followed by protonation via water that contributes to the rapid cleavage rate of the protective group by NTR to produce the active drug. Phenotypic and genotypic data are fully consistent with MXF-driven lethality of the prodrug in *Mtb* with the protective group being a relatively innocuous bystander. The prodrug increased intracellular concentrations of MXF than MXF alone and is more lethal than MXF in non-replicating persisters. Hence, arming drugs to improve permeability, accumulation and drug-target engagement is a new therapeutic paradigm to disarm phenotypic AMR.

## INTRODUCTION

Tuberculosis (TB), a deadly infectious disease caused by *Mycobacterium tuberculosis* (*Mtb*), is among the leading causes of death globally.^1^ Although curable through a chemotherapeutic regimen of 6 months, efforts are on to reduce treatment time, compliance, and relapse while addressing the growing incidences of drug resistance.^1,2^ A fraction of mycobacteria goes into a non-replicating state and become tolerant to clinically relevant TB drugs.^3^ These cells are genetically identical to the rest of the population but are phenotypically slow growing and enter a metabolically quiescent state.^4,5^ This phenotypic form of antimicrobial resistance (AMR) is associated with a reduced metabolic rate, activated stress response, altered cell-wall permeability, and heightened drug-efflux activity as compared to drug-susceptible mycobacteria (Figure 1a).^6–11^ These features of *Mtb* contribute to a protracted TB regimen and plays a significant role in relapse. In a recent promising clinical trial, reduction of therapy duration from 6 months to 4 months was achieved by the inclusion of rifapentine and moxifloxacin (MXF; Figure 1b) in the treatment regimen.^12^ MXF also features in the latest World Health Organization (WHO) guidelines for the treatment of TB in children and adolescents as well as individuals with multi-drug resistant (MDR) infections.^1^ This revised clinical regime is an important milestone for global TB elimination programs but further shortening of chemotherapy remains elusive. This could be due to the refractory nature of non-replicating *Mtb* to front-line drugs including MXF.^13,14^ Since MXF’s target, DNA gyrase is expressed in the non-replicating *Mtb*,^15,16^ diminished drug potency could be due to reduced drug-target engagement.^17–19^ Previous studies in non-replicating *Mtb* persisters support reduced intracellular accumulation of several clinical TB drugs, including MXF. This is attributable to the greater cell wall polysaccharide and lipid content leading to a thickening of the cell wall and a negative zeta potential.^6,13,20^ These attributes of the cell wall in non-replicating persisters work against MXF, which is both hydrophilic (clogP, -0.49) and negatively charged (pKa of the acid is 6.25).^21^ We therefore considered an “arm-to-disarm” approach to tackle this problem. Here, we make a modification to the drug to mask it as a prodrug form – “arming”; the prodrug is designed to have better permeability in non-replicating *Mtb*. Upon entry into *Mtb*, the prodrug is expected to be cleaved or by enzymes such as nitroreductases (NTRs)^22–24^ that are over-expressed in non-replicating mycobacteria. The increased drug accumulation should be lethal and“disarm” non-replicating *Mtb*. Since NTRs retain basal expression in replicating bacteria, the prodrug form is expected to retain potency against these bacteria as well (Figure 1b). We hypothesized that a suitably connected nitro(hetero)aryl group to MXF through an ester or carbamate linker allows for the release of MXF, only after reduction of the nitro group by NTR to an electron rich hydroxylamino or amino group followed by a self-immolation step (Figure 1b). Here, we describe the results of our design, synthesis, and development of prodrugs of MXF that led to the identification of a novel prodrug that undergoes rapid and efficient cleavage under bioreductive conditions to produce MXF; we provide chemical and biochemical insights into prodrug activation and release mechanism, and the lead compound had superior efficacy against non-replicating *Mtb* while retaining lethality against actively replicating *Mtb*.

**Figure 1.**
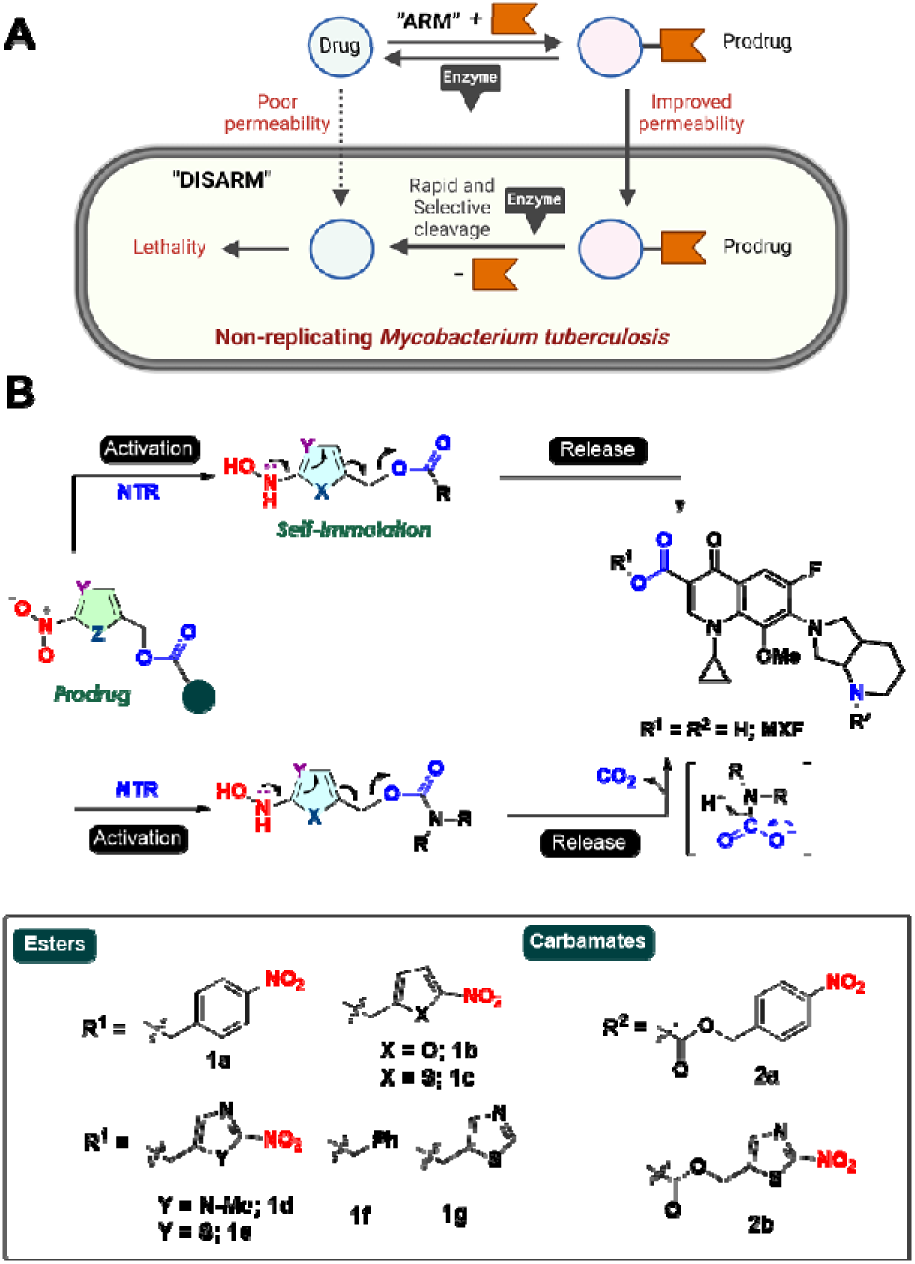
(a) Non-replicating *Mtb* have reduced drug uptake compared with replicating *Mtb*, due to an increased cell wall thickness. Proposed prodrug strategy to increase permeability and drug accumulation in non-replicating *Mtb*. (b) Proposed strategy for the generation of MXF following bioreductive activation of nitroheteroaryl ester and carbamate prodrugs and structures of MXF prodrugs (ester and carbamate) prepared in this study.

## RESULTS

### Synthesis, physicochemical properties, and reduction potential

Esters **1a-1g** (series **1**), with **1f-1g** acting as negative controls, and carbamates **2a-2b** (series **2**) were synthesized as reported in the supplementary information (Schemes S1-S2; Tables S1-S2). As expected, the clogP for derivatives was higher than MXF (Table S3). Next, guided by a recent and comprehensive study on the effect of physicochemical properties on drug accumulation in bacteria, we calculated molar refractivity (MR) and total polar surface area (TPSA) (Table S3).^25^ Both MR and TPSA were significantly higher for both series **1** and **2** when compared with MXF supporting better permeability in bacteria.

### MXF release from prodrugs under chemoreductive and bioreductive conditions

Since MXF is fluorescent, the ability of these derivatives to undergo cleavage under chemoreductive and bioreductive conditions was determined by monitoring MXF’s fluorescence (λ_ex_ = 289 nm and λ_em_ = 488 nm, quantum efficiency φ_F_= 0.21)^26^ (Figure S1). Chemoreductive conditions that simulate a hypoxic environment were next used to study reduction and release. In the presence of sodium dithionite (Na_2_S_2_O_4_) or zinc (Zn)/ammonium formate, yields of MXF were in the range 24-78% (Table 1; Figure S2).

**Table 1.** Reductive activation of MXF and reduction potentials of prodrugs.

| Cpd | Chemored. (%) | | $E^{\circ}_{\text{red}}$ (V) | | Biored. (%) | |
| --- | --- | --- | --- | --- | --- | --- |
| | $\text{Na}_2\text{S}_2\text{O}_4^{\text{a}}$ | $\text{Zn}^{\text{b}}$ | Expt.<sup>c</sup> | Theory<sup>d</sup> | NTR<sup>e</sup> | *Msm*<sup>f</sup> |
| **1a** | 55 | 68 | -0.96 | -0.88 | 13 | 6 |
| **1b** | 48 | 53 | -0.81 | -0.92 | 8 | 27 |
| **1c** | 57 | 56 | -0.88 | -0.96 | 64 | 25 |
| **1d** | 32 | 78 | -0.95 | -1.01 | 7 | 18 |
| **1e** | 53 | 78 | -0.63 | -0.55 | 86 | 61 |
| **2a** | 24 | 29 | -1.21 | -1.24 | 3 | 9 |
| **2b** | 39 | 71 | -0.83 | -1.03 | 49 | 53 |

<sup>a-b</sup>MXF (%) generated under chemoreductive conditions (<sup>a</sup>sodium dithionite (Na<sub>2</sub>S<sub>2</sub>O<sub>4</sub>) and <sup>b</sup>Zn/HCOONH<sub>4</sub>); <sup>c-d</sup>reduction potentials (<sup>c</sup>experimental and <sup>d</sup>computed); <sup>e-f</sup>MXF (%) generated under bioreductive conditions (<sup>e</sup>*E. coli* NTR (15 nM) and NADH (100 μM) and (<sup>f</sup>whole cell lysates of *Msm*).

The negative control compounds did not produce MXF under these conditions (Figure S2): hence, reduction of the nitroaryl group triggered the release of MXF. We computed the reduction potentials of our target molecules at wB97XD/aug-cc-pVDZ//B3LYP/6-31G(d)^27–30^ in acetonitrile against ferrocene as the standard electrode. Indeed, a correlation between the reduction potentials and reactivity was observed (Table 1). To validate our computed reduction potentials, experimental reduction potentials (E°_red,_ Expt.) for compounds **1a**-**2b** were also measured in an aprotic solvent (Figure S3). The computed reduction potentials (Table 1, E°_red_, Theory) agree with the experimentally measured potentials (E°_red_, Expt.) for compounds **1a**-**2b** with a mean deviation of 0.09 V (Figure S4).

Next, we assessed the formation of MXF under bioreductive conditions. In the presence of *E. coli* nitroreductase (NTR; NfsB), yields of MXF ranged from 3-86% (Table 1 and Figure S5A). Lastly, to ascertain the efficiency of MXF generation under bioreductive conditions relevant with mycobacteria, we utilized whole cell lysate of *Mycobacterium smegmatis* (*Msm*) and MXF yields ranging from 6-61% were observed (Table 1 and Figure S5B). As expected, the negative control compounds **1f** and **1g** did not produce MXF under any of the reductive conditions tested (Figure S5).

To better understand the differences in the rate of NTR catalyzed nitroreduction of prodrugs, the kinetics of MXF release was studied by recording the change in fluorescence at different substrate concentrations (Table 2 and Figures S6 and S7). We decided to use an enzyme concentration (15 nM) that is significantly lower than what has been previously reported, which allows us to study, with more physiological relevance, reaction efficiencies and rates.^31–36^ A Michaelis-Menten plot was constructed using initial reaction rates and a significant difference in rates among these compounds was recorded, with **1e** and **2b** being the most preferred substrates (Figure 2). Lineweaver-Burk plots were utilized to determine kinetic parameters (*k_cat_*, *K_m_*, *k_cat_*/*K_m_*and V_max_) that indicate the 2-nitrothiazolyl prodrugs **1e** and **2b** were the most catalytically efficient substrates for *E. coli* NTR, with **1e** giving a larger *k_cat_*/*K_m_* than **2b** (Table 2, entries 5 and 7; Figures S6E and S6G). Surprisingly, the 2-nitroimidazole MXF prodrug **1d**, contrary to previous work on NTR activated similar prodrugs,^31–37^ gave diminished yields of MXF, this discrepancy is likely due to lower concentration of NTR employed in the current study. Hence, the protocols developed in this work provide a protocol for the identification of viable substrates based on the correlation of reduction potential to catalytic efficiency.

**Figure 2.**
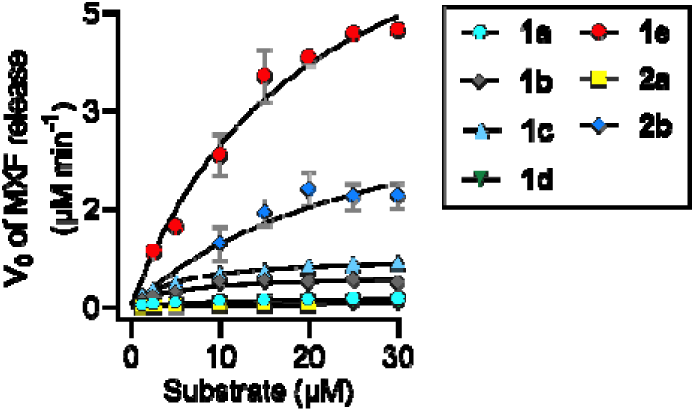
Michaelis-Menten plots of enzyme kinetics using prodrugs **1a-1e**, **2a** and **2b** with *E. coli* NTR. Values are averages (*n* = 3, mean ± SD).

**Table 2.**
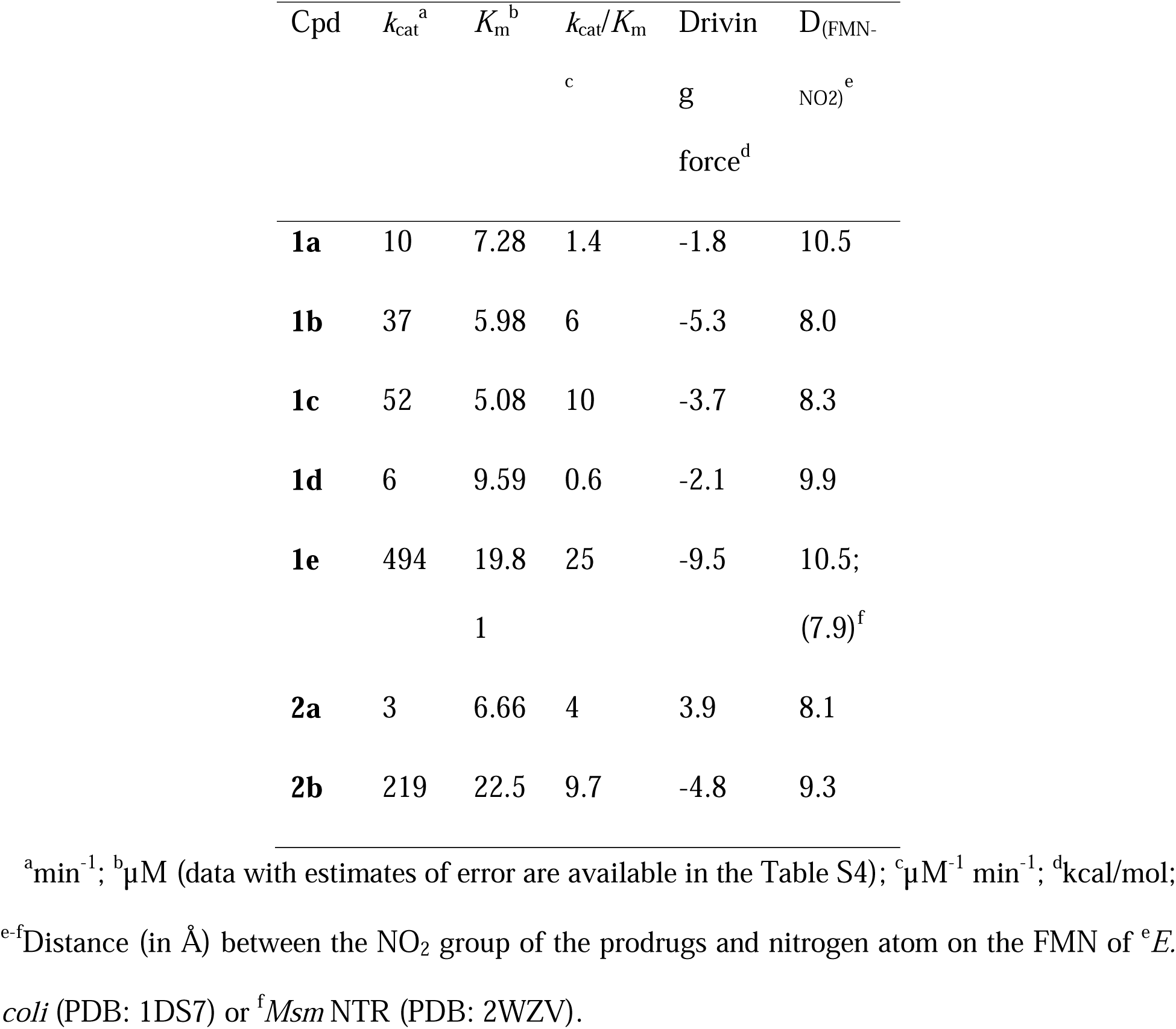
Comparison of experimental kinetic results with computational data for NTR-catalyzed reactions.

### Mechanistic studies

Driving force was computed from the experimentally measured reduction potentials (Table 2) for prodrugs **1a**-**1e** and **2a**-**2b**. A correlation between catalytic efficiency and driving force supports the importance of electron transfer in the mechanism of MXF release (R^2^ = 0.9, see Table 2 and Figure S8). We turned towards computational modelling of the prodrug-enzyme complexes to probe the mechanistic underpinnings of the bioreduction pathway. Prodrugs (**1a**-**1e**, **2a**, **2b**) were docked into the homodimer of an FMN-dependent NTR from *Escherichia coli* (PDB: 1DS7) and *Mycobacterium smegmatis* (PDB: 2WZV) using Swissdock (Figure S9).^38^

Three mechanistic scenarios consistent with the proposal by Wilke and coworkers^39^ for a similar NTR activated prodrug CB1954 were identified as possible pathways for the reduction of prodrugs **1a**-**1e** and **2a**-**2b** (as illustrated in Figure 3b) – Pathway a) proton transfer (PT) followed by a formal hydride transfer (HT), Pathway b) a formal HT followed by a PT, and Pathway c) two consecutive electron transfer (ET)−proton transfer (PT) sequences for the reduction of the nitro groups by NTR. The active sites for each of the final structures obtained from MD simulations were analyzed with these mechanistic scenarios under consideration.

**Figure 3.**
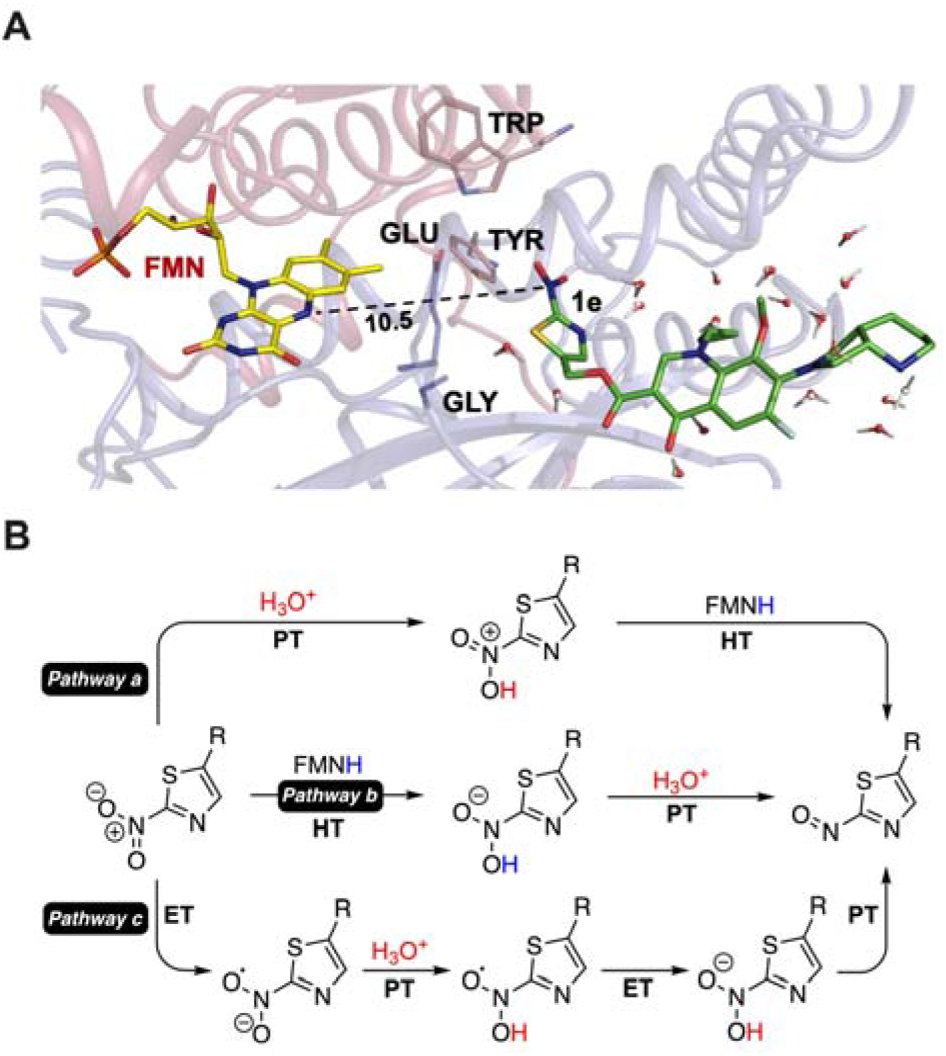
(a) Structure obtained from an MD simulation of **1e** in *E. coli* NTR (PDB: 1DS7). The **1e** active site is sandwiched between the two monomer subunits (blue and red colored cartoons). Key residues are shown. (b) Possible mechanistic scenarios for the reduction of the nitro group of **1e** in the presence of NTR. ET, PT and HT stands for electron transfer, proton transfer and hydride transfer respectively.

The active site for prodrug **1e** in 1DS7 obtained from MD simulations is shown in Figure 3a. The FMN molecule is localized between the two monomer subunits of the homodimer of 1DS7 (the distinct subunits are distinguished in Figure 3a by red and blue ribbons). Several residues (Glu383, Gly384, Tyr362 and Trp139) sterically hinder the prodrug **1e** from binding into close enough proximity of the FMN cofactor to facilitate a direct hydride transfer, eliminating pathways a and b as possible modes of MXF release. Furthermore, the MXF portion of the prodrug is not directly bound within the active site and localizes outside of the binding pocket.

A similar analysis was carried out using an average structure from the last 50 fs of each production run for the enzyme-prodrug complexes of **1a**-**1d** and **2a**-**2b** docked into 1DS7, as well as **1e** into 2WZV. The distances between the FMN cofactor and the nitro group of various prodrugs ranged from 8.0 – 10.5 Å with an average distance of 8.1 Å (Table 2). An overlay of the prodrug-enzyme complexes for 1DS7 suggest that all prodrugs investigated in this study dock and equilibrate similarly in the active site, suggesting that the prodrugs exhibit a similar mode of binding. The RMSD calculated for each of the prodrugs in 1DS7 with respect to **1e**, is between 1.1 Å and 1.5 Å (see SI, Figure S10 for an overlay of the enzyme-prodrug complex from all simulations, Table S5 and S6 for the corresponding RMSD values and binding energies of prodrug-enzyme complexes; Figures S11 and S12 for RMSD and RMSF analysis on **1e** bound to 1DS7). Based on the average distances of the bound prodrug to the FMN coenzyme, obtained from the MD simulations, a direct hydride transfer from FMN is unlikely. Furthermore, the abundance of water (Figure 3a) in the active site supports an electron transfer followed by protonation via water as a possible mechanistic scenario. As demonstrated in Table 2, the driving force exhibits a positive correlation with the catalytic efficiency (*k_cat_*/K_m_) measured for *E. coli* NTR suggesting rate-limiting influence of electron transfer,^40^ further validating our mechanistic hypothesis that the release of MXF occurs through pathway c. These results agree with the observations made by Wilke and coworkers^39^ in similar NTR activated prodrugs.

An analysis of the binding pocket of prodrug-enzyme complex **1e**-1DS7 reveals numerous stabilizing interactions by active site residues (Figure S13). The predominant interactions for **1e** include a stabilizing slip-stacking (4.4 Å) interaction with Phe385. Furthermore, the nearby Asp386 is involved in a bridging hydrogen bonding network with an adjacent water molecule and the two carbonyl groups of prodrug **1e**. There are numerous non-classical hydrogen bonding contacts between the nitro group and Lys142, Trp139, and Try362. Finally, Tyr362 can also participate in a t-stacking interaction with the π-cloud of **1e**. Similar interactions are present in the MD simulations of several other prodrugs, indicating that active site binding is dominated by non-covalent interactions (described in the SI, section 3.14).

### Bioreductive activation and stability of 1e

Based on a threshold of 50% yield in all reductive assays and rates of above 0.5 µM min^-1^, prodrug **1e** was identified as the lead molecule for further evaluation. Next, LC/MS analysis of a reaction mixture containing **1e** and NTR showed nearly quantitative conversion of **1e** to MXF in 30 min (Figure S14). Rapid activation and nearly complete conversion to MXF was seen in less than 30 s under these conditions (Figure S15). The prodrug **1e** remained largely stable in the presence of several oxidative and reductive conditions (Figure S16) as well as in mammalian cells (no MXF formation in 2 h, WT-MEF cells; Figure S17), human blood plasma (>75% recovery in 6 h; Figure S18A) and serum (13% MXF yield in 2 h; Figure S18B). On the contrary, treatment of bacterial lysates derived from *E. coli* and *M. smegmatis* with **1e** produced MXF in a nearly quantitative yield (Figure S19). Heat-treated lysate and lysate treated with NTR inhibitor (dicoumarol; DCOM)^41^ showed diminished yields supporting the role for catalysis by NTR in producing MXF from **1e** (Figure S19).

### Comparison of Potency and Mechanism of 1e and MXF in replicating bacteria

The prodrug was next compared with MXF in replicating *Mtb*. First, minimum inhibitory concentration (MIC) for the prodrug **1e** was determined using Alamar blue (AB) assay. In the laboratory strain, H37Rv, we find that MIC of the prodrug is similar to MXF (Figure 4a). Importantly, the inhibitory activity of the prodrug was comparable to MXF in patient-derived multi-drug resistant (MDR) and extensively drug resistant (XDR) strains (Figure 4a). Next, MXF and the prodrug were compared in a phorbol myristate acetate (PMA)-differentiated THP-1 macrophages infection model, and again, the bactericidal activity of **1e** and MXF were similar suggesting that the prodrug was effective in reducing mycobacterial burden inside macrophages (Figure 4b).

**Figure 4.**
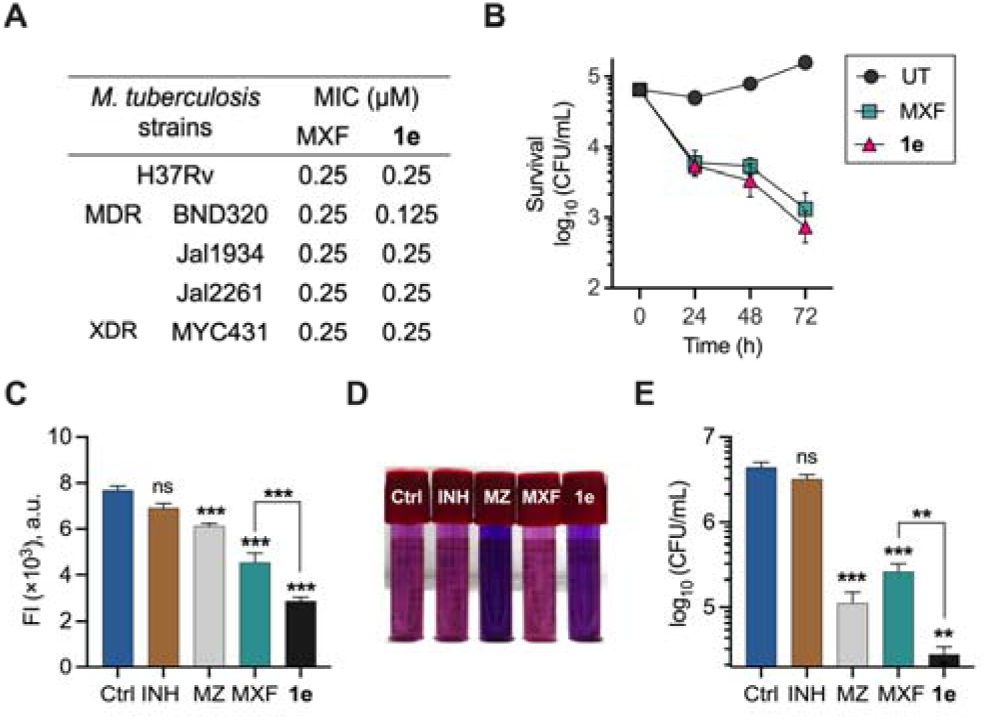
(A) Antimycobacterial activity of MXF and **1e** against drug-susceptible and patient-derived resistant strains of *Mtb*. (B) Survival kinetics after treatment of THP-1 macrophages infected with *Mtb* at 5× MIC of MXF or **1e**. (C) Fluorescence intensity (λ_ex_ = 289 nm and λ_em_ = 488 nm) of PBS starved *Mtb* either left untreated or treated with 20× MIC (5 µM) of MXF and **1e** for 5 days. Statistical significance was calculated against the untreated control (Ctrl) using two-way ANOVA Dunnett (ns indicate not significant, and *\*\*\* p* ≤ 0.001 ***). (D) Mycobactericidal effect of MXF and **1e** at indicated concentrations (10× MIC, 2.5 µM) in a visual hypoxia resazurin reduction assay (HyRRA). Isoniazid (INH, 10 µM) and Metronidazole (MZ, 10 mM) served as a negative and positive control respectively in (C-E). The pink colour in the tube indicates viable bacilli while blue colour indicates no growth. (E) Survival of hypoxic *Mtb* measured as log_10_ of colony-forming units (CFU) after treatment with 10× MIC (2.5 µM) of MXF and **1e** for 5 days. Results are expressed as mean ± SD from three independent experiments performed in triplicate. Statistical significance was established relative to untreated bacteria control (Ctrl) using unpaired *t*-test with Welch’s correction student’s (ns indicate not significant, ** *p* ≤ 0.01, and *\*\*\* p* ≤ 0.001).

The prodrug **1e** was the most potent *Mtb* inhibitor when compared with other analogues, **1a**-**1d** and **2a**-**2b** with only the nitrothiazole derivative **2b** having comparable MIC (Table S7). The control compounds lacking the nitro group, **1f** and **1g** were inactive (Table S7); and the derivative **3e** which had the nitrothiazole group but had a *t*-Boc attached was found to be similarly inactive (Table S8; entry 1). Lastly, the nitrothiazole alcohol **6e** was also found to be a poor inhibitor of *Mtb* (Table S8; entry 2). Together, these data indicate the importance of permeability and cleavage efficiency of the protective group to produce free MXF being critical aspects of efficacy.

We have recently shown that MXF induces oxidative stress and suppresses respiration and carbon catabolism in *Mtb*.^14^ We assessed if the prodrug elicits physiological changes similar to MXF within *Mtb*. First, we used a *Mtb* strain that expresses the redox biosensor Mrx1-roGFP2 from an episomal plasmid (strain Mtb-roGFP2). Mrx1-roGFP2 is a ratiometric fluorescent biosensor that reports the redox potential of the mycothiol redox couple (reduced mycothiol [MSH]/oxidized mycothiol [MSSM]) in the cytoplasm of *Mtb*.^42^ An increase in biosensor ratio indicates oxidative stress, whereas reductive stress decreases the biosensor ratio.^42^ A comparable increase in the biosensor ratio over time with both MXF and **1e** indicated elevated oxidative stress (Figure S20A). Second, using a Seahorse XFp analyzer,^43,44^ we compared dynamic changes in redox state of *Mtb* and found a similar profile in oxygen consumption rate (OCR) (Figure S20B) and extracellular acidification rate (ECAR) (Figure S20C). These phenotypic experiments support similarity in mechanism of action of **1e** and MXF.

For a genotypic comparison, we generated laboratory evolved mutants of *Mtb* in response to MXF and **1e** (Table S9) and confirmed that in both the cases mutations were mapped to the quinolone resistance determinant region (QRDR) of DNA gyrase A (gyrA)^45^ (Figure S21). Similar to MXF, a broad-spectrum antibiotic, the prodrug **1e** exhibited comparable potent activity against ESKAPE pathogens (including clinically derived drug-resistant pathogens), and showed similar efficacy in an animal model of infection (Figures S22-23; Tables S10-12). The lack of inhibitory activity against fluoroquinolone-resistant *E. coli* mutants (Table S13) supported that the prodrug **1e** has a mechanism of action similar to MXF.

### Comparison of 1e and MXF in non-replicating *Mycobacterium tuberculosis*

The prodrug was next compared with MXF by measuring the viability of nutrient-starved non-replicating *Mtb* using the resazurin microtiter assay (REMA). MXF and **1e** both exhibited a significant effect on the viability of nutrient-deprived bacilli, with the prodrug being notably more effective than MXF (Figure 4c). Under the same assay conditions, consistent with literature reports, these nutrient-starved bacilli displayed considerable tolerance to INH and susceptibility to metroindazole (MZ).^13,14,46^ The prodrug was next compared with MXF in non-replicating *Mtb* using hypoxia resazurin reduction assay (HyRRA). MXF had some effect on growth but the compound **1e** was found to have a significantly higher potency when compared with MXF (Figure 4d). Again, INH showed no significant growth inhibition while MZ served as a positive control.^46^ We also measured efficacy of **1e** by enumerating colony forming units (CFUs). Consistent with the HyRRA, MZ was more bactericidal than MXF, and **1e** was significantly better than both these compounds (Figure 4e). Collectively, these studies illustrate that the prodrug was more effective than MXF against non-replicating bacilli in both nutrient-starved and hypoxia models.

### Intracellular accumulation of MXF and 1e

A modified LC/MS protocol^13,47^ (Figure 5a; Figure S24 for detailed steps 1-8) was developed to quantitate the amount of MXF generated by replicating *Mtb* with 5 µM of MXF or **1e**. After 30 min of incubation, the bacilli were pelleted through silicone oil, lysed and then compounds were extracted from the resulting lysate (Figure 5a).

**Figure 5.**
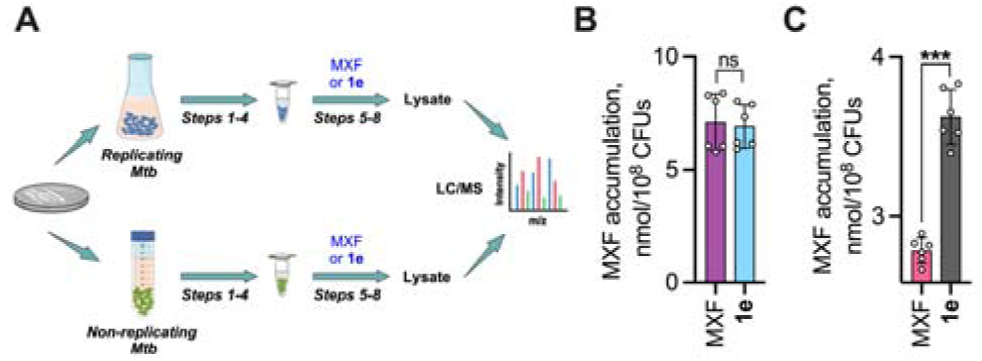
(A) Workflow for Accumulation assay. (B-C) LC/MS analysis of intracellular concentrations (nmol/CFU) of MXF (m/z 402.18) in (B) replicating and (C) non-replicating *Mtb* following incubation for 30 min with 5 μM of MXF or **1e**. The drug content is expressed as the amount of drug (nmol) per CFU. Statistical significance was established relative to MXF treated sample using unpaired *t*-test (ns indicate not significant and *\*\*\* p* ≤ 0.001).

The intracellular concentrations were determined using a standard calibration curve (Figures S25 and S26), was converted into accumulation values, then normalized with the bacterial count and were finally represented as MXF accumulation (nmol/10^8^ CFUs). Nearly identical values for MXF accumulation with free MXF as well as **1e** was recorded for replicating *Mtb* (Figure 5b). The intact prodrug **1e** could not be detected under these conditions, possibly due to rapid cleavage upon entry into cells. A similar workflow (Figure 5a and Figure S26) with free MXF in non-replicating *Mtb* gave diminished MXF accumulation, which is consistent with the previous report. With **1e** in non-replicating *Mtb*, we find a substantially higher MXF accumulation, with no evidence for intact **1e** (Figure 5c). The prodrug **1e** was stable and did not produce MXF in extracellular media derived from overnight cultured mycobacteria suggesting that activation of **1e** likely occurred upon permeation and intracellular activation (Figure S27).

## DISCUSSION

Given that drug accumulation is uniformly low across all anti-TB drugs,^13^ enhancing concentrations of the drug through an enzyme-prodrug approach can tackle phenotypic AMR and eliminate persisters. To identify a prodrug candidate that is sensitive to hypoxia and NTR-catalysis, a screen was conducted with various chemoreductive conditions as well as a significantly lower concentration of NTR. This comprehensive analysis gave 2-nitrothiazole ester prodrug **1e** as the lead candidate. This prodrug had superior sensitivity and drug release efficiency over other masked dertivates tested. Further insights into binding, orientation, and mechanism of prodrug reduction provide a robust framework for NTR-based prodrug design. Hence, together, a combination of experimental and computational data yields an excellent model for understanding catalysis in *E. coli* NTR. These findings can be extrapolated to mycobacterial NTR as well, and will therefore help identify similarly efficient prodrugs of other clinical TB drugs. A key feature of the developed prodrug **1e** is its efficient activation within replicating *Mtb* to produce MXF at levels that are comparable to free MXF and equipotency in several *in vitro* studies and in an animal model of infection with an ESKAPE pathogen. This prodrug significantly enhanced the accumulation and potency of active drug MXF in non-replicating *Mtb*. Since hypoxic non-replicating *Mtb* develops a thick outer layer that restricts the entry of anti-TB drugs,^6,10,13,20^ our study provides an example of how improving permeability and accumulation can lead to favorable outcomes.^48–50^ Our data suggest that the mechanism of killing by **1e** is similar to MXF in *Mtb in vitro*. MXF blocks the growth by forming reversible drug-gyrase complexes that rapidly inhibit DNA replication, followed by death due to chromosomal fragmentation, respiration modulation, and ROS.^14^ However, the clinical situation is complicated due to poor penetration of MXF into the caseous region of tubercular granulomas^51,52^ and diminished permeability inside hypoxic *Mtb*. Low local MXF concentrations likely promote the widely reported emergence of MXF resistance in TB patients.^53^ Thus, greater permeability and bioreductive activation of **1e** in the hypoxic non-replicating bacilli could increase the probability that MXF clears persisters, shorten TB therapy time, and reverse the emergence of MXF resistance. Previous studies that have developed prodrugs with a goal of improving permeability of the parent drug or drug-like molecule have focused on replicating pathogens whose characteristics are very different from non-replicating mycobacteria.^54,55^ Many TB drug discovery strategies screen for activity against replicating *Mtb* and it may take several years for the evaluation of new drugs in clinical settings, especially against persisters. Therefore, utilization of prodrug strategies to enhance the efficacy of existing standard TB drugs offers several advantages. Firstly, repurposing well-tolerated and widely used drugs reduces the potential concerns associated with new drug development. This approach also allows us to specifically target non-replicating bacteria while retaining efficacy in replicating bacteria and will thus improve the treatment outcomes. Given that the safety, efficacy, tolerability and toxicity of MXF is well documented, and this drug has been safely used in a large number of individuals, this first-in-class proof-of-concept study where MXF is armed to disarm non-replicating *Mtb* is a promising strategy in our efforts to treat this deadly pathogen.

## ASSOCIATED CONTENT

### Supporting Information

Experimental methods, spectra, assay protocols and other associated raw data. This material is available free of charge via the Internet at http://pubs.acs.org.

## AUTHOR INFORMATION

### Author Contributions

The manuscript was written through contributions of all authors. All authors have given approval to the final version of the manuscript.

## ACKNOWLEDGMENT

Financial support for this project was from IISER Pune, Department of Science and Technology (DST) Fund for Improvement of S&T Infrastructure (grant number SR/FST/LSII-043/2016) to the IISER Pune Biology Department for setting up the Biological Mass Spectrometry Facility and Ignite Life Sciences for funding LC/MS studies. Research fellowship for TAK and GK (DST-INSPIRE) and AA (UGC) are acknowledged. The authors thank Mr. Arnab Chakraborty (Dr. Siddhesh S. Kamat Lab, IISER Pune) and Mrs. Sunita (LC/MS facility in-charge, IISc, Bangalore). Financial support from NIH R35GM147183 (J.S.H.), and the XSEDE Science Gateway Program (under the NSF Grant Numbers ACI-1548562, CHE180061, and CHE210031) (J.S.H. and S.C.M.) was acknowledged.

---

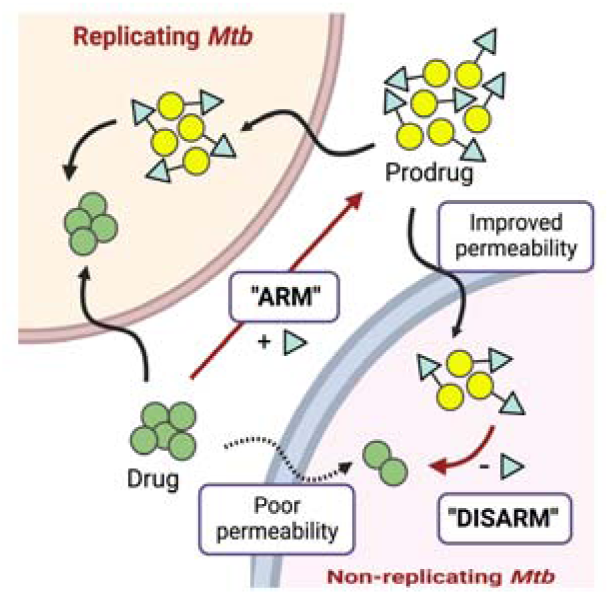

## Notes

### Competing Interest Statement

The authors have declared no competing interest.

### Summary of Updates

(1) Title, abstract, introduction, Fig 1a, discussion, conclusion, and TOC are updated to reflect arm-to-disarm prodrug strategy and demonstrate that non-replicating Mtb persisters can be sensitized to Moxifloxacin (MXF). (2) Significant changes were made to reduce the number of figures (Figures 1-7) in the revised manuscript (Figures 1-5) (2a) Synthetic schemes, and Figure 3 were moved to Supplementary File. Figure 4 was updated with a new figure (Figure 3 in revised manuscript) to probe the mechanistic underpinnings of the bioreduction pathway through MD simulations and possible mechanistic scenarios. (2b) Figure 5 and Figure 6 were merged and updated with a new figure (Figure 4) to reflect new results (the activity of MXF prodrug against both nutrient-starved and hypoxic non-replicating Mtb). (2c) Figure 7 was made to Figure 5. (3) Author affliations updated (4) Supplemental files updated

