## Supporting information for "An Arm-to-Disarm Strategy to Overcome Phenotypic AMR in *Mycobacterium tuberculosis*"

### for

### Table of contents


| | | |
|---|---|---|
| **1.** | **Synthesis and characterization** | S3 |
| **2.** | **Purity of prodrugs by HPLC** | S19 |
| **3.** | **Experimental protocols** | S23 |
| **4.** | **Figures** | S39 |
| **5.** | **Tables** | S55 |
| **6.** | **References** | S63 |
| **7.** | **NMR Spectral data** | S65 |
| **8.** | **HRMS Spectra of final compounds** | S104 |

#### 1. Synthesis and characterization

Preparation of ester prodrugs of MXF **(1a-1g):** The *t*-Boc protected MXF derivative **4** was prepared using a reported procedure.[1] Esterification of this compound using the corresponding nitroaryl or nitroheteroaryl alcohol afforded analogues **3a-3e**. Deprotection of the *t*-Boc group gave the desired compounds **1a-1e (Scheme S1)**. A similar two-step method was used to prepare compounds lacking nitro group **1f-1g (Scheme S1)**. Compounds **6b-6e** were synthesized following previously reported protocols and the analytical data for each compound was consistent with reported values.[2-5]

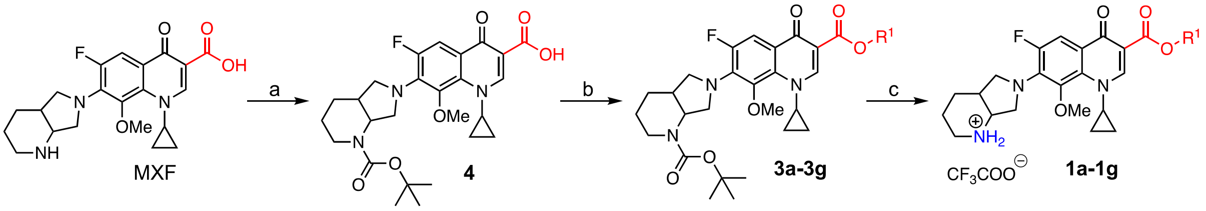


**Scheme S1.** Synthesis of **1a-1g**. *Reaction conditions*: (a) $(Boc)_2O$, NaOH, $H_2O$:dioxane (1:1 v/v); (b) $R^1OH$, HBTU, DMAP, DIPEA, DCM; (c) TFA, DCM.

##### A. General procedure for synthesis of *N*-boc protected esters of MXF (3a-3g)

To a stirred solution of **4**, HBTU (1.96 mmol) and DMAP (0.19 mmol) in dry DCM (100 mL), DIPEA (1.96 mmol) was added. The reaction mixture was stirred for 30 min at RT. The resulting solution turned yellow. To this reaction mixture, alcohol was added and heated to 38 °C for 12 h until the starting material was completely consumed (as monitored by TLC). After completion of the reaction, water (100 mL) was added and the aqueous layer was extracted with DCM (3 × 100 mL). The organic extracts were combined, washed with brine (2 × 50 mL), dried over anhydrous $Na_2SO_4$, filtered and the filtrate was concentrated under reduced pressure. The residue was purified by preparative HPLC using Kromasil®C-18 column at ambient temperature under the gradient elution with $H_2O$:ACN (20:80, v/v) as mobile phase at a flow rate of 12 mL/min.

**4-nitrobenzyl 7-((4aS,7aS)-1-(*tert*-butoxycarbonyl)hexahydro-1H-pyrrolo[3,4-b] pyridin-6(2H)-yl)-1-cyclopropyl-6-fluoro-8-methoxy-4-oxo-1,4-dihydroquinoline-3-carboxylate (3a):** Starting from **4** (0.197 g, 0.39 mmol) and commercially available **6a** (0.090 g, 0.58 mmol), **3a** (0.183 g, 73%) was isolated as a white solid; mp = 127-128 °C; $R_f$ (EtOAc:hexane = 70:30) = 0.4; FT-IR ($\nu_{max}$, $cm^{-1}$): 2924 (C-H stretch), 1729 (ester C=O

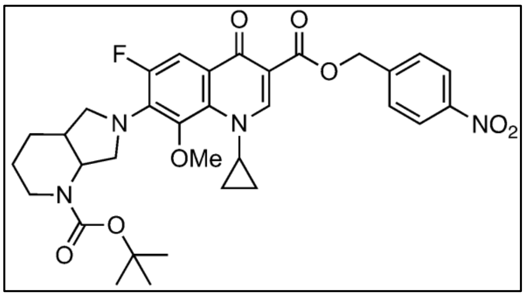


stretch), 1691 (conjugated ketone C=O stretch), 1457 (asymmetric $NO_2$ stretch), 1359 (symmetric $NO_2$ stretch); $^{19}F$ NMR (376 MHz, $CDCl_3$): δ -123.3; $^{1}H$ NMR (400 MHz, $CDCl_3$) : δ 8.58 (s, 1H), 8.26 – 8.22 (m, 2H), 7.85 (d, *J* = 14.2 Hz, 1H), 7.72 (d, *J* = 8.6 Hz, 2H), 5.52 – 5.44 (m, 2H), 4.78 (br, 1H), 4.08 – 4.03 (m, 2H), 3.93 – 3.88 (m, 1H), 3.84 (dt, *J* = 11.2, 4.9 Hz, 1H), 3.56 (s, 3H), 3.37 (br, 1H), 3.22 (s, 1H), 2.88 (t, *J* = 10.9 Hz, 1H), 2.28 – 2.21 (m, 1H), 1.77 (s, 2H), 1.48 (s, 11H), 1.24 – 1.22 (m, 1H), 1.08 – 1.00 (m, 2H), 0.84 – 0.76 (m, 1H); $^{13}C$ NMR (100 MHz, $CDCl_3$): δ 161.0, 155.4, 151.1, 147.6, 144.1, 140.6, 136.2, 134.6, 133.4, 130.9, 128.0, 123.9, 108.9 (d, *J* = 122.0 Hz), 80.2, 65.1, 61.2, 56.4, 56.3, 56.0, 39.9, 35.9, 28.6, 25.5, 24.3, 10.6, 8.5; HRMS (ESI-TOF) for $C_{33}H_{37}FN_4O_8$ $[M+H]^+$ : Calcd., 637.2673, Found, 637.2673.

**(5-nitrofuran-2-yl)methyl 7-(-1-(*tert*-butoxycarbonyl)hexahydro-1H-pyrrolo [3,4-b] pyridin-6(2H)-yl)-1-cyclopropyl-6-fluoro-8-methoxy-4-oxo-1,4-dihydroquinoline-3-carboxylate (3b):** Starting from **4** (0.160 g, 0.31 mmol) and **6b** (0.068 g, 0.47 mmol), **3b** (0.110 g, 56%) was isolated as a white solid; mp = 101-102 ºC; $R_f$ (EtOAc:hexane = 70:30) = 0.5; FT-IR ($\nu_{max}$, $cm^{-1}$): 2988 (C-H stretch), 1732 (ester C=O stretch), 1625 (conjugated ketone C=O stretch), 1449 (asymmetric $NO_2$ stretch), 1373 (symmetric $NO_2$ stretch); $^{19}F$ NMR (376 MHz, $CDCl_3$): δ -122.7; $^{1}H$ NMR (400 MHz, $CDCl_3$) : δ 8.57 (s, 1H), 7.78 (d, *J* = 14.0 Hz, 1H), 7.30 (s, 1H), 6.80 (d, *J* = 3.6 Hz, 1H), 5.40 – 5.31 (m, 2H), 4.78 (br, 1H), 4.07 – 4.02 (m, 2H), 3.94 - 3.81 (m, 1H), 3.84 (t, *J* = 9.6 Hz, 1H), 3.57 (s, 3H), 3.39 - 3.36 (m, 1H), 3.21 (d, *J* = 9.1 Hz, 1H), 2.88 (t, *J* = 11.2 Hz, 1H), 2.28 - 2.22 (m, 1H), 1.81 - 1.75 (m, 2H), 1.48 (s, 10H), 1.30 – 1.19 (m, 2H), 1.12 – 1.00 (m, 2H), 0.86 - 0.77 (m, 1H) ; $^{13}C$ NMR (100 MHz, $CDCl_3$): δ 172.8, 164.8, 155.4, 153.5, 152.3 (d, $J_{C-F}$ = 161.2 Hz), 151.0, 141.5, 136.0 (d, $J_{C-F}$ = 43.1 Hz), 133.3, 122.1, 113.5, 112.5, 108.8 (d, $J_{C-F}$ = 94.7 Hz), 108.5, 80.2, 60.9, 57.7, 56.4, 56.3, 39.8, 35.7, 28.5, 25.5, 24.3, 10.5, 8.5; DEPT-135 NMR (100 MHz, $CDCl_3$): δ 151.0 (CH), 113.6 (CH), 112.5 (CH), 108.5 (CH), 60.9 ($CH_3$), 57.6 ($CH_2$), 56.4 ($CH_2$), 56.3 ($CH_2$), 39.7 (CH), 35.7 (CH), 28.8 ($CH_3$), 25.8 ($CH_2$), 24.4 ($CH_2$), 10.5 ($CH_2$), 8.5 ($CH_2$); HRMS (ESI-TOF) for $C_{31}H_{35}FN_4O_9$ $[M+H]^+$ : Calcd., 627.2466, Found, 627.2469.

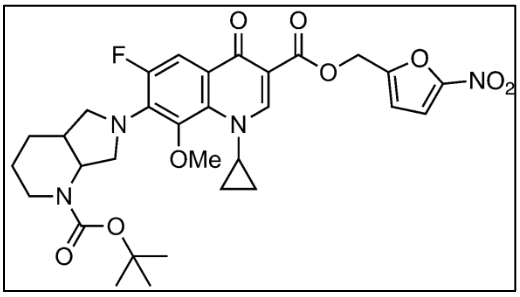

**(5-nitrothiophen-2-yl)methyl-7-(-1-(*tert*-butoxycarbonyl)hexahydro-1H-pyrrolo[3,4-b]pyridin-6(2H)-yl)-1-cyclopropyl-6-fluoro-8-methoxy-4-oxo-1,4-dihydro quinoline-3-carboxylate (3c):** Starting from **4** (0.150 g, 0.30 mmol) and **6c** (0.071 g, 0.44 mmol), **3c** (0.114 g, 59%) was isolated as a white solid; mp = 97-98 ºC; $R_f$ (EtOAc:hexane = 70:30) = 0.5; FT-IR ($\nu_{max}$, cm$^{-1}$): 1732 (ester C=O stretch), 1684 (conjugated ketone C=O stretch), 1455 (asymmetric $NO_2$ stretch), 1327 (symmetric $NO_2$ stretch); $^{19}$F NMR (376 MHz, $CDCl_3$): δ -122.5; $^1$H NMR (400 MHz, $CDCl_3$): δ 8.58 (s, 1H), 7.84 – 7.80 (m, 2H), 7.15 (d, $J$ = 4.1 Hz, 1H), 5.52 – 5.44 (m, 2H), 4.77 (br, 1H), 4.08 – 4.02 (m, 2H), 3.94 – 3.88 (m, 1H), 3.83 (td, $J$ = 9.9, 2.1 Hz, 1H), 3.56 (s, 3H), 3.36 (br, 1H), 3.21 (d, $J$ = 9.8 Hz, 1H), 2.88 (t, $J$ = 12.2 Hz, 1H), 2.28 – 2.21 (m, 1H), 1.82 – 1.75 (m, 2H), 1.48 (s, 10 H), 1.28 – 1.22 (m, 2H), 1.11 – 1.01 (m, 2H), 0.85 – 0.78 (m, 1H); $^{13}$C NMR (100 MHz, $CDCl_3$): δ 172.7, 165.5, 155.4, 152.3 (d, $J_{C\text{-}F}$ = 144.6 Hz), 151.1, 147.0, 141.5, 136.0 (d, $J_{C\text{-}F}$ = 44.6 Hz), 133.3, 128.4, 126.9, 122.1, 122.0, 108.9 (d, $J_{C\text{-}F}$ = 95.5 Hz), 108.6, 100.1, 80.2, 60.9, 60.7, 56.5, 56.4, 56.3, 39.9, 35.8, 28.6, 25.5, 24.3, 10.5, 8.5; DEPT-135 NMR (100 MHz, $CDCl_3$): δ 151.1 (CH), 128.4 (CH), 126.9 (CH), 109.0 (CH), 61.0 ($CH_3$), 60.7 ($CH_2$), 56.4 ($CH_2$), 56.3 ($CH_2$), 39.9 (CH), 35.7 (CH), 28.6 ($CH_3$), 25.5 ($CH_2$), 24.3 ($CH_2$), 10.5 ($CH_2$); HRMS (ESI-TOF) for $C_{31}H_{35}FN_4O_8S$ $[M+H]^+$ : Calcd., 643.2238, Found, 643.2242.

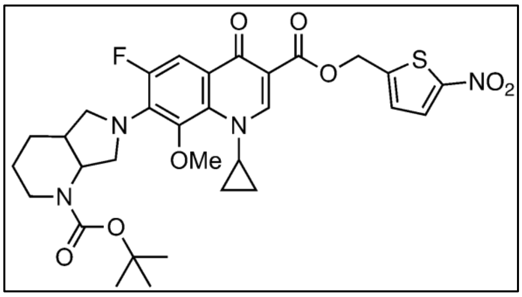


**(1-methyl-2-nitro-1H-imidazol-5-yl)methyl 7-(-1-(*tert*-butoxy carbonyl) octa hydro-6H-pyrrolo[3,4-b]pyridin-6-yl)-1-cyclopropyl-6-fluoro-8-methoxy-4-oxo-1,4-dihydroquinoline-3-carboxylate (3d):** Starting from **4** (0.170 g, 0.33 mmol) and **6d** (0.080 mg, 0.50 mmol), **3d** (0.124 mg, 57%) was isolated as a gray solid; mp = 144-145 ºC; $R_f$ (MeOH:$CHCl_3$ = 05:95) = 0.2; FT-IR ($\nu_{max}$, cm$^{-1}$): 2974 (C-H stretch), 1731 (ester C=O stretch), 1684 (conjugated ketone C=O stretch), 1539 (asymmetric $NO_2$ stretch), 1364 (symmetric $NO_2$ stretch); $^{19}$F NMR (376 MHz, $CDCl_3$): δ -122.7; $^1$H NMR (400 MHz, $CDCl_3$) : δ 8.54 (s, 1H), 7.76 (d, $J$ = 14.1 Hz, 1H), 7.27 (d, $J$ = 10.1 Hz, 1H), 5.36 (q, $J$ = 13.6 Hz, 2H), 4.77 (s, 1H), 4.21 (s, 3H), 4.07 – 4.02 (m, 2H), 3.93 – 3.88 (m, 1H), 3.83 (td, $J$ = 10.0, 2.7 Hz, 1H), 3.56 (s, 3H), 3.35 (br, 1H), 3.21 (d, $J$ = 8.6 Hz, 1H), 2.88 (t, $J$ = 11.2 Hz, 1H), 2.27 – 2.22 (m, 1H), 1.80 – 1.75 (m, 2H), 1.48

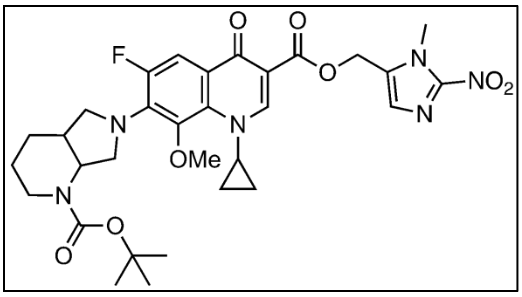

(s, 10H), 1.32 – 1.19 (m, 2H), 1.10 – 1.01 (m, 2H), 0.84 – 0.76 (m, 1H); $^{13}$C NMR (100 MHz, $CDCl_3$): δ 172.6, 165.5, 155.4, 155.0, 152.5, 151.1, 146.3, 141.5 (d, $J_{C\text{-}F}$ = 25.0 Hz), 136.1 (d, $J_{C\text{-}F}$ = 44.0 Hz), 133.3, 132.6, 129.6, 122.0, 121.9, 108.9, 108.6, 80.2, 60.9, 56.4, 56.3, 55.2, 39.9, 35.7, 34.7, 28.5, 25.4, 24.3, 10.5, 8.5; DEPT-135 NMR (100 MHz, $CDCl_3$): δ 151.0 (CH), 129.5 (CH), 108.6 (CH), 60.8 ($CH_3$), 56.3 ($CH_2$), 56.2 ($CH_2$), 55.1 ($CH_2$), 39.8 (CH), 35.6 (CH), 34.6 ($CH_3$), 28.4 ($CH_3$), 25.3 ($CH_2$), 24.2 ($CH_2$), 10.4 ($CH_2$), 8.4 ($CH_2$); HRMS (ESI-TOF) for $C_{31}H_{37}FN_6O_8$ $[M+H]^+$ : Calcd., 641.2735, Found, 641.2737.

**(2-nitrothiazol-5-yl)methyl 7-(-1-(*tert*-butoxycarbonyl)octahydro-6H-pyrrolo [3,4-b]pyridin-6-yl)-1-cyclopropyl-6-fluoro-8-methoxy-4-oxo-1,4-dihydroquinoline-3-carboxylate (3e):** Starting from **4** (0.125 g, 0.25 mmol) and **6e** (0.060 g, 0.37 mmol), **3e** (0.125 g, 78%) was isolated as yellowish white solid; mp = 110-111 ºC; FT-IR ($\nu_{max}$, cm$^{-1}$): 2992 (C-H stretch), 1739 (ester C=O stretch), 1681 (conjugated ketone C=O stretch), 1358 (symmetric $NO_2$ stretch); $R_f$ (EtOAc:hexane = 70:30) = 0.5; $^{19}$F NMR (376 MHz, $CDCl_3$): δ -122.6; $^1$H NMR (400 MHz, $CDCl_3$) : δ 8.58 (s, 1H), 7.91 (s, 1H), 7.80 (d, $J$ = 14.0 Hz, 1H), 5.54 (s, 2H), 4.76 (s, 1H), 4.07 – 4.02 (m, 2H), 3.94 – 3.89 (m, 1H), 3.83 (td, $J$ = 9.8, 1.9 Hz, 1H), 3.56 (s, 3H), 3.36 (s, 1H), 3.21 (d, $J$ = 9.2 Hz, 1H), 2.88 (t, $J$ = 11.5 Hz, 1H), 2.28 – 2.23 (m, 1H), 1.80 – 1.75 (m, 2H), 1.48 (s, 11H), 1.32 – 1.23 (m, 1H), 1.11 – 1.00 (m, 2H), 0.83 – 0.77 (m, 1H); $^{13}$C NMR (100 MHz, $CDCl_3$): δ172.6, 166.8, 165.8, 155.4, 151.3, 142.1, 142.0, 136.1 (d, $J_{C\text{-}F}$ = 43.7 Hz), 133.3, 122.0, 108.8 (d, $J_{C\text{-}F}$ = 95.0 Hz), 108.2, 80.2, 60.9, 58.1, 56.4, 56.3, 39.9, 35.7, 28.6, 25.5, 24.3, 10.5, 8.5; DEPT-135 NMR (100 MHz, $CDCl_3$): δ 156.0 (CH), 141.9 (CH), 108.8 (CH), 60.8 ($CH_3$), 58.0 ($CH_2$), 56.2 ($CH_2$), 39.8 (CH), 35.6 (CH), 28.4 ($CH_3$), 25.4 ($CH_2$), 24.2 ($CH_2$), 10.4 ($CH_2$), 8.4 ($CH_2$); HRMS (ESI-TOF) for $C_{30}H_{34}FN_5O_8S$$[M+H]^+$ : Calcd., 644.2190, Found, 644.2184.

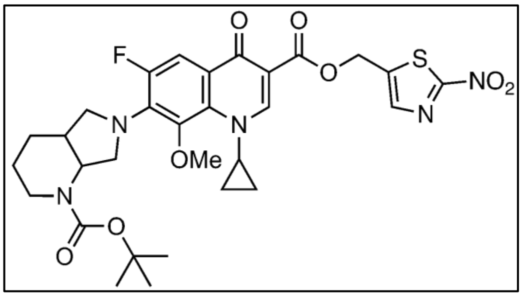


**Benzyl 7-(-1-(*tert*-butoxycarbonyl)hexahydro-1H-pyrrolo[3,4-b]pyridin-6(2H)-yl)-1-cyclopropyl-6-fluoro-8-methoxy-4-oxo-1,4-dihydroquinoline-3-carboxylate (3f):** Starting from **4** (0.150 g, 0.30 mmol) and **6f** (0.048 g, 0.44 mmol), **3f** (0.115 g, 65%) was isolated as a white solid; mp = 109-110 ºC; $R_f$ (EtOAc:hexane = 70:30) = 0.5; FT-IR ($\nu_{max}$, cm$^{-1}$): 2932 (C-H stretch), 1685 (conjugated ketone C=O stretch), 1614 (conjugated ketone C=O stretch); $^{19}$F

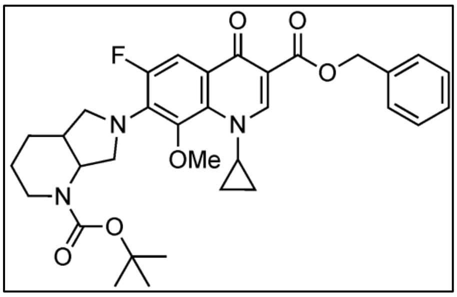

NMR (376 MHz, $CDCl_3$): δ -123.2; $^{1}$H NMR (400 MHz, $CDCl_3$): δ 8.55 (s, 1H), 7.84 (d, *J* = 14.2 Hz, 1H), 7.53 – 7.50 (m, 2H), 7.39 – 7.35 (m, 2H), 7.32 – 7.29 (m, 1H), 5.42 – 5.34 (m, 2H), 4.76 (s, 1H), 4.06 – 4.01 (m, 2H), 3.89 – 3.80 (m, 2H), 3.55 (s, 3H), 3.35 (s, 1H), 3.20 (d, *J* = 8.4 Hz, 1H), 2.88 (t, *J* = 12.0 Hz, 1H), 2.27 – 2.20 (m, 1H), 1.83 – 1.75 (m, 4H), 1.48 (s, 9H), 1.24 – 1.17 (m, 1H), 1.08 – 0.97 (m, 2H), 0.80 – 0.73 (m, 1H); $^{13}$C NMR (100 MHz, $CDCl_3$): δ 173.0, 165.7, 155.4, 150.7, 136.6, 135.8 (d, $J_{C\text{-}F}$ = 44.2 Hz), 133.4, 128.6, 128.1, 128.0, 122.2, 122.1, 109.7, 108.9 (d, $J_{C\text{-}F}$ = 96.5 Hz), 80.2, 66.4, 60.9, 56.4, 56.3, 39.6, 35.7, 28.6, 25.5, 24.3, 10.4, 8.5; DEPT-135 NMR (100 MHz, $CDCl_3$): δ 150.7 (CH), 128.6 (CH), 128.0 (CH), 127.9 (CH), 108.9 (CH), 66.4 ($CH_2$), 60.8 ($CH_3$), 56.3 ($CH_2$), 56.2 ($CH_2$), 39.6 (CH), 35.7 (CH), 28.5 ($CH_3$), 25.4 ($CH_2$), 24.3 ($CH_2$), 10.4 ($CH_2$), 8.5 ($CH_2$); HRMS (ESI-TOF) for $C_{33}H_{38}FN_3O_6$ $[M+H]^+$ : Calcd., 592.2817, Found, 592.2818.

**Thiazol-5-ylmethyl 7-(1-(*tert*-butoxycarbonyl)octahydro-6*H*-pyrrolo[3,4-*b*]pyridin-6-yl)-1-cyclopropyl-6-fluoro-8-methoxy-4-oxo-1,4-dihydroquinoline-3-carboxylate (3g):**

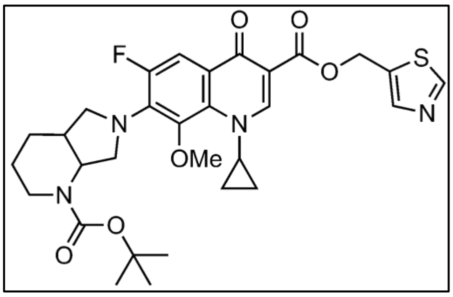

Starting from **4** (0.130 g, 0.26 mmol) and **6g** (0.045 g, 0.38 mmol), **3g** (0.105 g, 67%) was isolated as yellowish white solid; mp = 129-130 ºC; $R_f$ (EtOAc:hexane = 70:30) = 0.5; FT-IR ($\nu_{max}$, cm$^{-1}$): 2971 (C-H stretch), 1742 (ester C=O stretch), 1668 (conjugated ketone C=O stretch); $^{19}$F NMR (376 MHz, $CDCl_3$): δ -123.0; $^{1}$H NMR (400 MHz, $CDCl_3$): δ 8.80 (s, 1H), 8.54 (s, 1H), 7.97 (s, 1H), 7.81 (d, *J* = 14.2 Hz, 1H), 5.60 – 5.53 (m, 2H), 4.76 (s, 1H), 4.07 – 4.01 (m, 2H), 3.90 – 3.85 (m, 1H), 3.83 (td, *J* = 9.8, 2.0 Hz, 1H), 3.55 (s, 3H), 3.35 (s, 1H), 3.20 (d, *J* = 8.2 Hz, 1H), 2.87 (t, *J* = 11.7 Hz, 1H), 2.28 – 2.20 (m, 1H), 1.81 – 1.75 (m, 2H), 1.48 (s, 11H), 1.26 – 1.19 (m, 1H), 1.07 – 0.97 (m, 2H), 0.81 – 0.74 (m, 1H); $^{13}$C NMR (100 MHz, $CDCl_3$): δ 172.7, 165.4, 155.5, 155.0, 154.7, 152.5, 150.9, 144.0, 141.5 (d, $J_{C\text{-}F}$ = 24.3 Hz), 135.9 (d, $J_{C\text{-}F}$ = 43.9 Hz), 133.3, 122.2, 122.1, 109.0, 108.8, 80.2, 60.9, 58.0, 56.4, 56.3, 39.7, 35.7, 28.5, 25.5, 24.3, 10.4, 8.5; DEPT-135 NMR (100 MHz, $CDCl_3$): δ 150.8 (CH), 143.9 (CH), 108.8 (CH), 60.8 ($CH_3$), 57.9 ($CH_2$), 56.2 ($CH_2$), 39.6 (CH), 35.6 (CH), 28.4 ($CH_3$), 25.4 ($CH_2$), 24.2 ($CH_2$), 10.3 ($CH_2$), 8.4 ($CH_2$); HRMS (ESI-TOF) for $C_{30}H_{35}FN_4O_6S[M+H]^+$ : Calcd., 599.2339, Found, 599.2343.

**B. General procedure for *N*-boc deprotection of NTR-MXF derivatives (1a-1g)**

To a solution of compound in dry DCM (15 mL), TFA (26.47 mmol) was added dropwise at 0 ºC under nitrogen atmosphere. The resulting reaction mixture was stirred at RT (~ 4-6 h) until the starting material had been completely consumed (determined by TLC) and the solvent was concentrated *in vacuo* at ~5 ºC . The product was precipitated by diethyl ether on sonication and the ether solution was then carefully decanted. The obtained solid was washed twice with diethyl ether, and the ethereal solution was concentrated *in vacuo* to afford prodrugs **1a-1g** as pure products.

**6-(1-cyclopropyl-6-fluoro-8-methoxy-3-(((4-nitrobenzyl)oxy)carbonyl)-4-oxo-1,4-dihydroquinolin-7-yl)octahydro-1H-pyrrolo[3,4-b]pyridin-1-ium 2,2,2-trifluoroacetate**

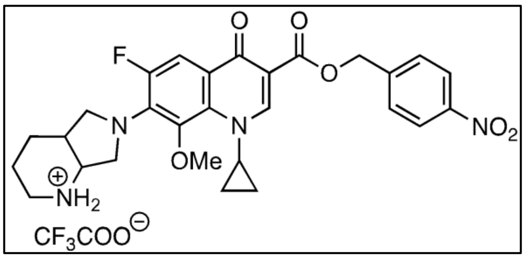


**(1a):** Starting from **3a** (0.100 g, 0.16 mmol), **1a** (0.090 g, 83%) was obtained as a white solid; mp = 217-218 ºC; FT-IR ($\nu_{max}$, cm$^{-1}$): 3394 (N-H stretch), 2924 (C-H stretch), 1691 (ester C=O stretch), 1616 (conjugated ketone C=O stretch); $^{19}$F NMR (376 MHz, DMSO-*d6*): δ -74.4, -122.8; $^{1}$H NMR (400 MHz, DMSO-*d6*) : δ 9.24 (d, *J* = 9.6 Hz, 1H), 8.59 (d, *J* = 10.1 Hz, 1H), 8.55 (s, 1H), 8.27 (d, *J* = 8.7 Hz, 2H), 7.79 (d, *J* = 8.7 Hz, 2H), 7.60 (d, *J* = 14.4 Hz, 1H), 5.47 – 5.39 (m, 2H), 4.03 – 4.00 (m, 2H), 3.90 (s, 1H), 3.79 (td, *J* = 9.8, 4.6 Hz, 1H), 3.70 (t, *J* = 7.7 Hz, 1H), 3.57 (s, 3H), 3.50 (d, *J* = 11.7 Hz, 1H), 3.24 (d, *J* = 9.4 Hz, 1H), 2.99 (q, *J* = 9.5, 8.0 Hz, 1H), 2.68 – 2.65 (m, 1H), 1.83 -1.70 (m, 4H), 1.18 – 1.10 (m, 1H), 1.07 – 0.94 (m, 2H), 0.85 – 0.78 (m, 1H); $^{13}$C NMR (100 MHz, DMSO-*d6*): δ 171.3, 164.4, 153.5, 151.2, 147.0, 144.7, 141.0 (d, *J* = 26.6 Hz), 134.9 (d, *J* = 41.3 Hz), 133.3, 128.1, 123.6, 120.9 (d, *J* = 29.4 Hz), 108.2, 107.3 (d, *J* = 94.1 Hz), 64.2, 61.5, 54.4, 54.1 (d, *J* = 31.2 Hz), 51.6 (d, *J* = 38.0 Hz), 41.9, 34.3, 20.7, 17.8, 9.5, 8.4; DEPT-135 NMR (100 MHz, DMSO-*d6*): δ 150.9 (CH), 127.8 (CH), 123.3 (CH), 107.0 (CH), 63.9 ($CH_2$), 61.3 ($CH_3$), 54.2 (CH), 54.0 ($CH_2$), 51.3 ($CH_2$), 41.6 ($CH_2$), 34.0 (CH), 20.4 ($CH_2$), 17.5 ($CH_2$), 9.2 ($CH_2$), 8.1 ($CH_2$); HRMS (ESI-TOF) for $C_{28}H_{30}FN_4O_6$ $[M]^+$ : Calcd., 537.2144, Found, 537.2146.

**6-(1-cyclopropyl-6-fluoro-8-methoxy-3-(((5-nitrofuran-2-yl)methoxy) carbonyl)-4-oxo-1,4-dihydroquinolin-7-yl)octahydro-1H-pyrrolo[3,4-b]pyridin-1-ium 2,2,2-trifluoro acetate (1b):** Starting from **3b** (0.100 mg, 0.15 mmol), **1b** (0.092 mg, 85%) was obtained as a

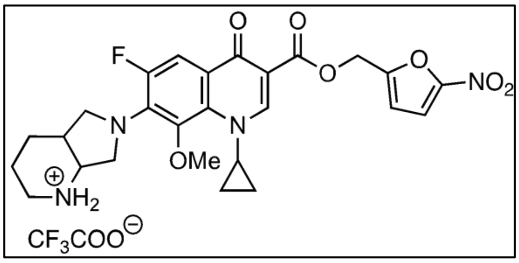


reddish yellow solid; mp = 198-199 ºC; FT-IR ($\nu_{max}$, $cm^{-1}$): 3427 (N-H stretch), 2971 (C-H stretch), 1690 (ester C=O stretch), 1660 (conjugated ketone C=O stretch), 1451 (asymmetric $NO_2$ stretch), 1361 (symmetric $NO_2$ stretch); $^{19}F$ NMR (376 MHz, DMSO-*d*$_6$): δ -74.6, -122.8; $^1H$ NMR (400 MHz, DMSO-*d*$_6$) : δ 9.21 (d, *J* = 11.4 Hz, 1H), 8.55 (d, *J* = 9.0 Hz, 1H), 8.51 (s, 1H), 7.71 (d, *J* = 3.7 Hz, 1H), 7.55 (d, *J* = 13.5 Hz, 1H), 6.99 (d, *J* = 3.7 Hz, 1H), 5.38 – 5.31 (m, 2H), 4.02 – 3.98 (m, 2H), 3.89 (s, 1H), 3.77 (td , *J* = 10.0, 4.5 Hz, 1H), 3.72 – 3.67 (m, 1H), 3.56 (s, 3H), 3.51 (d, *J* = 11.9 Hz, 1H), 3.25 – 3.22 (m, 1H), 2.98 (q, *J* = 9.6 Hz, 1H), 2.67 – 2.63 (m, 1H), 1.83 – 1.66 (m, 4H), 1.16 – 1.07 (m, 1H), 1.04 – 0.96 (m, 2H), 0.82 – 0.77 (m, 1H); $^{13}C$ NMR (100 MHz, DMSO-*d*$_6$): δ 171.3, 163.7, 158.4 (d, *J* = 140.6 Hz), 153.9, 153.5, 151.3, 151.1, 141.0 (d, *J* = 26.7 Hz), 135.0 (d, *J* = 41.6 Hz), 133.3, 120.8 (d, *J* = 29.4 Hz), 114.1, 113.9, 107.8, 107.2 (d, *J* = 95.1 Hz), 61.6, 57.2, 54.5, 54.2 (d, *J* = 12.4 Hz), 51.5 (d, *J* = 37.7 Hz), 41.9, 34.3, 20.7, 17.7, 9.5, 8.4; DEPT-135 NMR (100 MHz, DMSO-*d*$_6$): δ 151.0 (CH), 113.7 (CH), 113.5 (CH), 106.9 (CH), 61.3 ($CH_3$), 56.8 ($CH_2$), 54.1 (CH), 53.9 ($CH_2$), 51.2 ($CH_2$), 41.5 ($CH_2$), 34.0 (CH), 20.4 ($CH_2$), 17.4 ($CH_2$), 9.2 ($CH_2$), 8.1 ($CH_2$); HRMS (ESI-TOF) for $C_{28}H_{28}FN_4O_7$ $[M]^+$ : Calcd., 527.1937, Found, 527.1938.

**6-(1-cyclopropyl-6-fluoro-8-methoxy-3-(((5-nitrothiophen-2-yl)methoxy)carbonyl)-4-oxo -1,4-dihydroquinolin-7-yl)octahydro-1H-pyrrolo[3,4-b]pyridin-1-ium 2,2,2-trifluoroacetate (1c)**: Starting from **3c** (0.100 mg, 0.15 mmol), **1c** (0.094 mg, 85%) was

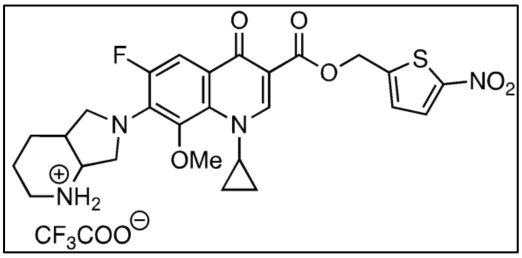


obtained as a yellow solid; mp = 211-212 ºC;FT-IR ($\nu_{max}$, $cm^{-1}$): 3425 (N-H stretch), 2999 (C-H stretch), 1670 (ester C=O stretch), 1659 (conjugated ketone C=O stretch), 1423 (asymmetric $NO_2$ stretch), 1314 (symmetric $NO_2$ stretch); $^{19}F$ NMR (376 MHz, DMSO-*d*$_6$): δ -74.8, -122.7; $^1H$ NMR (400 MHz, DMSO-*d*$_6$): δ 9.13 (s, 1H), 8.53 (s, 2H), 8.07 (d, *J* = 3.8 Hz, 1H), 7.59 (d, *J* = 14.4 Hz, 1H), 7.36 (d, *J* = 4.4 Hz, 1H), 5.53 – 5.46 (m, 2H), 4.06 – 3.98 (m, 2H), 3.88 (t, *J* = 5.4 Hz, 1H), 3.76 (td , *J* = 10.0, 4.5 Hz, 1H), 3.69 (dt , *J* = 8.2 Hz, 2.9 Hz, 1H), 3.57 (s, 3H), 3.50 (d, *J* = 12.0 Hz, 1H), 3.23 (d, *J* = 12.9 Hz, 1H), 2.99 (d, *J* = 11.4 Hz, 1H), 2.67 – 2.60 (m, 1H), 1.84 – 1.66 (m, 4H), 1.17 – 1.10 (m, 1H), 1.06 – 0.94 (m,

2H), 0.87 – 0.78 (m, 1H); $^{13}C$ NMR (100 MHz, DMSO-$d_6$): δ 171.2, 164.0, 153.5, 151.3, 151.0, 150.8, 148.1, 141.0 (d, $J$ = 26.7 Hz), 134.9 (d, $J$ = 41.9 Hz), 133.2, 129.7, 127.5, 120.9 (d, $J$ = 29.1 Hz), 107.6, 107.2 (d, $J$ = 95.3 Hz), 61.5, 60.3, 54.4, 54.1 (d, $J$ = 16.3 Hz), 51.5 (d, $J$ = 36.4 Hz), 41.8, 34.2, 20.6, 17.7, 9.4, 8.4; DEPT-135 NMR (100 MHz, DMSO-$d_6$): δ 151.1 (CH), 129.4 (CH), 127.3 (CH), 107.0 (CH), 61.3 ($CH_3$), 60.1 ($CH_2$), 54.2 (CH), 53.9 ($CH_2$), 51.2 ($CH_2$), 41.5 ($CH_2$), 34.0 (CH), 20.4 ($CH_2$), 17.4 ($CH_2$), 9.2 ($CH_2$), 8.1 ($CH_2$); HRMS (ESI-TOF) for $C_{26}H_{28}FN_4O_6S$ $[M]^+$ : Calcd., 543.1708, Found, 543.1716.

**6-(1-cyclopropyl-6-fluoro-8-methoxy-3-(((1-methyl-2-nitro-1*H*-imidazol-5-yl)methoxy) carbonyl)-4-oxo-1,4-dihydroquinolin-7-yl)octahydro-1*H*-pyrrolo[3,4-*b*]pyridin-1-ium 2,2,2-trifluoroacetate (1d):** Starting from **3d** (0.180 g, 0.28 mmol), **1d** (0.178 g, 97%) was

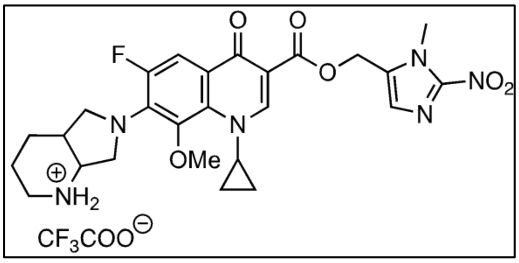


obtained as a white solid; mp = 189-190 ºC; FT-IR ($\nu_{max}$, $cm^{-1}$): 2963 (C-H stretch), 1718 (ester C=O stretch), 1688 (conjugated ketone C=O stretch), 1617 (N-H bend), 1539 (asymmetric $NO_2$ stretch), 1320 (symmetric $NO_2$ stretch); $^{19}F$ NMR (376 MHz, DMSO-$d_6$): δ -74.3, -122.8; $^{1}H$ NMR (400 MHz, DMSO-$d_6$): δ 9.23 (d, $J$ = 11.5 Hz, 1H), 8.60 – 8.55 (m, 1H), 8.48 (s, 1H), 7.56 (d, $J$ = 14.4 Hz, 1H), 7.33 (s, 1H), 5.38 (s, 2H), 4.04 (s, 3H), 4.02 - 3.97 (m, 2H), 3.90 (s, 1H), 3.76 (td, $J$ = 10.1, 4.8 Hz, 1H), 3.72 – 3.67 (m, 1H), 3.56 (s, 3H), 3.50 (d, $J$ = 10.1 Hz, 1H), 3.23 (d, J= 13.4 Hz, 1H), 2.98 (d, $J$ = 10.6 Hz, 1H), 2.68 – 2.65 (m, 1H), 1.86 – 1.72 (m, 4H), 1.16 – 1.07 (m, 1H), 1.05 – 0.94 (m, 2H), 0.81 – 0.76 (m, 1H); $^{13}C$ NMR (100 MHz, DMSO-$d_6$): δ 171.3, 163.9, 158.5, 158.2, 153.5, 151.2, 151.0, 146.1, 141.0 (d, $J$ = 26.2 Hz), 134.9 (d, $J$ = 42.2 Hz), 133.4, 133.3, 128.6, 120.8 (d, $J$ = 29.1 Hz), 108.0, 107.2 (d, $J$ = 93.2 Hz), 61.5, 55.1, 54.4, 54.1 (d, $J$ = 14.1 Hz), 51.4 (d, $J$ = 36.4 Hz), 41.8, 34.4, 34.2, 20.6, 17.7, 9.4, 8.4; DEPT-135 NMR (100 MHz, DMSO-$d_6$): δ 150.9 (CH), 128.4 (CH), 107.0 (CH), 61.3 ($CH_3$), 54.8 ($CH_2$), 54.2 (CH), 53.9 ($CH_2$), 51.3 ($CH_2$), 41.6 ($CH_2$), 34.1 (CH), 34.0 (CH), 20.4 ($CH_2$), 17.4 ($CH_2$), 9.2 ($CH_2$), 8.1 ($CH_2$); HRMS (ESI-TOF) for $C_{26}H_{30}FN_6O_6$ $[M]^+$ : Calcd., 541.2205, Found, 541.2209.

**6-(1-cyclopropyl-6-fluoro-8-methoxy-3-(((2-nitrothiazol-5-yl)methoxy) carbonyl)-4-oxo-1,4-dihydroquinolin-7-yl)octahydro-1*H*-pyrrolo[3,4-*b*]pyridin-1-ium 2,2,2-trifluoroacetate (1e):** Starting from **3e** (0.180 g, 0.27 mmol), **1e** (0.178 g, 97%) was obtained as orange yellow solid; mp = 207-208 ºC; FT-IR ($\nu_{max}$, $cm^{-1}$): 2924 (C-H stretch), 1716 (ester C=O stretch), 1680 (conjugated ketone C=O stretch), 1617 (N-H bend), 1543 (asymmetric $NO_2$

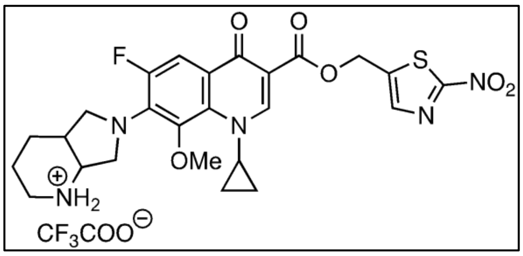

stretch), 1322 (symmetric $NO_2$ stretch);$^{19}$F NMR (376 MHz, DMSO-$d_6$): δ -74.6, -122.7; $^1$H NMR (400 MHz, DMSO-$d_6$): δ 9.25 (d, $J$ = 9.7 Hz, 1H), 8.58 (d, $J$ = 9.1 Hz, 1H), 8.53 (s, 2H), 8.16 (s, 1H), 7.57 (d, $J$ = 14.2 Hz, 1H), 5.56 (s, 2H), 4.03 – 3.98 (m, 2H), 3.89 (t, $J$ = 5.7 Hz, 1H), 3.77 (td, $J$ = 10.0, 4.6 Hz, 1H), 3.72 – 3.67 (m, 1H), 3.56 (s, 3H), 3.50 (d, $J$ = 12.2 Hz, 1H), 3.23 (d, $J$ = 12.8 Hz, 1H), 2.98 (t, $J$ = 10.6 Hz, 1H), 2.68 – 2.64 (m, 1H), 1.83 – 1.70 (m, 4H), 1.15 – 1.10 (m, 1H), 1.04 – 0.96 (m, 2H), 0.83 – 0.79 (m, 1H); $^{13}$C NMR (100 MHz, DMSO-$d_6$): δ 171.2, 166.3, 164.2, 153.5, 151.4, 151.0, 142.8, 141.9, 141.0 (d, $J$ = 26.5 Hz), 135.0 (d, $J$ = 41.8 Hz), 120.8 (d, $J$ = 28.3 Hz), 107.5, 107.2 (d, $J$ = 94.7 Hz), 61.5, 57.9, 54.4, 54.0 (d, $J$ = 13.6 Hz), 51.5 (d, $J$ = 36.8 Hz), 41.8, 34.2, 20.6, 17.7, 9.5, 8.4; DEPT-135 NMR (100 MHz, DMSO-$d_6$): δ 151.1 (CH), 141.6 (CH), 107.0 (CH), 61.3 ($CH_3$), 57.7 ($CH_2$), 54.2 (CH), 53.9 ($CH_2$), 51.2 ($CH_2$), 41.5 ($CH_2$), 34.0 (CH), 20.4 ($CH_2$), 17.4 ($CH_2$), 9.2 ($CH_2$), 8.1 ($CH_2$); HRMS (ESI-TOF) for $C_{25}H_{27}FN_5O_6S[M]^+$ : Calcd., 544.1661, Found, 544.1669.

**6-(3-((benzyloxy)carbonyl)-1-cyclopropyl-6-fluoro-8-methoxy-4-oxo-1,4-dihydroquinolin-7-yl)octahydro-1H-pyrrolo[3,4-b]pyridin-1-ium 2,2,2-trifluoroacetate**

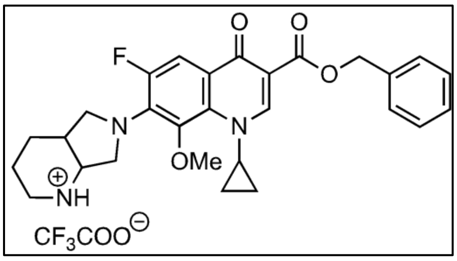

**(1f):** Starting from **3f** (0.100 g, 0.17 mmol), **1f** (0.090 g, 88%) was isolated as a yellow solid; mp = 147-148 ºC; FT-IR ($\nu_{max}$, $cm^{-1}$): 3522 (N-H stretch), 2981 (C-H stretch), 1709 (ester C=O stretch), 1610 (conjugated ketone C=O stretch); $^{19}$F NMR (376 MHz, DMSO-$d_6$): δ -74.2, -122.9; $^1$H NMR (400 MHz, DMSO-$d_6$): δ 9.16 (d, $J$ = 9.2 Hz, 1H), 8.55 (d, $J$ = 11.2 Hz, 1H), 8. 50 (s, 1H), 7.58 (d, $J$ = 14.3 Hz, 1H), 7.50 – 7.47 (m, 2H), 7.42 – 7.37 (m, 2H), 7.35 – 7.31 (m, 1H), 5.31 – 5.24 (m, 2H), 4.02 – 3.97 (m, 2H), 3.90 (br, 1H), 3.77 (td, $J$ = 10.0, 4.4 Hz, 1H), 3.72 – 3.67 (m, 1H), 3.56 (s, 3H), 3.50 (d, $J$ = 11.3 Hz, 1H), 3.23 (d, $J$ = 12.4 Hz, 1H), 2.98 (d, $J$ = 10.9 Hz, 1H), 2.68 – 2.62 (m, 1H), 1.84 – 1.66 (m, 4H), 1.16 – 1.07 (m, 1H), 1.04 – 0.94 (m, 2H), 0.84 – 0.76 (m, 1H); $^{13}$C NMR (100 MHz, DMSO-$d_6$): δ 171.3, 164.3, 151.0, 150. 9, 141.0 (d, $J$ = 26.0 Hz), 136.7, 134.8 (d, $J$ = 42.2 Hz), 133.3, 128.4, 127.8, 127.6, 120.9 (d, $J$ = 29.7 Hz), 108.5, 107.2 (d, $J$ = 95.5 Hz), 65.2, 61.5, 54.4, 54.1 (d, $J$ = 12.2 Hz), 51.4 (d, $J$ = 35.8 Hz), 41.8, 40.2, 34.2, 20.6, 17.7, 9.4, 8.4; DEPT-135 NMR (100 MHz, DMSO-$d_6$): δ 150.6 (CH), 128.1 (CH), 127.5 (CH), 127.4 (CH), 106.9 (CH), 64.9 ($CH_2$), 61.2 ($CH_3$), 54.1

(CH), 53.8 ($CH_2$), 51.1 ($CH_2$), 41.5 ($CH_2$), 33.9 (CH), 20.4 ($CH_2$), 17.4 ($CH_2$), 9.2 ($CH_2$), 8.1 ($CH_2$); HRMS (ESI-TOF) for $C_{28}H_{31}FN_3O_4$ $[M]^+$ : Calcd., 492.2293, Found, 492.2297.

**6-(1-cyclopropyl-6-fluoro-8-methoxy-4-oxo-3-((thiazol-5-ylmethoxy)carbonyl)-1,4-dihydroquinolin-7-yl)octahydro-1*H*-pyrrolo[3,4-*b*]pyridin-1-ium 2,2,2-trifluoroacetate**

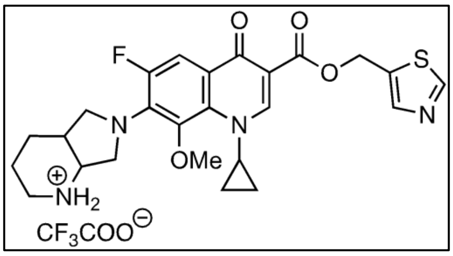


**(1g):** Starting from **3g** (0.100 g, 0.16 mmol), **1g** (0.090 g, 88%) was isolated as a yellow solid; mp = 188-189 ºC; FT-IR ($\nu_{max}$, $cm^{-1}$): 2952 (C-H stretch), 1716 (ester C=O stretch), 1686 (conjugated ketone C=O stretch), 1617 (N-H stretch); $^{19}$F NMR (376 MHz, DMSO-$d_6$): δ -74.3, -122.8; $^1$H NMR (400 MHz, DMSO-$d_6$): δ 9.18 – 9.12 (m, 2H), 8.53 – 8.48 (m, 2H), 8.02 (s, 1H), 7.54 (d, $J$ = 14.3 Hz, 1H), 5.49 (s, 2H), 4.01 – 3.97 (m, 2H), 3.89 (br, 1H), 3.76 (td, $J$ = 9.9, 4.3 Hz, 1H), 3.71 – 3.66 (m, 1H), 3.55 (s, 3H), 3.49 (d, $J$ = 11.2 Hz, 1H), 3.23 (d, $J$ = 14.3 Hz, 1H), 2.98 (q, $J$ = 10.8 Hz, 1H), 2.67 – 2.64 (m, 1H), 1.84 – 1.66 (m, 4H), 1.15 – 1.07 (m, 1H), 1.03 – 0.93 (m, 2H), 0.83 – 0.75 (m, 1H); $^{13}$C NMR (100 MHz, DMSO-$d_6$): δ 171.2, 164.0, 158.4, 158.1, 155.9, 153.4, 151.1, 151.0, 143.7, 141.0 (d, $J$ = 25.8 Hz), 135.0 (d, $J$ = 41.8 Hz), 133.3, 120.8 (d, $J$ = 30.1 Hz), 115.7, 108.0, 107.2 (d, $J$ = 94.3 Hz), 61.5, 57.5, 54.4, 54.1 (d, $J$ = 16.0 Hz), 51.4 (d, $J$ = 38.4 Hz), 41.8, 34.2, 20.6, 17.7, 9.4, 8.4; DEPT-135 NMR (100 MHz, DMSO-$d_6$): δ 150.8 (CH), 143.4 (CH), 106.9 (CH), 61.3 ($CH_3$), 57.2 ($CH_2$), 54.2 (CH), 53.9 ($CH_2$), 51.2 ($CH_2$), 41.6 ($CH_2$), 34.0 (CH), 20.4 ($CH_2$), 17.4 ($CH_2$), 9.2 ($CH_2$), 8.1 ($CH_2$); HRMS (ESI-TOF) for $C_{25}H_{28}FN_4O_4S$$[M]^+$ : Calcd., 499.1810, Found, 499.1816.

**Preparation of carbamates of MXF (2a-2b):** The reaction of 4-nitrophenyl carbonates **5a** and **5b** with MXF in the presence of triethylamine provided **2a** and **2b** in good yields**.**

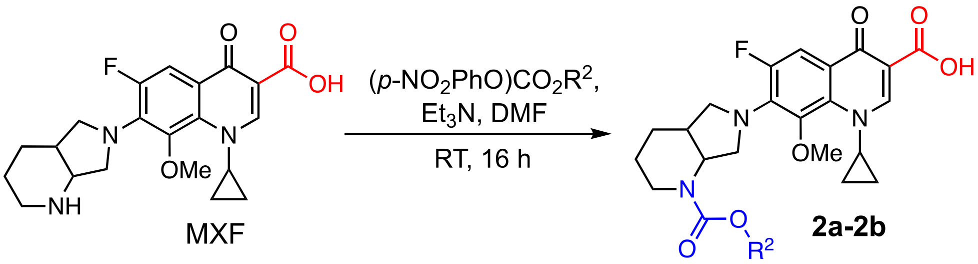


**Scheme S2.** Synthesis of **2a-2b**

#### C. General procedure for synthesis of nitroheterocyclic carbamates of MXF (2a-2b)

To a dried two-neck round bottom flask charged with a stir bar and carbonate derivative **5a** or **5b** (0.383 mmol) in anhydrous DMF (2 mL), $Et_3N$ (0.106 mL, 0.766 mmol) was added. This was followed by dropwise addition of a solution of MXF (0.200 g, 0.498 mmol) in anhydrous DMF (2 mL). The reaction mixture was left to stir at RT overnight under a positive pressure of nitrogen. After completion of the reaction (determined by TLC), the crude product was adsorbed onto celite. The resulting filtrate was diluted with water (10 mL), and the aqueous layer was extracted with EtOAc (3 × 10 mL). The organic extracts were combined, washed with brine (2 × 10 mL), dried over anhydrous $Na_2SO_4$, filtered and filtrate was concentrated under reduced pressure. The residue was subjected to purification by column chromatography over silica gel (60-120) with 5-10% MeOH/$CHCl_3$ to provide MXF carbamates **2a-2b** as pure products.

**1-cyclopropyl-6-fluoro-8-methoxy-7-(1-(((4-nitrobenzyl)oxy)carbonyl)octahydro-6*H*-pyrrolo[3,4-*b*]pyridin-6-yl)-4-oxo-1,4-dihydroquinoline-3-carboxylic acid (2a):** Starting

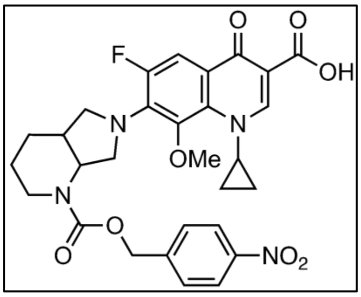


from **5a** (0.122 g, 0.383 mmol) and MXF, **2a** (0.126 g, 56%) was isolated as a yellow solid; mp = 183-184 ºC; $R_f$ (EtOAc:hexane = 50:50) = 0.5; FT-IR ($\nu_{max}$, $cm^{-1}$): 3502 (N-H stretch), 2930 (C-H stretch), 1727 (ester C=O stretch), 1701 (conjugated ketone C=O stretch), 1619 (N-H stretch), 1442 (asymmetric $NO_2$ stretch), 1319 (symmetric $NO_2$ stretch); $^{19}$F NMR (376 MHz, $CDCl_3$): δ -121.0; $^{1}$H NMR (400 MHz, $CDCl_3$): δ 14.94 (s, 1H), 8.78 (s, 1H), 8.23 (d, $J$ = 6.7 Hz, 2H), 7.80 (d, $J$ = 13.8 Hz, 1H), 7.53 (d, $J$ = 8.8 Hz, 2H), 5.27 (s, 2H), 4.88 (s, 1H), 4.20 – 4.09 (m, 2H), 4.02 – 3.96 (m, 1H), 3.93 (td, $J$ = 10.1, 2.6 Hz, 1H), 3.58 (s, 3H), 3.29 (s, 1H), 3.02 (s, 1H), 2.38 – 2.30 (m, 1H), 1.89 – 1.82 (m, 2H), 1.60 – 1.50 (m, 2H), 1.35 – 1.27 (m, 1H), 1.17 – 1.05 (m, 2H), 0.90 – 0.79 (m, 2H); $^{13}$C (100 MHz, $CDCl_3$): δ 176.9, 167.1, 155.5, 149.9, 147.8, 144.0, 134.5, 128.4, 124.0, 119.1 (d, $J$ = 35.0 Hz), 108.2 (d, $J$ = 94.7 Hz), 107.9, 66.1, 61.3, 56.5 (d, $J$ = 30.6 Hz), 52.9, 40.5, 39.9, 35.7, 29.8, 25.3, 10.7, 8.6; DEPT-135 NMR (100 MHz, $CDCl_3$): δ 149.8 (CH), 128.3 (CH), 123.9 (CH), 108.1 (CH), 66.0 ($CH_2$), 61.2 ($CH_3$), 56.5 ($CH_2$), 52.8 (CH), 40.4 (CH), 39.8 ($CH_2$), 35.6 (CH), 25.1 ($CH_2$), 10.6 ($CH_2$), 8.5 ($CH_2$); HRMS (ESI-TOF) for $C_{29}H_{29}FN_4O_8$ $[M+H]^+$ : Calcd., 581.2047, Found, 581.2051.

**1-cyclopropyl-6-fluoro-8-methoxy-7-(1-(((2-nitrothiazol-5-yl) methoxy) carbonyl) octahydro-6*H*-pyrrolo[3,4-*b*]pyridin-6-yl)-4-oxo-1,4-dihydroquinoline-3-carboxylic acid (2b):**Starting from **5b** (0.124 g, 0.383 mmol) and MXF, **2b** (0.140 g, 62%) was isolated as a

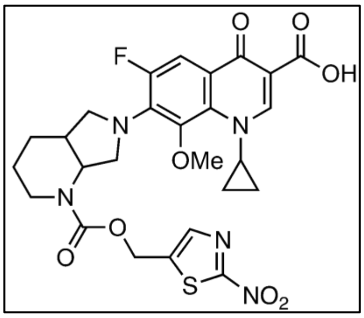


yellow solid; mp = 131-132 ºC; $R_f$ (EtOAc:hexane = 50:50) = 0.35; FT-IR ($\nu_{max}$, $cm^{-1}$): 3647 (N-H stretch), 2933 (C-H stretch), 1725 (ester C=O stretch), 1701 (conjugated ketone C=O stretch), 1619 (N-H stretch), 1439 (asymmetric $NO_2$ stretch), 1319 (symmetric $NO_2$ stretch); $^{19}F$ NMR (376 MHz, $CDCl_3$): δ -121.1; $^{1}H$ NMR (400 MHz, $CDCl_3$): δ 14.93 (s, 1H), 8.78 (s, 1H), 7.84 (s, 1H), 7.79 (d, *J* = 13.8 Hz, 1H), 5.35 (d, *J* = 4.7 Hz, 2H), 4.87 – 4.84 (m, 1H), 4.14 – 4.09 (m, 1H), 4.02 – 3.96 (m, 1H), 3.91 (td, *J* = 10.1, 2.6 Hz, 1H), 3.59 (s, 3H), 3.28 (d, *J* = 7.9 Hz, 1H), 3.01 (s, 1H), 2.36 – 2.30 (m, 1H), 1.86 – 1.82 (m, 2H), 1.58 – 1.49 (m, 2H), 1.35 – 1.27 (m, 1H), 1.18 – 1.03 (m, 3H), 0.90 – 0.80 (m, 2H); $^{13}C$ (100 MHz, $CDCl_3$): δ 176.9, 167.1, 155.2, 149.9, 142.2, 141.9, 134.5, 119.1 (d, *J* = 35.0 Hz), 108.3 (d, *J* = 94.7 Hz), 107.8, 61.3, 59.1, 56.5 (d, *J* = 30.6 Hz), 52.9, 40.5, 40.0, 35.6, 29.8, 25.1, 10.7, 8.6; DEPT-135 NMR (100 MHz, $CDCl_3$): δ 149.9 (CH), 141.9 (CH), 108.2 (CH), 61.3 ($CH_3$), 58.7 ($CH_2$), 56.5 ($CH_2$), 52.9 (CH), 40.5 (CH), 40.0 ($CH_2$), 35.6 (CH), 25.1 ($CH_2$), 24.1 ($CH_2$), 10.7 ($CH_2$); HRMS (ESI-TOF) for $C_{26}H_{26}FN_5O_8S[M+H]^+$ : Calcd., 588.1564, Found, 588.1572.

**D. Synthesis of carbonates**

**4-nitrobenzyl (4-nitrophenyl) carbonate (5a):**[6] A solution of 4-nitrophenyl chloroformate (0.987 g, 4.90 mmol) in anhydrous DCM (3 mL) was slowly added after 10 min to a stirred solution of **1a** (0.5 g, 3.27 mmol) and $Et_3N$ (0.911 mL, 6.53 mmol) in anhydrous DCM (6 mL). The reaction mixture was stirred under nitrogen atmosphere at RT for 16 h and then partitioned between DCM (20 mL) and water (20 mL). The organic layer was washed with water (30 mL) and brine (30 mL). The combined organic extracts were dried and the solvent was evaporated. The residue was purified by silica gel (60-120 mesh) column chromatography using 12% EtOAc/hexane to obtain **5a** (0.702 g, 68%) as a white solid. The analytical data is consistent with the reported literature.

**4-nitrophenyl ((2-nitrothiazol-5-yl)methyl) carbonate (5b):** A solution of 4-nitrophenyl chloroformate (1.36 g, 6.75 mmol) in anhydrous THF (3 mL) was slowly added to a stirred solution of **6e** (0.901 g, 5.63 mmol) and pyridine (0.543 mL, 6.75 mmol) in anhydrous THF (6 mL) after 20 min at room temperature. The reaction mixture was allowed to stir under nitrogen

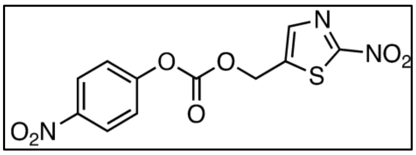


atmosphere at RT for 12 h and then partitioned between EtOAc (20 mL) and water (20 mL). The organic layer was washed with water (30 mL) and brine (30 mL). The combined extracts were dried and the solvent was evaporated. The residue was purified by silica gel (60-120 mesh) using 10% EtOAc/hexane to obtain **5b** (0.28 g, 16%) as yellow solid. $R_f$ (EtOAc:hexane = 20:80) = 0.5; $^1$H NMR (400 MHz, $CDCl_3$): δ 8.32 – 8.29 (m, 2H), 7.96 (s, 1H), 7.42 – 7.38 (m, 2H), 5.51 (s, 2H). HRMS (ESI-TOF) for $C_{11}H_7N_3O_7S[M+H]^+$ : Calcd., 326.0083, Found, 326.0078.

#### E. Synthesis of heterocyclic alcohols (6b-6e)

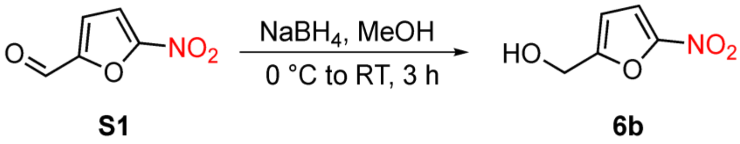


**Scheme S3.** Synthesis of (5-nitrofuran-2-yl)methanol **(6b)**

**(5-nitrofuran-2-yl)methanol (6b):**[2] $NaBH_4$ (0.294 g, 7.80 mmol) was slowly added to a vigorously stirred one-neck round bottom flask charged with **S1** (1 g, 7.09 mmol) in dry MeOH (15 mL) under nitrogen atmosphere at 0 °C. The reaction mixture was allowed to stir at RT for 3 h. After completion of the reaction (as monitored by TLC), ice cold water (20 mL) was added and the solution was acidified to pH 7.0 with 3 M HCl. The reaction mixture was extracted with EtOAc water (3 × 25 mL) and the combined organic extracts were dried over anhydrous $Na_2SO_4$, filtered and the filtrate was concentrated *in vacuo*. The residue was purified by silica gel (60-120 mesh) column chromatography using 20-25% EtOAc/hexane as an eluant to afford **6b** (0.890 g, 89%)as a yellow oil. All analytical data are consistent with reported values.

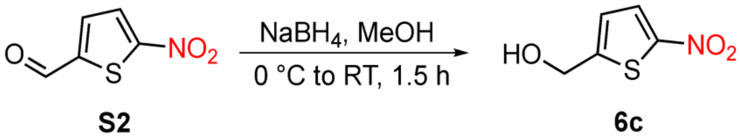


**Scheme S4.** Synthesis of (5-nitrothiophen-2-yl)methanol **(6c)**

**(5-nitrothiophen-2-yl)methanol (6c):**[2] To a stirred solution of **S2** (1 g, 6.36 mmol) in dry MeOH (15 mL), $NaBH_4$ (0.265 g, 7.00 mmol) was added at 0 °C and the reaction mixture was allowed to stir at RT for 1.5 h. After completion of the reaction (as monitored by TLC), ice cold water (20 mL) was added and the solution was acidified to pH 7.0 with 3 M HCl. The reaction mixture was extracted with EtOAc water (3 × 25 mL) and the combined organic extracts were dried over anhydrous $Na_2SO_4$, filtered and the filtrate was concentrated *in vacuo*. The residue was purified by silica gel (60-120 mesh) column chromatography using 25-30% EtOAc/hexane as an eluant to afford **6c** (0.920 g, 91%)as a brown oil. All analytical data are consistent with reported values.

**(1-methyl-2-nitro-1*H*-imidazol-5-yl)methanol (6d):**[3] The synthesis of **6d** commenced with compound **7**, that was generated through a Marckwald cyclisation of **S3** and KSCN. Following a modified reported protocol, $NaNO_2$-mediated desulfurization of **7** under acidic conditions gave **8**. Selective protection of **8** with TBDMSCl provided the silyl ether **9**. This intermediate was reacted with $TsN_3$, then deprotected with HCl/MeOH, and finally hydrogenatedby Pd/C to afford **12**. The diazotization of **12** produced **6d** in good yield (**Scheme S5**). All analytical data of **6d**, **7-12** are consistent with reported values.

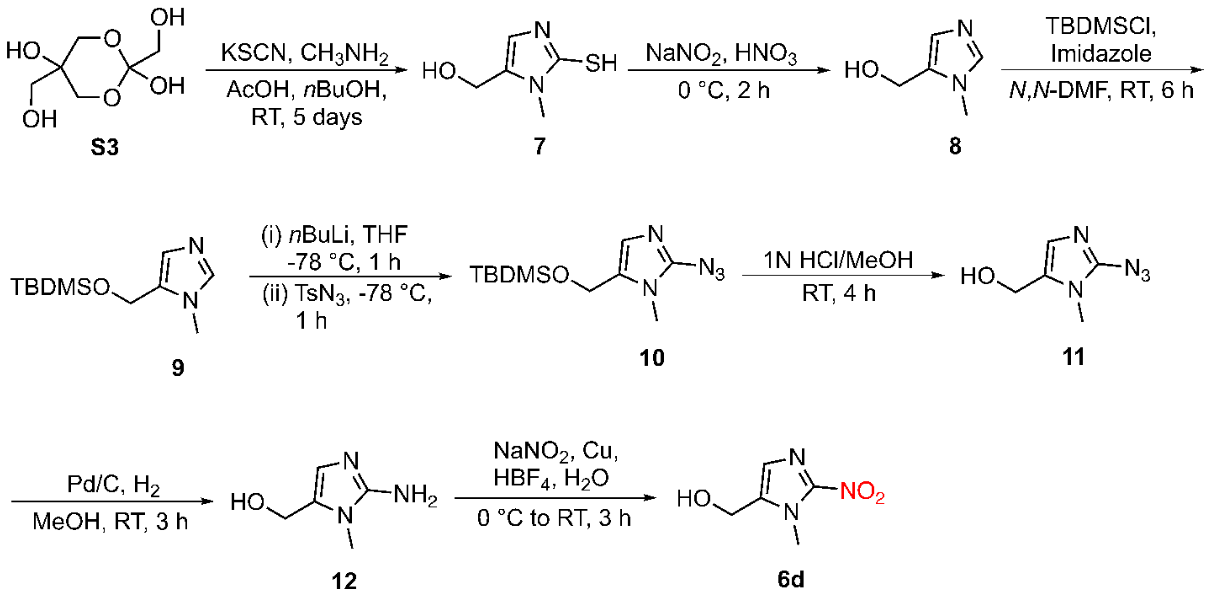


**Scheme S5.** Synthesis of (1-methyl-2-nitro-1*H*-imidazol-5-yl)methanol **(6d)**

**(2-mercapto-1-methyl-1*H*-imidazol-5-yl)methanol (7):**[4,5] To a mixture of *n*BuOH (200 mL) and AcOH (30 mL), dihydroxyacetone dimer (**S3**) (25 g, 1.38 mol), KSCN (40.4 g, 4.16 mol) and methylamine hydrochloride (10.7 g, 3.46 mol) were added and stirred at 25 ºC. After 5 days, water (50 mL) was added and the solid was isolated by filtration. The solid residue was washed with water (125 mL) and $Et_2O$ (125 mL) and dried in vacuum to afford compound **7** (11.9 g, 59%)as a white powder.

**(1-methyl-1*H*-imidazol-5-yl)methanol (8):**[4,5] The intermediate **7** (8.16 g, 56.62 mmol) was added in small portions to a vigorously stirred solution of $NaNO_2$ (0.160 g, 2.32 mmol) in aq. $HNO_3$ (20 mL, 2.4 M) at 0 ºC over a period of 1 h. The resulting yellow solution was stirred at RT for 2 h. The reaction mixture was adjusted to pH 9.0 by adding solid $NaHCO_3$ and evaporated to dryness in vacuum. The residue was extracted with hot $CHCl_3$ (3 × 25 mL), dried over anhydrous $Na_2SO_4$, filtered and the filtrate was concentrated *in vacuo* to give compound **8** (1.30 g, 20%)as a pale yellow solid.

**5-(((*tert*-butyldimethylsilyl)oxy)methyl)-1-methyl-1*H*-imidazole (9):**[6] To a solution of compound **8** (1.32 g, 11.77 mmol) in *N,N*-DMF (10 mL), imidazole (2.40 g, 35.32 mmol) and TBDMSCl (1.95 g, 12.95 mmol) was added. The reaction was stirred at RT for 6 h under nitrogen atmosphere, and the resulting mixture was poured into water (20 mL) and extracted with DCM (3 × 20 mL). The organic layer was dried over anhydrous $Na_2SO_4$, concentrated in *vacuo,* and purified by silica gel (60-120 mesh) using 1% EtOAc/hexane as an eluant to yield **9** (0.710 g, 60%)as a yellow solid.

**2-azido-5-(((*tert*-butyldimethylsilyl)oxy)methyl)-1-methyl-1*H*-imidazole (10):**[3] *n*BuLi (1.5 mL, 3.75 mmol, 2.4 N in hexane) was added dropwise at -78 ºC to a solution of compound **9** (1 g, 4.42 mmol) in anhydrous THF (10 mL). $TsN_3$ (0.609 mL, 3.98 mmol) was then added dropwise and the reaction was continued stirring at -78 ºC for 1 h. The reaction mixture was quenched with aq. $NH_4Cl$ (15 mL) and extracted with EtOAc water (3 × 20 mL) and the combined organic extracts were washed with brine, dried over anhydrous $Na_2SO_4$, filtered and the filtrate was concentrated *in vacuo* to afford the crude product. The residue was purified by column chromatography on silica gel (60-120 mesh) using 2% EtOAc/hexane as an eluant to give **10** (0.710 g, 60%)as yellow oil.

**(2-azido-1-methyl-1*H*-imidazol-5-yl)methanol (11):**[3] The intermediate **10** (0.5 g, 1.87 mmol) was stirred in 1 N HCl/MeOH (5 mL) at RT for 4 h. The pH of the reaction was adjusted to 7.0 by adding a few drops of $Et_3N$. The reaction mixture was concentrated and then the crude product was washed with $Et_2O$ (5 mL) to afford compound **11** (0.225 g, 78%) as a yellow solid.

**(2-amino-1-methyl-1*H*-imidazol-5-yl)methanol (12):**[3] The compound **11** (0.220 g, 1.44 mmol) was dissolved in MeOH (5 mL). Pd/C (10%, 22 mg) was then added and the mixture was hydrogenated under $H_2$ balloon at RT for 3 h. After reaction, the catalyst was removed by filtration and the filtrate was concentrated *in vacuo* to provide compound **12** (0.162 g, 89%) as a gray solid.

**(1-methyl-2-nitro-1*H*-imidazol-5-yl)methanol (6d):**[3] Compound **12** (0.160 g, 1.26 mmol) was dissolved in fluoroboric acid (48% w/v, 0.986 mL) and the resulting solution was cooled to -15 ºC, followed by dropwise addition of $NaNO_2$ (0.130 g, 1.89 mmol) in water (1 mL). The solution was stirred at -15ºC for 30 min, then added dropwise into a solution of Cu powder (80 mg, 1.26 mmol) and sodium nitrite (1.17 g, 16.9 mmol) in water (2 mL).The reaction mixture was stirred at RT for 1 h, filtered to remove Cu and then the filtrate was extracted with EtOAc (3 × 10 mL). The combined organic extracts were dried over anhydrous $Na_2SO_4$, filtered and filtrate was evaporated in vacuum. The residue was purified by silica gel (60-120 mesh) column chromatography using 5% MeOH/$CHCl_3$ as an eluant to afford **6d** (0.123 g, 62%) as a gray solid.

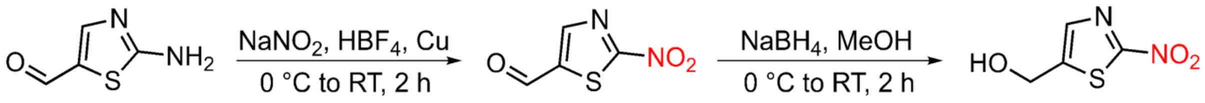


**Scheme S6.** Synthesis of (2-nitrothiazol-5-yl)methanol **(6e)**

**(2-nitrothiazol-5-yl)methanol (6e):**[7,8] To a solution of **S4** in fluoroboric acid (48% w/v, 6.12 mL), $NaNO_2$ (0.538 g, 7.80 mmol) in water (10 mL) was added dropwise at 0 ºC for 1 h. The resulting solution was then added dropwise into a mixture of Cu powder (1.61 g, 25.3 mmol) and 30 % w/v solution of sodium nitrite in water (10 mL). The reaction mixture was stirred at RT for 1 h, filtered to remove Cu, and the filtrate was extracted with EtOAc (3 × 20 mL). The combined organic extracts were dried over anhydrous $Na_2SO_4$ and the filtrate was evaporated in vacuum. The residue was purified by silica gel (60-120 mesh) column chromatography using 10-12% EtOAc/hexane as an eluant to afford **13** (0.990 g, 80%)as a dark reddish solid. This intermediate (0.5 g, 3.16 mmol) was dissolved in dry MeOH (10 mL). $NaBH_4$ (0.143 g, 3.79 mmol) was then added at 0 ºC and the reaction mixture was allowed to stir at RT for 1.5 h. After completion of the reaction (as monitored by TLC), ice cold water (10 mL) was added and the solution was acidified to pH 7.0 with 3 M HCl. The reaction mixture was extracted with EtOAc water (3 × 10 mL) and the combined organic extracts were dried over anhydrous $Na_2SO_4$, filtered and the filtrate was concentrated *in vacuo*. The residue was purified by silica gel (60-120 mesh) column chromatography using 30-32% EtOAc/hexane as an eluant to give **6e** (0.435 g, 85%) as a dark reddish solid. All analytical data are consistent with reported values.

#### 2. Purity of prodrugs by HPLC

Stock solutions (1 mM) of **1a-1e**, **2a** and **2b** were independently prepared in DMSO. A solution of 25 μM of respective prodrug (5 μL, 1 mM stock) was added to 195 μL of ACN and then injected (25 μL) in an HPLC instrument attached with a diode-array detector (detection wavelength 280 nm). The stationary phase used was C-18 reversed phase column (4.6 mm × 100 mm, 5 μm). The mobile phase used was 0.1 % HCOOH in $H_2O$:ACN at a flow rate of 0.5 mL/min with a run time of 18 min starting with 40:60 → 0-4 min, 20:80 → 4-12 min, 40:60 → 12-15 min, 40:60 → 15-18 min. The purity of the compounds were found to be >95% pure and HPLC traces are shown below.

(1) HPLC trace of **1a**

**1a,** $R_t$ = 3.03 min

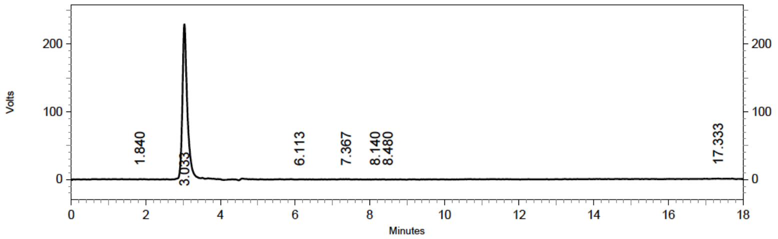


**DAD: Signal A,**
**280 nm/Bw:4 nm**
**Results**

| Retention Time | Area | Area % | Height | Height % |
|---|---|---|---|---|
| 1.840 | 21342 | 0.46 | 1156 | 0.24 |
| 3.033 | 4382848 | 94.90 | 479452 | 98.53 |
| 6.113 | 9281 | 0.20 | 924 | 0.19 |
| 7.367 | 64021 | 1.39 | 1635 | 0.34 |
| 8.140 | 21134 | 0.46 | 1017 | 0.21 |
| 8.480 | 18011 | 0.39 | 596 | 0.12 |
| 17.333 | 101557 | 2.20 | 1809 | 0.37 |
| Totals | 4618194 | 100.00 | 486589 | 100.00 |

#### (2) HPLC trace of **1b**

**1b,** $R_t$ = 3.01 min

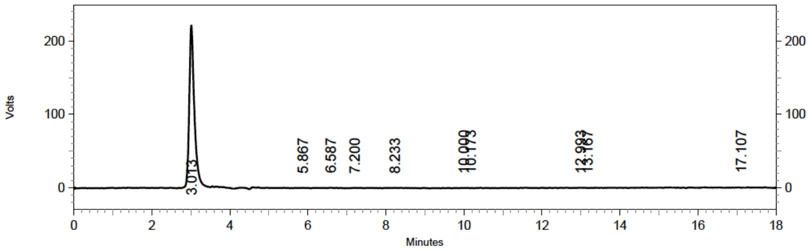


**DAD: Signal A,**
**280 nm/Bw:4 nm**
**Results**

| Retention Time | Area | Area % | Height | Height % |
|---|---|---|---|---|
| 3.013 | 4249534 | 95.33 | 464587 | 98.29 |
| 5.867 | 18237 | 0.41 | 1100 | 0.23 |
| 6.587 | 47590 | 1.07 | 1756 | 0.37 |
| 7.200 | 48370 | 1.09 | 1342 | 0.28 |
| 8.233 | 60716 | 1.36 | 1430 | 0.30 |
| 10.000 | 14193 | 0.32 | 910 | 0.19 |
| 10.173 | 52 | 0.00 | 0 | 0.00 |
| 12.993 | 10661 | 0.24 | 642 | 0.14 |
| 13.167 | 24 | 0.00 | 0 | 0.00 |
| 17.107 | 8411 | 0.19 | 917 | 0.19 |
| Totals | 4457788 | 100.00 | 472684 | 100.00 |

#### (3) HPLC trace of **1c**

**1c,** $R_t$ = 3.03 min

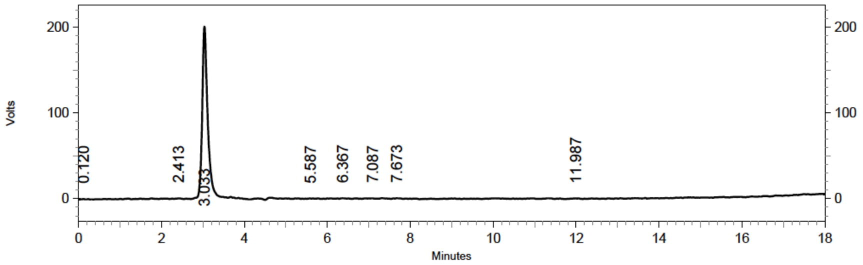


**DAD: Signal A,**
**280 nm/Bw:4 nm**
**Results**

| Retention Time | Area | Area % | Height | Height % |
|---|---|---|---|---|
| 0.120 | 8733 | 0.21 | 968 | 0.22 |
| 2.413 | 16169 | 0.40 | 1723 | 0.40 |
| 3.033 | 3864401 | 94.54 | 419994 | 97.47 |
| 5.587 | 12252 | 0.30 | 934 | 0.22 |
| 6.367 | 38289 | 0.94 | 1776 | 0.41 |
| 7.087 | 72793 | 1.78 | 1836 | 0.43 |
| 7.673 | 55056 | 1.35 | 2274 | 0.53 |
| 11.987 | 19815 | 0.48 | 1379 | 0.32 |
| Totals | 4087508 | 100.00 | 430884 | 100.00 |

(4) HPLC trace of **1d**

**1d,** $R_t$ = 2.98 min

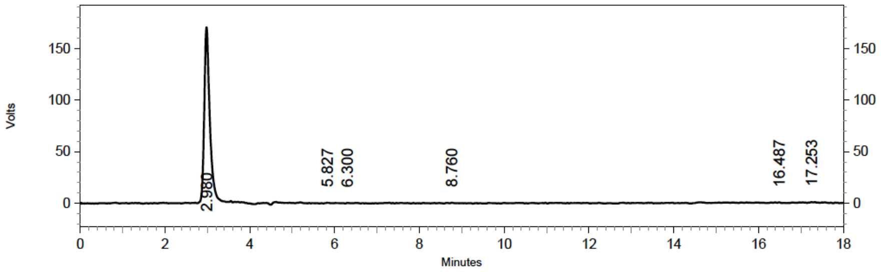


**DAD: Signal A,**
**280 nm/Bw:4 nm**
**Results**

| Retention Time | Area | Area % | Height | Height % |
|---|---|---|---|---|
| 2.980 | 3265485 | 95.52 | 357050 | 97.92 |
| 5.827 | 23674 | 0.69 | 1381 | 0.38 |
| 6.300 | 14342 | 0.42 | 1229 | 0.34 |
| 8.760 | 28198 | 0.82 | 1389 | 0.38 |
| 16.487 | 18731 | 0.55 | 1576 | 0.43 |
| 17.253 | 68362 | 2.00 | 2028 | 0.56 |
| Totals | 3418792 | 100.00 | 364653 | 100.00 |

(5) HPLC trace of **1e**

**1e,** $R_t$ = 3.00 min

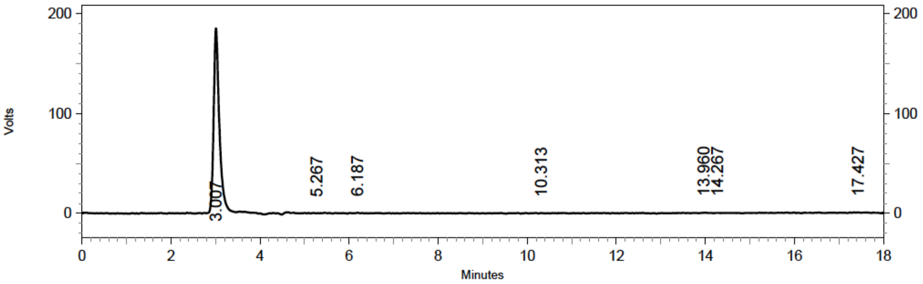


**DAD: Signal A,**
**280 nm/Bw:4 nm**
**Results**

| Retention Time | Area | Area % | Height | Height % |
|---|---|---|---|---|
| 3.007 | 3435313 | 96.56 | 387705 | 98.32 |
| 5.267 | 7606 | 0.21 | 689 | 0.17 |
| 6.187 | 18837 | 0.53 | 1559 | 0.40 |
| 10.313 | 24065 | 0.68 | 927 | 0.24 |
| 13.960 | 13319 | 0.37 | 1205 | 0.31 |
| 14.267 | 18752 | 0.53 | 821 | 0.21 |
| 17.427 | 39816 | 1.12 | 1411 | 0.36 |
| Totals | 3557708 | 100.00 | 394317 | 100.00 |

(6) HPLC trace of **2a**

**2a,** $R_t$ = 11.46 min

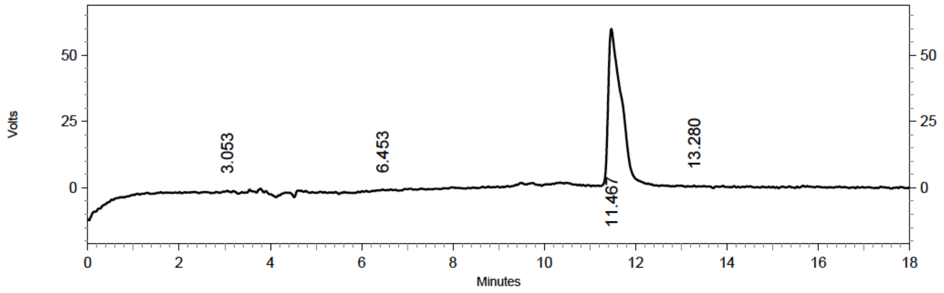


**DAD: Signal A,**
**280 nm/Bw:4 nm**
**Results**

| Retention Time | Area | Area % | Height | Height % |
|---|---|---|---|---|
| 3.053 | 27623 | 1.09 | 1876 | 1.46 |
| 6.453 | 26293 | 1.04 | 1334 | 1.04 |
| 11.467 | 2476755 | 97.55 | 124152 | 96.81 |
| 13.280 | 8286 | 0.33 | 876 | 0.68 |
| Totals | 2538957 | 100.00 | 128238 | 100.00 |

(7) HPLC trace of **2b**

**2b,** $R_t$ = 10.80 min

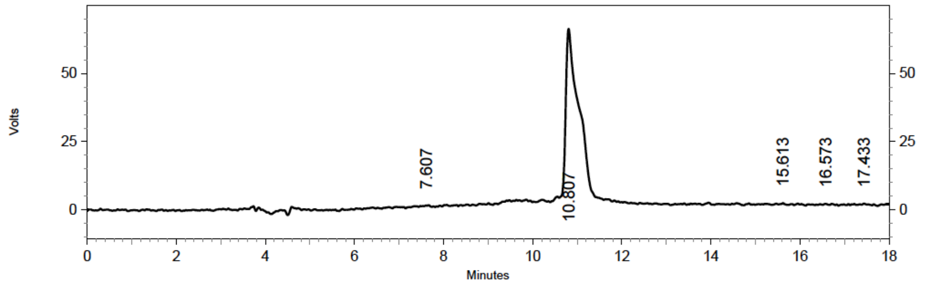


**DAD: Signal A,**
**280 nm/Bw:4 nm**
**Results**

| Retention Time | Area | Area % | Height | Height % |
|---|---|---|---|---|
| 7.607 | 35855 | 1.29 | 1480 | 1.08 |
| 10.807 | 2626734 | 94.66 | 130403 | 95.55 |
| 15.613 | 30280 | 1.09 | 1487 | 1.09 |
| 16.573 | 35370 | 1.27 | 1359 | 1.00 |
| 17.433 | 46566 | 1.68 | 1748 | 1.28 |
| Totals | 2774805 | 100.00 | 136477 | 100.00 |

#### 3. Experimental protocols:

##### 3.1. General methods:

Stock solutions of MXF (1 mM), compounds **1a-1h, 2a** and **2b** (1 mM) were prepared in DMSO whereas the stock solutions of *E. coli* NTR (1 mg/mL) and NADH (5 mM) were prepared in phosphate buffer (pH 7.4, 10 mM). Unless otherwise specified, the reaction mixtures were prepared in 10 mM pH 7.4 phosphate buffer and the fluorescence corresponding to MXF ($\lambda_{ex}$ = 289 nm and $\lambda_{em}$ = 488 nm) was measured using an EnSight Multimode Plate Reader (PerkinElmer).

##### 3.2. Monitoring the fluorescence of MXF by fluorimetry

MXF (10 µL, 1 mM stock) was diluted with 990 µL of buffer in a 1.5 mL eppendorf tube and then transferred into a micro-fluorescence cell (Hellma, path length 1.0 cm). Fluorescence spectra ($\lambda_{ex}$ = 289 nm and $\lambda_{em}$ = 488 nm) was recorded using a HORIBA Jobin Yvon Fluorolog fluorescence spectrophotometer with an excitation and emission slit width of 1 nm.

##### 3.3. Evaluating the stability of NTR-MXF prodrugs in buffer

10 µM of prodrug (10 µL, 1 mM stock) was added to a 1.5 mL eppendorf tube containing 990 µL of buffer and placed in an Eppendorf thermomixer comfort (700 rpm) at 37 ºC. Aliquots (100 µL) were dispensed into the 96-well microtiter plate at indicated time points. The fluorescence signal attributable to MXF was measured using a microplate reader (EnSight).

##### 3.4. Monitoring the release of MXF upon chemoreduction of prodrugs

**(A) Sodium dithionite induced chemoreduction of NTR-MXF prodrugs**

A stock solution of sodium dithionite (38.2 mM) was prepared in 15 mL of methanol-water (1:1 v/v). A solution of 10 µM of prodrug (2 µL, 1 mM stock) was added to 198 µL of 38.2 mM $Na_2S_2O_4$ solution in a 96-well plate. The plate was incubated at 25 ºC inside microplate reader (EnSight) with shaking (60 rpm) and the fluorescence response was recorded after 30 min.

**(B) Zinc/Ammonium formate mediated chemoreduction of prodrugs**

Chemoreduction assays were conducted as previously reported with some modifications.[9] A stock solution of ammonium formate (3 mM) was prepared in deionised water. The reaction samples were prepared by adding 30 µM of ammonium formate (18 µL, 3 mM stock) followed by zinc dust (15 mg) to a 2 mL eppendorf tubes containing a solution of 10 µM of prodrugs (18 µL, 1 mM stock) in 1764 µL of methanol-water (1:1 v/v). The reaction mixtures were incubated at 37 ºC in an Eppendorf thermomixer comfort (800 rpm). Aliquots (250 µL) were periodically taken from the reaction mixture, centrifuged (9,391 × *g* for 2 min), and the supernatant (200 μL) was then carefully transferred to 96-well plate (Tarson). Reduction of the prodrugs were monitored by measurement of fluorescence using a microplate reader (EnSight).

##### 3.5. Kinetics of dithionite-dependent reduction of NTR-MXF prodrugs

Varying concentrations of sodium dithionite ($Na_2S_2O_4$; 2.5-10 mM) in 15 mL of methanol-water (1:1 v/v) were independently prepared. Reduction reactions were performed in a 96-well microtiter plate by addition of 10 µM of compound (2 µL; 1 mM stock) to 198 µL of $Na_2S_2O_4$ solution with the final concentration ranged from 2.5 mM to 10 mM. The kinetics of dithionite-dependent changes in fluorescence was recorded at 25 ºC for a period of 4 min (frequency of measurement was 1 s) using a microplate reader (EnSight).

##### 3.6. Investigating the release of MXF from prodrugs using a fluorescence-based assay

The reactions in 96-well plate were initiated by the addition of 15 nM *E. coli* NTR (30 µL; 0.1 µM) and 100 µM of NADH (4 µL; 5 mM stock) to a solution of 10 µM prodrugs (2 µL, 1 mM stock) in 164 µL of buffer. The enhancement in fluorescence signal corresponding to the release of MXF relative to NTR untreated control was recorded at 37 ºC using a microplate reader (EnSight) for 1 h.

##### 3.7. *In situ* detection of MXF generated from prodrugs in *M. smegmatis* lysates using fluorescence-based assay

*M. smegmatis* $mc^2155$ bacteria was grown with agitation of 180 rpm in Middlebrook 7H9 broth media supplemented with glycerol (0.2%), and tween-80 (0.1%) at 37 ºC for 24 h. The bacterial cells were harvested by centrifugation at 2486 × *g* for 15 min at 4 ºC. The bacterial pellets were washed twice with PBS buffer (1x, pH 7.4), resuspended in PBS (1x, 10 mL) and transferred into a microcentrifuge tube. The cells were lysed by sonication using (130 W ultrasonic processor, VX 130W) stepped microtip for 5 min pulse on time (with 3 s ON and 3 s OFF pulse, 60% amplitude, 20 kHz frequency) under ice cold conditions. The total protein concentration of the whole cell lysate was determined by Bradford assay using bovine serum albumin (BSA) and further adjusted to 1 mg/mL with PBS (1x). The reactions were conducted by treatment of 10 µM of various prodrugs (5 µL, 1 mM stock) with 495 µL of *M. smegmatis* lysate (1 mg/mL) at 37 ºC on an Eppendorf thermomixer comfort (800 rpm). An aliquot (100 µL) was taken post 1 h of incubation, transferred to 96-well microplate and the fluorescence for the release of MXF was recorded using a microplate reader (EnSight).

##### 3.8. Theoretical calculation of reduction potentials of prodrugs

Density functional theory (DFT) was used to compute reduction potentials for the substrates of interest. B3LYP,[10] functional as implemented in Gaussian16[11] with 6-31G(d)[12] basis set was used for geometry optimizations and computing vibrational frequencies. This level of theory was shown to be appropriate for computing reduction potentials via benchmarks in literature.[13] Chlorobenzene was used as the solvent to mimic the enzymatic environment and was approximated via polarized continuum model. Furthermore, reduction potentials in acetonitrile were also computed. Thermal corrections were computed using quasi-rigid rotor harmonic oscillator approximation.[14] Single point energies were computed at PCM(Chlorobenzene)-UWB97XD/aug-cc-pVDZ.[15-18] Molecular graphics were generated using CYLView.

The Nerst equation was used to compute the redox potentials (equation S1) where n is the number of electrons (1 for all the species investigated), $\mathscr{F}$ is the Faraday constant (a value of 23.061 kcal/mol-V), $E_{calc}^0$ is the computed reduction potential, and $\Delta G^‡$ is the free energy difference between the reduced form and the oxidized form of the species under investigation. The computed $E_{calc}^0$ was further corrected by subtracting the absolute value for the standard hydrogen electrode (4.281 V) and ferrocene/ferrocenium couple (-0.559 V). These are then compared to the literature values for the experimental redox potentials in acetonitrile.[19]

$$\Delta G^‡ = -\mathrm{n}.\mathcal{F}.E_{\mathrm{calc}}^0$$

**Equation S1.** Nernst equation used for computing reduction potentials.

A correlation plot between the experimental reduction potentials and computed reduction potentials further validate the authenticity of the chosen level of theory (Figure S4). Driving force was computed from the experimentally measured reduction potentials of the substrates and FMN. Correlation plot between the driving force and $k_{cat}/K_m$ reveal a weak correlation between the electron transfer rates and the release of MXF (Figure S8). However, accurate electron transfer rate computations weren’t performed owing to the complexity in computing the solvent reorganization energies inside the enzyme active site. The computed driving forces demonstrate a reasonable agreement with the observed release profiles of MXF.

##### 3.9. Electrochemical studies by Cyclic voltammetry

Cyclic voltammetry studies were conducted using a standard three-electrode setup connected to a CHI760E electrochemical workstation. Glassy carbon and platinum wire were used as the working electrode and counter electrode respectively. Ag/AgCl was used as a reference electrode. These experiments were conducted under an atmosphere of argon in anhydrous ACN with tetra-butyl ammonium hexafluorophosphate (TBAP; 0.1 M) as supporting electrolyte. Stock solutions of compounds **1a-1e, 2a** and **2b** (10 mM in DMSO) were prepared, added to respective 25 mL volumetric flasks and diluted (20-fold; final conc. 0.5 mM) with electrolyte solution. The cyclic voltammograms (CV) were recorded at 20 ºC in a potential range between -2 V to +2 V at a scan rate of 100 mV/s with 10 sweep segments, 0.001 V sample interval, 2 s quiet time and $1 \times 10^{-5}$ A/V sensitivity. Potentials (V) were calibrated using an internal standard $Fc/Fc^{+}$ redox couple and are reported vs. Ag/AgCl.

##### 3.10. Determination of steady state apparent kinetic parameters for NTR-mediated release of MXF from ester and carbamate derivatives (1a-1e and 2a-2b)

Two-fold serial dilutions were prepared from compound stocks (10 mM in DMSO) yielding a concentration range of 0 mM – 4 mM. Kinetic measurements were carried out in total volume of 200 µL/well containing 164 µL of buffer, 100 µM of NADH (4 µL; 5 mM stock) with varying substrate concentrations ranged from 0 µM - 40 µM (2 µL; 0 to 4 mM stock). The final DMSO concentration was 1%. Reactions were initiated by the addition of 15 nM *E. coli* NTR (30 µL; 0.1 µM stock) with a multichannel pipette. Control wells contained only substrates with or without either NADH or NTR. Fluorescence attributable to formation of MXF was followed using a microplate reader in a 96-well plate format. The following parameters were used for fluorescence measurement: Readings were collected from the top at every 37 s intervals for a period of 40 min (for prodrugs **1a-1d** and **2a** at 37 ºC) or at every 1 s interval for a period of 4 min (for prodrug **1e** and **2b** at 25 ºC) with 15 flashes per well and a focus height adjusted to 9.5 mm. The background fluorescence was subtracted for each time point from the total fluorescence signal to obtain corrected relative fluorescence intensity (RFI) values. Next, the change in fluorescence (ΔF, RFI/min) upon addition of enzyme was obtained by subtracting the fluorescence signal in the absence of enzyme ($F_o$) of the varying concentrations of substrates for given time points from the total fluorescence signal (F). The change in fluorescence (ΔF) vs. time was plotted. The standard curve of MXF was generated by plotting the corrected RFI values vs. concentration (Figure S7). The slope and intercept were determined using linear regression ($R^2$ = 0.9934) and used to calculate the amount of MXF formed. The RFI/min was converted into µM/min of MXF produced from various concentrations of substrates using equation S2.

$$\mu M/min = \frac{\Delta F\ (RFU/min) - Intercept\ (Std.curve)}{Slope\ of\ Std.\ curve\ (RFU/\mu M)} \quad \text{...equation S2}$$

Linear regression analysis was performed to determine the initial reaction rates by taking the slope during the linear stage (0-12 min for **1a**; 0-2 min for **1b** and **1c**; 0-6 min for **1d**; 0-0.3 min for **1e** and **2b**; 0-12.5 min for **2a**) of the resulting curves of varying concentrations of substrates using OriginPro 8.5.1. These initial reaction velocities were replotted against substrate concentrations and kinetic parameters ($V_{max}$, $K_m$ and $k_{cat}/K_m$) for each concentration were calculated by non-linear regression using the GraphPad Prism 9.1 according to the Michaelis-Menten equation (equation S3). All data are shown as mean ± SD for three biological replicates.

$$V = \frac{V_{max}\,[S]}{K_m+[S]} \qquad \text{...equation S3}$$

##### 3.11. Monitoring the release of MXF from 1d

###### (A) Fluorimetry studies

To an eppendorf tube (1.5 mL) containing 10 µM solution of **1d** (10 µL, 1 mM stock) in 870 µL of buffer, 100 µM of NADH (20 µL, 5 mM stock) and varied concentrations of NTR ranged from 0 to 60 nM (100 µL, 0 to 0.6 µM stock) were added. Control reactions containing only **1d** or lacking NADH or NTR were also carried out. These reaction mixtures were transferred to a micro-fluorescence cell (Hellma, path length 1.0 cm) following incubation at 37 ºC on an Eppendorf thermomixer comfort (700 rpm) for 30 min. The fluorescence measurements were carried out using a HORIBA Jobin Yvon Fluorolog fluorescence spectrophotometer.

###### (B) Fluorescence-based analysis

A similar experiment as mentioned in Section 3.10.A was carried out in a 96-well plate to monitor the enzyme catalysed reaction of **1d** with different concentrations of NTR. The reaction mixture either contained only 10 µM of **1d** (2 µL, 1 mM stock) or 10 µM of **1d** (2 µL, 1 mM stock), 100 µM of NADH (4 µL, 5 mM stock) with varied concentrations of NTR ranged from 0 to 60 nM (30 µL, 0 to 0.4 µM stock) in 164 µL of buffer. The reactions were incubated at 37 ºC, and the progression of the NTR-dependent reaction was assessed by change in fluorescence over time using a microplate reader (EnSight) with readings collected from top at every 30 s intervals for a period of 60 min with the same parameters as stated earlier in Section 3.10.

##### 3.12. Sequence alignment

The sequences of NfsB (*E. coli*; UniProtKB ID: P38489) and NfnB (*M. smegmatis*; UniProtKB ID: A0R6D0) were obtained from Uniprot database. The unweighted sequence alignments between NfsB and NfnB from *E. coli* and *M. smegmatis* were performed using T-coffee at the

European Bioinformatics Institute website (https://www.ebi.ac.uk) using the default settings and displayed using Jalview. The sequence name indicates the organism of origin, the Uniprot code and the numbers indicate the amino acid residues displayed. The consensus symbols: ‘*’, ‘:’ and “.” under the amino acids indicate identical, conserved and semi-conserved residues respectively.

##### 3.13. *In silico* molecular docking studies

The structures of compounds (**1a-1e, 2a** and **2b**) were built with standard bond length and angles using ChemDraw and then energy minimized with Chem3D using the integrated MM2 energy minimization script. The X-ray crystal structures of FMN-bound oxidized *E. coli* NTR (PDB ID: 1DS7; resolution = 2.0 Å), and FMN-bound *M. smegmatis* NfnB (PDB ID: 2WZW; resolution = 1.8 Å) were retrieved from PDB. The protein and ligand PDBQT files were prepared using AutoDock Tools 1.5.6 (ADT) following the standard protocol. The following docking parameters of FMN-bound active sites were employed for (a) *E. coli* NTR (PDB ID: 1DS7): grid box ($9.52 \times 9.52 \times 9.52$ Å$^3$) at the coordinates (x = 18.533, y = -32.728, z = 48.114); and (b) *M. smegmatis* NfnB (PDB ID: 2WZW): grid box ($9.6 \times 9.6 \times 9.6$ Å$^3$) at the coordinates (x = -14.253, y = -81.909, z = -25.468) with default settings: exhaustiveness = 24, energy range = 3 kcal/mol and number of modes = 20. The best-scored docking pose with the lowest binding energy was selected for analysis and figures were visualized using PyMOL (The PyMOL Molecular Graphics System, Version 2.0 Schrödinger, LLC).

##### 3.14. Molecular dynamics simulations

Molecular dynamics simulations were performed using Amber 20. Parameter files for the protein were generated using ff14SB and are provided as separate attachments. Parameter files were generated using Antechamber with the general amber force field (gaff) and the BCC charge model. Protonation states were predicted using H++ server. Each protein was immersed in a pre-equilibrated cubic box with an 8 Å buffer of OPC water molecules using tLeap resulting in a box size of 8.5nm. A total of 16,000 water molecules were added to the system. 12-14 sodium atoms were added using tLeap to neutralize the charges in the system. A multi-step relaxation was performed on the system. First, the position of solvent molecules and ions was minimized by adding positional restraints on the solute by a harmonic potential with a force constant of 50 kcal/mol/Å$^2$. The position restraint is then lowered to 10 kcal/mol/Å$^2$ on the solute and water molecules and the system was gradually brought to a temperature of 300 K over 1 ns with a 1fs time step. The resulting system was then equilibrated for another 1 ns with reduced restraints of 1 kcal/mol/Å$^2$ to reach a constant pressure of 1 atm. Finally, production runs of 10 ns with a time step of 2 fs was performed at constant volume and temperature (300 K). SHAKE algorithm was used for water molecules. Long range electrostatic effects were modelled using the particle mesh Ewald method. The protein structures thus obtained were then overlayed (Figure S10) and the RMSD values obtained for these overlays were around 1 Å compared to the structure obtained after the production run for 1DS7 with **1e** docked. The RMSDs for each of the proteins are tabulated in Table S5. Trajectory analysis was performed using cpptraj. RMSD over the entire 10 ns run was evaluated using cpptraj for C, CA, and N atoms (Figure S11). The RMSD values for these atoms were found to be around 1.5 Å across prodrugs **1a-1e** and **2a-2b**. Similarly, an RMSF analysis was carried out using cpptraj on **1e** bound to 1DS7 (Figure S12). This analysis revealed that over the course of the MD simulation, the residues that fluctuated the most are situated far away from the active site in the outer most loops. Furthermore, the prodrugs themselves suffered significant deviation from the equilibrated structure as indicated by the large fluctuation of about 2.5 Å in residue 1 (prodrug **1e**). Binding energies for these systems were computed at HF-3c level of theory using ORCA 5.0.1. All the residues within 5 Å of the prodrug were extracted and capped with hydrogen atoms. FMN residue was also included in each of these computations. No clear trend was observed in the binding energies. Similarly, the dispersion contributions to the binding energies were also evaluated using Grimme's DFTD3 scheme with Becke-Johnson corrections.

##### 3.15. LC-MS analysis of NTR-dependent decomposition of 1e

The reaction mixture was prepared by adding compound 20 µM of **1e** (10 µL, 2 mM stock) to a solution of 100 µM NADH (20 µL, 5 mM stock) and 15 nM of NTR (150 µL; 0.1 µM) in 820 µL of buffer. The reaction samples containing **1e** or MXF served as reference controls, while the reaction mixture with NADH and NTR was used as blank. The reaction mixtures were incubated at 37 ºC on an Eppendorf thermomixer comfort (700 rpm) for 30 min. An aliquot (100 µL) was withdrawn from the reaction mixtures and centrifuged (9,391 × *g*) at RT for 5 min. The supernatants (100 µL) were carefully sampled, diluted with ACN (100 µL) and assessed thereafter by LC/MS. All measurements were done using a LC-MS method in the positive ion mode using high resolution multiple reaction monitoring (MRM-HR) analysis on a Sciex X500R quadrupole time-of flight (QTOF) mass spectrometer fitted with an Exion UHPLC system using a Kinetex 2.6 mm hydrophilic interaction liquid chromatography (HILIC) column with 100 Å particle size, 150 mm length and 3 mm internal diameter

(Phenomenex). Nitrogen was the nebulizer gas, with the nebulizer pressure set at 50 psi. MS was calibrated in positive mode and samples were injected (50 µL) and analyzed with the following parameters: Mode: Electrospray ionization (ESI), ion source gas 1 = 40 psi, ion source gas 2 = 50 psi, curtain gas = 30, CAD gas = 7, spray voltage = 5500 V and temperature = 500 ºC. The MRM-HR mass spectrometry parameters are: MXF (Q1, M + $H^+$) = 402.18, **1e** (Q2, M + $H^+$) = 544.17, **6a** (Q3, M + $H^+$) = 514.19, **6b** (Q4, M + $H^+$) = 530.19, **11** (Q5, M + $H^+$) = 131.03, declustering potential = 80 V, declustering potential spread = 20 V, collision energy = 10 V, collision exit potential = 5 V and accumulation time = 0.24 s. The LC runs were for 30 min with gradient of 100% solvent A (0.1% HCOOH in milliQ water) for 5 min, linear gradient of solvent B (ACN, 0% to 100%) for 25 min followed by 100% solvent A for 5 min all at a flow rate of 0.5 mL per min.

##### 3.16. Time- and concentration-dependent kinetics of 1e with NTR

The reaction mixture contained 10 µM of **1e** (10 µL, 1 mM stock), with or without 100 µM of NADH (20 µL, 5 mM stock) in 870 µL of buffer were independently added to a micro-fluorescence cell (Hellma, path length 1.0 cm). The cuvette was placed in slowly stirring condition using magnetic stirred equipped in fluorescence instrument ($t$ = 0 s). The fluorescence response was continuously recorded at an $\lambda_{em}$ = 488 nm ($\lambda_{ex}$ = 289 nm) either at 25 ºC or 37 ºC on a Fluoromax-4 spectrophotometer (Jobin Yvon Edison) following addition of NTR ranged from 0 to 15 nM (100 µL, 0 to 0.15 µM stock) at ($t$ = 50 s) for 150 s. The time-axis was normalized according to equation S4 and fluorescence intensity vs. time was plotted.

$$t = t - 50 \qquad \ldots\textbf{equation S4}$$

##### 3.17. Evaluating the selectivity of 1e

Stock solutions of porcine liver esterase (50 U/mL), GSH (10 mM), Cys (10 mM), vitamin-C (10 mM), histidine (10 mM), and $H_2O_2$ (30%, 10 mM) in phosphate buffer were prepared independently from commercial sources. In a typical reaction, compound 10 µM of **1e** (2 µL, 1 mM stock), 100 µM of NADH (4 µL, 5 mM stock) and 15 nM of NTR (30 µL; 0.1 µM stock) were added to 164 µL of buffer. Similarly, a reaction mixture of 10 µM of **1e** (2 µL, 1 mM stock) and 100 µM of various analytes (2 µL, 10 mM stock) were prepared in 196 µL of buffer. In a separate experiment, the reaction mixture was prepared by adding compound 10 µM of **1e** (2 µL, 1 mM stock) along with 1 U/mL esterase (4 µL, 50 U/mL stock) in 194 µL of buffer. The reaction mixtures were incubated for 15 min at 37 ºC, and then fluorescence response was recorded using a microplate reader (EnSight).

##### 3.18. Assessing the stability of 1e in mammalian cellular lysate by fluorescence

MEF cells were grown in culture flask in complete DMEM medium supplemented with 5% FBS (fetal bovine serum) and 1% antibiotic solution in an atmosphere of 5% $CO_2$ at 37 ºC. When the cells were 70% confluent, old media was removed and the cells were washed with PBS buffer (1x). The cells were then detached by trypsinization, subsequently resuspended in PBS (1x) and transferred to a microcentrifuge tube. The cells were lysed by sonication using (130 W ultrasonic processor, VX 130W) stepped microtip for 2 minutes (with 5 s ON and 10 s OFF pulse, 60% amplitude) under ice cold conditions. The total protein concentration of the whole cell lysate was determined by Bradford assay and further adjusted to 1 mg/mL with PBS (1x). The stability of **1e** was assessed by treatment of 10 µM of **1e** (5 µL, 1 mM stock) with 495 µL of whole cell lysate in PBS (pH 7.4, 10 mM) at 37 ºC on an Eppendorf Thermomixer comfort (800 rpm). At predetermined time points, aliquots (100 µL) were transferred to a 96-well microplate and the fluorescence was recorded using a microplate reader (EnSight).

##### 3.19. Stability of 1e in human plasma and serum

The use of human blood plasma and serum was approved by Institutional Human Ethics Committee at Indian Institute of Science (IISc), Bangalore, India (Approval number: 16/17.03.2022). Fresh whole blood (5 mL) was collected separately in commercially available red (without anticoagulant) or lavender coloured (with EDTA as anticoagulant) vacutainers. The whole blood in the red coloured vacutainer was allowed to clot, centrifuged at 21,130 ×*g* for 10 min to obtain the supernatant serum. Similarly, the whole blood from lavender coloured vacutainer was immediately centrifuged at 21,130 × *g* for 10 min at 4 ºC, and the resultant supernatant plasma was collected. The plasma (970 µL) or serum (242.5 µL) was then incubated with 30 µM of **1e** (30 µL or 7.5 µL respectively, 1 mM stock). Aliquots (50 µL) were taken at different time points (0, 0.5, 2, 4, 6, 18 and 24 h for plasma and 0, 2 and 6 h for serum), mixed vigorously with ACN (100 µL), centrifuged (3×) at 21,130 × *g* for 10 min at 4 ºC. The clear supernatant was collected and assessed thereafter by LC-MS with identical parameters described in Section 3.15.

##### 3.20. *In situ* detection of MXF generated from 1e in *E. coli, M. smegmatis* (WT and ∆mshA) lysates using fluorescence-based assay

*E.coli* ATCC 25922 was grown in LB media at 37 ºC in a rotary shaker for overnight. The bacterial cells were harvested by centrifugation at 2486 × *g* for 15 min at 4 ºC. The bacterial pellets were washed twice with PBS buffer (1x, pH 7.4), resuspended in PBS (1x, 2 mL) and transferred into a microcentrifuge tube. The cells were lysed by sonication using (130 W ultrasonic processor, VX 130W) stepped microtip for 2 minutes pulse on time (with 5 s ON and 10 s OFF pulse, 60% amplitude, 20 kHz frequency under ice cold conditions. The total protein concentration of the whole cell lysate was determined by bradford assay using bovine serum albumin (BSA) and further adjusted to 1 mg/mL with PBS (1x).The whole cell lysates (1 mg/mL) of *M. smegmatis* mc$^2$155 (WT and Δ*mshA*) were prepared following the protocol described in Section 3.7. The reactions were conducted by treatment of 10 µM of **1e** (5 µL, 1 mM stock) with 495 µL of *E.coli* or *M. smegmatis* lysate (1 mg/mL) at 37 ºC on an Eppendorf thermomixer comfort (800 rpm). An aliquot (100 µL) was taken post 1 h of incubation (or post 2 h in *E. coli* lysate), transferred to 96-well microplate and the fluorescence was recorded. A similar protocol was followed to determine the specificity of **1e** towards *E.coli* or *M. smegmatis* NTR by treatment of 485 µL bacterial lysate with 100 µM of DCOM, a NTR inhibitor(10 µL, 10 mM) for 10 min prior to addition of 10 µM of **1e** (5 µL, 1 mM stock). Furthermore, a control experiment was performed by incubating **1e** in heat-inactivated bacterial lysates. These lysates were prepared by heating the bacterial lysates at 90 ºC for 45 min on an Eppendorf Thermomixer comfort (800 rpm) followed by cooling down to 37 ºC.

##### 3.21. Determination of Minimum Inhibitory Concentrations (MICs) of prodrugs

**(A) Against *M. tuberculosis*:**

All *Mtb* strains were grown in Middlebrook 7H9 broth (BD, Difco) supplemented with 0.2% glycerol, 0.1% Tween-80 and ADS (0.5% albumin, 0.2% dextrose and 0.085% NaCl) with shaking (180 rpm) in a rotary shaker incubator (Lab Therm LT-X). Minimal inhibitory concentration (MIC) of *Mtb* strains were determined by a resazurin microtiter assay (REMA) using 96-well flat-bottom plates.[20] Briefly, 100 µL of Middlebrook 7H9 broth supplemented with 0.2% glycerol and 10% (v/v) ADS was dispensed in each well of a 96-well plate (except peripheral wells), and serial 2-fold dilutions of each drug were prepared directly in the plate. Bacteria, cultured in 7H9 (supplemented with 0.2% glycerol, 0.1% Tween-80)−ADS medium and grown until exponential phase ($OD_{600}$ = 0.4−0.8) were diluted in experimental medium. An inoculum (100 µL) from each strain (approximately $1\times10^5$ bacteria/well, by OD) was added in triplicate wells. A growth control and a media control were also included for each strain. After 5 days of incubation at 37 ºC in the presence of drug, 30 µL of 0.02% (w/v) resazurin was added, and plates were incubated for an additional 24 h. Fluorescence intensity

($\lambda_{ex}$ = 530 nm and $\lambda_{em}$ = 590 nm) was measured using a SpectraMax M3 plate reader in the top-reading mode and the percentage growth inhibition was calculated relative to an untreated control. MIC was taken as the lowest drug concentration that resulted in at least 90% reduction in fluorescence compared to an untreated growth control.

**(B) Against ESKAPE pathogens:**

Antibiotic susceptibility testing of compounds against *S. aureus* ATCC 29213, *E. coli* ATCC 25922, *P. aeruginosa* ATCC 27853 and resistant clinical isolates of *E. coli* and *S. aureus* was carried out according to the CLSI (Clinical and Laboratory Standards Institute) guidelines for broth microdilution assay.10 mg/mL stock solutions of test compounds were prepared in DMSO. Bacterial cultures were inoculated in Muller Hilton broth II (MHB II) and OD of the cultures was measured at 600 nm, followed by dilution to achieve ~$10^5$ CFU/mL. The compounds were tested ranging from 64 to 0.03 mg/L in a 2-fold serial diluted fashion with 2.5 μL of each concentration added to each well of a 96-well round-bottom microtiter plate. Later, 97.5 μL of bacterial suspension was added to each well containing the test compound along with appropriate controls. The plates were incubated at 37 ºC for 18−24 h, following which, the MIC was identified. MIC of a compound is defined as the lowest concentration of compound inhibiting complete visible growth.

##### 3.22. Determination of bacterial survival (*M. tuberculosis* H37Rv) with 1e

The bactericidal activity was assessed by the time-kill method. Time-kill curves were obtained by treating exponentially growing cultures of *Mtb* ($OD_{600}$ of 0.1 or ~$1.6\times10^7$ cells/mL) with minimum bactericidal concentration (MBC) of MXF and **1e**. Tubes (50 mL) were incubated with shaking (180 rpm) for 8 days at 37 ºC. Aliquots were taken at various intervals, serially diluted, and plated for CFU enumeration.

##### 3.23. Determination of mycobactericidal activity of 1e on *Mtb* infected macrophages

The human monocytic cell line (THP-1) was grown in RPMI-1640 medium supplemented with 10% heat-inactivated (55 ºC) fetal bovine serum (MP Biomedical). A total of $3\times10^4$ cells/well was seeded into a 96-well cell-culture plate. THP-1 monocytes were differentiated into macrophages by a 16-18 h treatment with 20 ng/mL phorbol-12-myristate 13-acetate (PMA). Cells were then rested for 48 h post-PMA treatment to reduce the influence of PMA on the activation status of THP-1 cells.[21] PMA-differentiated THP-1 cells were infected with *Mtb* H37Rv at multiplicity of infection (MOI) of 2 and incubated at 37 ºC in 5% $CO_2$. After 4 h of infection, cells were washed twice with warm RPMI and replenished with complete RPMI containing 0.2 mg/mL amikacin (AMK) for 2 h to remove any remaining extracellular bacteria. Subsequently, cells were washed and maintained in complete RPMI for the rest of the experiment. Compounds were diluted in complete media, added to respective wells at different concentrations and were incubated for various times. For colony-forming unit (CFU) determination, infected cells were lysed in PBS containing 0.06% sodium dodecyl sulphate (SDS); dilutions were prepared using 7H9 medium, and aliquots were plated on 7H11-OADC agar plates. Plates were incubated at 37 ºC for 2-4 weeks and colonies were counted.

##### 3.24. Effect of MXF and 1e on biosensor oxidation

Log phase cultures (10 mL) of *Mtb*-roGFP2 ($OD_{600}$ = 0.3) were treated with different concentrations of **1e** and MXF (2.5 μM; 10× MIC) for 6, 12 and 24 h and incubated in shaker incubator (180 rpm, 37 ºC). The fluorescence was measured at the fixed emission (510 nm) after excitation at 405 and 488 nm. The biosensor response was acquired using the BD FACSAria Fusion flow cytometer. The ratiometric analysis of the biosensor was normalized using the culture treated with cumene hydroperoxide (10 mM) which is reported to oxidise the biosensor to maximal and dithiothreitol (20 mM) which is reported to reduce the biosensor to maximal. Ten thousand events were acquired per sample.

##### 3.25. OCR and ECAR measurements

To measure basal oxygen concentration rate (OCR) and extracellular acidification rate (ECAR), exponential phase *Mtb* cultures (OD ~ 0.6) were incubated in 7H9 media containing non-metabolizable detergent tyloxapol devoid of ADS/carbon source for 24 h. The single cell suspensions were prepared by 26-gauge syringe passage and 2 × $10^6$ cells/well were placed in the bottom of the Cell-Tak-coated XF culture plate (Agilent/Seahorse Biosciences). The measurements were done in Seahorse XFp analyzer (Agilent/Seahorse Biosciences) with cells in unbuffered 7H9 growth medium (pH 7.35 lacking $KH_2PO_4$ and $Na_2HPO_4$) containing glucose (2 mg/mL) as a carbon source. OCR and ECAR measurements were recorded for ~21 min (3 initial baseline readings) before addition of drug (**1e** or MXF) (2.5 μM; 10× MIC), which was delivered through the drug ports of the sensor cartridge (Wave Software, Agilent Technologies). CCCP was added through the drug ports at 416 min of the run. The OCR and ECAR were measured in the absence (untreated) and presence of the drug and CCCP. Changes in the OCR and ECAR readings by the drugs were calculated as a percentage of the third baseline reading for OCR and ECAR taken before addition of the drugs.

##### 3.26. Selection of MXF and 1e resistant colonies of *Mtb*

Exponentially growing *Mtb* ($5 \times 10^9$ cells/mL) were plated on 7H11-OADC agar containing different concentration of drugs. Resistant colonies were identified after incubation at 37 ºC for 4-6 weeks. The drug-resistant phenotype identified colonies were confirmed by plating them on drug-containing 7H11-OADC agar plates. MICs of different resistant colonies were calculated using the method described in Section 3.21.A.

##### 3.27. DNA sequencing of MXF and 1e resistant colonies of *Mtb*

The quinolone resistance determinant region (QRDR) was PCR amplified using genomic DNA isolated from drug-resistant colonies. The sequences of the primer pair used for this study is listed below.

| Entry | Primer | Primer sequence |
|---|---|---|
| 1 | Forward | 5'- GCAGCTACATCGACTATGCGATGAG - 3' |
| 2 | Reverse | 5'- CCTCGTCGATTTCCCTCAGCATCTC - 3' |

The PCR products were purified using Qiagen PCR purification kit and sent for DNA sequencing. The sequencing data were analysed and compared using MEGA (version 11.0.11).

##### 3.28. Time-kill kinetics of 1e against *E. coli* ATCC 25922 and *S. aureus* ATCC 29213

The bactericidal activity was assessed by the time-kill method. In brief, *S. aureus* ATCC 29213 or *E. coli* ATCC 25922 was diluted to $\sim 10^5$ CFU/mL in MHBII, treated with 1× and 10× MIC of **1e** and MXF and then incubated at 37 ºC with shaking for 24 h. Aliquots (100 µL) were collected at time intervals of 0, 1, 6 and 24 h, serially diluted and plated on MHA (Mueller Hinton Agar) followed by incubation at 37 ºC for 18−20 h. Kill curves were constructed by counting the colonies from plates and plotting the CFU/mL of surviving bacteria at each time point, in the presence and absence of a compound. Each experiment was repeated three times in duplicate and the mean data is plotted.

##### 3.29. Neutropenic murine thigh infection model

The use of mice for infectious studies (IAEC/2014/139 dated 03.12.2014) was approved by Institutional Animal Ethics Committee at CSIR-CDRI, Lucknow. Male balb/C mice weighing 24–26 g were housed 2 days prior to the start of the experiment, kept in groups of 4 per cage and given food and water *ad libitum*. To cause neutropenia in mice, cyclophosphamide was injected 4 days before infection, through the intraperitoneal route (150 mg/kg body weight), and 1 day before (100 mg/kg body weight) infection. 100 μL intramuscular infection of ~$10^7$ $log_{10}$ CFU/mL was given in the thigh of each mouse. The drug/compound was dosed through the intraperitoneal route, 3 and 6 h post-infection, to a group of five mice at a concentration of MXF (10 mg/kg body weight) and prodrug **1e** (10 mg/kg body weight). MXF was used as a reference compound, while untreated mice were the controls. Mice were sacrificed 24 h post-infection, thighs were removed, homogenized and dilutions were plated aseptically to estimate the infection load in each mouse.

##### 3.30. Determination of lethality against nutrient-starved non-replicating *Mtb* by REMA assay

The cultures of *Mtb* H37Rv were grown to exponential phase in roller-culture bottle containing Middlebrook 7H9 medium, supplemented with ADS, 0.2% glycerol and 0.05% Tween 80, at 37 ºC with rolling (at 6 rpm) in a roller incubator. Cultures grown to an $OD_{600}$ of ~0.6 were harvested by centrifugation (5000 rpm for 5 min) followed by two washes with PBS (pH 7.4). Bacterial cells were diluted to a final $OD_{600}$ of 0.3 in PBS supplemented with 0.02% tyloxapol. Eighty millilitres of this suspension was transferred into a roller culture bottle and incubated for 14 days to achieve starvation. Post starvation the culture was either left untreated or treated with isoniazid (INH; 10 μM), metronidazole (MZ; 10 mM), MXF (5 μM) and **1e** (5 μM) for 5 days. For the resazurin microtiter assay (REMA) the culture was pretreated in Middlebrook 7H9 medium, supplemented with ADS, at 37°C in a micro-titre plate for 2 days. After the incubation, resazurin solution was added to achieve a final concentration of 0.002% (w/v), and the plate was incubated for additional 48 h post which the fluorescence intensity was measured using SpectraMax M3 plate reader (Molecular Devices, San Jose, CA) in the top-reading mode ($\lambda_{ex}$ = 530 nm and $\lambda_{em}$ = 590 nm).

##### 3.31. Determination of lethality against *Mtb* under hypoxic conditions by HyRRA and CFU enumeration

For determination of lethality of the drugs under hypoxic conditions, bacterial cultures ($OD_{600}$ = 0.1) were placed in vacutainer tubes (Becton Dickinson) followed by incubation for 10 days at 37 ºC.[22] A high cell density ($OD_{600}$ = 0.1) was used for rapid achievement of hypoxia, which was observed as decolorization of methylene blue. Drugs were added to cultures at day 10 once hypoxia was established. Metronidazole (10 mM; MZ) and Isoniazid (10 µM; INH) were used as positive and negative controls, respectively. Drugs were injected in volumes of 100 µL in PBS following passage of argon through the drug solution to remove residual oxygen. Hypoxic cultures were treated with drugs (**1e** and MXF) for five days, similar to the incubation time for MIC determination with aerobically growing cells. After treatment, resazurin solution was added to achieve final concentration of 0.002% (w/v), and tubes were incubated for an additional 48 h to check visual change of colour from blue (absence or inhibition of growth) to pink (presence of growth). At this point, vacutainer tubes were unsealed, and end-point bacterial survival was determined by plating on drug-free 7H11-OADC agar, incubating for 3-4 weeks at 37 ºC, and determining CFU.

##### 3.32. Evaluation of accumulation of compounds within bacteria using LC-MS

**(A) Replicating *Mtb***

Exponential growing cells of *Mtb* ($OD_{600}$ = 0.6-0.7) were harvested as mentioned in Section 3.21.A and the pooled bacterial pellets were resuspended in PBS (pH 7.4, 10 mM with 0.1% Tween 80) to adjust an $OD_{600}$ 2-4. The bacterial suspension was aliquoted to different tubes. Drugs were added (5 µM) to the tubes and incubated at 37 ºC for 30 minutes. For CFU enumeration, bacterial suspension was collected from control tube, plated on 7H11-OADC agar and incubated at 37 ºC. After incubation, equal volume of pre-chilled (-80 ºC) silicone oil mix (9:1 AR20/sigma high temperature) was added to each tube containing the suspension and centrifuged (13,000 × *g*) for 5 min. The pellet was collected, suspended in PBS  and lysed through bead beating. The compounds were extracted from the resultant lysate using PBS-MeOH and filtered through a 3 kDa Amicon® centrifugal filter. The concentration of compounds were quantified using a LC-MS method in the positive ion mode on a Bruker Daltonics ESI-QTOF (Maxis Impact) mass spectrometer connected to a Thermo Dionex (Ultimate 3000) micro-LC system. Chromatographic separation was performed on C18 reverse phase column (4.6 × 150 mm, 2.7 µm; Agilent Poroshell 120) using a mobile phase of solvent A (0.1 % HCOOH in milliQ water) and solvent B (ACN) with a run time of 64 min and a multistep gradient 95:5 → 0–3 min, 95:5 to 5:95 → 3–50 min, 5:95 to 5:95 → 50–55 min, 5:95 to 95:5 → 55–64 min at a flow rate of 0.3 mL/min. The TOF MS/MS parameters are: MXF (Q1, M + $H^+$; m/z = 402.18 to Q3, M + $H^+$; m/z = 384.17), **1e** (Q1, M + $H^+$; m/z = 544.17 to Q3, M + $H^+$; m/z = 384.17 and Q1, M + $H^+$; m/z = 544.17 to Q3, M + $H^+$; m/z = 402.18 ) and CIP (Q1, M + $H^+$; m/z = 332.14 to Q3, M + $H^+$; m/z = 314.13). Blank untreated cell lysates were extracted using the procedure described above and were used to construct calibration curves to quantitate the intracellular concentration (nM) of compound/antibiotic in each respective sample. These are reported as accumulation values (nmol/CFUs) following normalization with CFU/mL of untreated bacteria.

**(B) Non-replicating *Mtb***

For preparing *Mtb* under hypoxic condition, modified protocol of Section 3.31 was used. 40 mL ($OD_{600}$ 0.1) bacterial culture were added in 50 mL conical tubes followed by incubation at 37 ºC in shaking condition. Achievement of hypoxia was observed as decolorization of methylene blue. The culture were grown till $OD_{600}$ ~0.6, harvested and were treated with compounds (MXF and **1e**) as mentioned above. The samples were prepared and assessed thereafter by LC-MS with identical parameters as described above.

#### 4. Figures

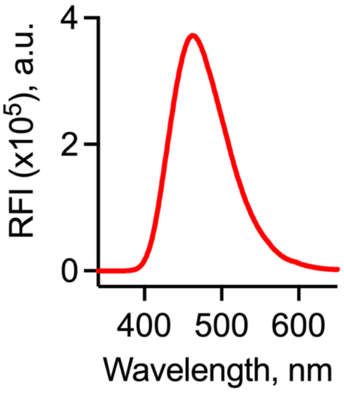


**Figure S1.** Fluorescence emission spectra of MXF (10 μM, $\lambda_{ex}$ = 289 nm) in phosphate buffer (pH 7.4, 10 mM) at 37 ºC. MXF exhibits a strong emission peak at 488 nm.

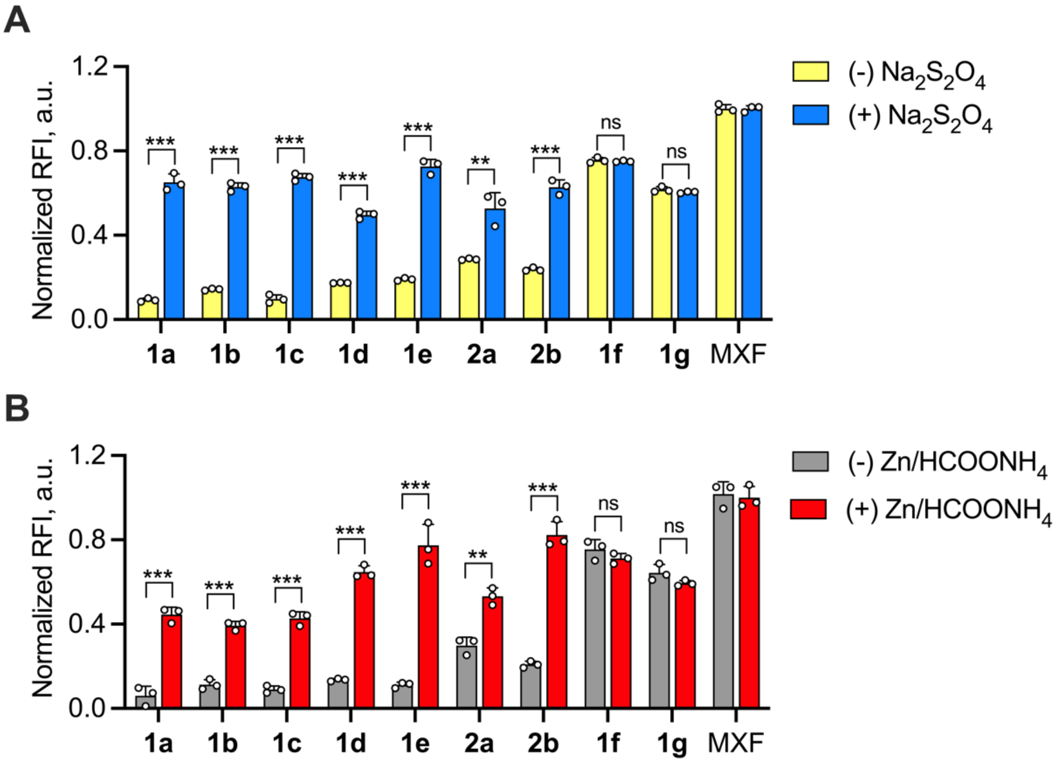


**Figure S2**. Screening and evaluation for release of MXF following (A) sodium dithionite ($Na_2S_2O_4$) and (B) Zinc/ammonium formate ($Zn/HCOONH_4$) based chemoreduction of compounds (10 μM) in $H_2O$:MeOH (1:1) in a fluorescence-assay. All the data represent mean ± SD from quadruplicate experiments performed in triplicate. $p$ values were determined by student’s two-tailed unpaired parametric $t$-test relative to untreated control. (** $p \leq 0.01$, *** $p \leq 0.001$ and ns indicate not significant).

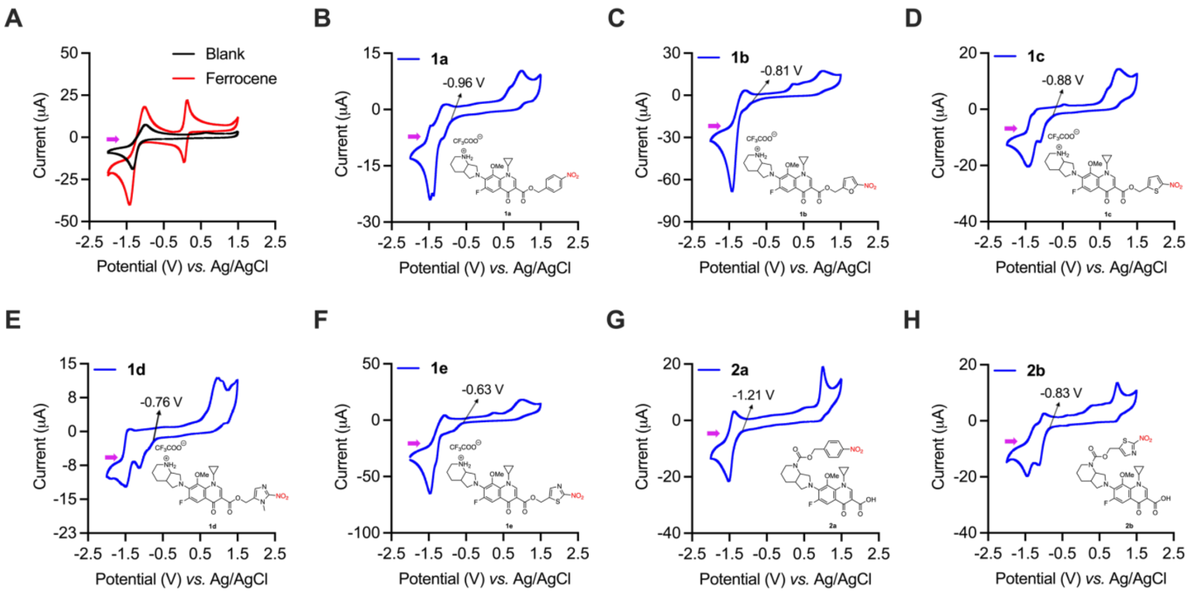

**Figure S3.** Cyclic voltammograms of (A) controls (blank electrolyte and ferrocene), (B) **1a**, (C) **1b**, (D) **1c**, (E) **1d**, (F) **1e**, (G) **2a**, and (H) **2b** recorded in argon saturated 0.1 M TBAP solution in ACN with an initial positive scan (shown by a purple-coloured arrow) at a sweep rate of 100 mV/s, sample interval of 1 mV, quiet time of 2 s and a sensitivity of 1 × $10^{-5}$ A/V. The final concentration of the analytes were 0.5 mM. The onset reduction potentials ($E^{\circ}_{red}$ Expt.) are shown with arrows in the voltammograms.

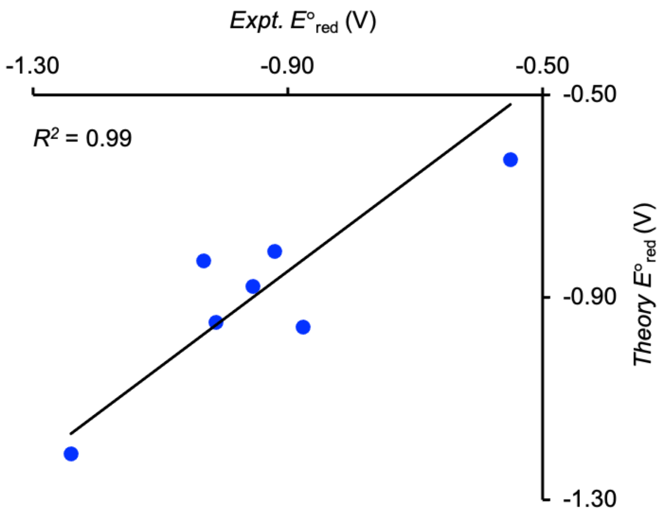


**Figure S4.** Correlation plot for experimental reduction potentials against computed reduction potentials at WB97XD/aug-cc-pVDZ//B3LYP/6-31G(d) in acetonitrile.

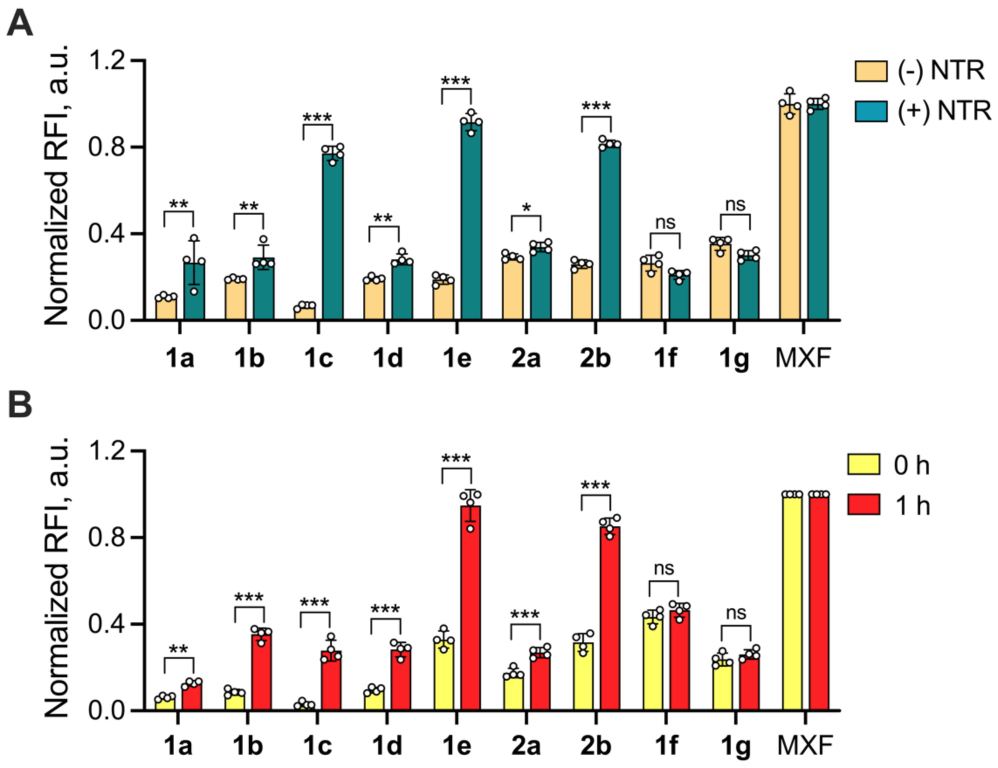


**Figure S5.** Monitoring the bioreductive activation of prodrugs by fluorescence ($\lambda_{ex}$ = 289 nm and $\lambda_{em}$ = 488 nm) after 1 h of incubation with (A) *E. coli* NTR (15 nM) and NADH (100 µM) in pH 7.4 phosphate buffer (10 mM) and (B) *M. smegmatis* whole cell lysate (1 mg/mL) at 37 ºC. All the data represent mean ± SD from quadruplicate independent experiments performed in triplicate. *p* values were determined using the student's two-tailed unpaired parametric *t*-test relative to the NTR untreated control and 0 h respectively. (* $p< 0.05$, ** $p \leq 0.01$,*** $p \leq 0.001$ and ns indicate not significant).

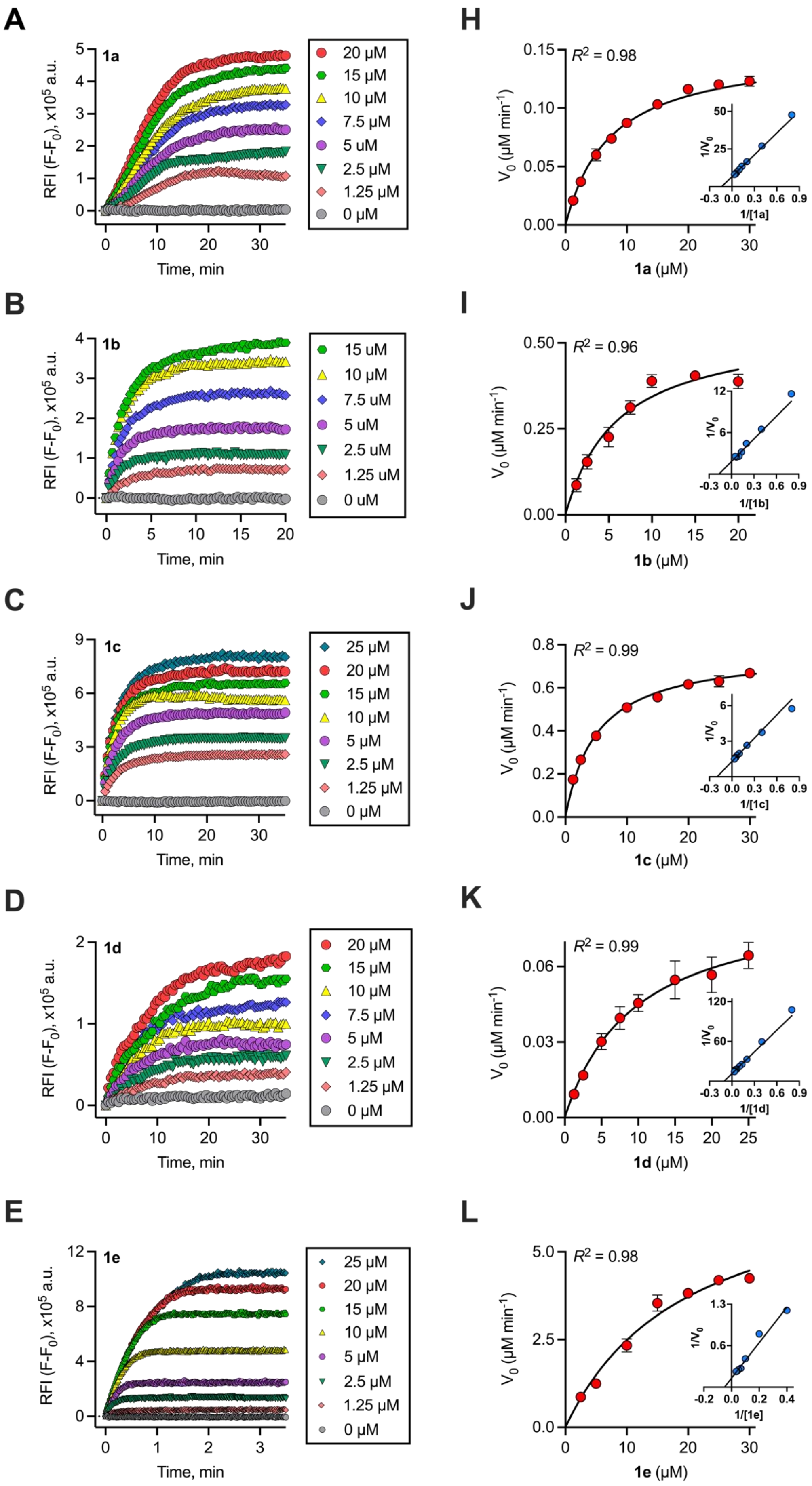
A
1a
RFI (F-F0), x10^5 a.u.
Time, min
20 µM
15 µM
10 µM
7.5 µM
5 uM
2.5 µM
1.25 µM
0 µM
B
1b
15 uM
10 uM
7.5 uM
5 uM
2.5 uM
1.25 uM
0 uM
C
1c
25 µM
20 µM
15 µM
10 µM
5 µM
2.5 µM
1.25 µM
0 µM
D
1d
20 µM
15 µM
10 µM
7.5 µM
5 µM
2.5 µM
1.25 µM
0 µM
E
1e
25 µM
20 µM
15 µM
10 µM
5 µM
2.5 µM
1.25 µM
0 µM
H
R2 = 0.98
V0 (µM min-1)
1a (µM)
1/V0
1/[1a]
I
R2 = 0.96
1b (µM)
1/[1b]
J
R2 = 0.99
1c (µM)
1/[1c]
K
R2 = 0.99
1d (µM)
1/[1d]
L
R2 = 0.98
1e (µM)
1/[1e]

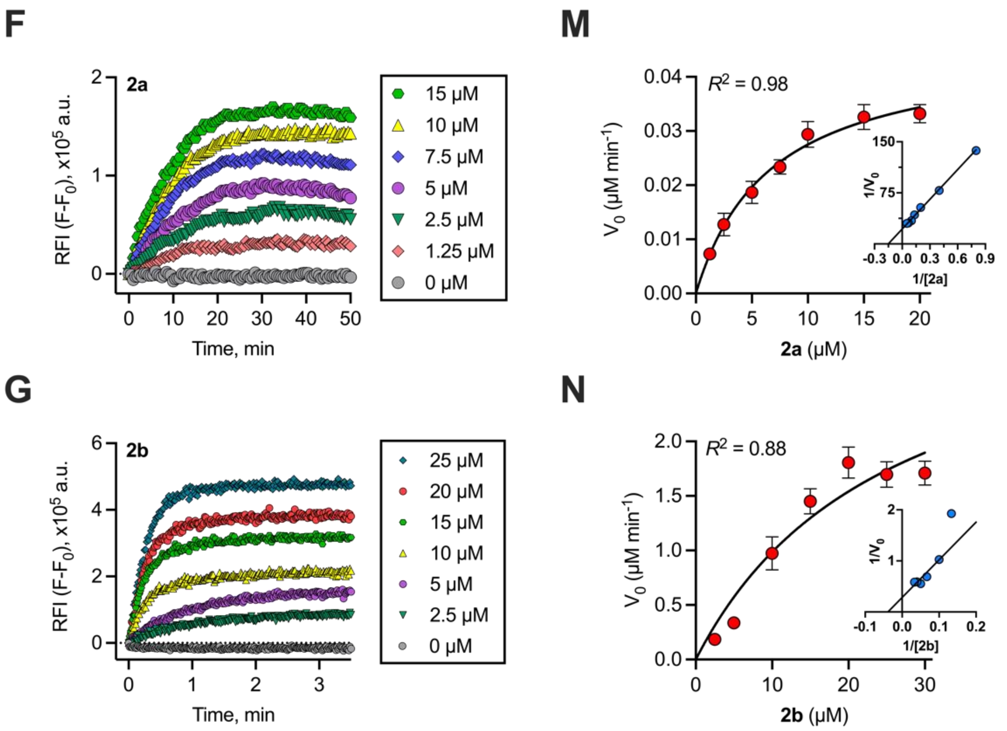


**Figure S6.** Determination of enzymatic kinetic parameters for nitro-heterocyclic ester and carbamate derivatives of MXF with *E.coli* NTR.

(A-F) Representative reaction progress curves obtained by monitoring the release of MXF ($\lambda_{ex}$ = 289 nm and $\lambda_{em}$ = 488 nm) as a function of time from (A) **1a**, (B) **1b**, (C) **1c**,(D) **1d**, (E) **1e,** (F) **2a** and (G) **2b** using a fluorescence-based assay. The reactions were performed with a broad range of substrate concentrations (reported on right of each curve) in the presence of NADH (100 µM) and a fixed concentration of *E.coli* NTR (15 nM) in phosphate buffer (pH 7.4, 10 mM). The kinetic experiments were performed at either 37 °C (for **1a-1d** and **2a**)or 25 °C(for **2b**).(H-N) Michaelis-Menten plots of NTR-catalyzed nitro-reduction and release of MXF from substrates: (H) **1a**, (I) **1b**, (J) **1c**, (K) **1d**, (L) **1e**, (M) **2a** and (N) **2b**. Each concentration represents rates of formation of MXF calculated from three biological replicates from a linear regression analysis. The data are plotted as mean ± SD for each concentration, and the lines connecting the points represent a non-linear curve fit ($R^2$ ~ 0.88-0.99) to a classical Michaelis-Menten enzyme kinetics equation. *inset*: The same data is represented as Lineweaver-Burk double reciprocal plots for all the substrates.

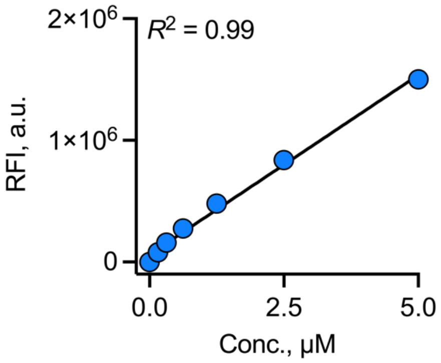


**Figure S7.** Calibration curve for MXF in pH 7.4 phosphate buffer (10 mM) at 37 ºC.

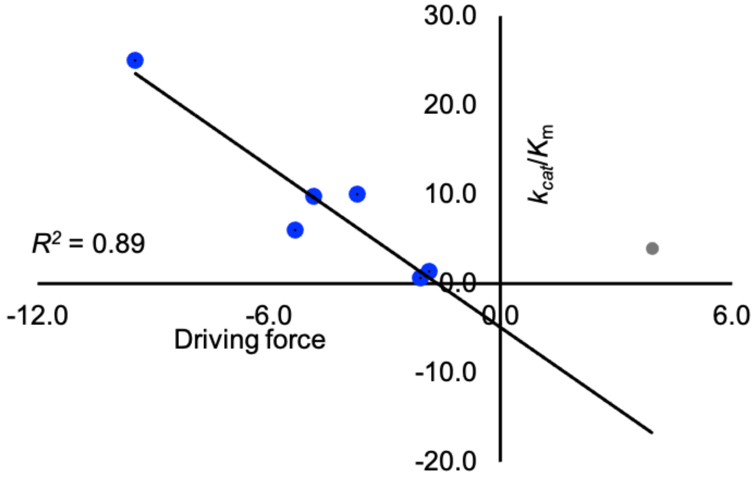


**Figure S8.** Correlation plot for driving force for electron transfer against and $k_{cat}/K_m$. Point in grey is excluded in the regression analysis.

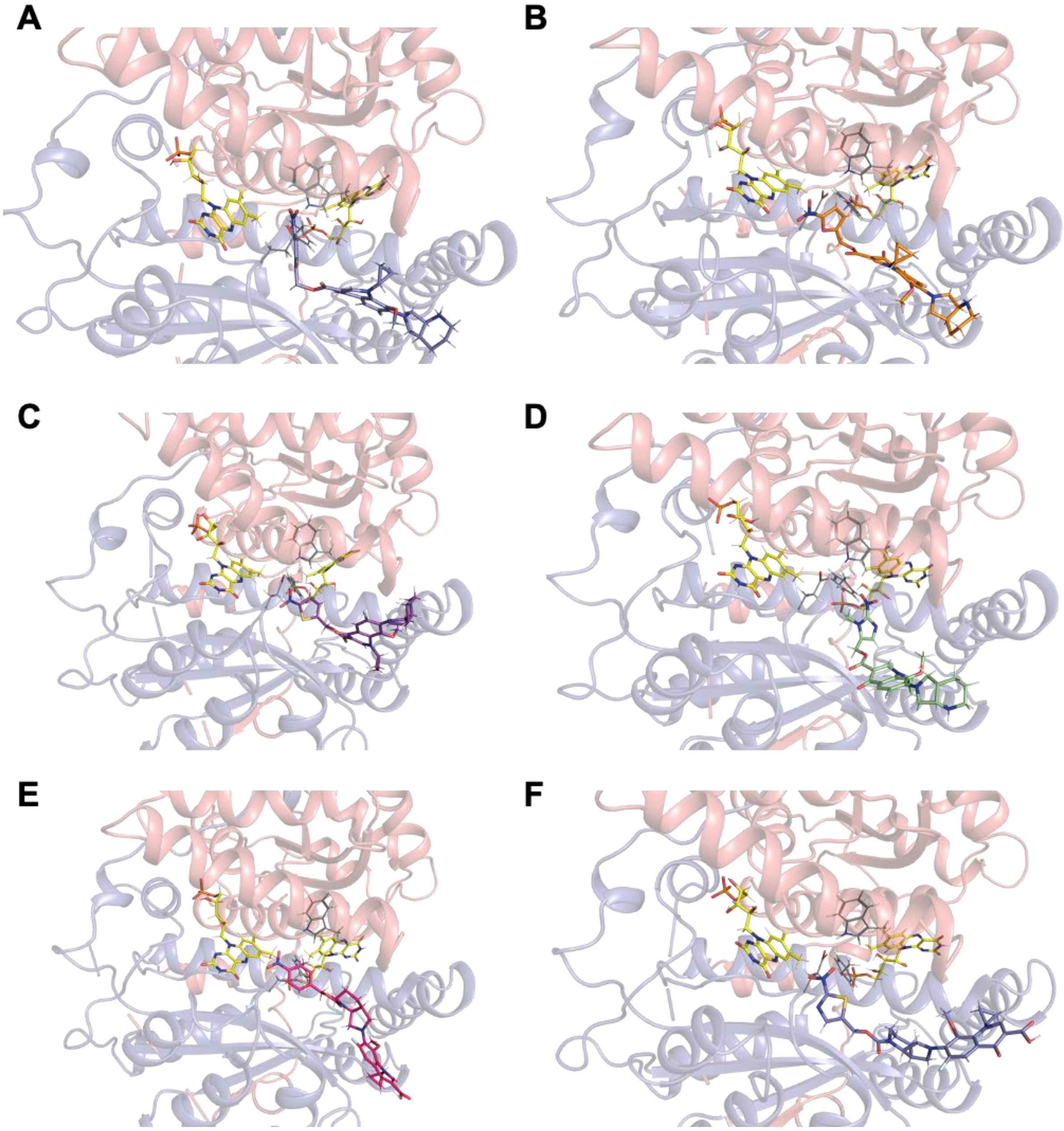


**Figure S9.** Docking based MD simulation of prodrugs (A) **1a**, (B) **1b**, (C) **1c**, (D) **1d**, (E) **2a** and (F) **2b** in *E. coli* NTR (PDB: 1DS7).

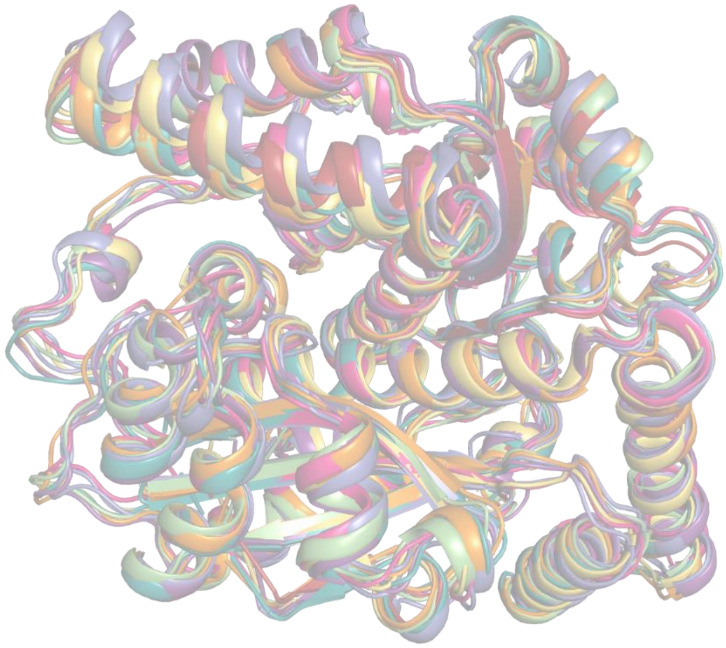

**Figure S10.** An overlay of the proteins obtained after 10 ns MD simulation for prodrugs docked in 1DS7.

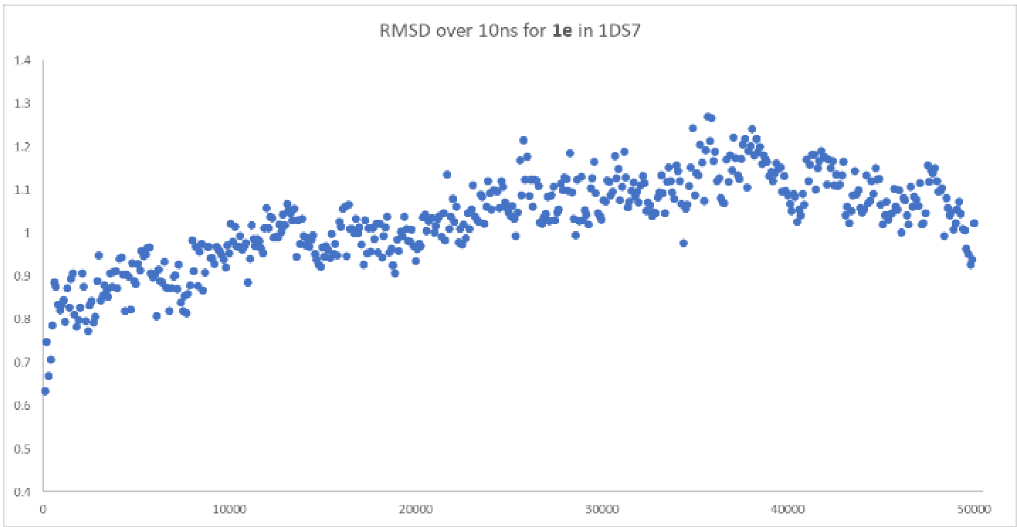


**Figure S11.** RMSD with respect to the equilibrated structure over a 10 ns simulation for **1e** bound to 1DS7.

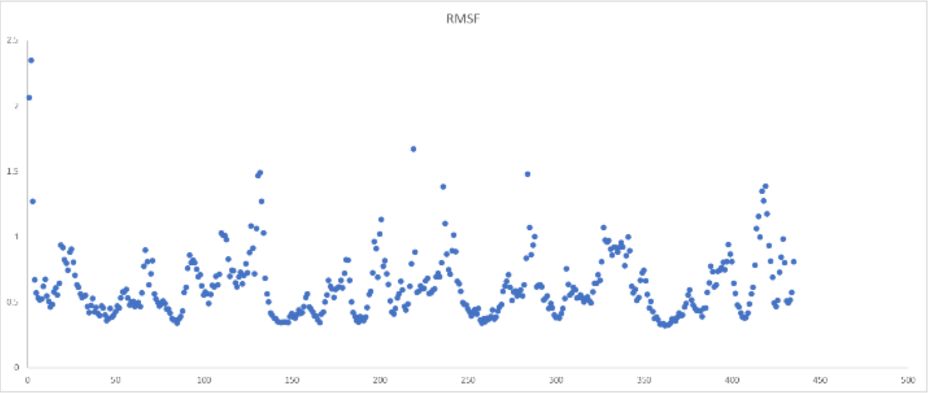


**Figure S12.** RMSF with respect to the equilibrated structure over a 10 ns simulation for **1e** bound to 1DS7.

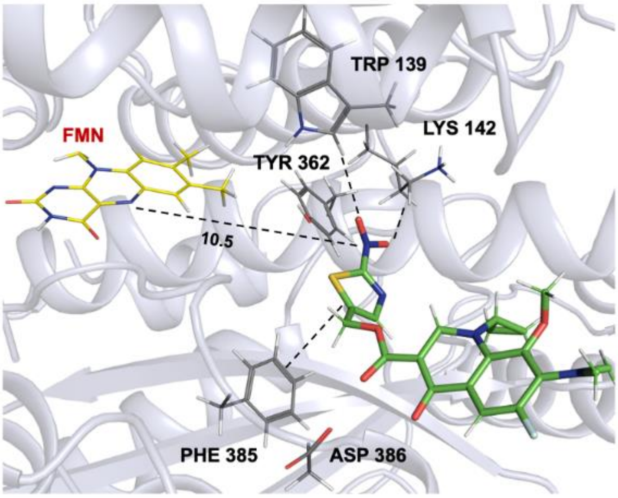


**Figure S13.** Key active site interactions for the binding of prodrug **1e** in *E. coli* NTR (PDB: 1DS7).

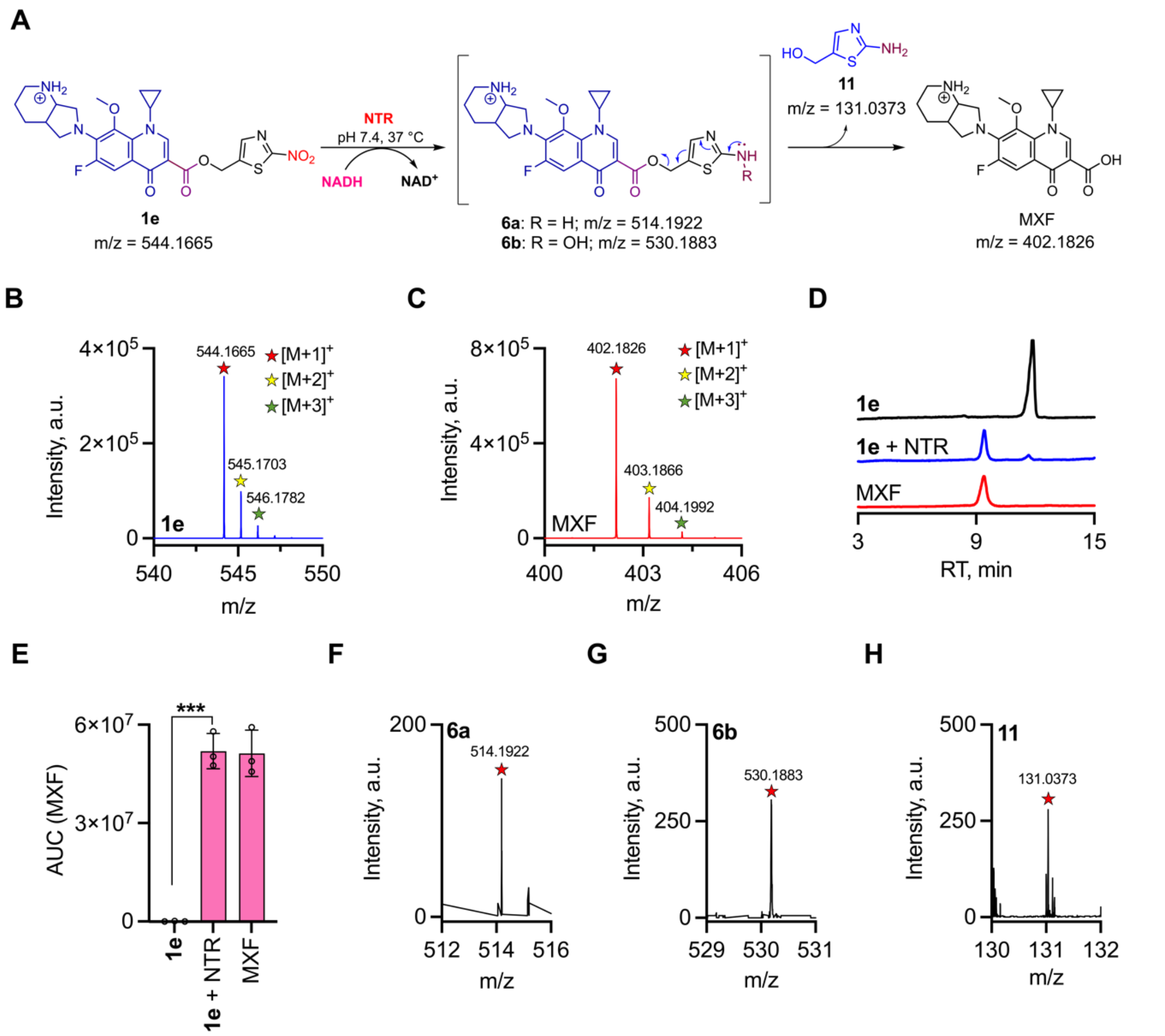

**Figure S14.** *E. coli* NTR mediated nitro-reduction of **1e** resulted in the release of MXF. (A) Schematic of generation of MXF and other reaction products from **1e** in the presence of NADH (100 µM) and *E. coli* NTR (15 nM) in phosphate buffer (10 mM) at 37 ºC. (B-C) Positive-ion-mode mass spectrum of the protonated singly $[M+1]^+$, doubly $[M+2]^+$, and triply charged $[M+3]^+$ molecular ions of authentic standards: (B) **1e** (expected, $[M+H]^+$ = 544.1661; observed, $[M+H]^+$ = m/z 544.1664) (C) MXF $(M+H]^+$, expected = m/z 402.1824; observed = m/z 402.1826). (D) LC-MS analysis showing the representative total ion chromatograms (TICs) corresponding to the generation of MXF (eluted at 11.8 min) upon reacting **1e** (eluted at 9.4 min) with NTR (15 nM) and NADH (100 µM) in PB (pH 7.4, 10 mM). (E) Area under the curve (AUC) for the peaks corresponding to formation of MXF after incubation of **1e** in the presence of *E. coli* NTR. During the enzymatic reaction, a quantitative conversion of **1e** to MXF (~98% yield) was observed. All data presented as means $\pm$ SD of peak areas for extracted ion chromatograms from triplicate experiments. *p* value was determined by unpaired two-tailed student's *t*-test analyzed relative to **1e** alone. (***; $p \leq$ 0.001). (F-H) Positive-ion-mode mass spectrum of the protonated singly charged $[M+1]^+$ molecular ion of the reaction products (F) **6a**, (G) **6b** and (H) **11** formed in the enzymatic reaction of **1e** with *E. coli* NTR.

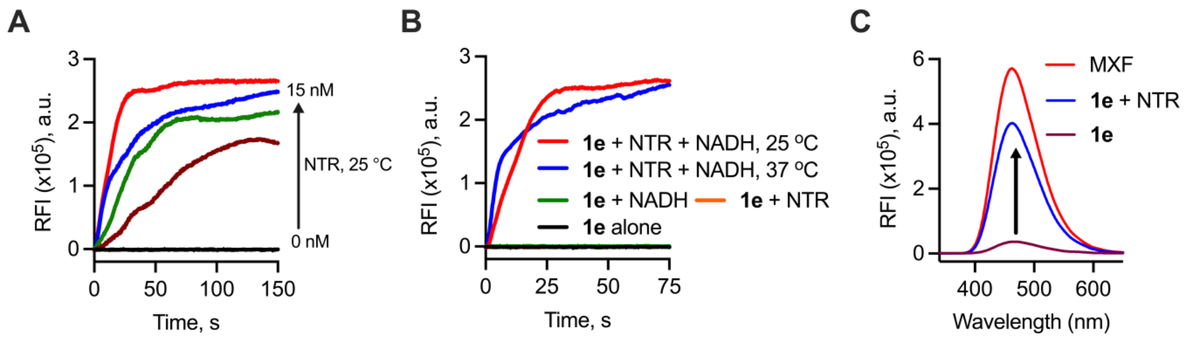


**Figure S15.** Time-dependent fluorimetric analysis of (A) reaction mixtures containing **1e** with or without NTR at different concentrations (0, 1.875, 3.75, 7.5 and 15 nM) in the presence of NADH (100 µM) in phosphate buffer (pH 7.4, 10 mM) at 25 ºC and (B) reaction mixtures containing **1e** with a fixed concentration of NTR (15 nM) at 25 ºC or 37 ºC in the presence of NADH (100 µM). The generation of MXF in the reaction mixtures were monitored at $\lambda_{ex}$ = 289 nm and $\lambda_{em}$ = 488 nm. The data was smoothed using second order with twenty neighbouring points in GraphPad Prism 9. (C) Fluorescence spectra before and after incubation of **1e** (10 µM) with *E. coli* NTR (15 nM) and NADH (100 µM) in PB (pH 7.4, 10 mM) at 37 ºC.

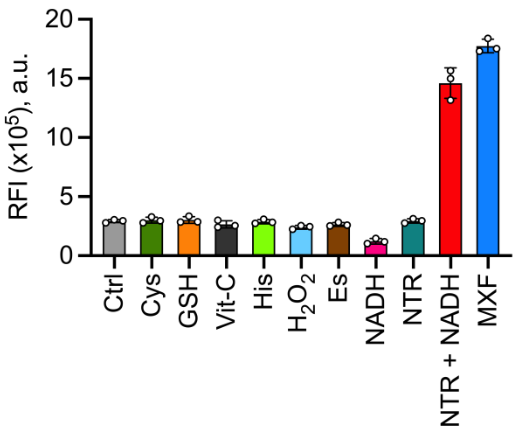


**Figure S16.** Fluorescence response of **1e** (10 µM) to various biological analytes (500 µM) in phosphate buffer (pH 7.4, 10 mM) after 15 min of incubation at 37 ºC. Ctrl = **1e** alone; Cys = cysteine; GSH = glutathione; Vit-C = ascorbic acid; His = histidine; $H_2O_2$ = hydrogen peroxide; Es = esterase 1 U/mL; NADH = reduced nicotinamide adenine dinucleotide; NTR = *E. coli* nitroreductase (15 nM). All data presented as means ± SD from three independent experiments performed in triplicate. The prodrug **1e** uniquely responded to *E. coli* NTR in the presence of NADH and produced an intense fluorescence response.

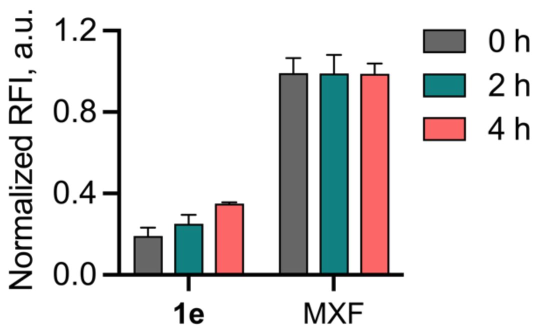


**Figure S17.** Evaluation of the stability of **1e** in lysates (1 mg/mL) of mammalian cell line (MEF; mouse embryonic fibroblast) by measuring fluorescence ($\lambda_{ex}$ = 289 nm and $\lambda_{em}$ = 488 nm) corresponding to MXF.

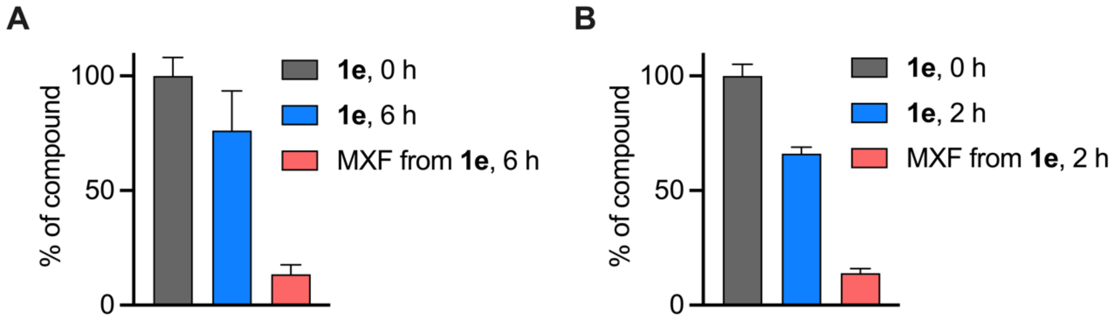


**Figure S18.** Monitoring the stability of **1e** (30 µM) by LC-MS upon incubation in human blood (A) plasma after 6 h and (B) serum after 2 h. The prodrug was recovered in an excellent yield (>75% in plasma and >65% in serum) along with the generation of MXF (13 + 4%)

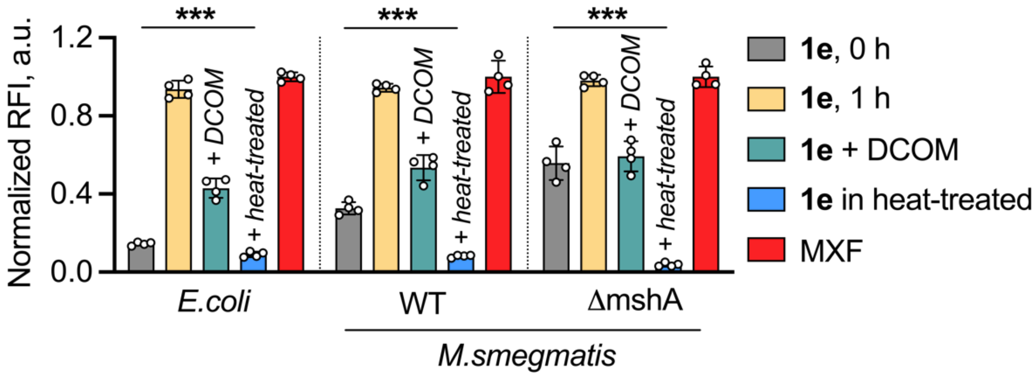


**Figure S19.** Monitoring the release of MXF from **1e** (10 µM) by measuring enhancement in fluorescence in lysates (1 mg/mL) of *E. coli* ATCC 25922, wild-type (WT)and MshA deficient (ΔmshA) *M. smegmatis* (mc$^2$155) alone or in the presence of NTR inhibitor, dicoumarol (DCOM; 100 µM) and heat-inactivated lysates.

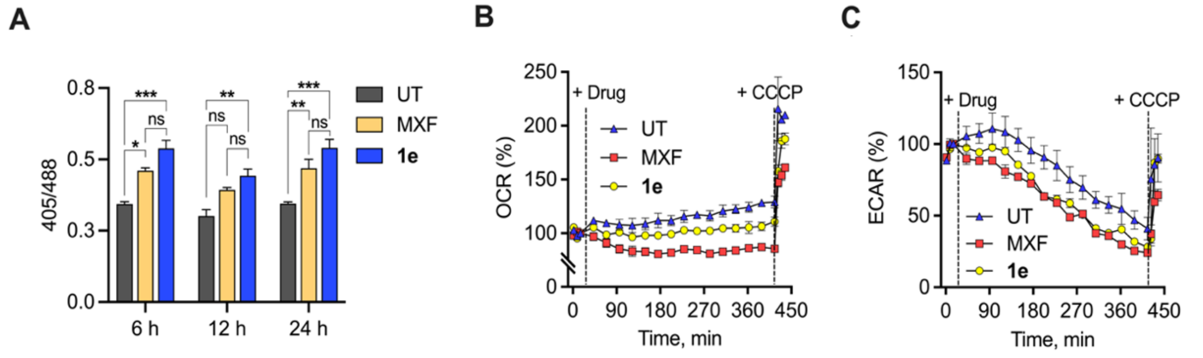


**Figure S20.** (A) Ratiometric biosensor response following exposure of *Mtb*-roGFP2 to 10× MICs of MXF and **1e** at different time intervals. Data in are represented as mean ± SD, $n = 3$ independent experiments (ns indicate not significant, * $p \leq 0.05$, ** $p \leq 0.01$, *** $p \leq 0.001$ and are determined against UT). Determination of (B) OCR (pmol/min) and (C) ECAR (mpH/min) shown as percentage of baseline values of replicating *Mtb* left untreated (UT) or treated with drug (MXF or **1e**, 10× MIC) or CCCP (10 µM) added at indicated times shown by dotted lines.

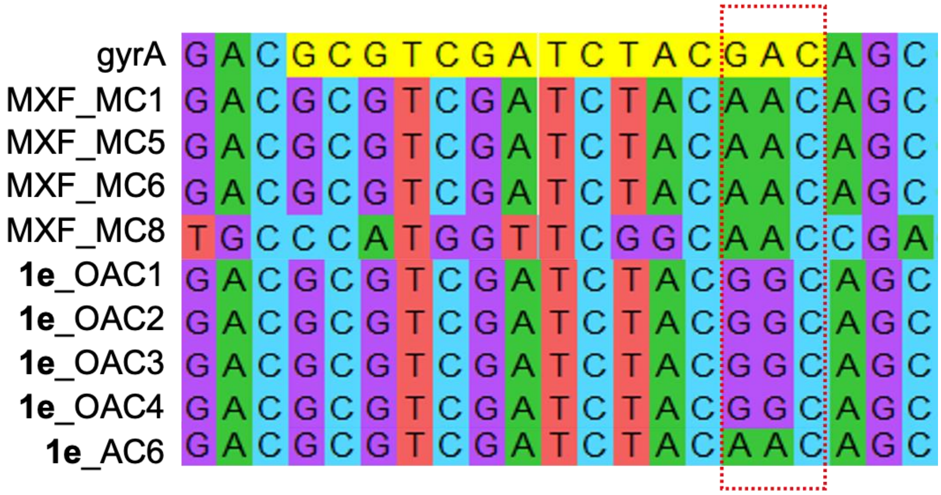


**Figure S21.** Sequences of PCR amplified quinolone resistance determinant region (QRDR) using genomic DNA in gyrA of WT and resistant isolates from MXF (MC1, MC5, MC6 and MC8) and **1e** (OAC1-4 and AC6) mutant prevention concentration (MPC) plates. The dotted box highlights the mutation from GAC (aspartic acid) to AAC (asparagine) in resistant colonies of MXF (MC1) and **1e** (OAC1) as well as to GGC (glycine) in resistant colony of **1e** (AC6).

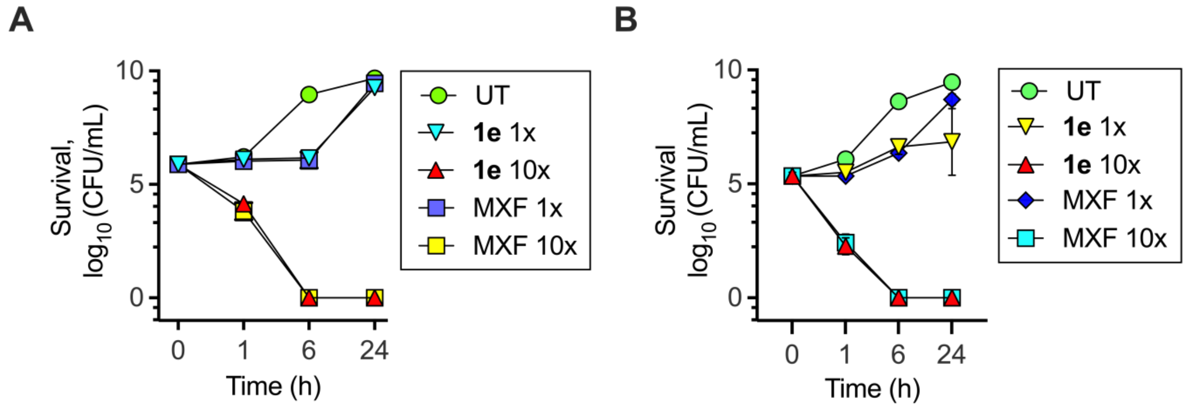


**Figure S22.** Time-kill curves for (A) *S. aureus* ATCC 29213 (~$10^5$ CFU/mL) and (B) *E. coli* ATCC 25922 (~$10^5$ CFU/mL) and treated with 1× and 10× MIC of **1e** and MFX.

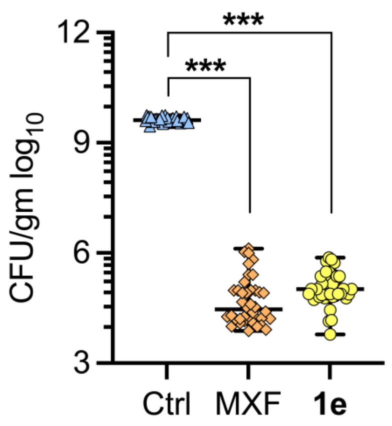


**Figure S23.** *In vivo* efficacy of **1e** (10 mg/kg) against *E. coli* ATCC 25922 in a neutropenic murine thigh infection model. MXF (10 mg/kg) was used as a reference compound while Ctrl is untreated mice. Statistical significance was established with respect to Ctrl (***$p < 0.001$).

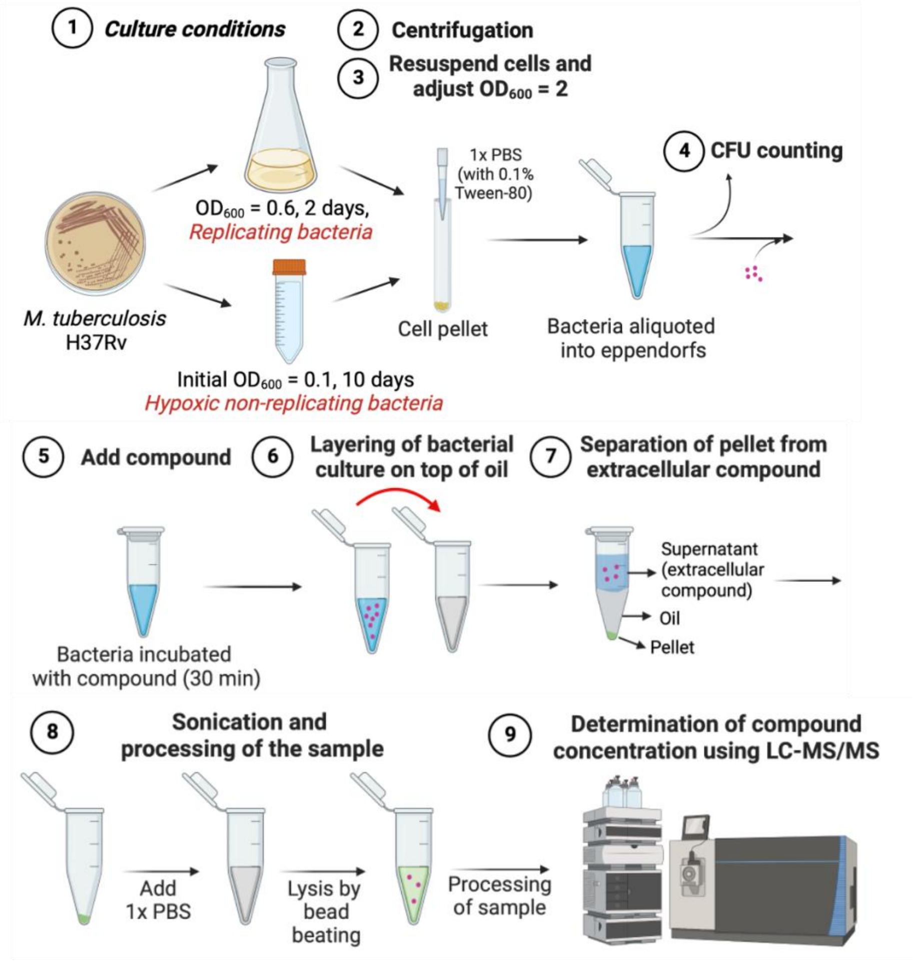


**Figure S24.** Schematic workflow for LC-MS based accumulation assay in replicating and non-replicating *Mtb*.

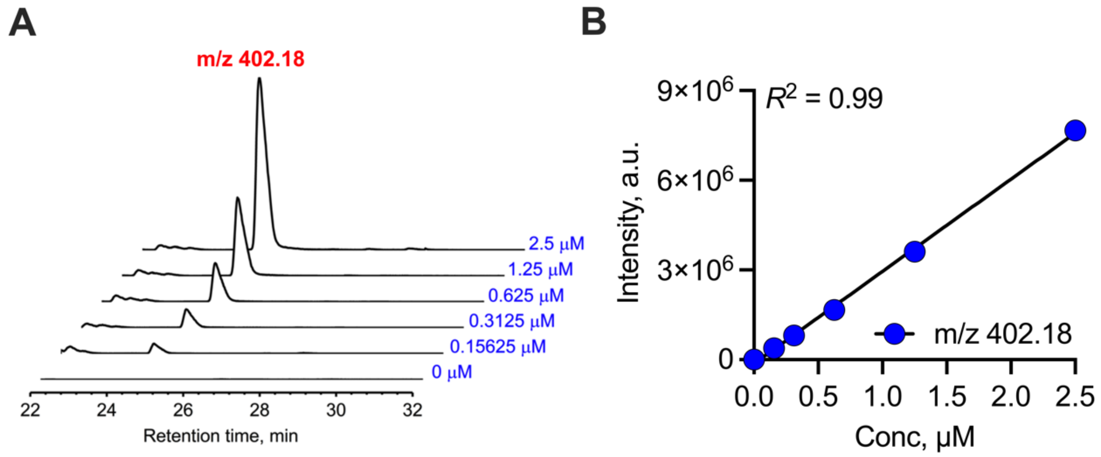


**Figure S25.** (A) Representative extracted ion chromatograms (EIC) and (B) standard calibration curve obtained with different concentrations of MXF (m/z 402.18).

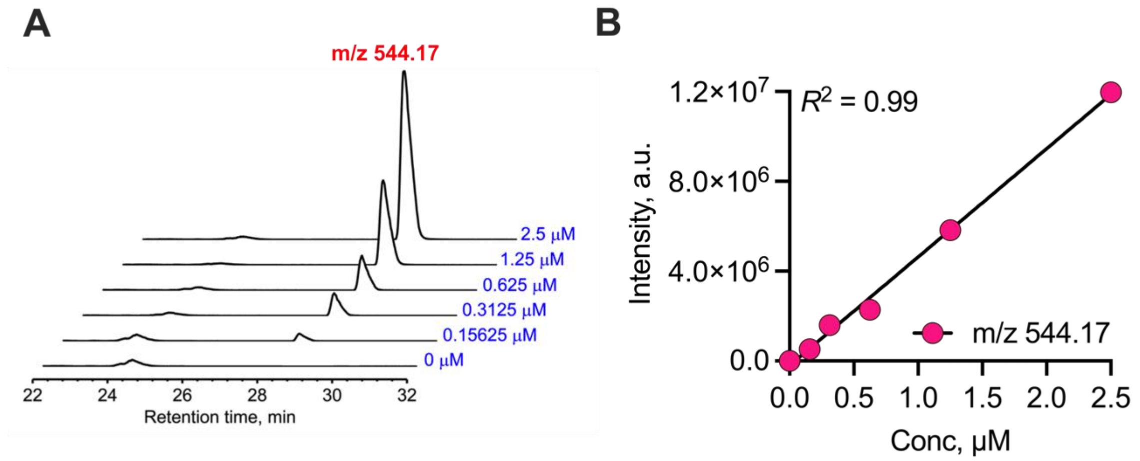


**Figure S26.** (A) Representative extracted ion chromatograms (EIC) and (B) standard calibration curve obtained with different concentrations of **1e** (m/z 544.17).

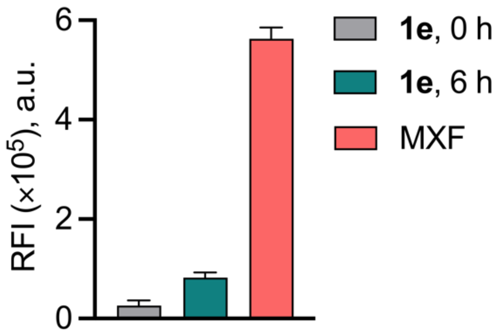


**Figure S27.** Monitoring the stability of **1e** (10 µM) by change in fluorescence in supernatant of *M. smegmatis (mc$^2$155)* at 0 h and 6 h.

#### 5. Tables

**Table S1. Synthesis of *N*-boc protected esters of MXF**

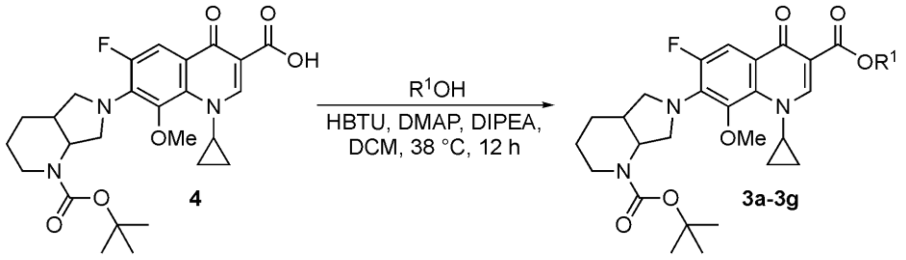


| Entry | $R^1$ | $R^1OH$ | Product | Yield, % |
|---|---|---|---|---|
| 1 | 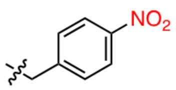 | **6a** | **3a** | 73 |
| 2 | 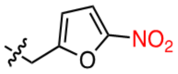 | **6b** | **3b** | 61 |
| 3 | 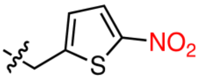 | **6c** | **3c** | 60 |
| 4 | 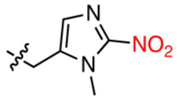 | **6d** | **3d** | 81 |
| 5 | 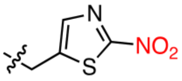 | **6e** | **3e** | 78 |
| 6 | 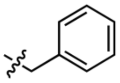 | **6f** | **3f** | 65 |
| 7 | 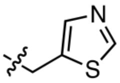 | **6g** | **3g** | 55 |

**Table S2. Synthesis of *N*-boc deprotected esters of MXF**

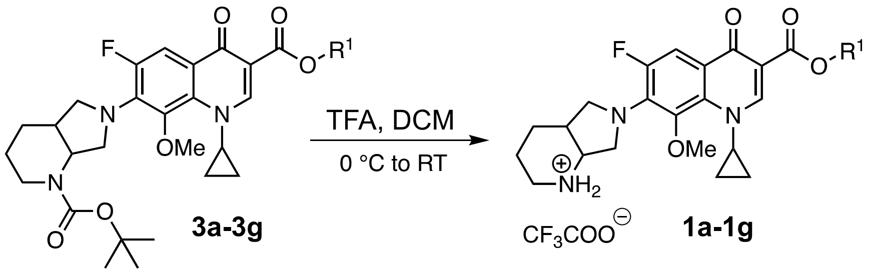


| Entry | R | Product | Yield, % |
|---|---|---|---|
| 1 | 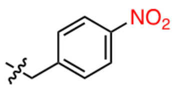 | **1a** | 83 |
| 2 | 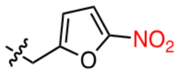 | **1b** | 85 |
| 3 | 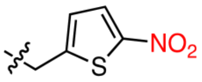 | **1c** | 87 |
| 4 | 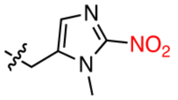 | **1d** | 95 |
| 5 | 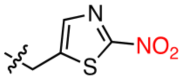 | **1e** | 95 |
| 6 | 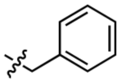 | **1f** | 85 |
| 7 | 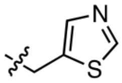 | **1g** | 83 |

**Table S3.** Physicochemical properties and reduction potentials of compounds

| Cpd | clogP[a] | MR[b] | TPSA[c] |
|---|---|---|---|
| **1a** | 3.0 | 164.3 | 163.3 |
| **1b** | 2.0 | 156.6 | 176.4 |
| **1c** | 2.5 | 162.2 | 191.5 |
| **1d** | 0.7 | 161.4 | 181.1 |
| **1e** | 1.0 | 160.0 | 204.4 |
| **2a** | 4.4 | 158.7 | 147.1 |
| **2b** | 2.6 | 154.4 | 188.2 |
| MXF | -0.49 | 120.8 | 88.3 |

[a]clogP values were obtained using ChemDraw 19.1 for prodrugs/drug; [b]MR = molar refractivity (represents the molar volume of the molecule); [c]TPSA = topological polar surface area (Represents the surface sum over all polar atoms or molecules). MR and TPSA were calculated from SwissADME webservice.[23]

**Table S4.** Comparative analysis of kinetic parameters of NTR-MXF prodrugs

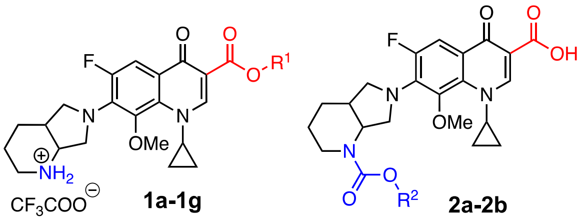


| Entry | Prodrug | R | Initial rate V ($\mu M\ min^{-1}$) | $k_{cat}$ ($min^{-1}$) | $K_m$ ($\mu M$) | $k_{cat}/K_m$ ($\mu M^{-1}\ min^{-1}$) | Relative rate* |
|---|---|---|---|---|---|---|---|
| 1 | **1a** | 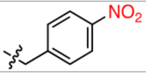 | 0.15 ± 0.005 | 10.0 | 7.28 ± 0.75 | 1.37 | 2 |
| 2 | **1b** | 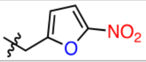 | 0.54 ± 0.056 | 36.6 | 5.98 ± 1.59 | 6.11 | 7 |
| 3 | **1c** | 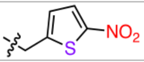 | 0.77 ± 0.016 | 51.6 | 5.08 ± 0.39 | 10.16 | 10 |
| 4 | **1d** | 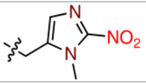 | 0.08 ± 0.003 | 5.85 | 9.59 ± 0.87 | 0.61 | 1 |
| 5 | **1e** | 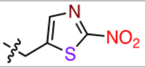 | 7.41 ± 0.98 | 494.3 | 19.81 ± 5.18 | 24.95 | 93 |
| 6 | **2a** | 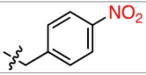 | 0.045 ± 0.002 | 3.05 | 6.66 ± 0.93 | 0.45 | 0.6 |
| 7 | **2b** | 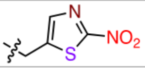 | 3.28 ± 1.14 | 218.9 | 22.5 ± 14.95 | 9.72 | 41 |

*Initial rates were recorded relative to the rate of NTR-catalyzed nitro-reduction for **1d** under the defined conditions

**Table S5.** RMSD values for different prodrugs in 1DS7

| System (prodrugs in 1DS7) | RMSD in Å |
|---|---|
| **1a** | 1.47 |
| **1b** | 1.16 |
| **1c** | 1.07 |
| **1d** | 1.10 |
| **2a** | 1.16 |
| **2b** | 1.25 |
| **1e** | Reference |

**Table S6.** Binding energies and dispersion interactions for various prodrug-enzyme complexes

| Prodrug | Full_Sys (H) | Only_Protein (H) | Prodrug (H) | Binding_HF3C (kcal/mol) | D3BJ (kcal/mol) |
|---|---|---|---|---|---|
| **1a** | -10079.66366 | -8247.750559 | -1831.852825 | -37.8 | -23.0 |
| **1b** | -10077.73807 | -8247.830973 | -1829.805004 | -64.1 | -38.7 |
| **1c** | -10398.90989 | -8247.739884 | -2150.998077 | -107.9 | -36.9 |
| **1d** | -10112.54917 | -8247.724042 | -1864.721574 | -65.0 | -23.1 |
| **1e** | -10414.72916 | -8247.797284 | -2166.835597 | -60.4 | -23.8 |
| **2a** | -10265.65324 | -8247.737849 | -2017.870752 | -28.0 | -29.2 |
| **2b** | -10600.69929 | -8247.777665 | -2352.811509 | -69.1 | -18.4 |

**Table S7.** Antimycobacterial activity of NTR-MXF prodrugs against *M. tuberculosis* H37Rv (*Mtb*)

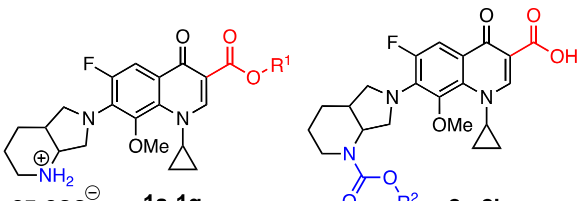


**1a-1g** **2a-2b**

| Entry | Prodrug | R | MIC (μM) |
|---|---|---|---|
| 1 | **1a** | 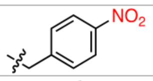  | 2 |
| 2 | **1b** | 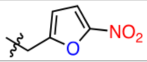  | 8 |
| 3 | **1c** | 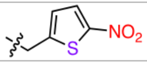  | 2-8 |
| 4 | **1d** | 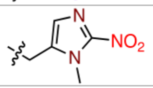  | 1 |
| 5 | **1e** | 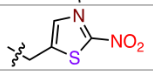  | 0.25 |
| 6 | **1f** | 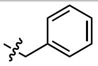 | 1 |
| 7 | **1g** | 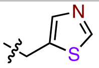  | 2-8 |
| 8 | **2a** | 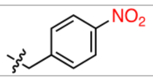  | >8 |
| 9 | **2b** | 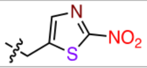  | 0.25 |

**Table S8.** Antimycobacterial activity of 2-nitrothiazole intermediates against *M. tuberculosis* H37Rv (*Mtb*)

| Entry | Compound | MIC (μM) |
|---|---|---|
| 1 | 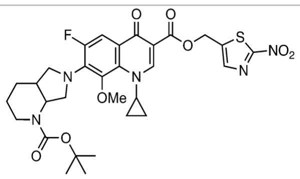  **3e** | >8 |
| 2 | 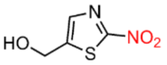  **6e** | >64 |

**Table S9.** MICs against *Mtb* resistant isolates from MXF and **1e** MPC plates

| Entry | Isolates | | MIC (μM) | |
|---|---|---|---|---|
| | | | MXF | **1e** |
| 1 | Isolates from MXF MPC plate | MC1 | 4 | 8 |
| 2 | | MC5 | 4 | 8 |
| 3 | | MC6 | 4 | 8 |
| 4 | | MC8 | 4 | 8 |
| 5 | Isolates from **1e** MPC plate | OAC1 | 8 | 8 |
| 6 | | OAC2 | 8 | 8 |
| 7 | | OAC3 | 8 | 8 |
| 8 | | OAC4 | 8 | 8 |
| 9 | | AC6 | 8 | 16 |

**Table S10.** MIC of **1e** against pathogenic bacteria

| Entry | Bacterial species | MIC (μM) | | Fold difference |
|---|---|---|---|---|
| | | **1e** | MXF | |
| 1 | *Escherichia coli* ATCC 25922 | 0.09 | 0.03 | 3 |
| 2 | *Staphylococcus aureus* ATCC 29213 | 0.09 | 0.07 | 1.2 |
| 3 | *Pseudomonas aeruginosa* ATCC 27853 | 6 | 4 | 1.5 |
| 4 | *Mycobacterium abscessus* ATCC 19977 | 1 | 1 | - |
| 5 | *Mycobacterium fortuitum* ATCC 6841 | 0.12 | 0.03 | 4 |
| 6 | *Mycobacterium chelonae* ATCC 35752 | 0.06 | 0.02 | 3 |

**Table S11.** MIC of **1e** against clinically resistant strains of *E. coli*

| **Prodrug/Drug** | **MIC (μg/mL)** | | | |
|---|---|---|---|---|
| | *E.coli* ATCC 25922 | *E.coli* NR 17661 | *E.coli* NR 17663 | *E.coli* NR 17666 |
| **1e** | 0.125 | >64 | 0.25 | 0.25 |
| MXF | 0.015 | 32 | 0.015 | 0.015 |
| CIP | 0.0075 | 64 | 0.0075 | 0.0075 |
| LVX | 0.015 | 32 | 0.015 | 0.015 |
| NFX | 0.03 | >64 | 0.06 | 0.06 |
| Meropenem | 0.03 | 0.03 | 0.03 | 0.03 |
| Ceftazidime | 0.5 | 0.5 | 0.125 | 0.125 |
| Amikacin | 0.5 | 1 | 1 | 1 |
| Clarithromycin | 64 | >64 | 16 | 64 |
| Tobramycin | 0.25 | 0.5 | 0.25 | 0.5 |
| Streptomycin | 2 | >64 | 2 | 4 |
| Minocycline | 1 | 4 | 1 | 0.5 |
| Polymixin-B | 0.125 | 0.125 | 0.125 | 0.125 |

Drug susceptible Drug resistant

**Table S12.** Antibacterial activity of **1e** against clinically resistant strains of *S. aureus*

| ***S.aureus*** | | **Prodrug/Drug, MIC (μg/mL)** | | | | | | |
|---|---|---|---|---|---|---|---|---|
| | | **1e** | MXF | LVX | DAP | MER | VAN | MET |
| **MSSA** | ATCC 29213 | 0.125 | 0.03 | 0.125 | 1 | 0.0625 | 1 | 1 |
| **VRSA** | VRS 1 | 8 | 8 | 32 | 1 | 64 | >64 | >64 |
| | VRS 4 | 16 | 64 | >64 | 0.5 | 64 | >64 | >64 |
| | VRS 12 | 16 | 4 | 32 | 0.5 | 16 | >64 | >64 |
| **MRSA** | NRS 100 | 0.25 | 0.06 | 0.25 | 1 | 32 | 2 | >64 |
| | NRS 119 | 8 | 4 | 16 | 1 | 64 | 2 | >64 |
| | NRS 129 | 0.25 | 0.06 | 0.25 | 0.5 | 4 | 1 | >64 |
| | NRS 186 | 4 | 1 | 8 | 0.5 | 4 | 1 | >64 |
| | NRS 191 | 8 | 4 | 16 | 1 | 64 | 1 | >64 |
| | NRS 192 | 2 | 1 | 8 | 0.5 | 8 | 1 | >64 |
| | NRS 193 | 8 | 4 | 32 | 1 | 64 | 2 | >64 |
| | NRS 194 | 0.125 | 0.03 | 0.125 | 0.5 | 1 | 1 | 32 |
| | NRS 198 | 16 | 4 | 32 | 1 | 64 | 2 | >64 |

Drug susceptible Drug resistant

**Table S13.** MIC of MXF and **1e** against Fluoroquinolone-resistant strains

| Entry | *E. coli* strains | MIC (μM) | |
|---|---|---|---|
| | | MXF | **1e** |
| 1 | NR 17661 | 73 | >97 |
| 2 | NR 48983 | 73 | >97 |
| 3 | NR 51487 | 73 | >97 |

#### 7. NMR spectra of compounds

$^{19}$F NMR spectrum (376 MHz, DMSO-$d_6$) of **1a**

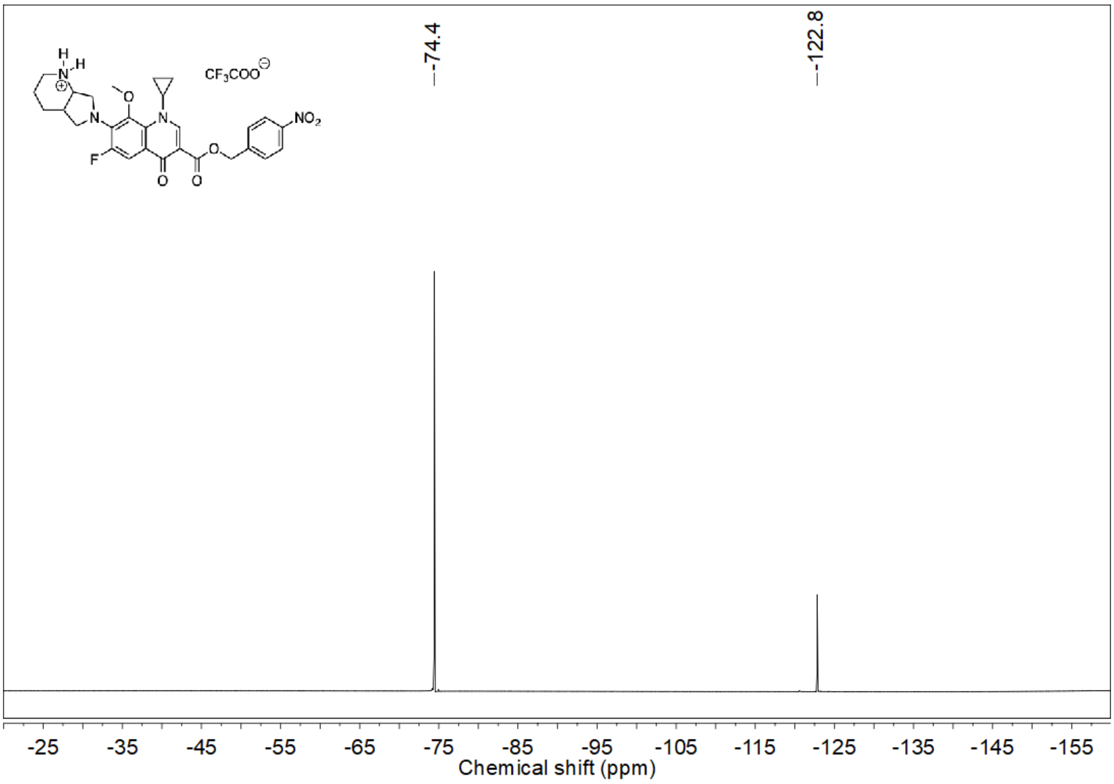


$^{1}$H NMR spectrum (400 MHz, DMSO-$d_6$) of **1a**

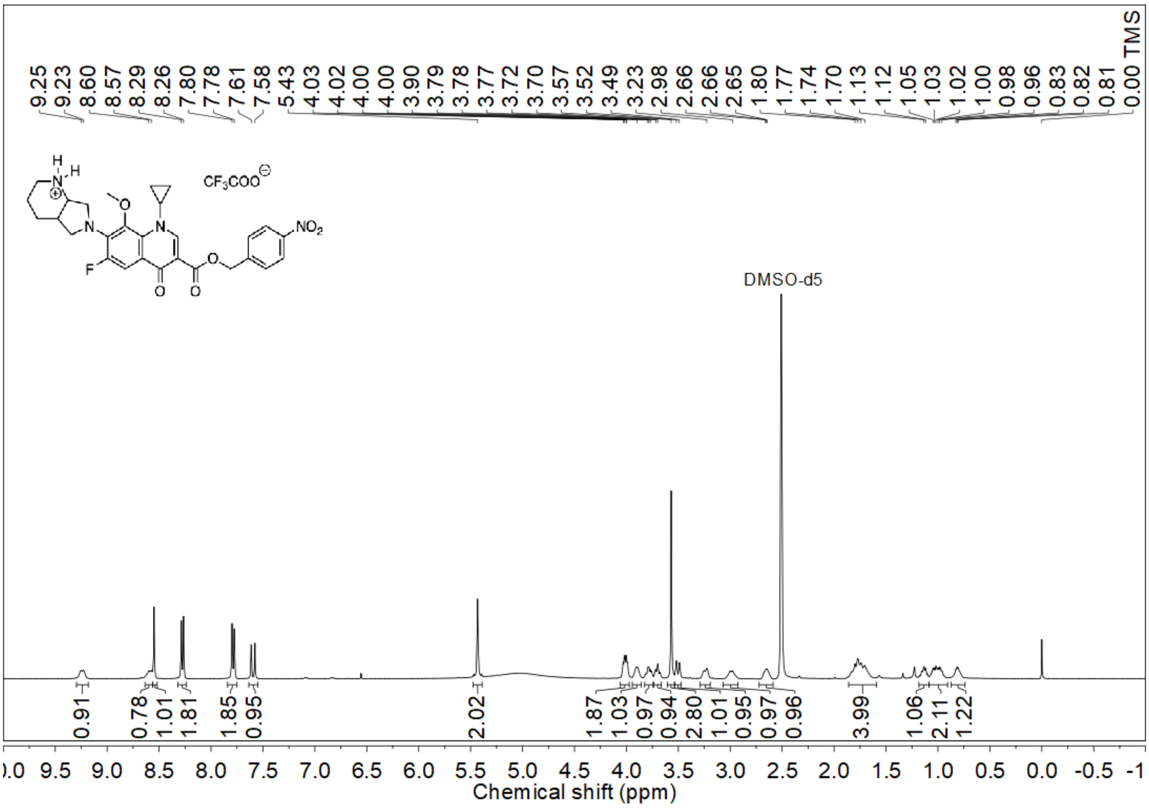

$^{13}C$ NMR spectrum (100 MHz, DMSO-$d_6$) of **1a**

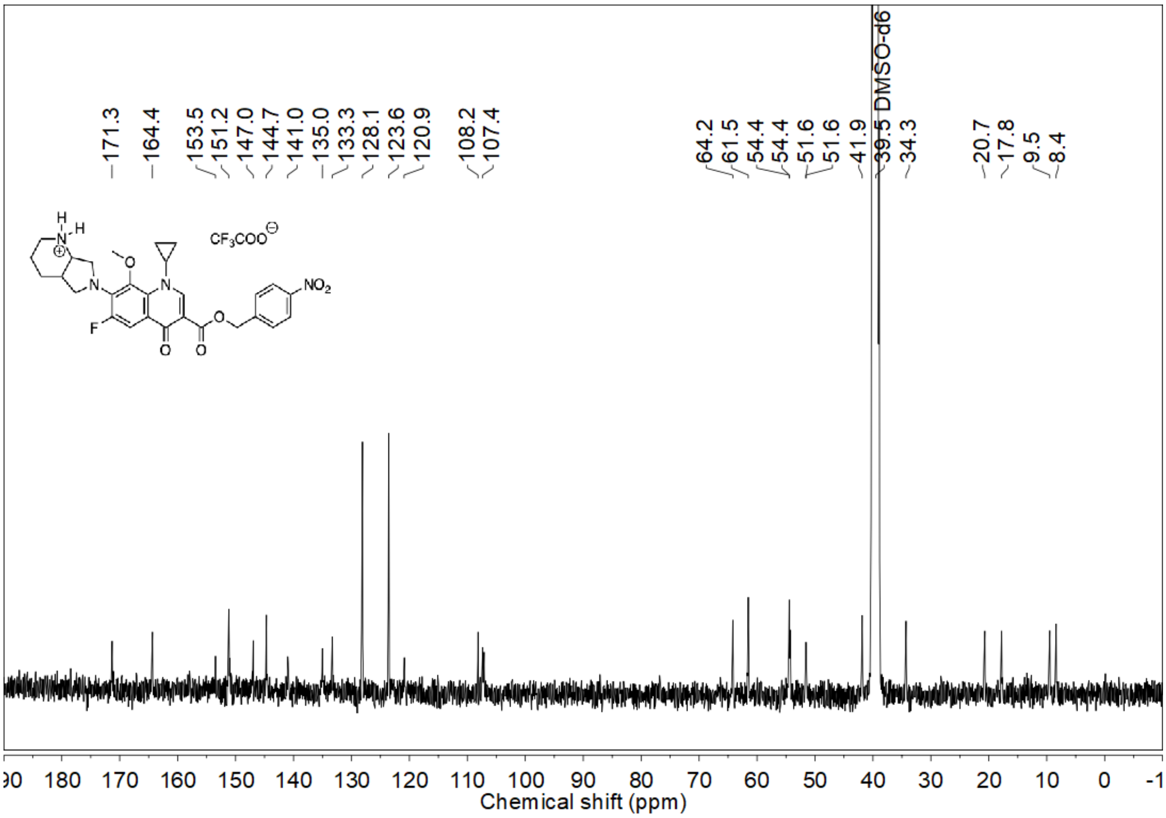


DEPT-135 NMR spectrum (100 MHz, DMSO-$d_6$) of **1a**

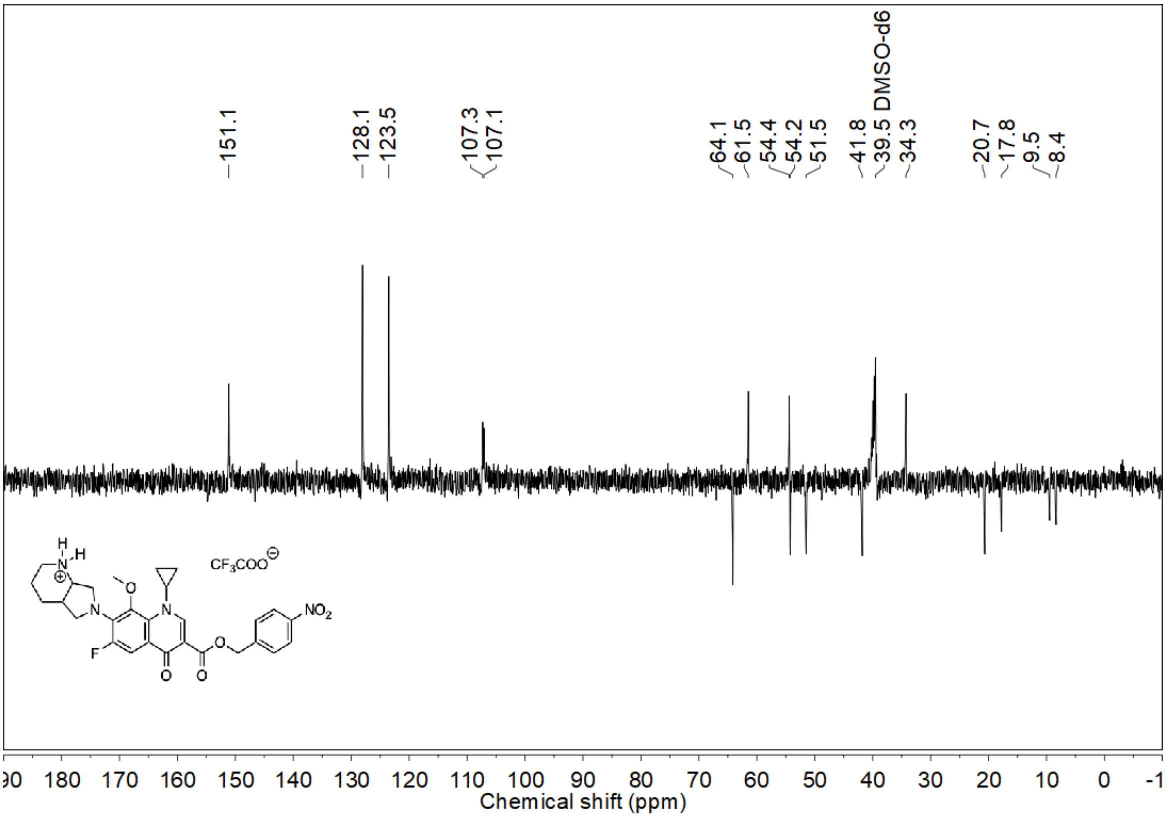

[19]F NMR spectrum (376 MHz, DMSO-$d_6$) of **1b**

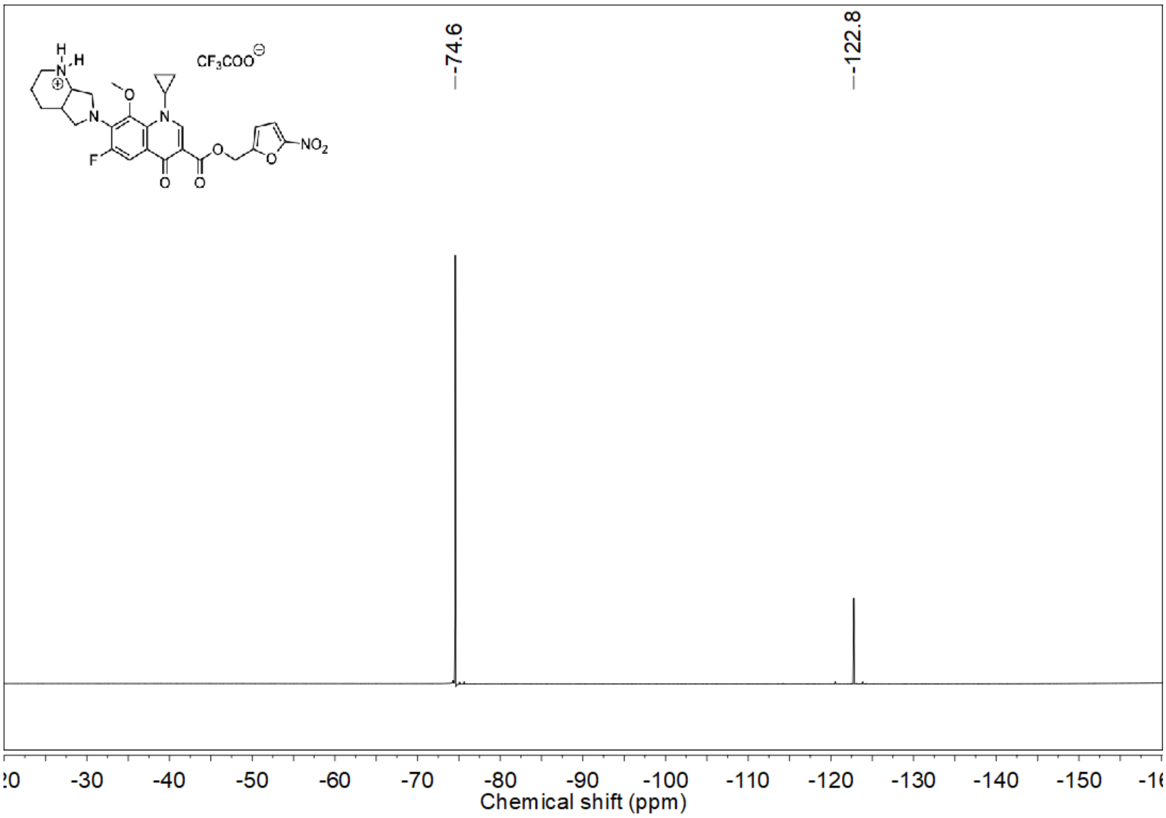


[1]H NMR spectrum (400 MHz, DMSO-$d_6$) of **1b**

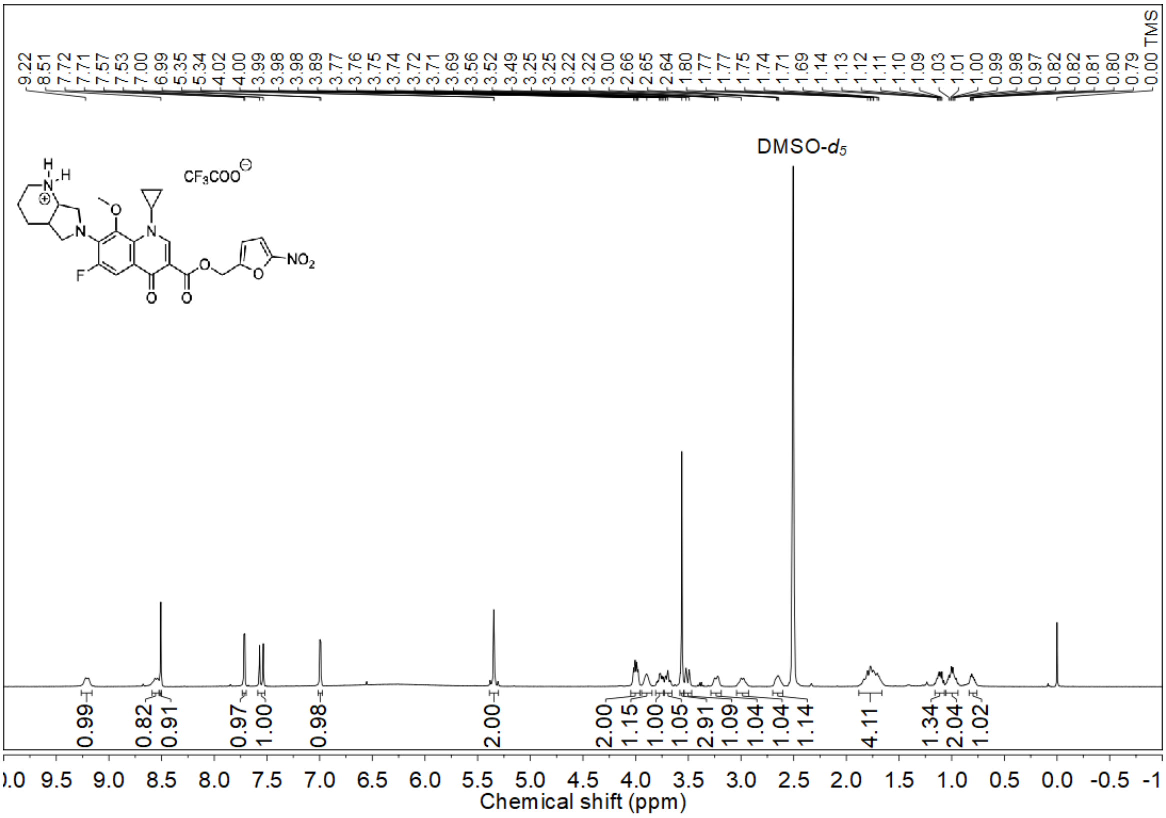

$^{13}C$ NMR spectrum (100 MHz, DMSO-$d_6$) of **1b**

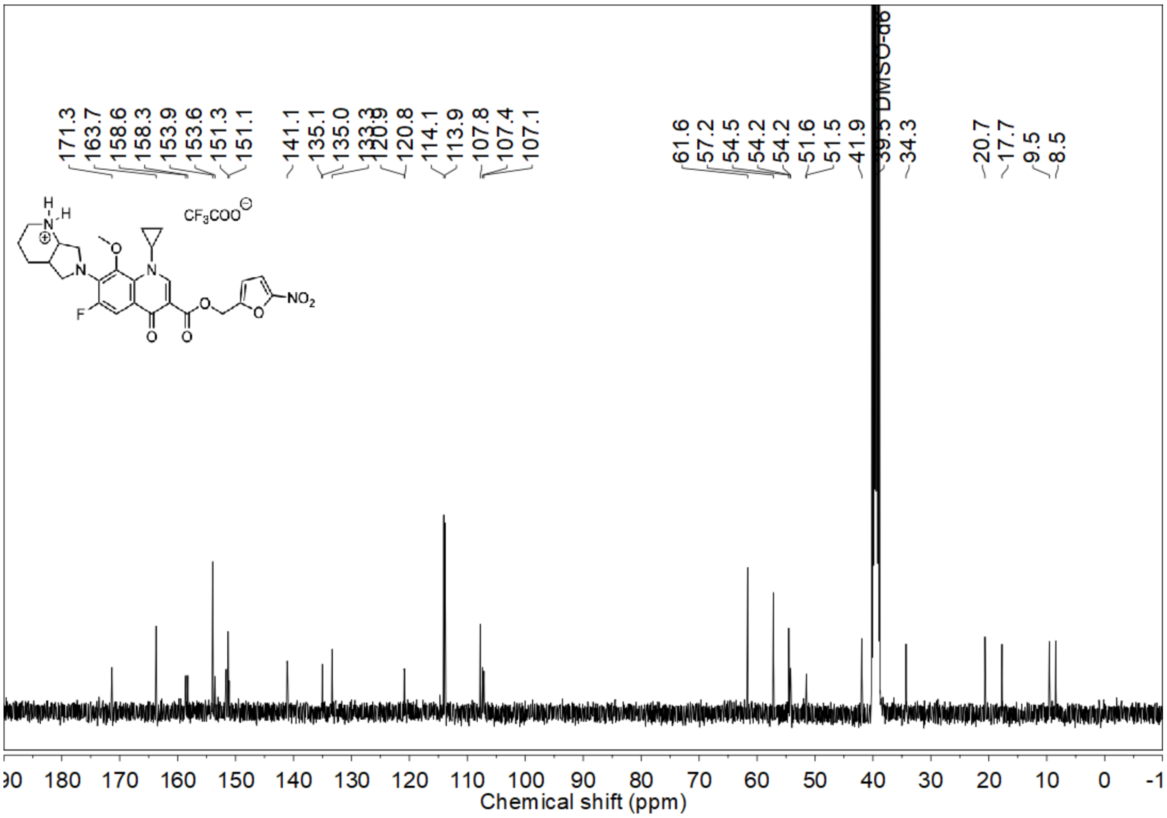


DEPT-135 NMR spectrum (100 MHz, DMSO-$d_6$) of **1b**

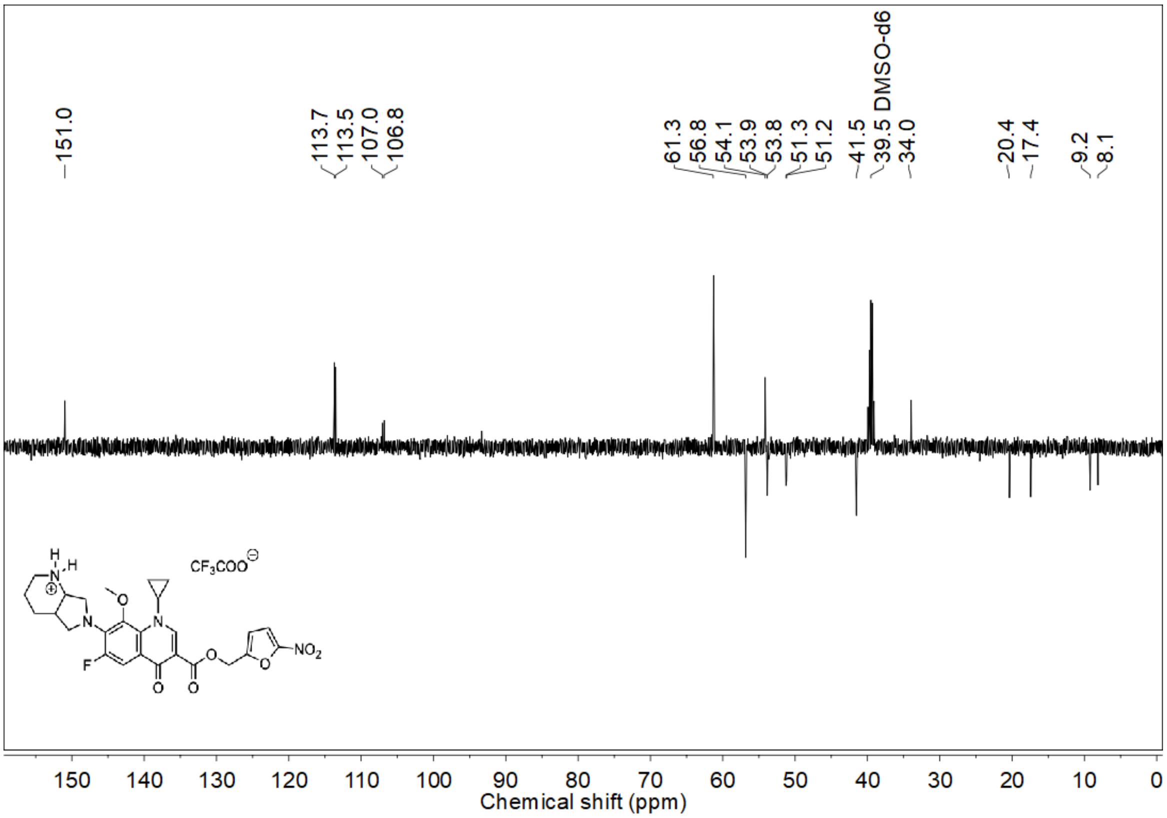

$^{19}$F NMR spectrum (376 MHz, DMSO-$d_6$) of **1c**

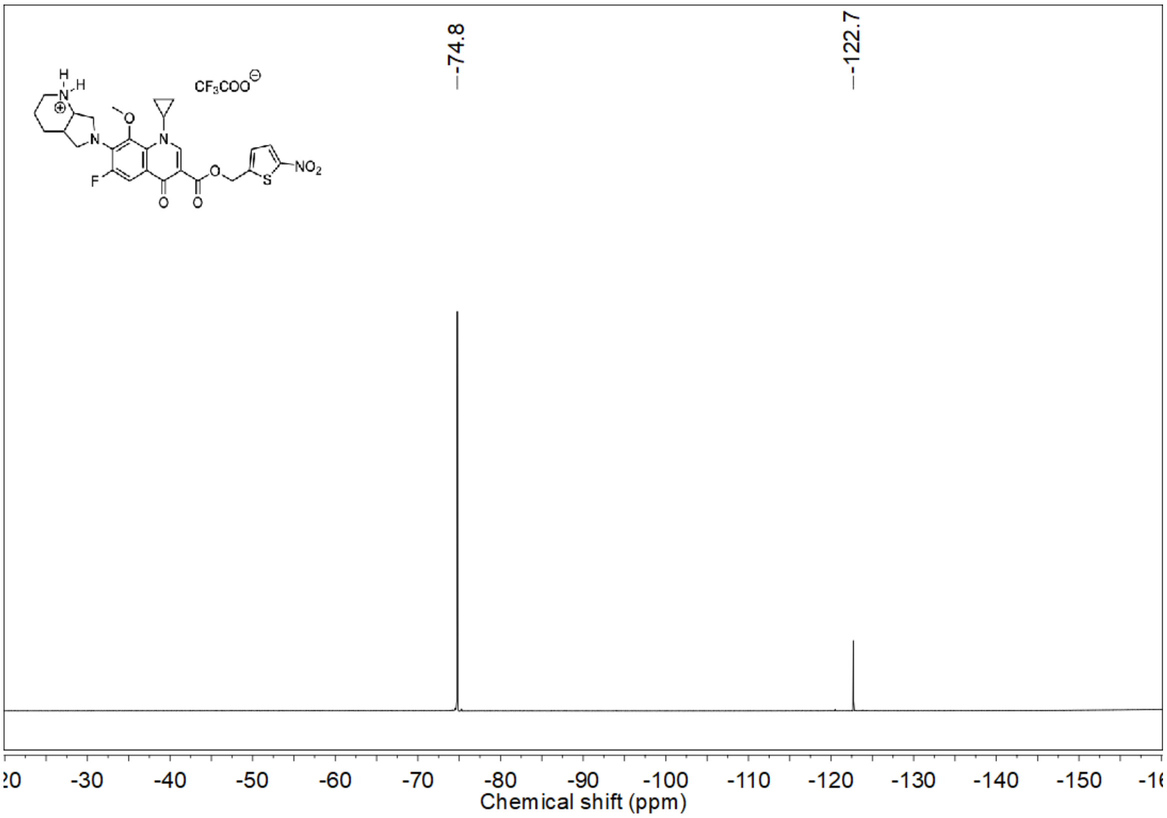


$^{1}$H NMR spectrum (400 MHz, DMSO-$d_6$) of **1c**

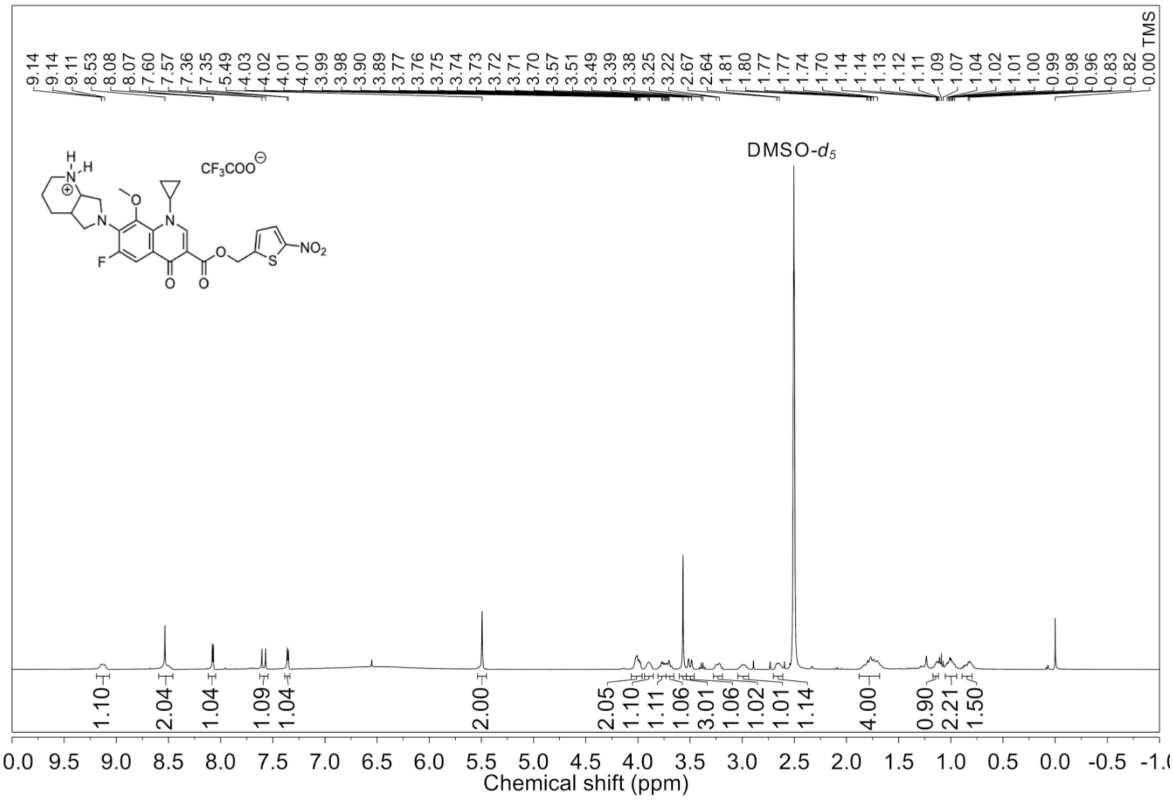

<sup></sup>$^{13}$C NMR spectrum (100 MHz, DMSO-$d_6$) of **1c**

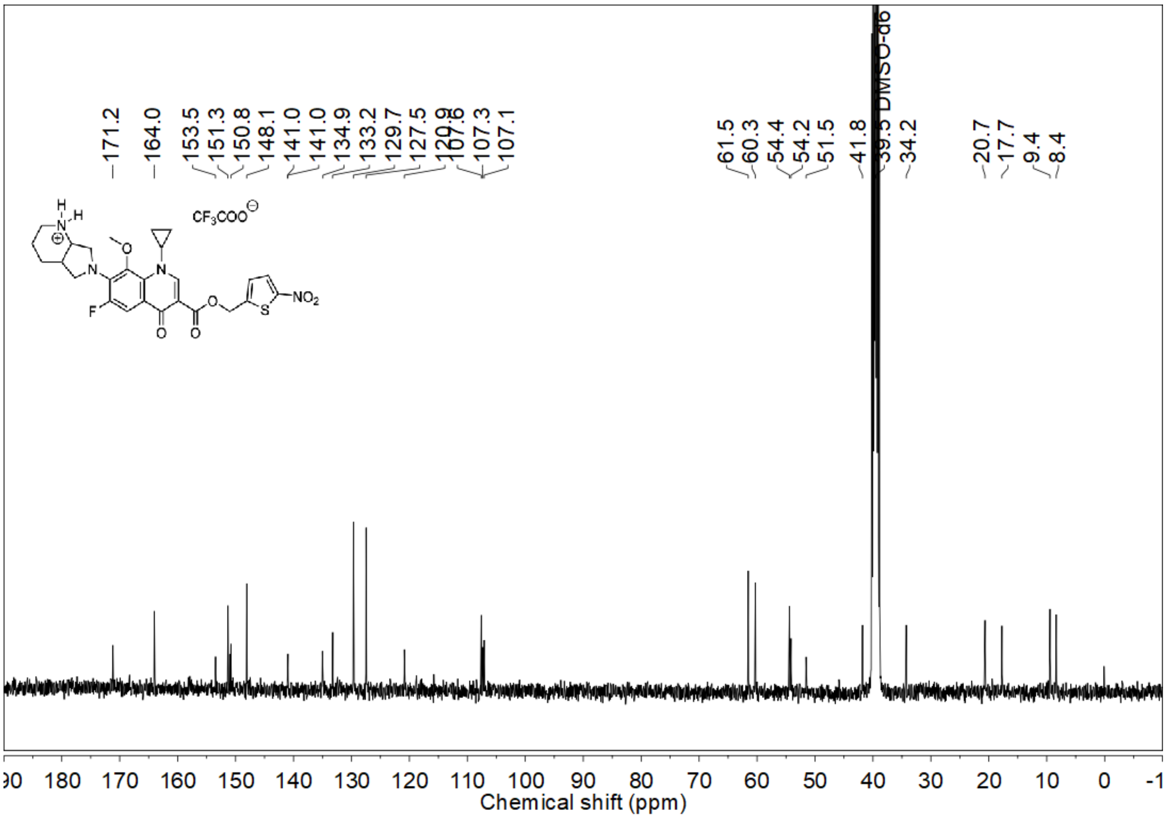


DEPT-135 NMR spectrum (100 MHz, DMSO-$d_6$) of **1c**

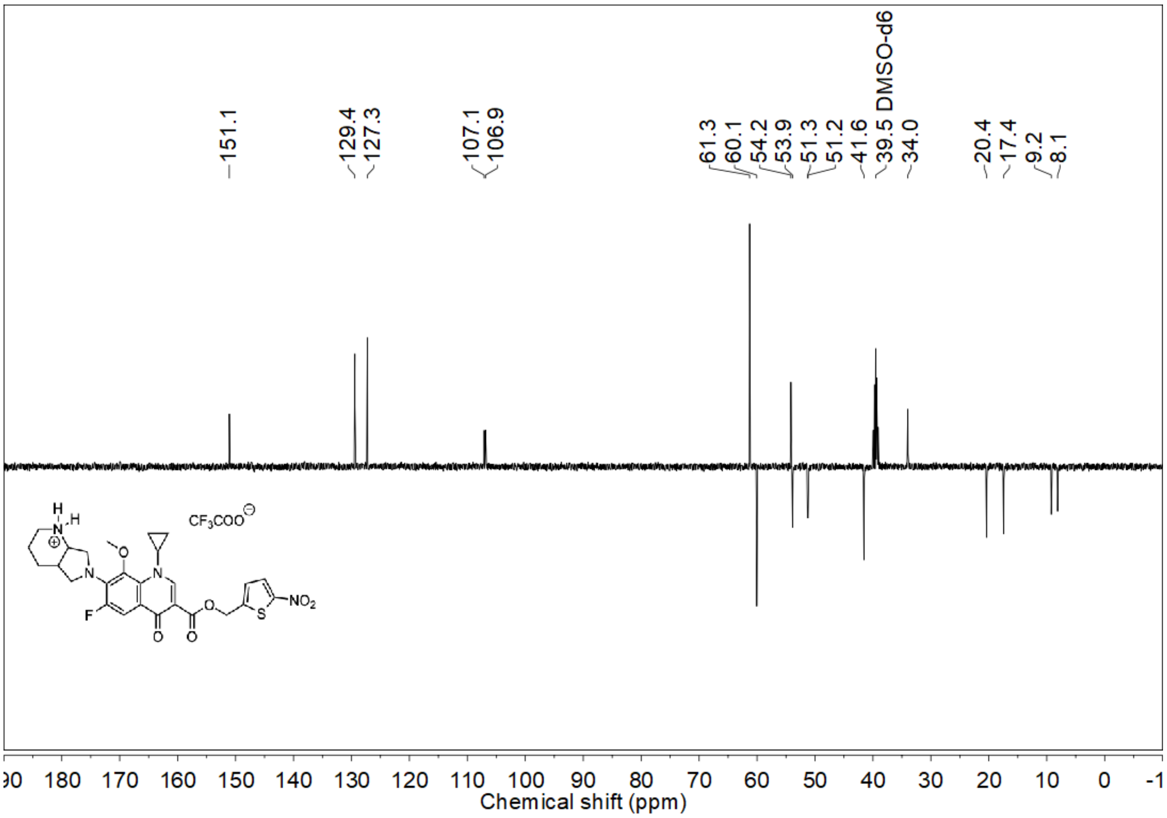

$^{19}$F NMR spectrum (376 MHz, DMSO-$d_6$) of **1d**

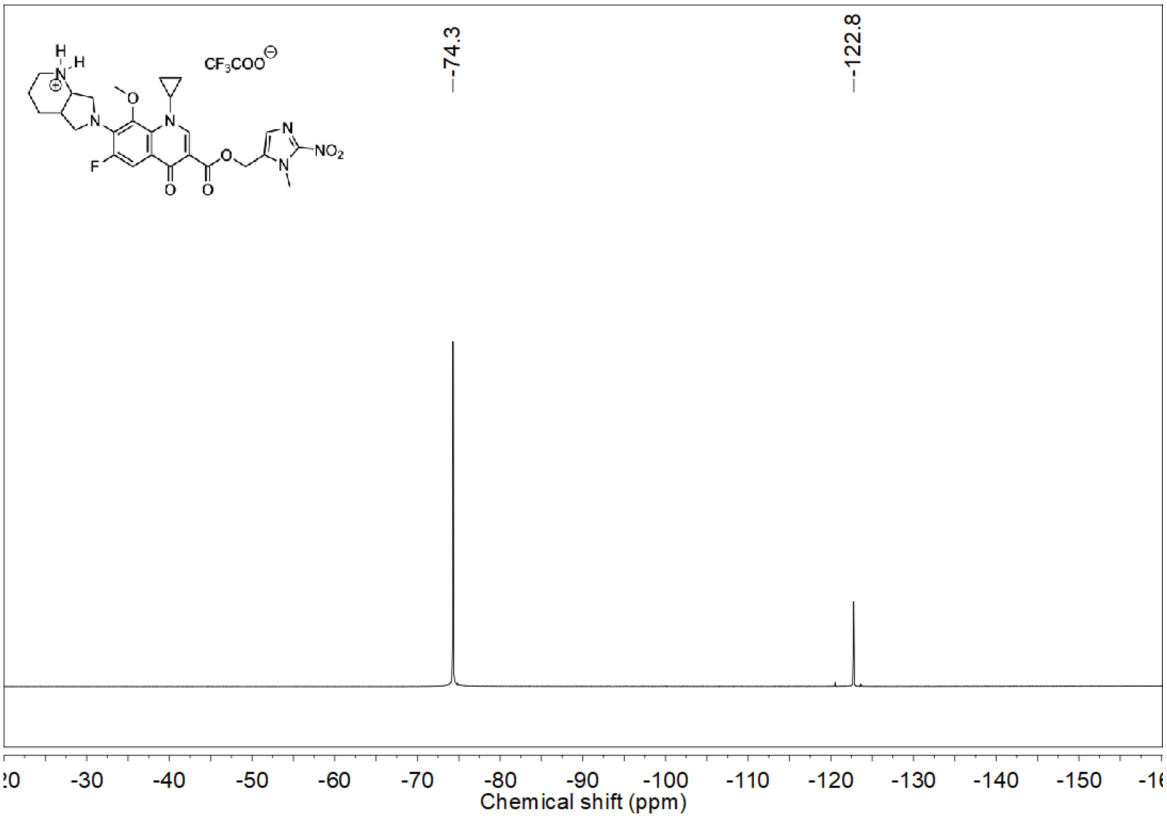

$^{1}$H NMR spectrum (400 MHz, DMSO-$d_6$) of **1d**

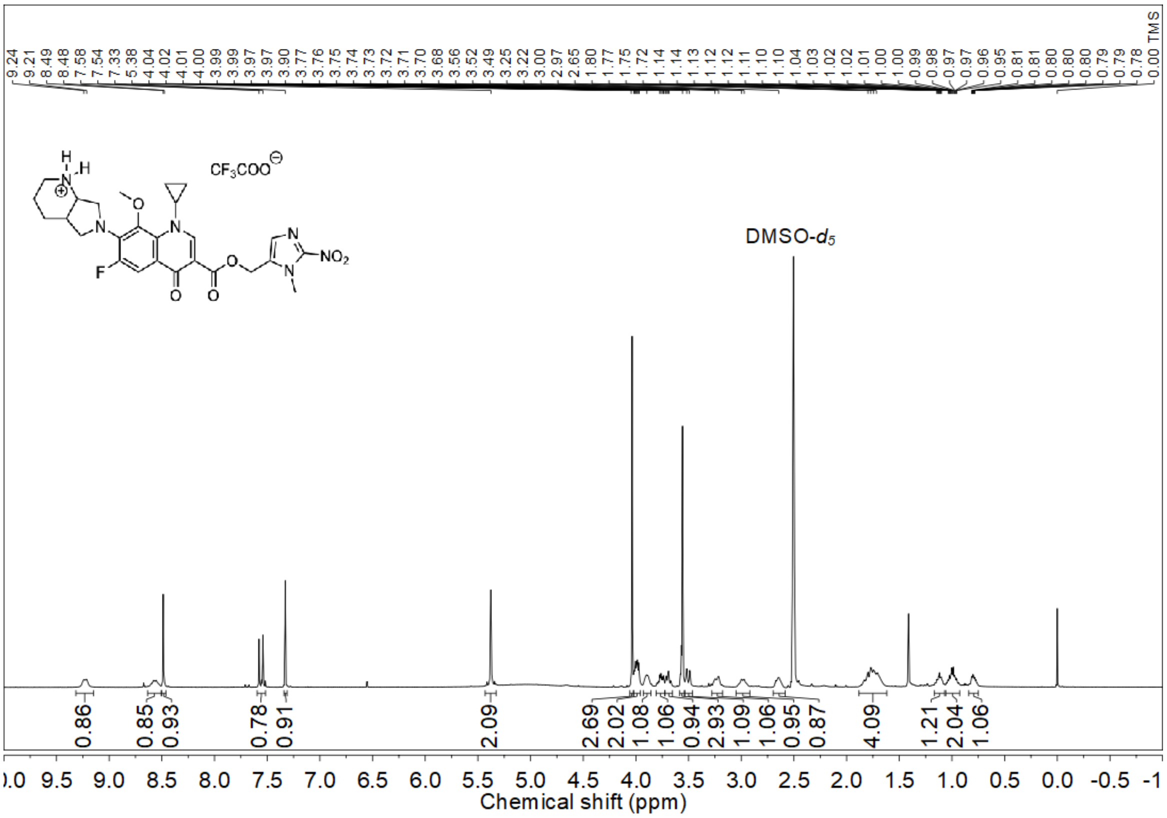

$^{13}$C NMR spectrum (100 MHz, DMSO-$d_6$) of **1d**

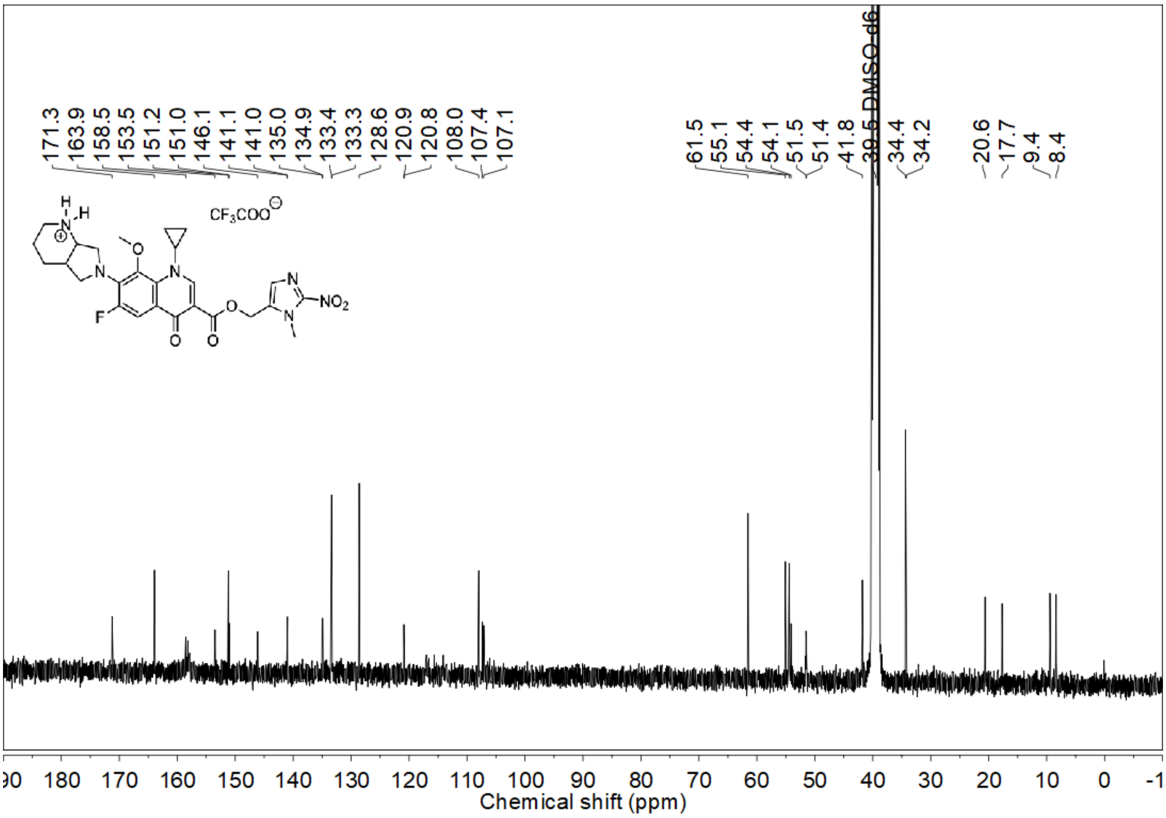


DEPT-135 NMR spectrum (100 MHz, DMSO-$d_6$) of **1d**

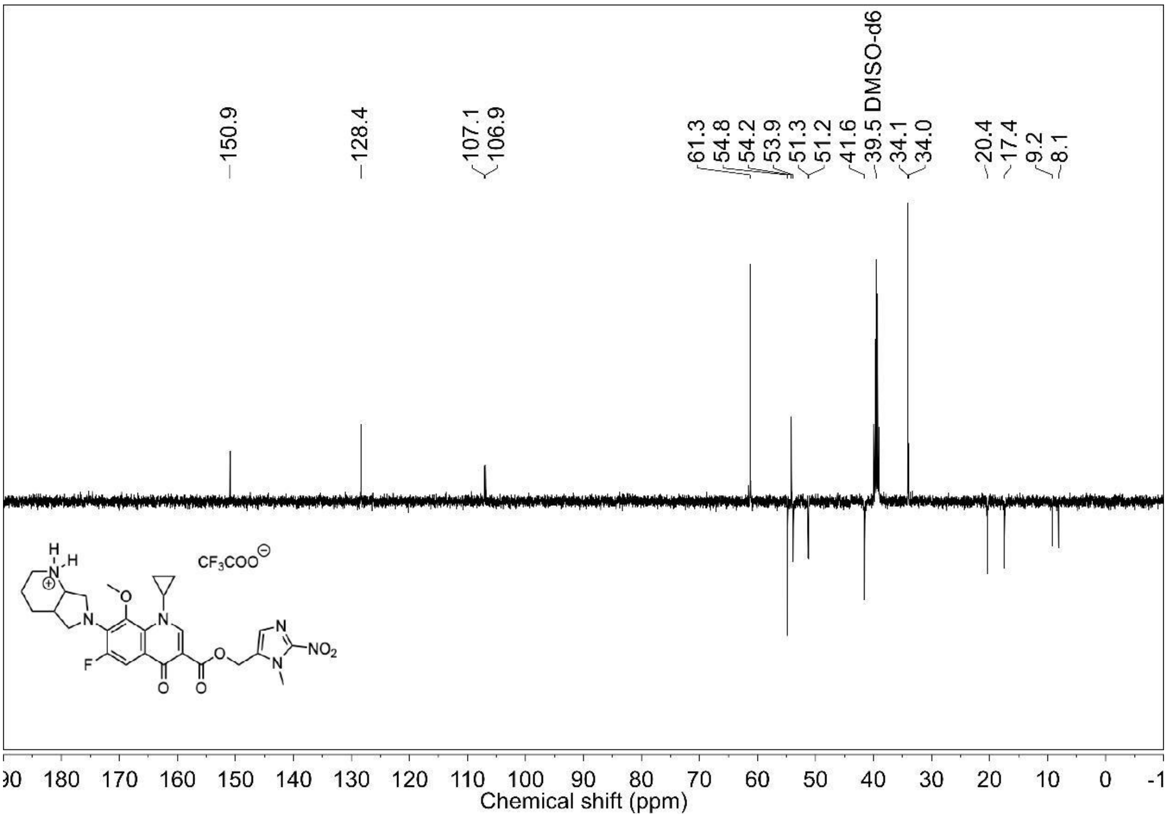

[19]F NMR spectrum (376 MHz, DMSO-$d_6$) of **1e**

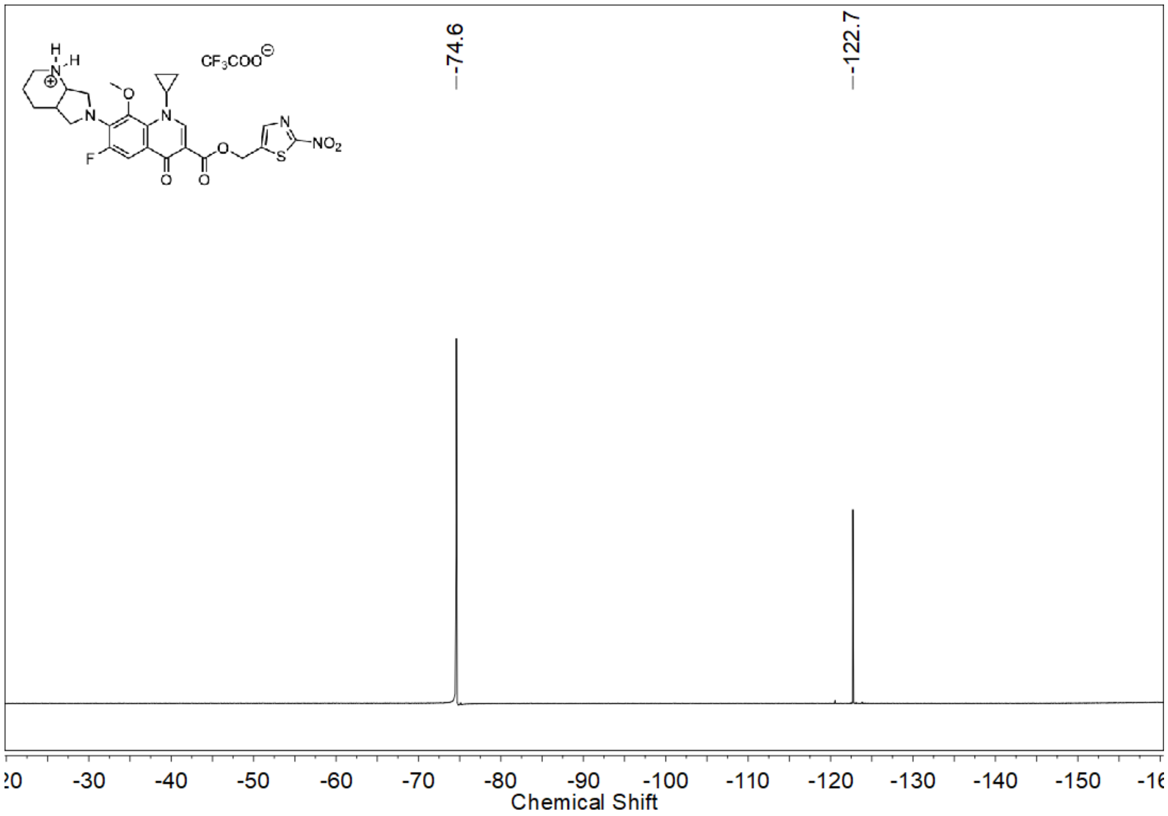


[1]H NMR spectrum (400 MHz, DMSO-$d_6$) of **1e**

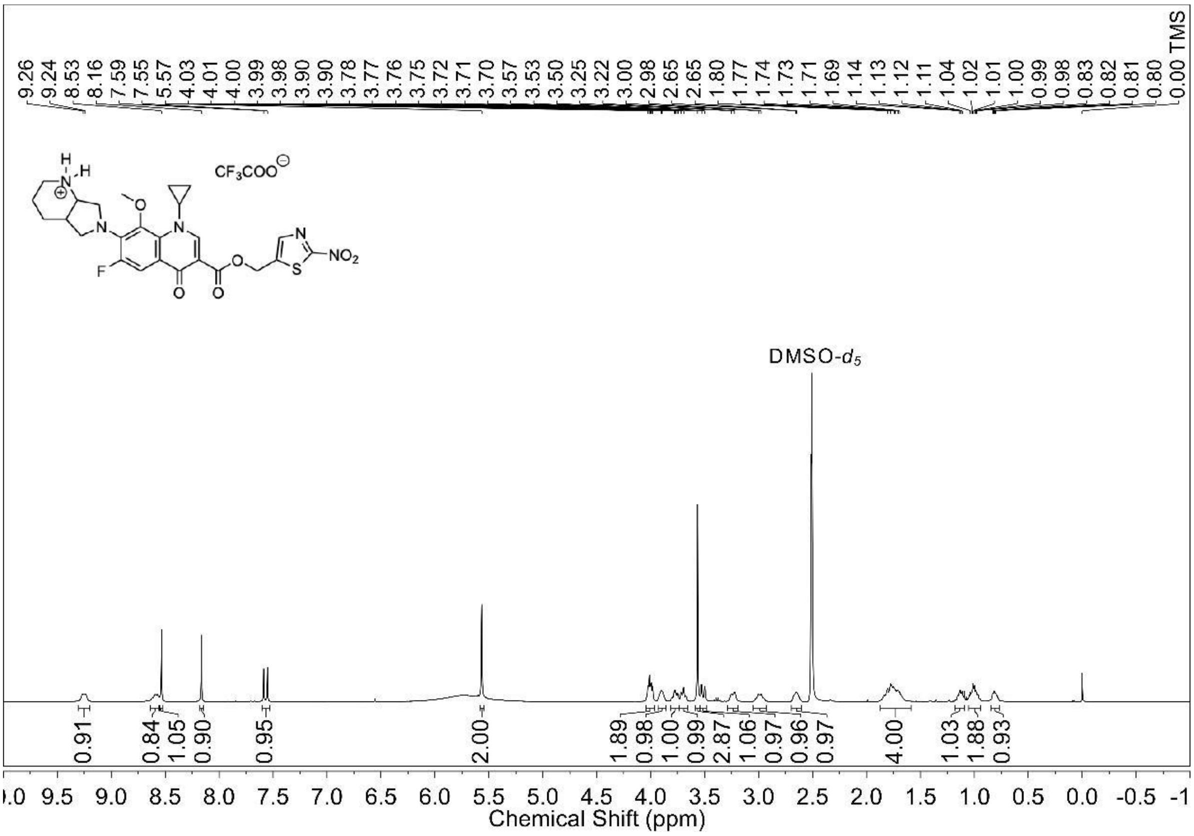

$^{13}C$ NMR spectrum (100 MHz, DMSO-$d_6$) of **1e**

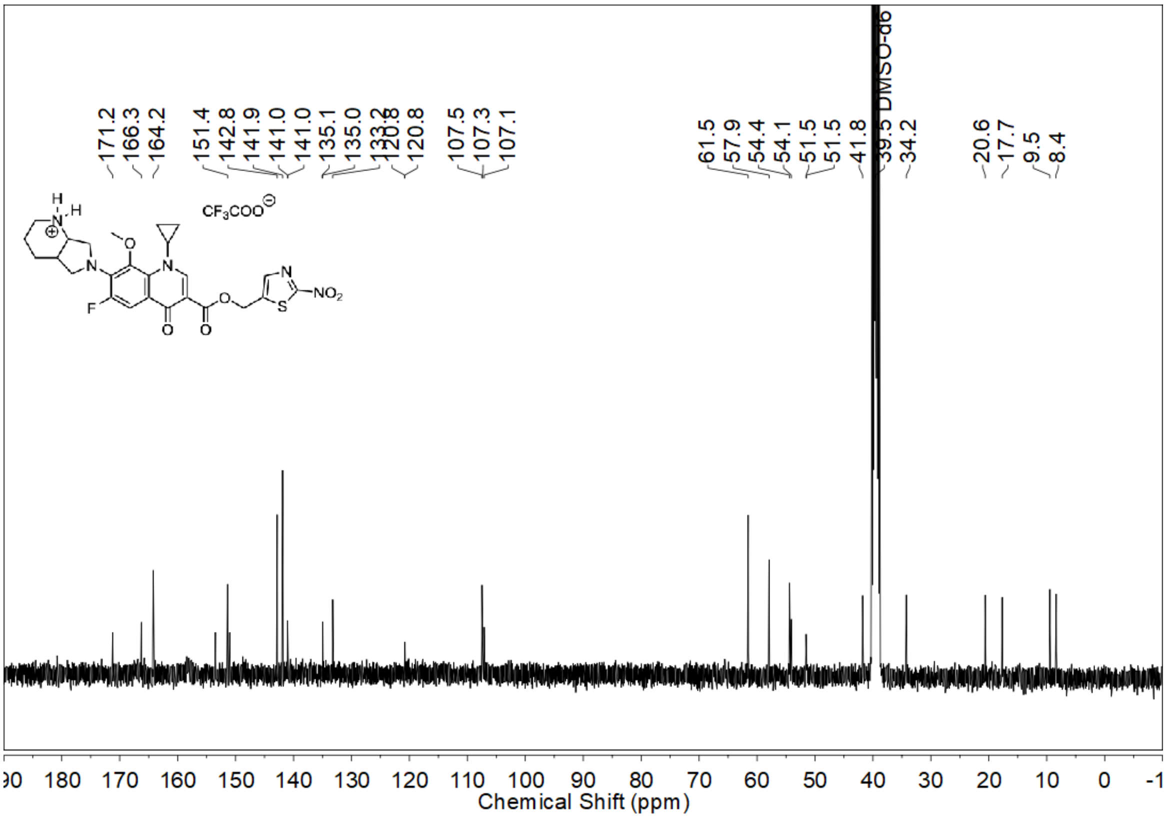

DEPT-135 NMR spectrum (100 MHz, DMSO-$d_6$) of **1e**

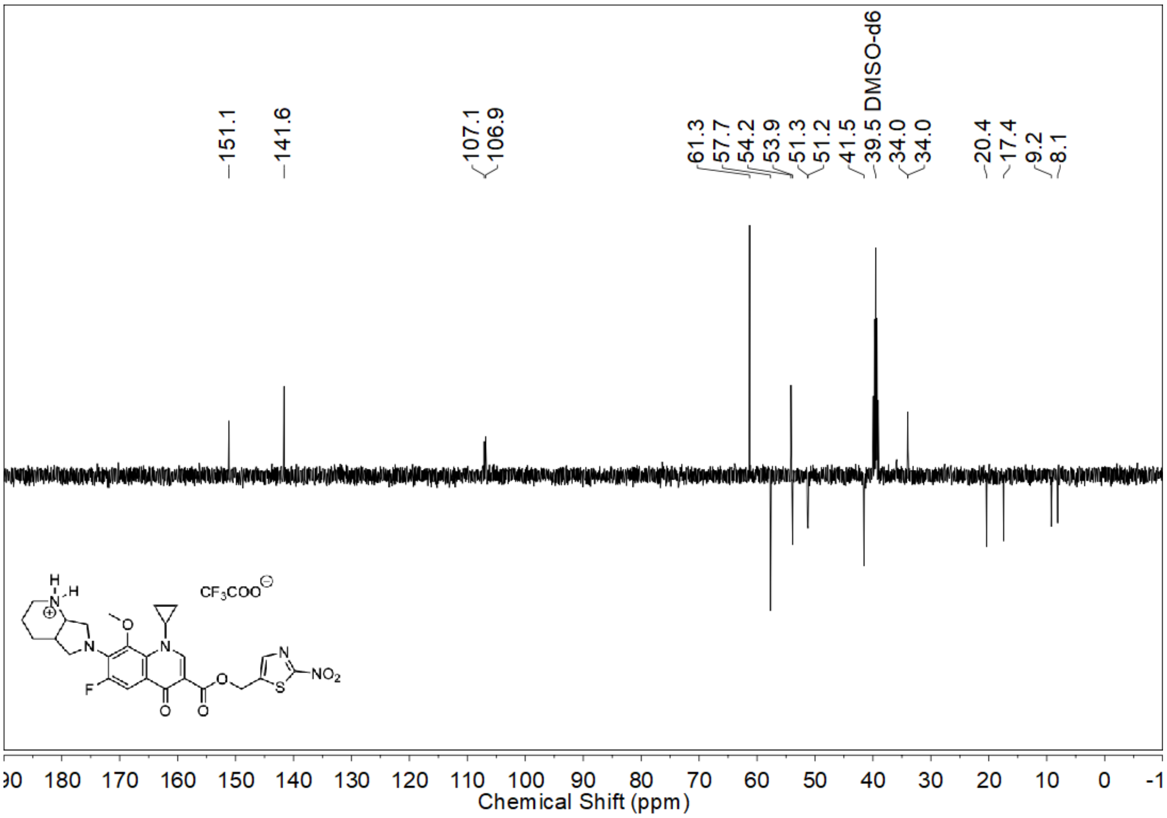

$^{19}$F NMR spectrum (376 MHz, DMSO-$d_6$) of **1f**

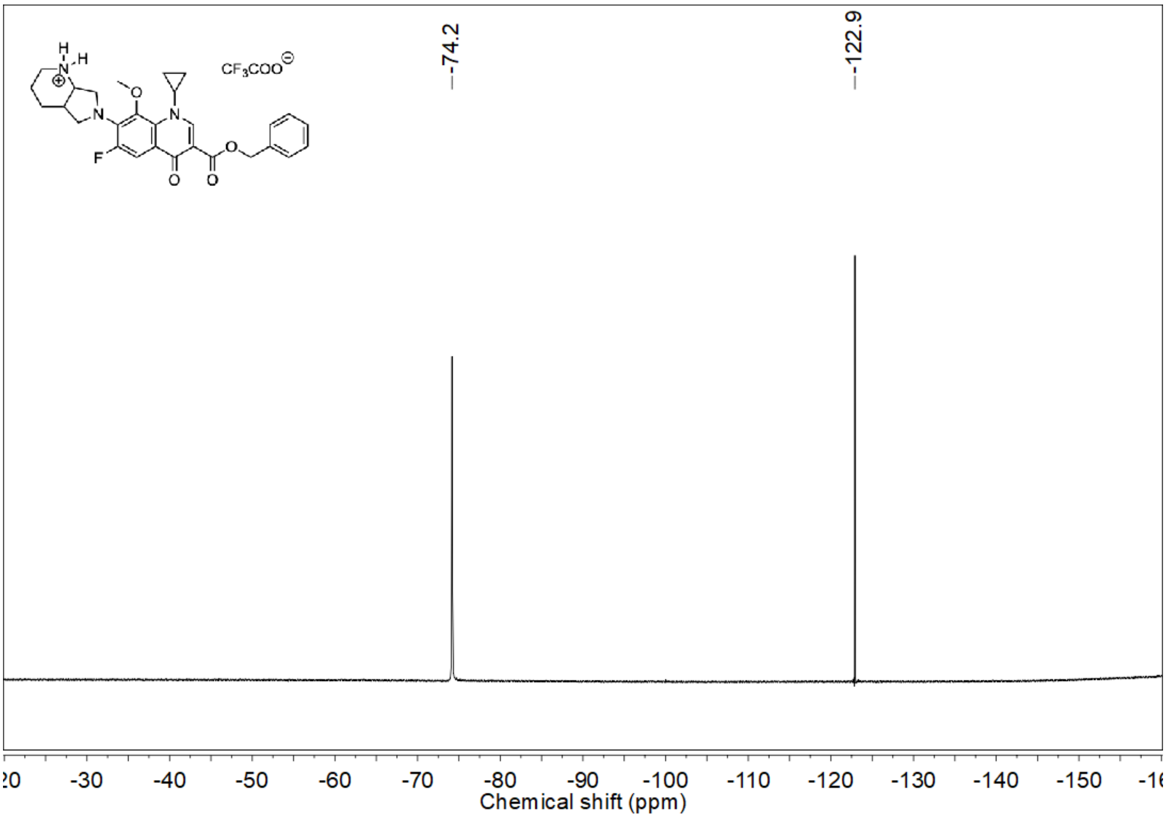


$^{1}$H NMR spectrum (400 MHz, DMSO-$d_6$) of **1f**

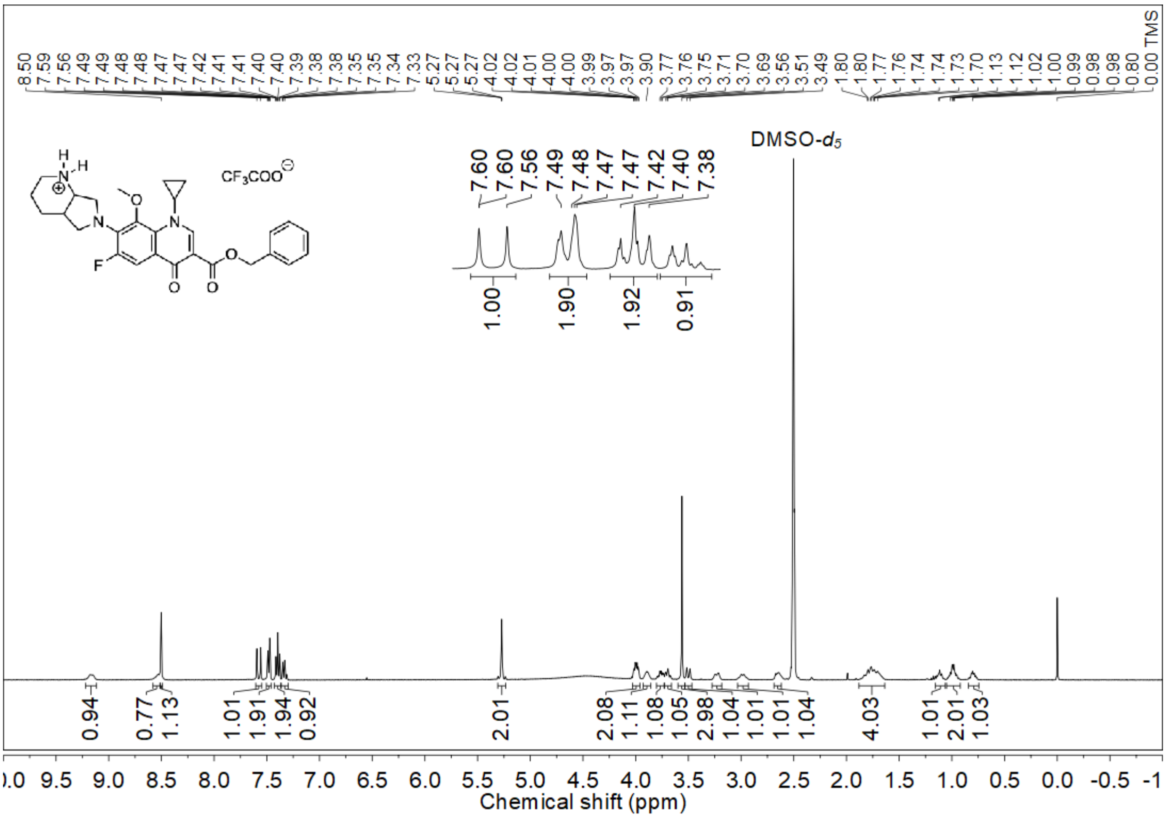

$^{13}$C NMR spectrum (100 MHz, DMSO-$d_6$) of **1f**

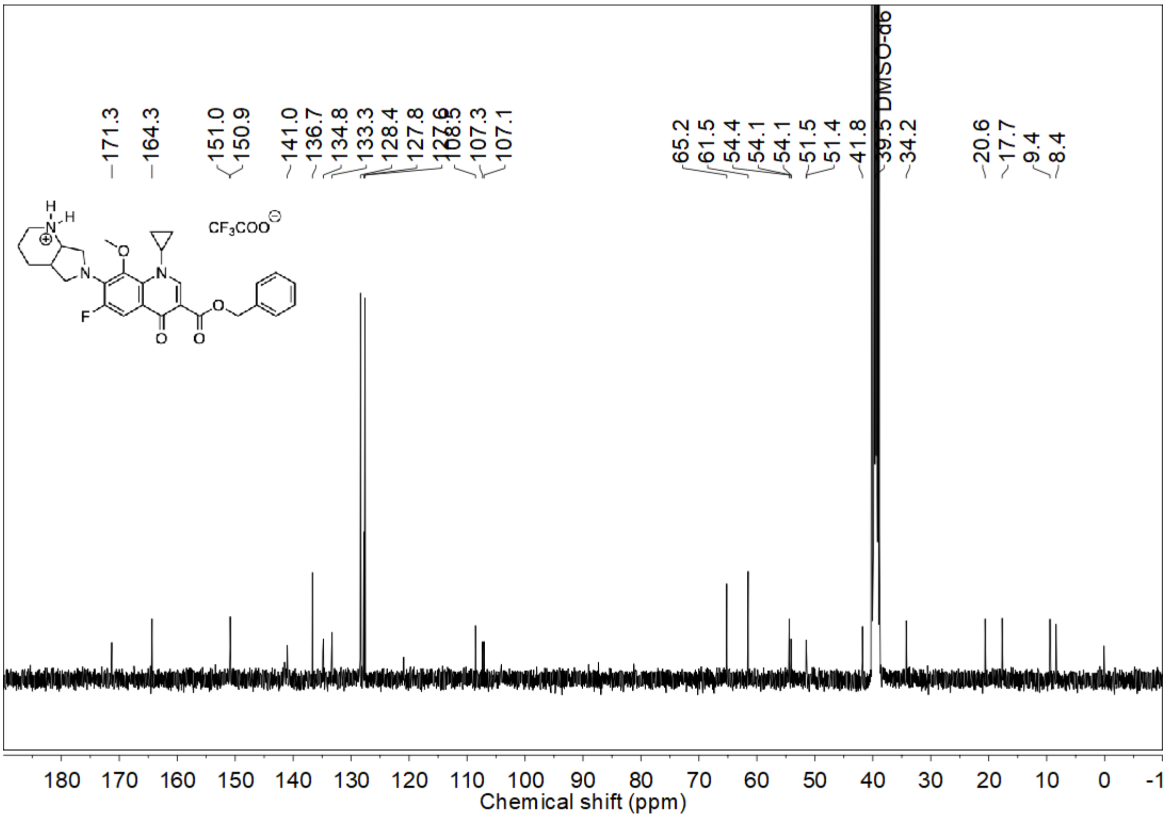


DEPT-135 NMR spectrum (100 MHz, DMSO-$d_6$) of **1f**

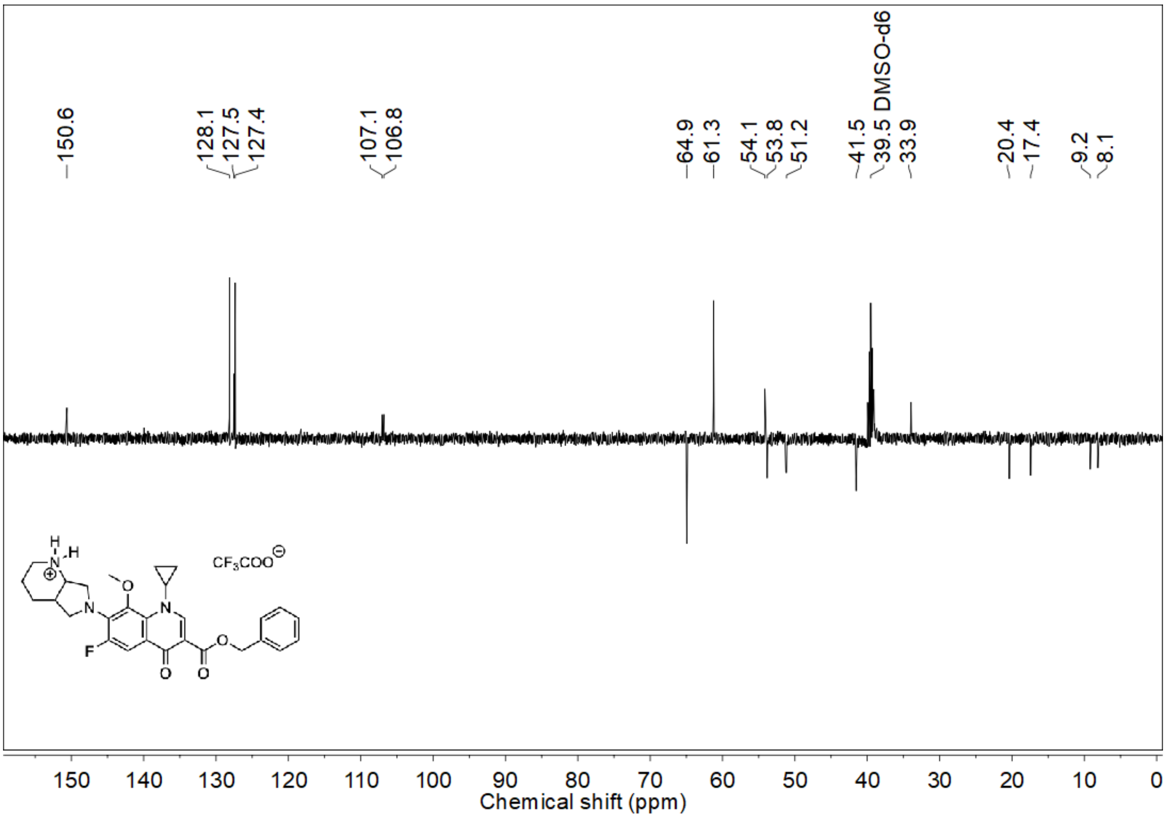

[19]F NMR spectrum (376 MHz, DMSO-$d_6$) of **1g**

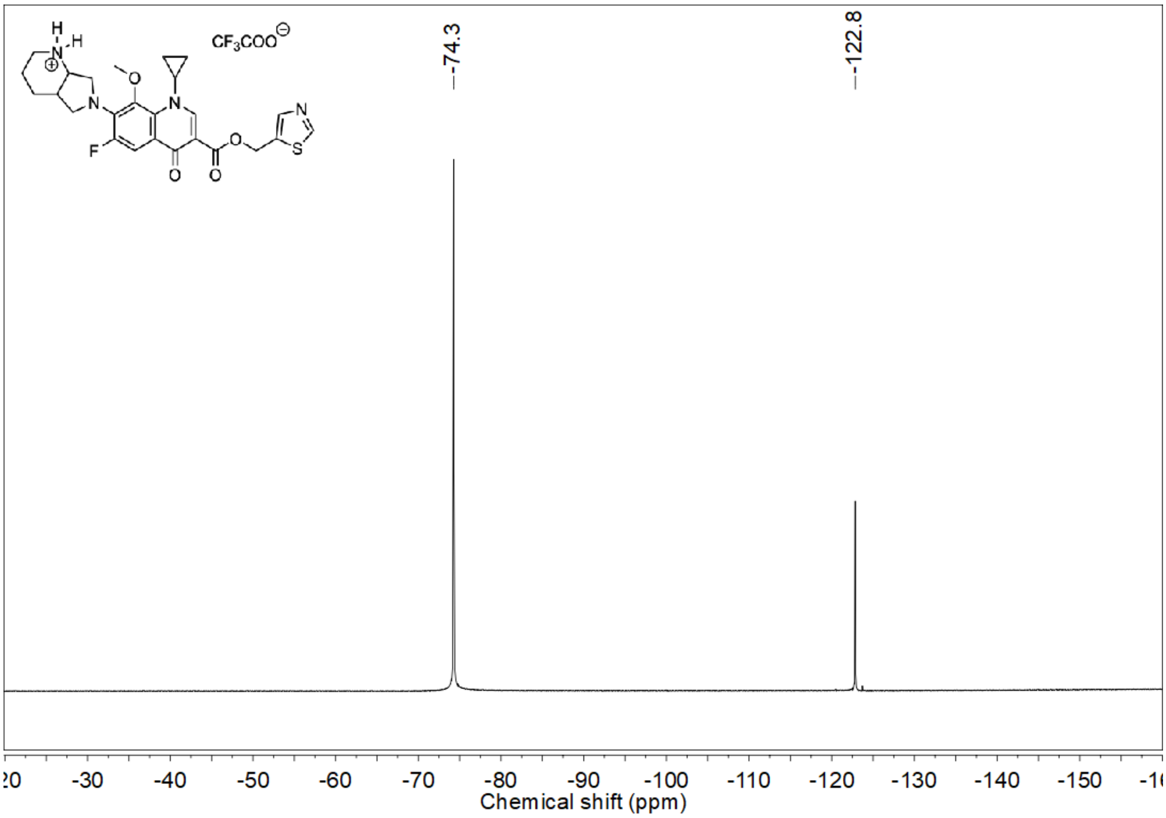


[1]H NMR spectrum (400 MHz, DMSO-$d_6$) of **1g**

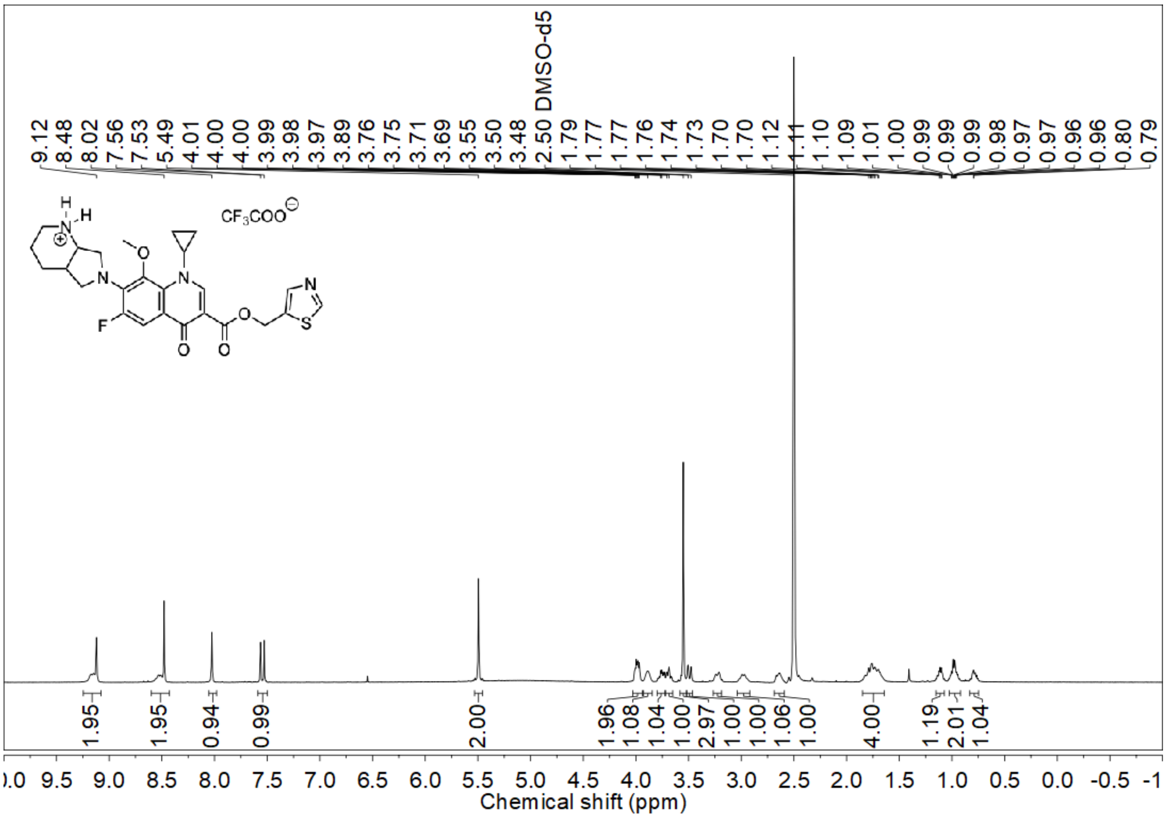

[13]C NMR spectrum (100 MHz, DMSO-$d_6$) of **1g**

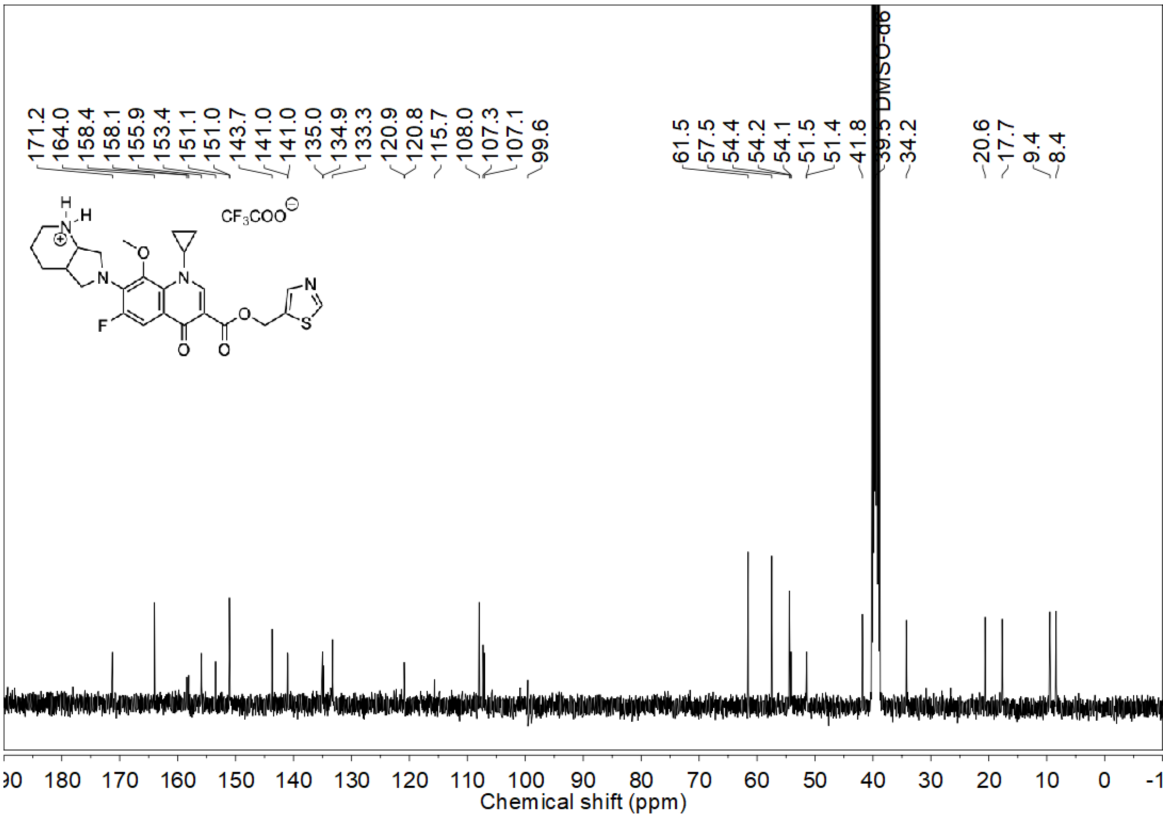


DEPT-135 NMR spectrum (100 MHz, DMSO-$d_6$) of **1g**

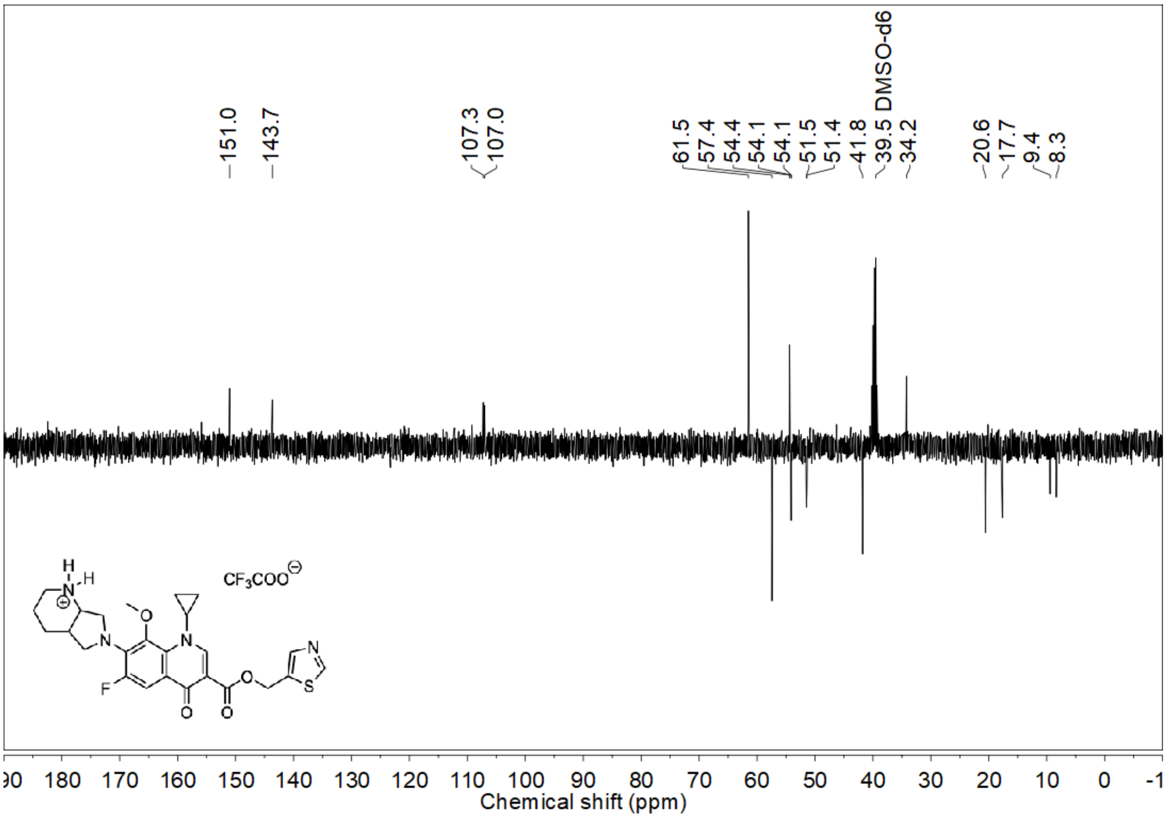

$^{19}$F NMR spectrum (376 MHz, $CDCl_3$) of **2a**

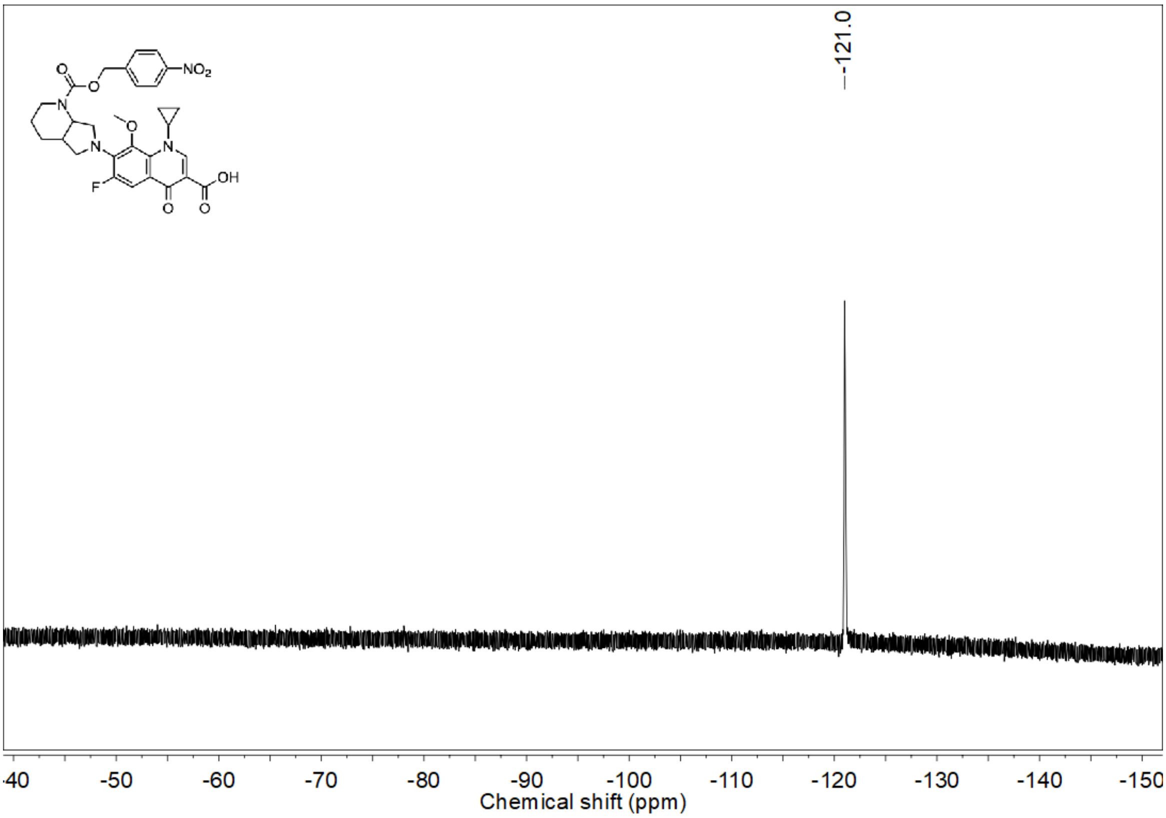


$^{1}$H NMR spectrum (400 MHz, $CDCl_3$) of **2a**

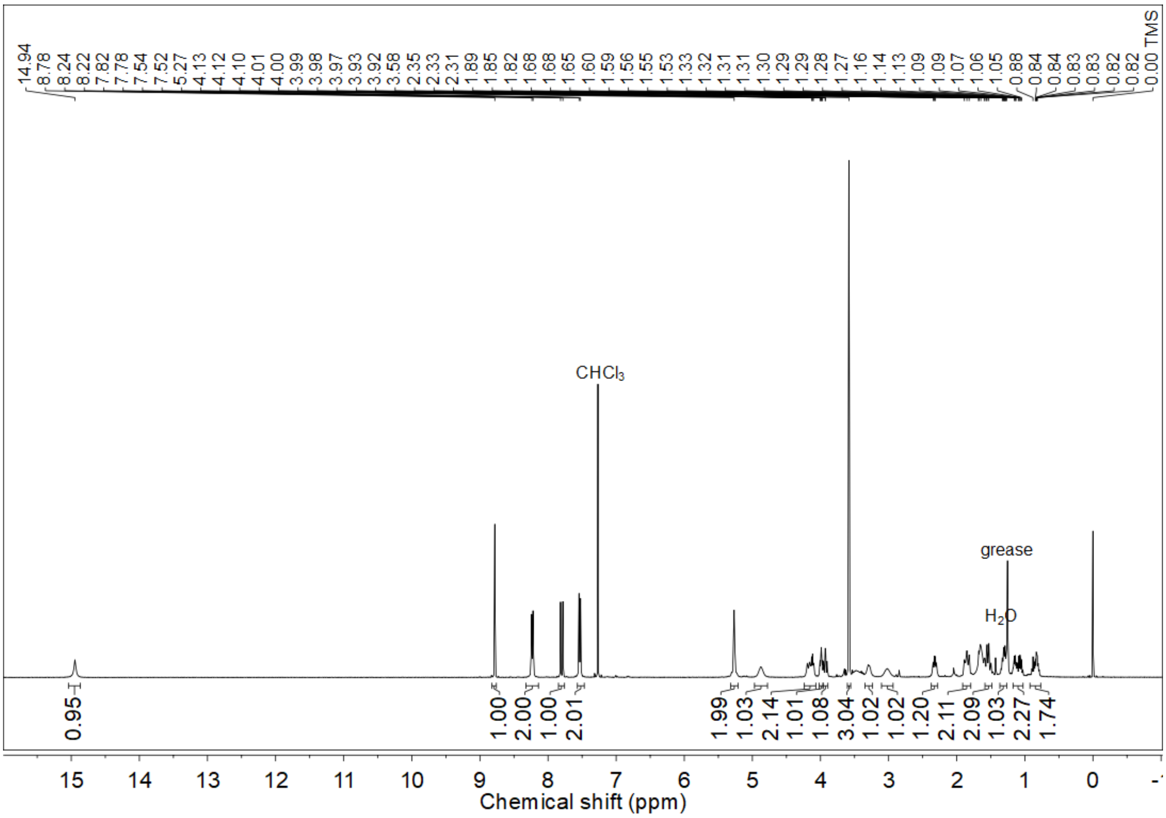

$^{13}$C NMR spectrum (100 MHz, CDCl$_3$) of **2a**

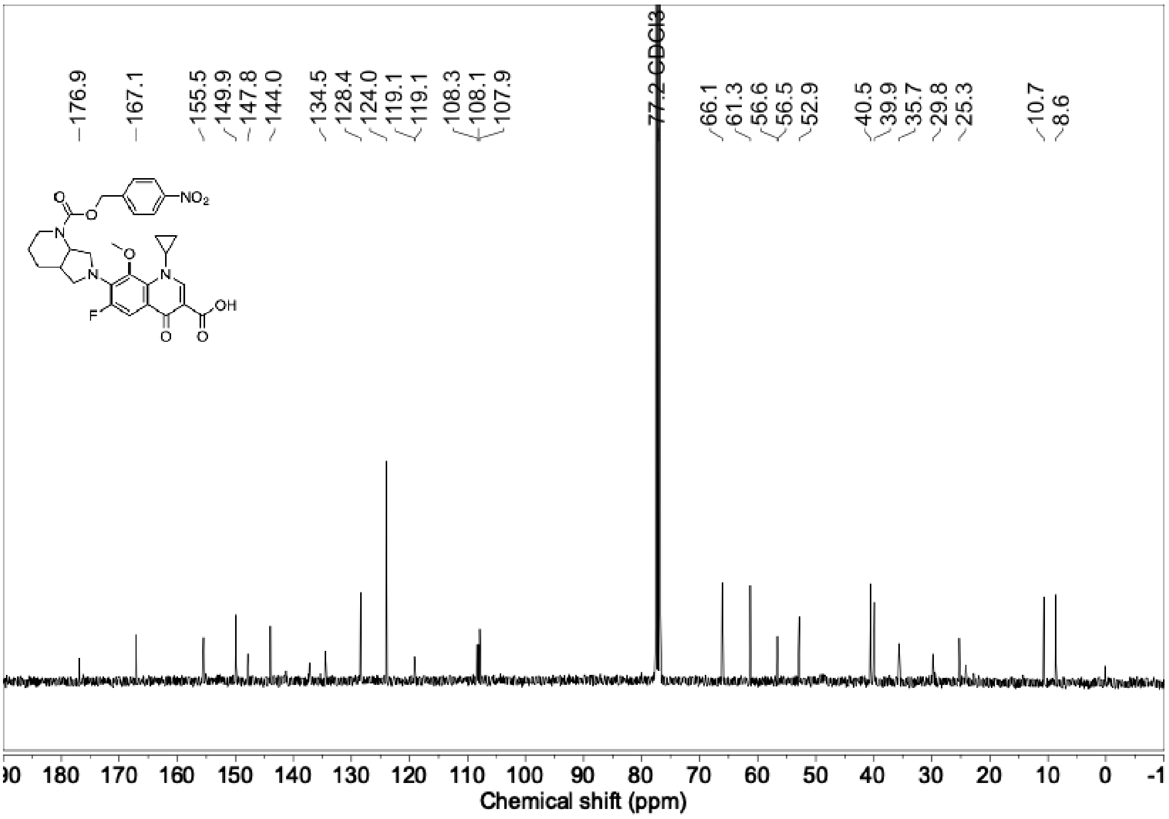


DEPT-135 NMR spectrum (100 MHz, CDCl$_3$) of **2a**

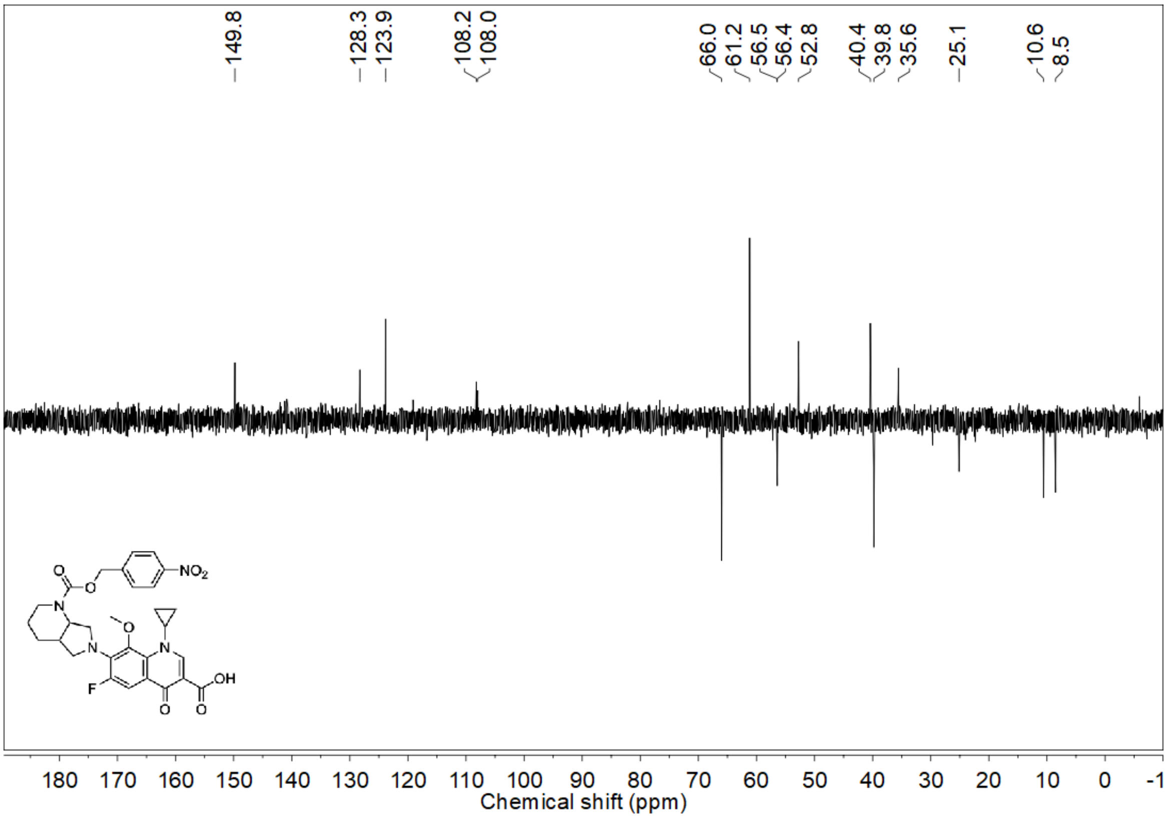

$^{19}$F NMR spectrum (376 MHz, $CDCl_3$) of **2b**

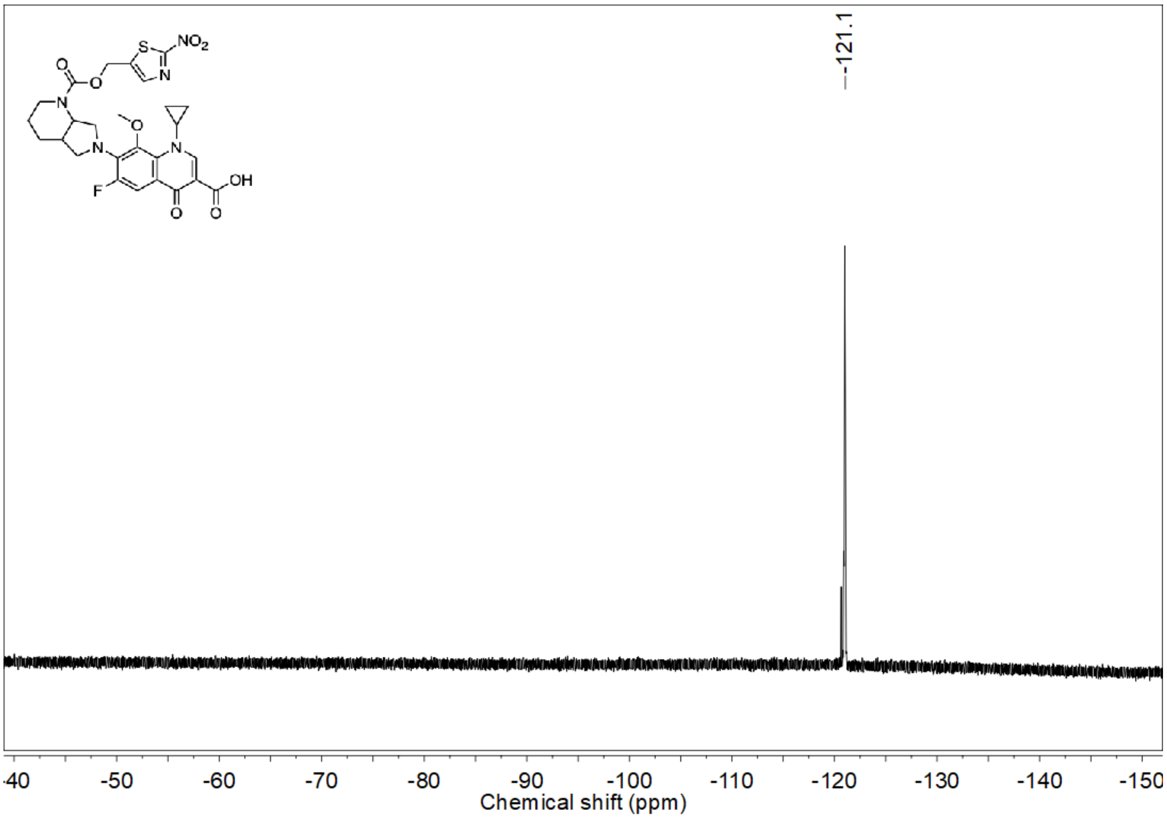


$^{1}$H NMR spectrum (400 MHz, $CDCl_3$) of **2b**

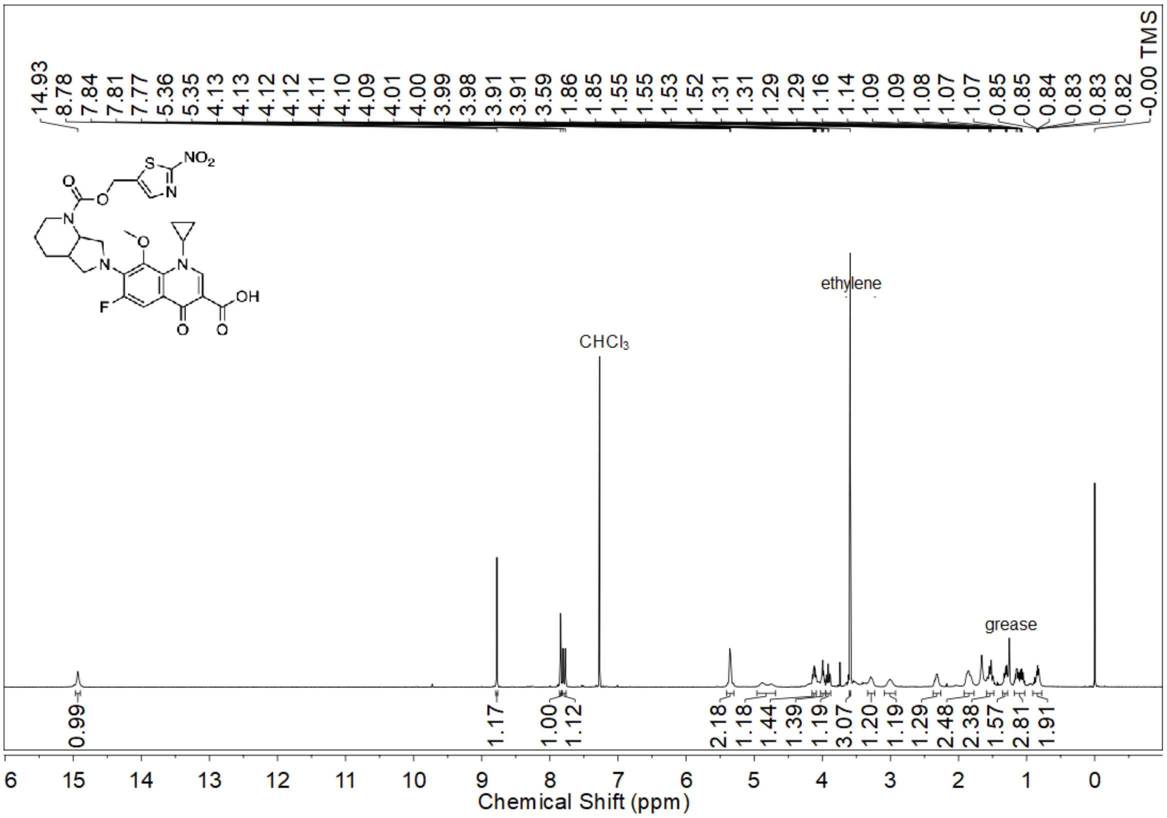

$^{13}C$ NMR spectrum (100 MHz, $CDCl_3$) of **2b**

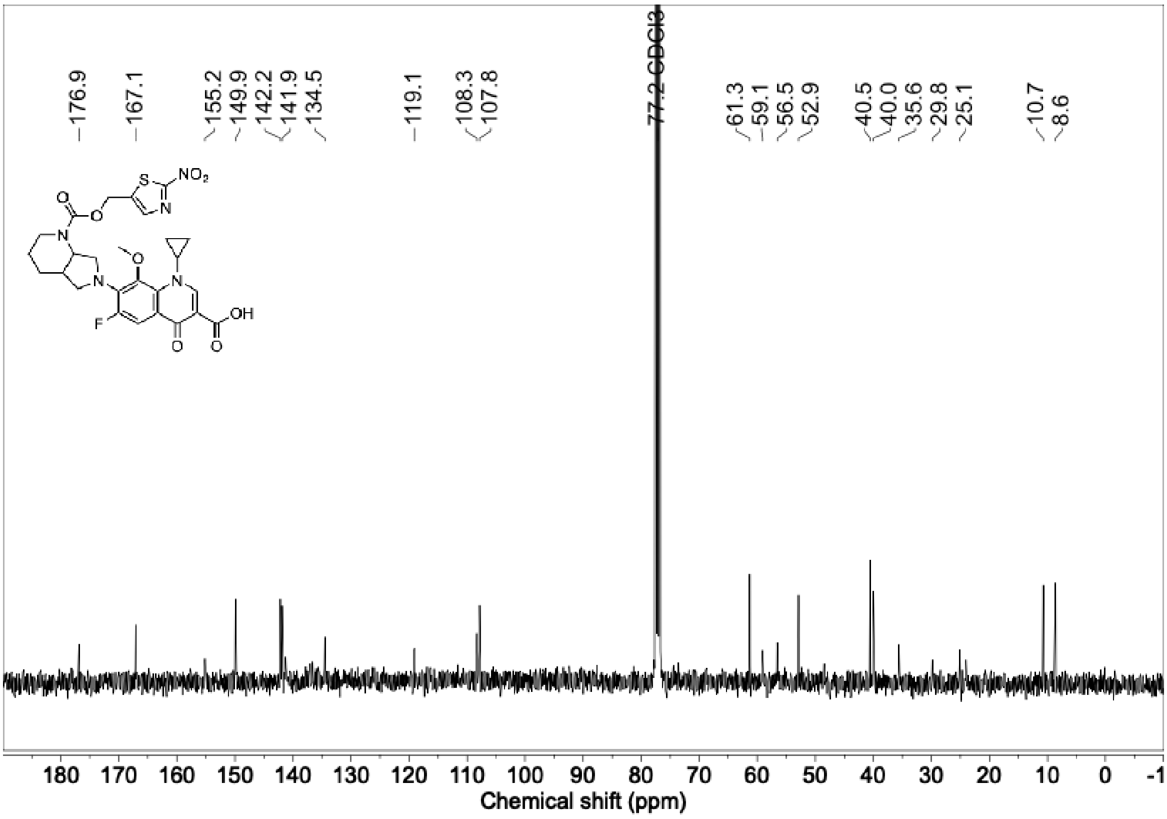


DEPT-135 NMR spectrum (100 MHz, $CDCl_3$) of **2b**

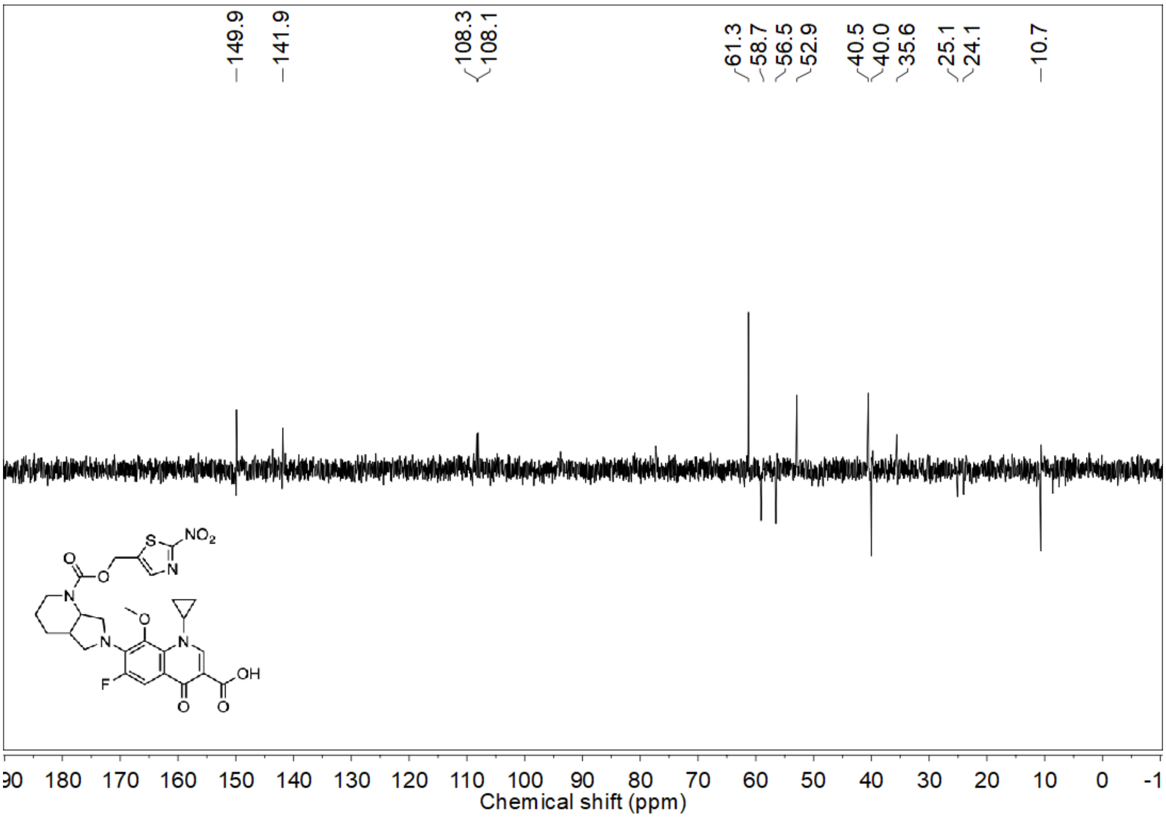

$^{19}F$ NMR spectrum (376 MHz, $CDCl_3$) of **3a**

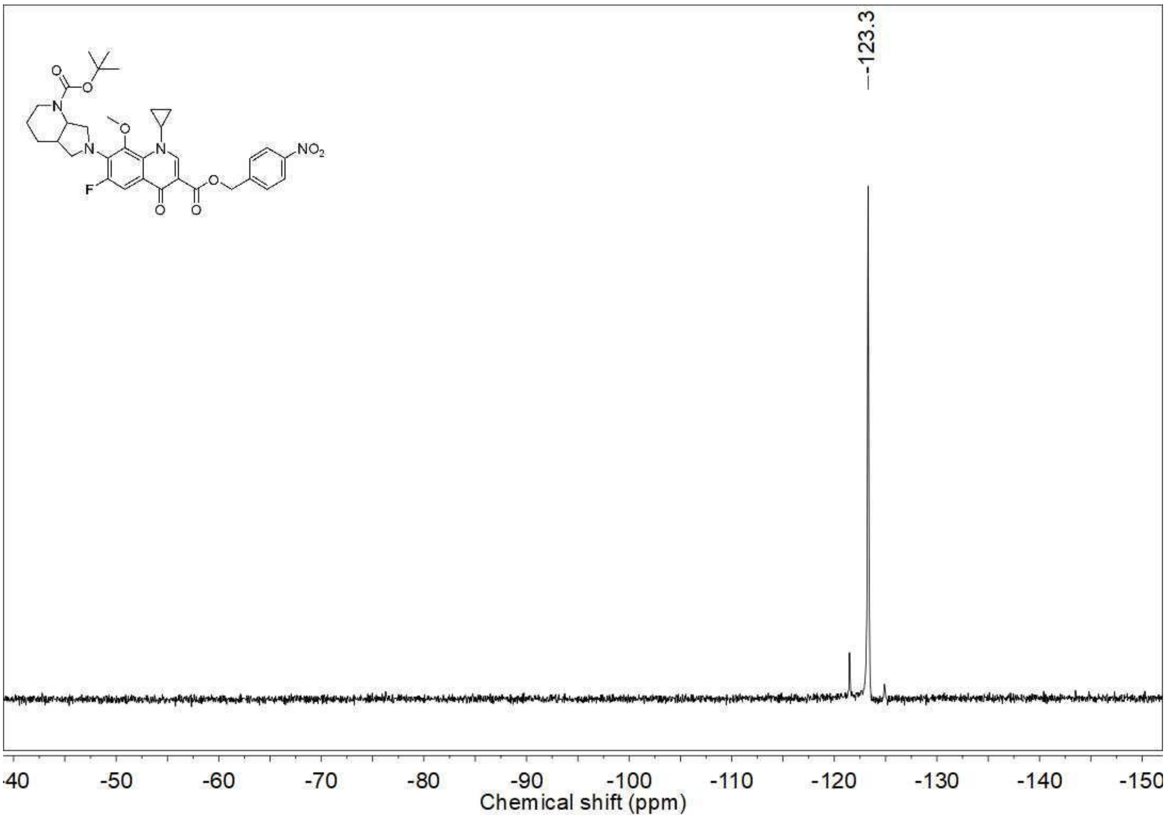


$^{1}H$ NMR spectrum (400 MHz, $CDCl_3$) of **3a**

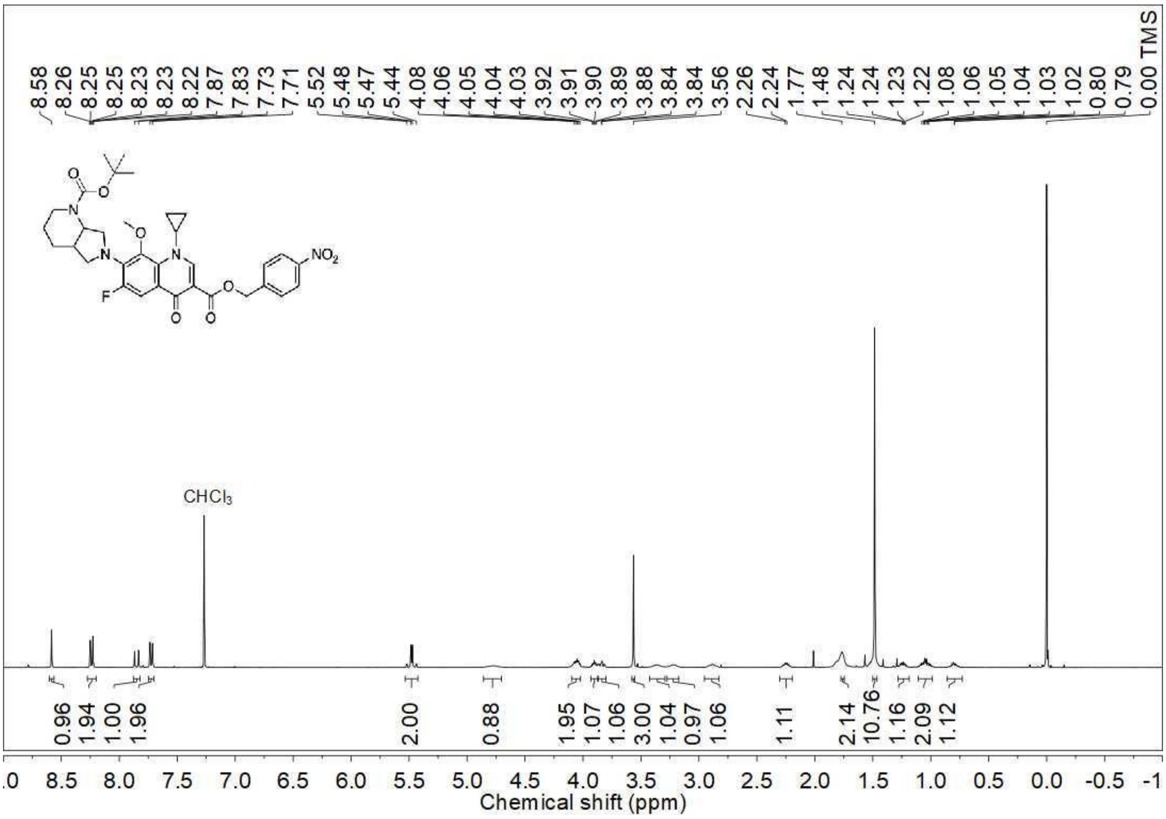

$^{13}$C NMR spectrum (100 MHz, $CDCl_3$) of **3a**

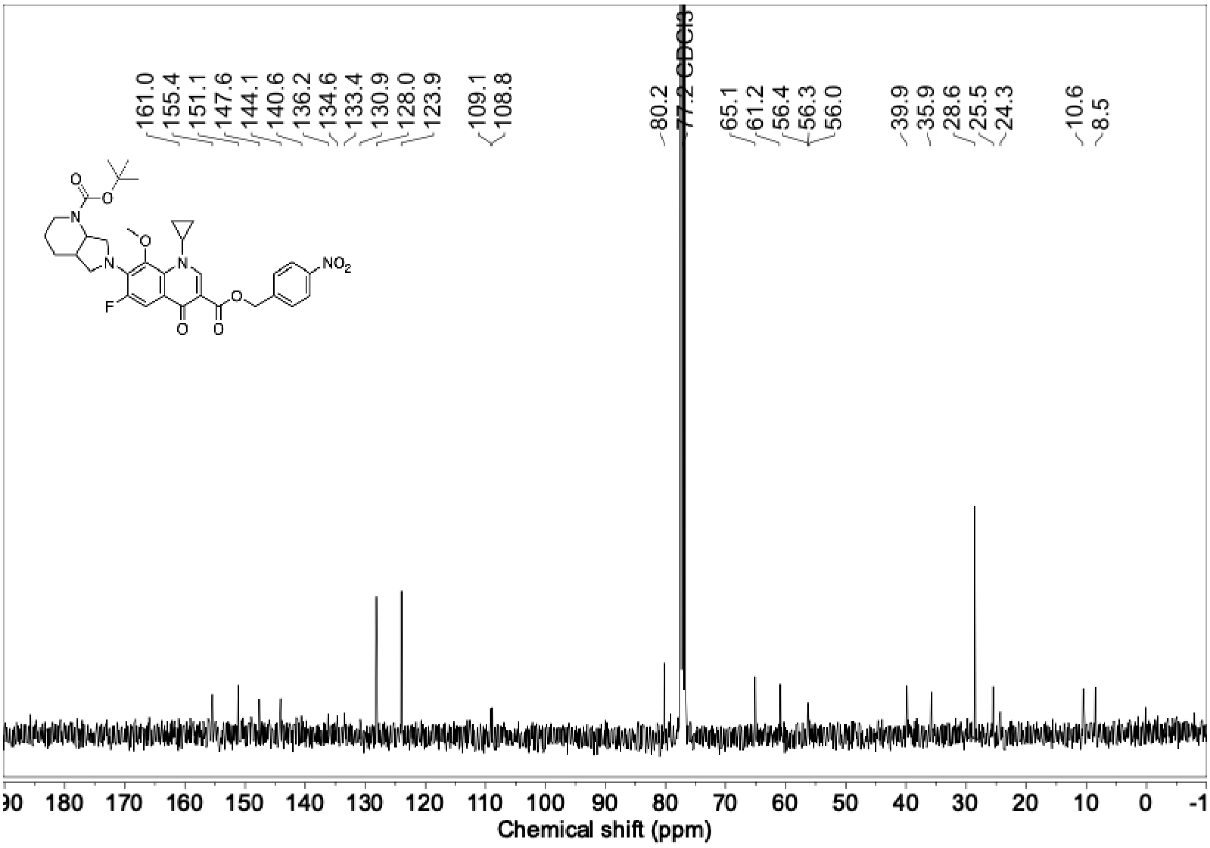


$^{19}$F NMR spectrum (376 MHz, $CDCl_3$) of **3b**

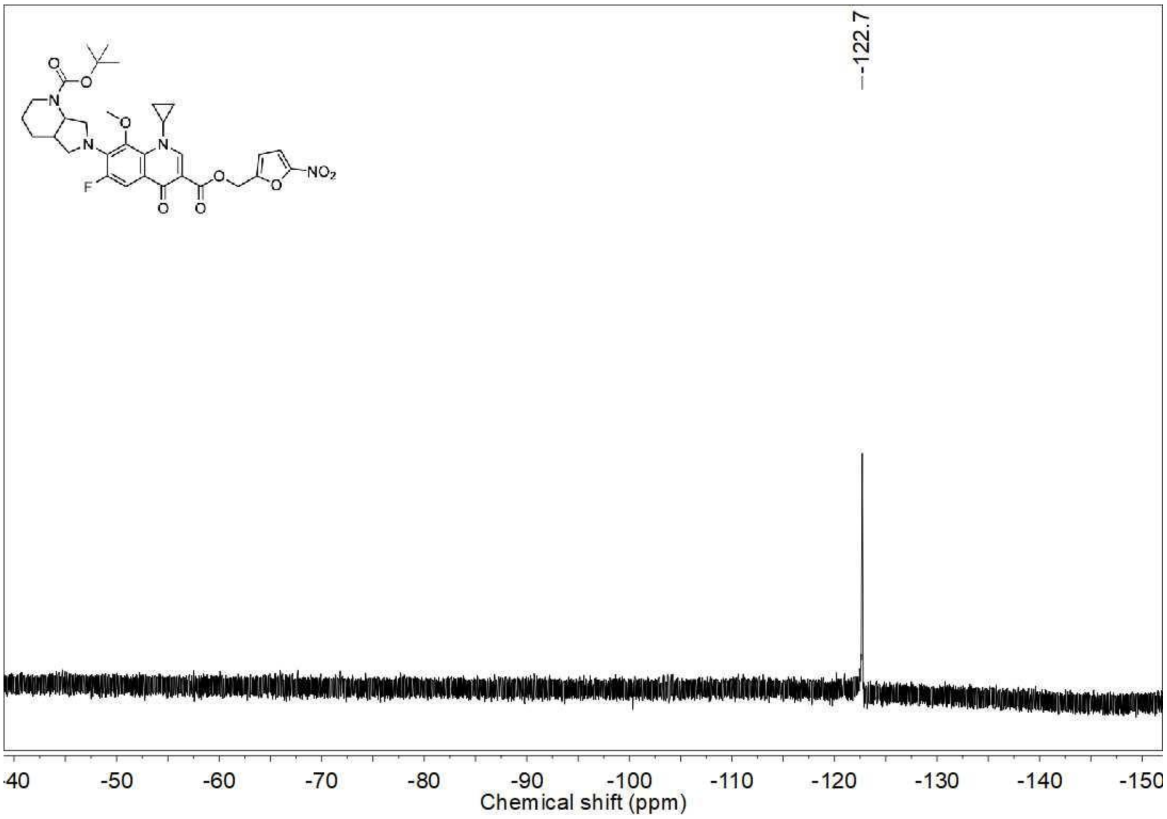

$^{1}H$ NMR spectrum (400 MHz, $CDCl_3$) of **3b**

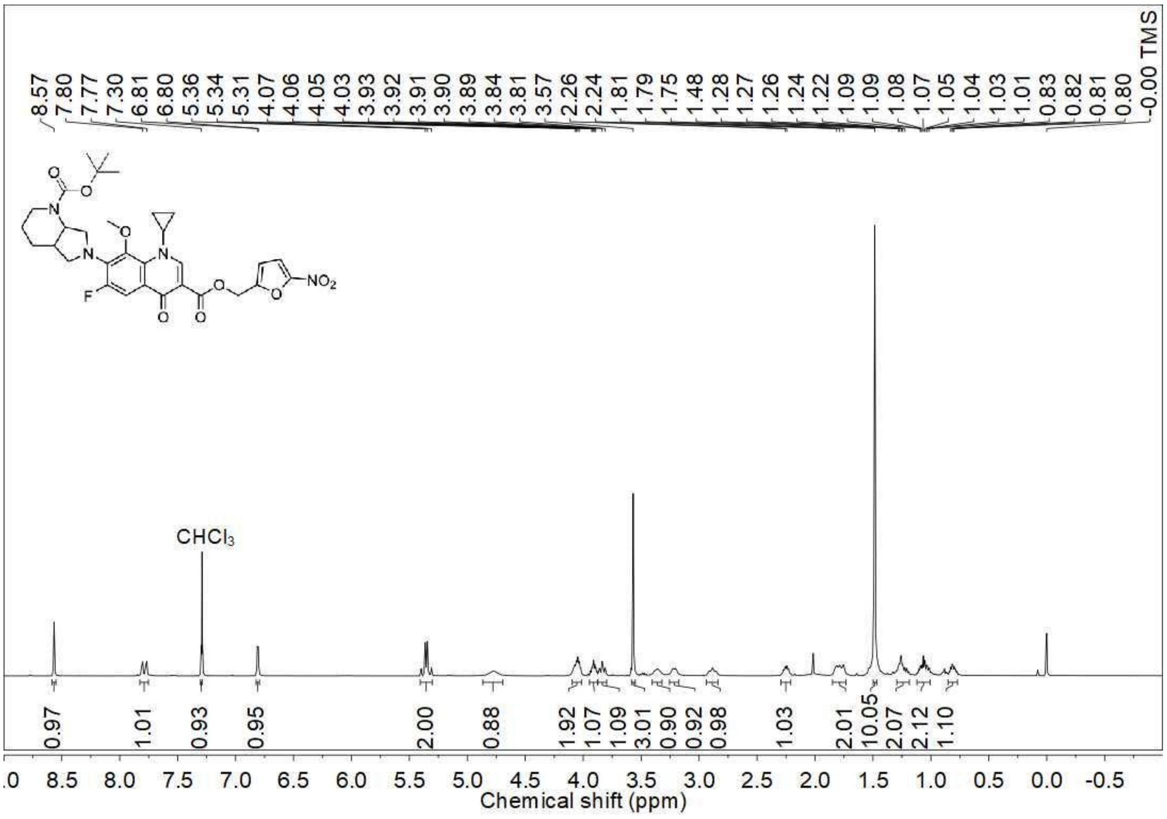

$^{13}C$ NMR spectrum (100 MHz, $CDCl_3$) of **3b**

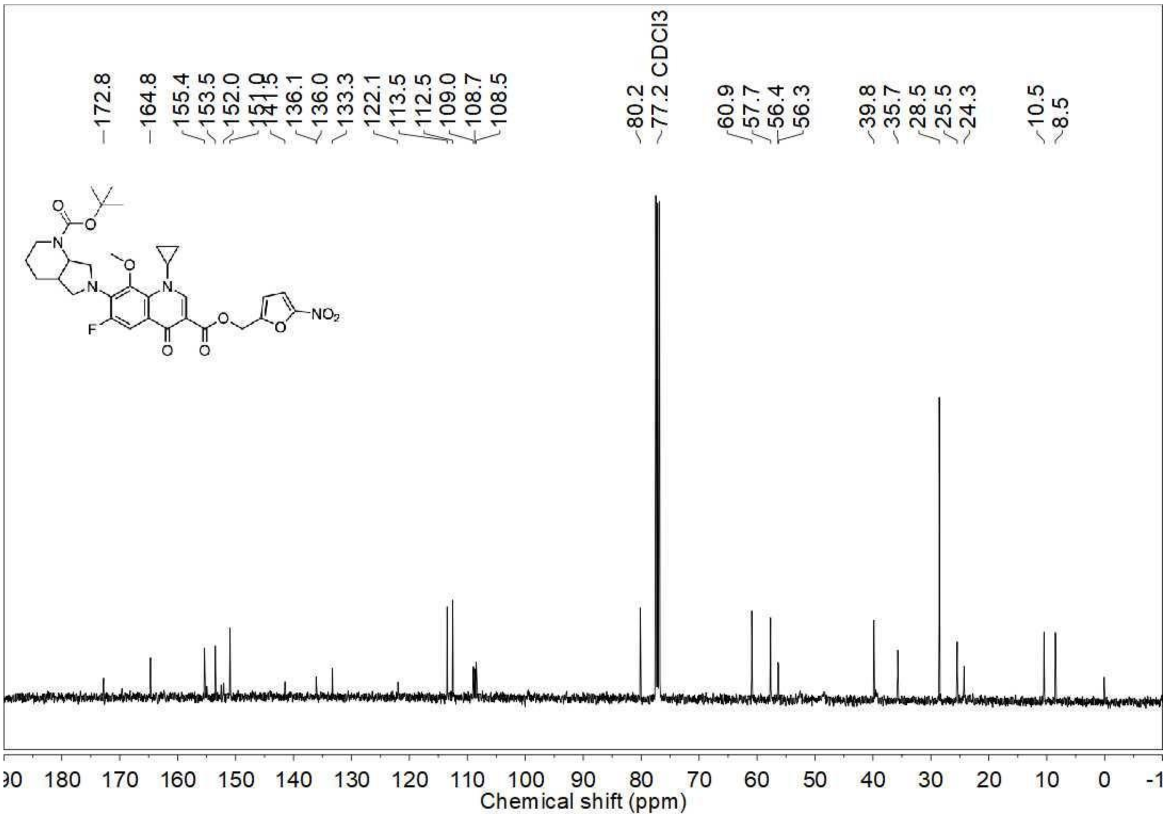

DEPT-135 NMR spectrum (100 MHz, $CDCl_3$) of **3b**

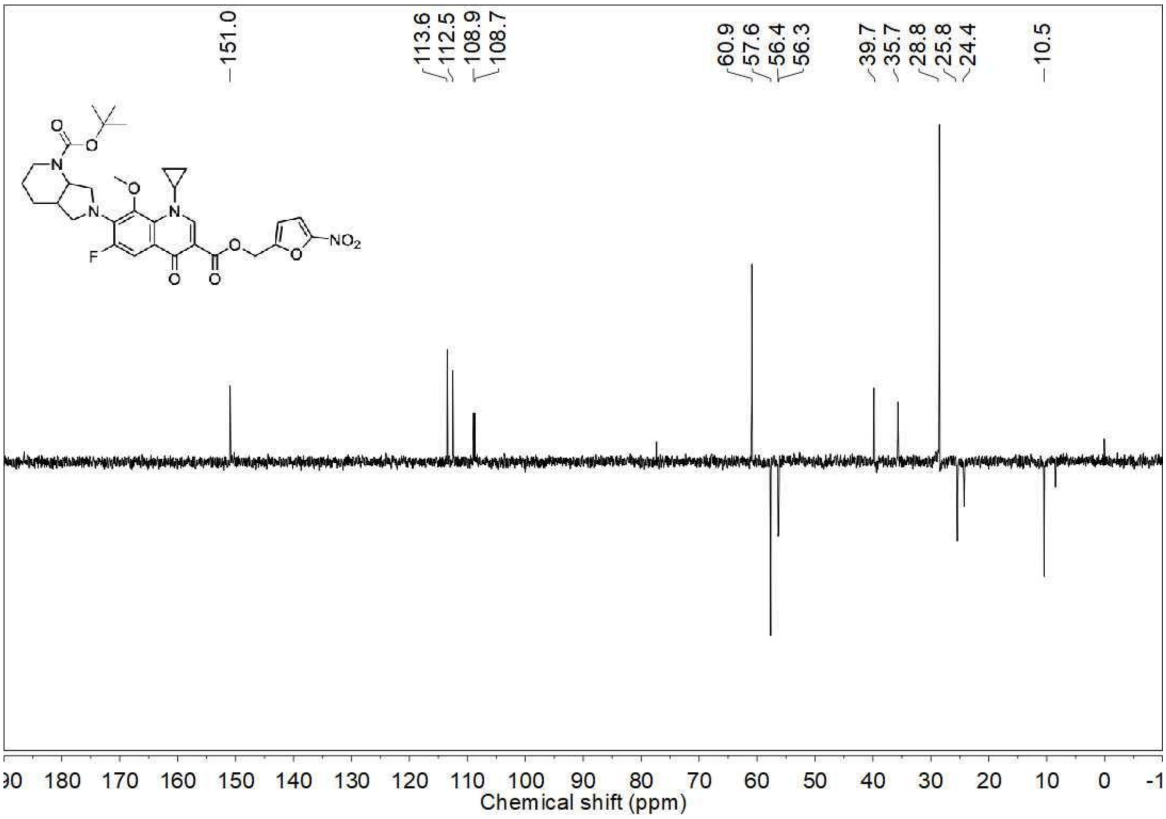


$^{19}$F NMR spectrum (376 MHz, $CDCl_3$) of **3c**

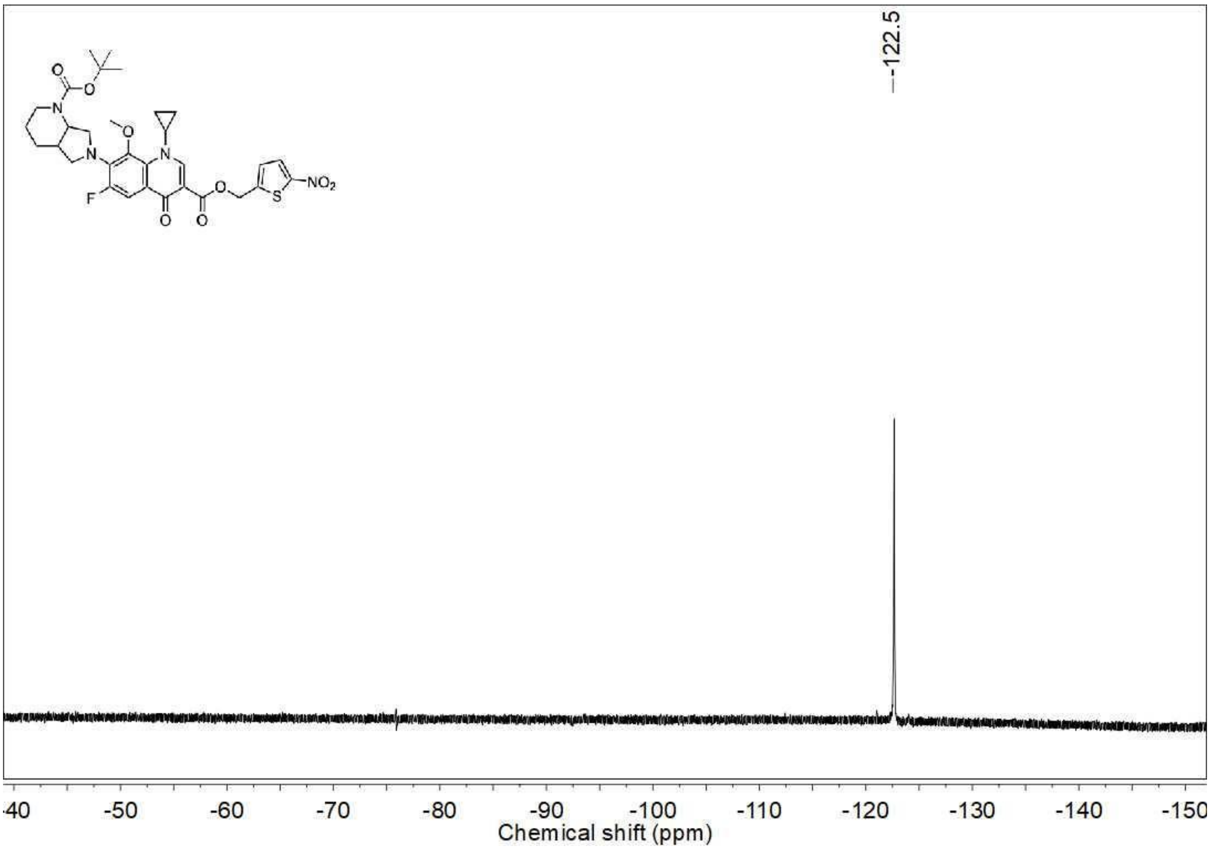

$^1$H NMR spectrum (400 MHz, $CDCl_3$) of **3c**

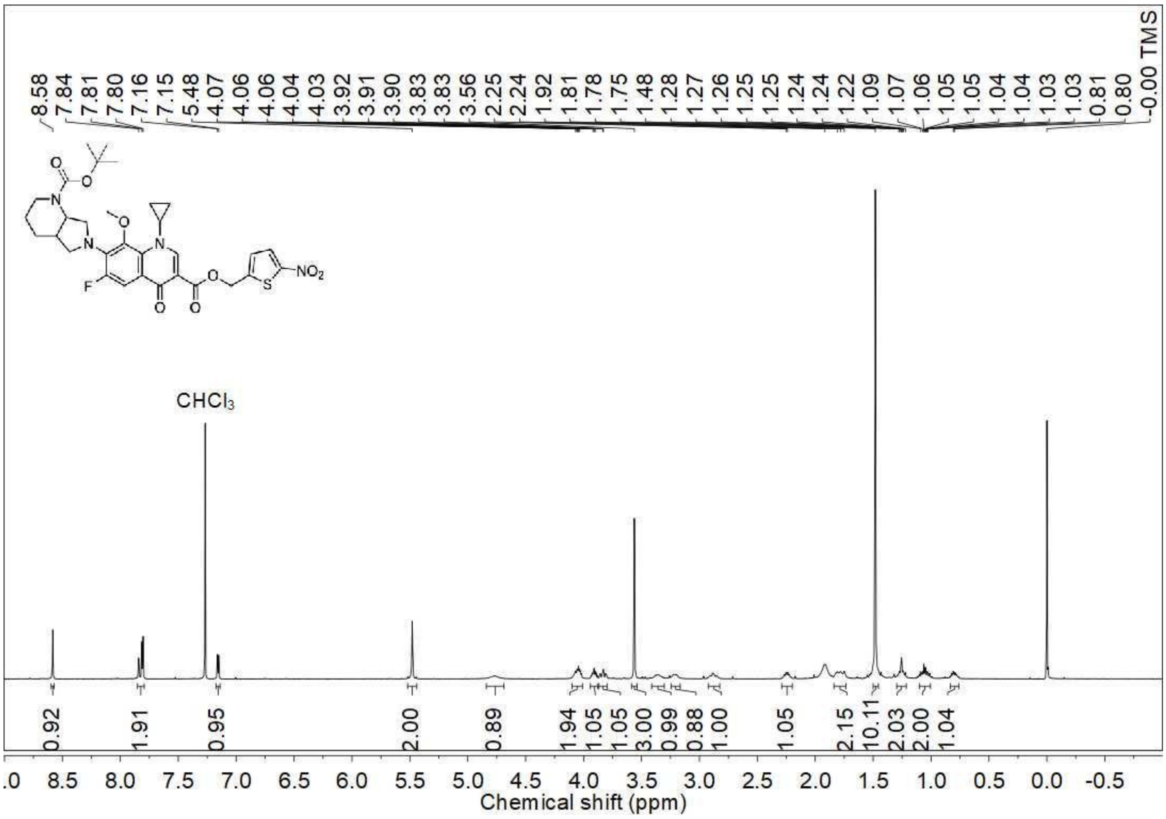


$^{13}$C NMR spectrum (100 MHz, $CDCl_3$) of **3c**

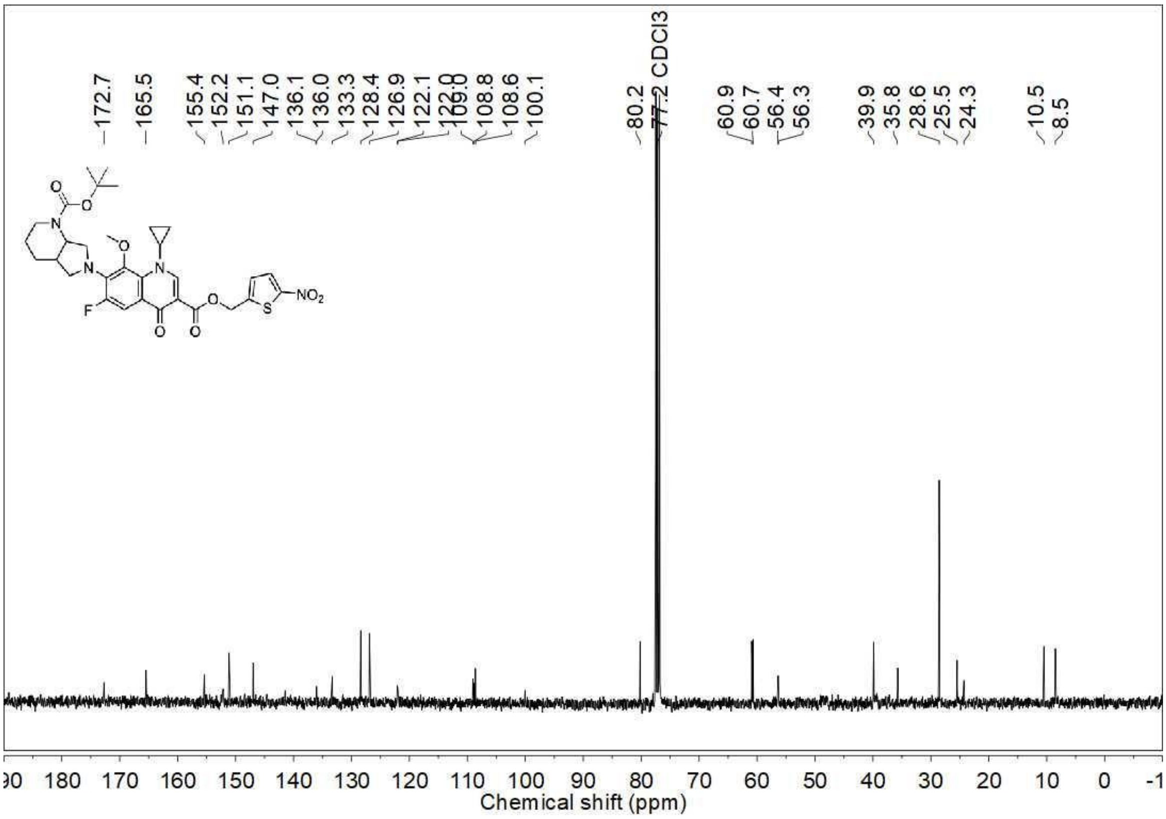

DEPT-135 NMR spectrum (100 MHz, $CDCl_3$) of **3c**

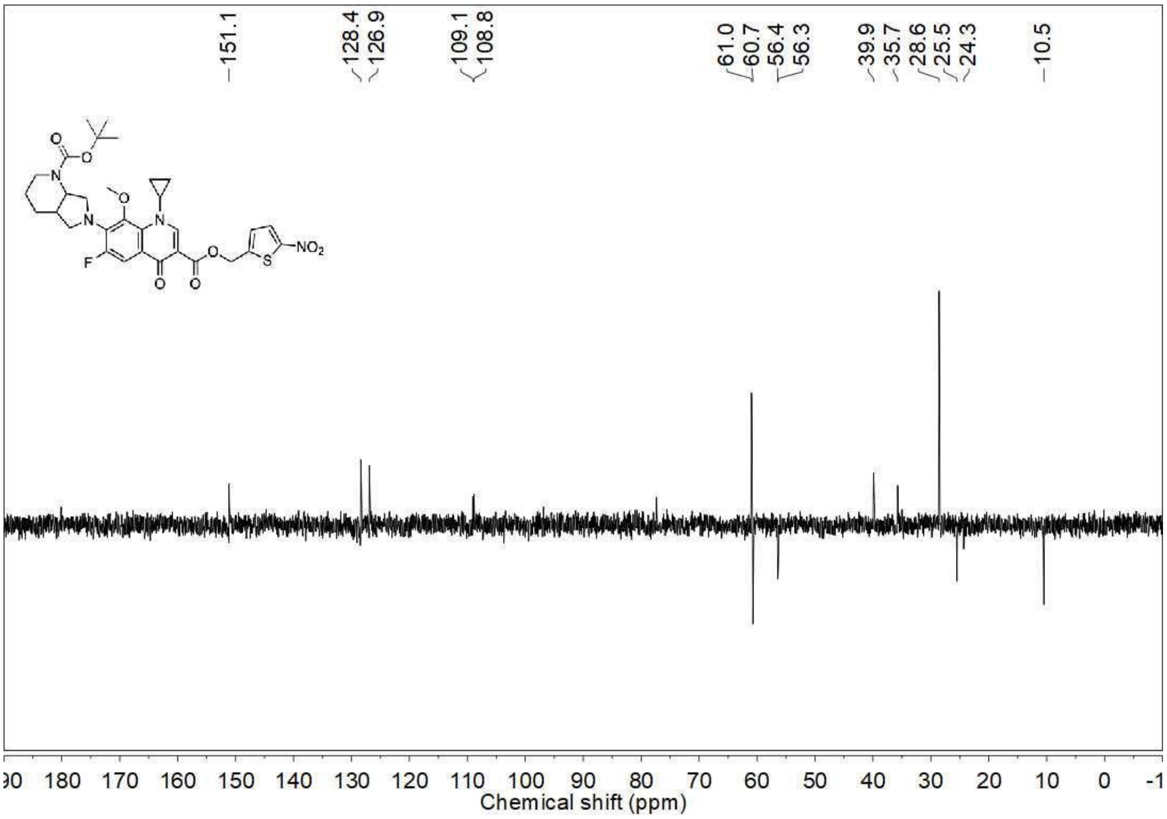


$^{19}F$ NMR spectrum (376 MHz, $CDCl_3$) of **3d**

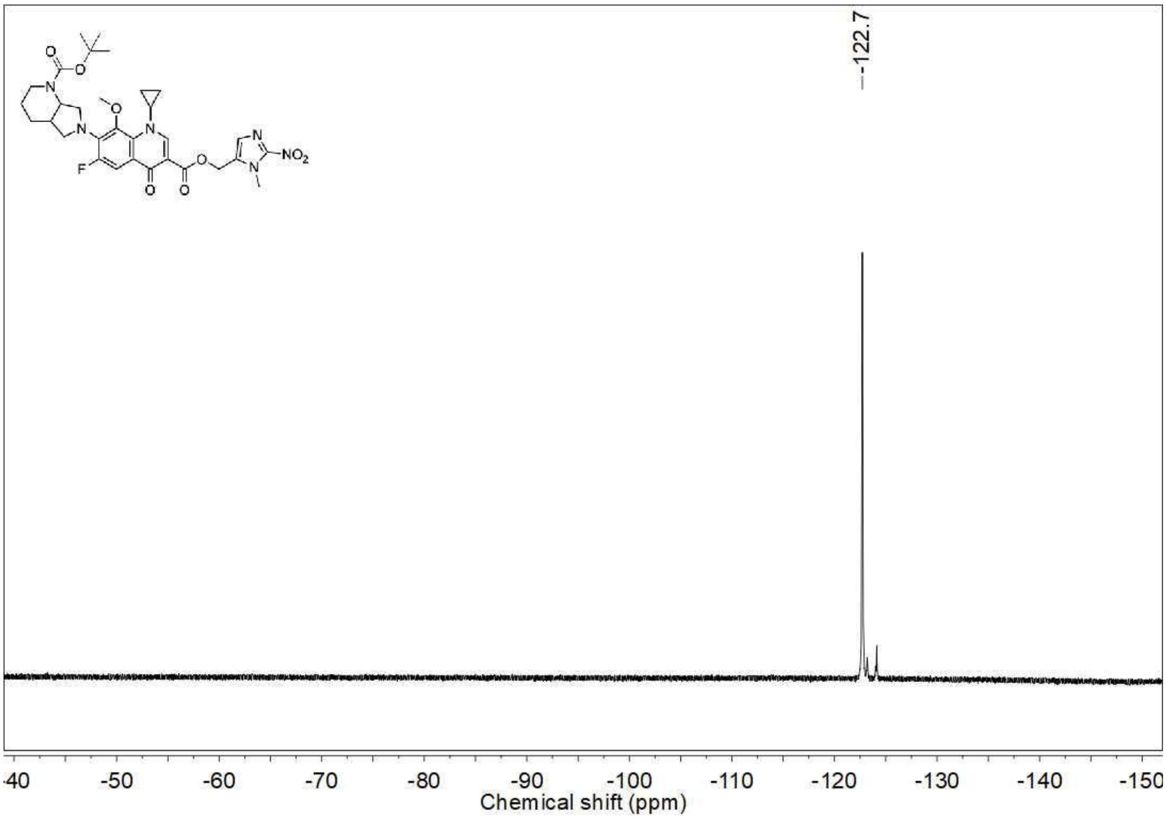

[1]H NMR spectrum (400 MHz, $CDCl_3$) of **3d**

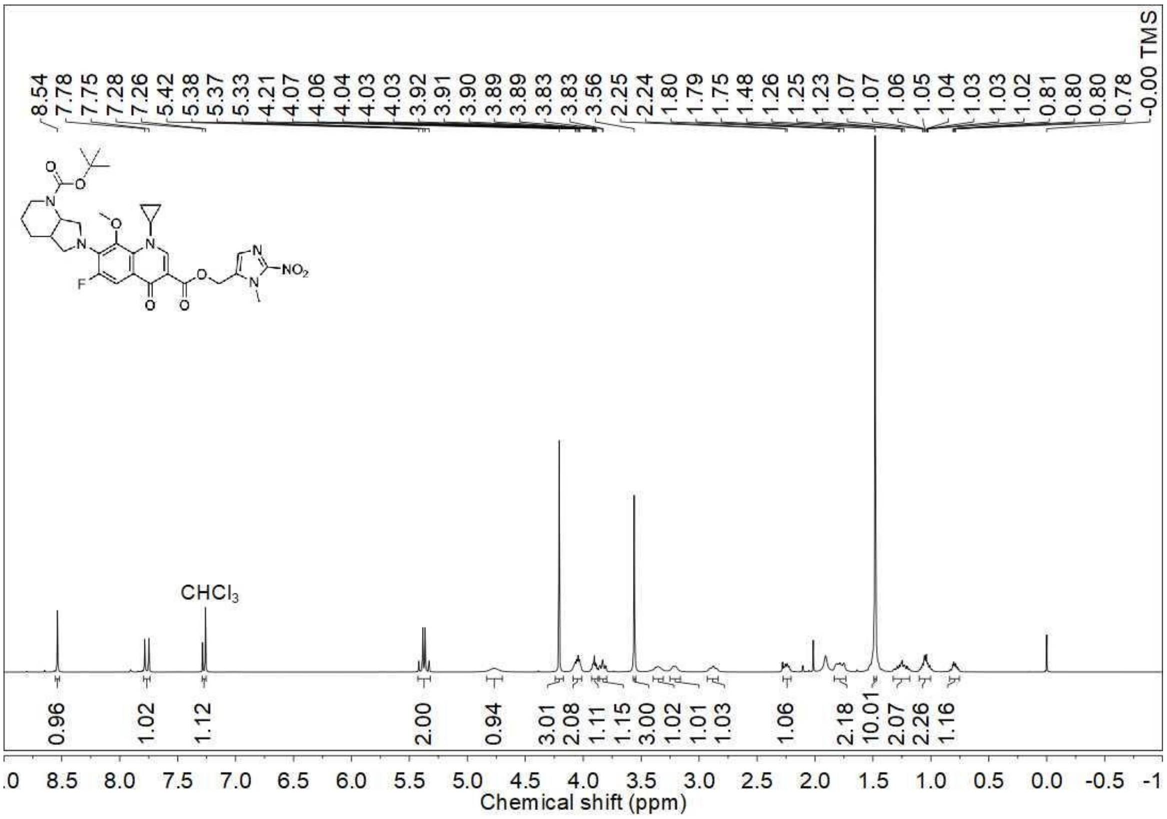

[13]C NMR spectrum (100 MHz, $CDCl_3$) of **3d**

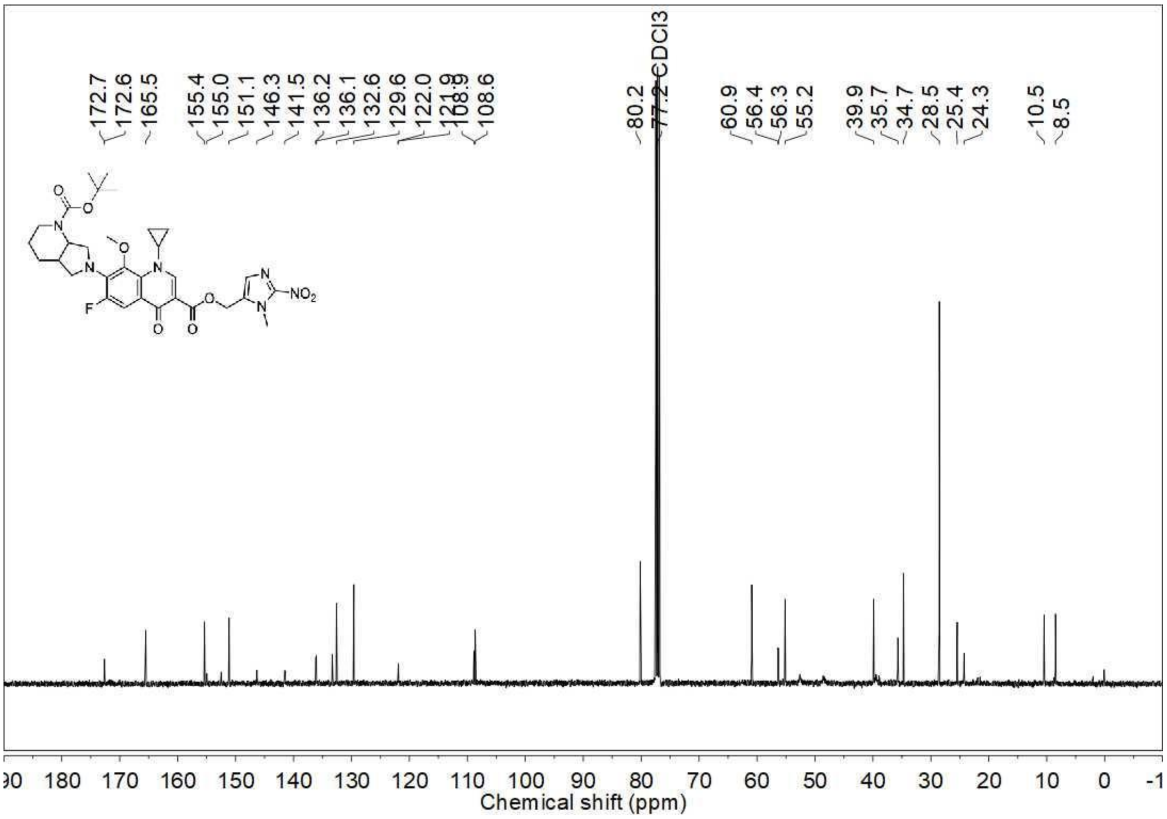

DEPT-135 NMR spectrum (100 MHz, $CDCl_3$) of **3d**

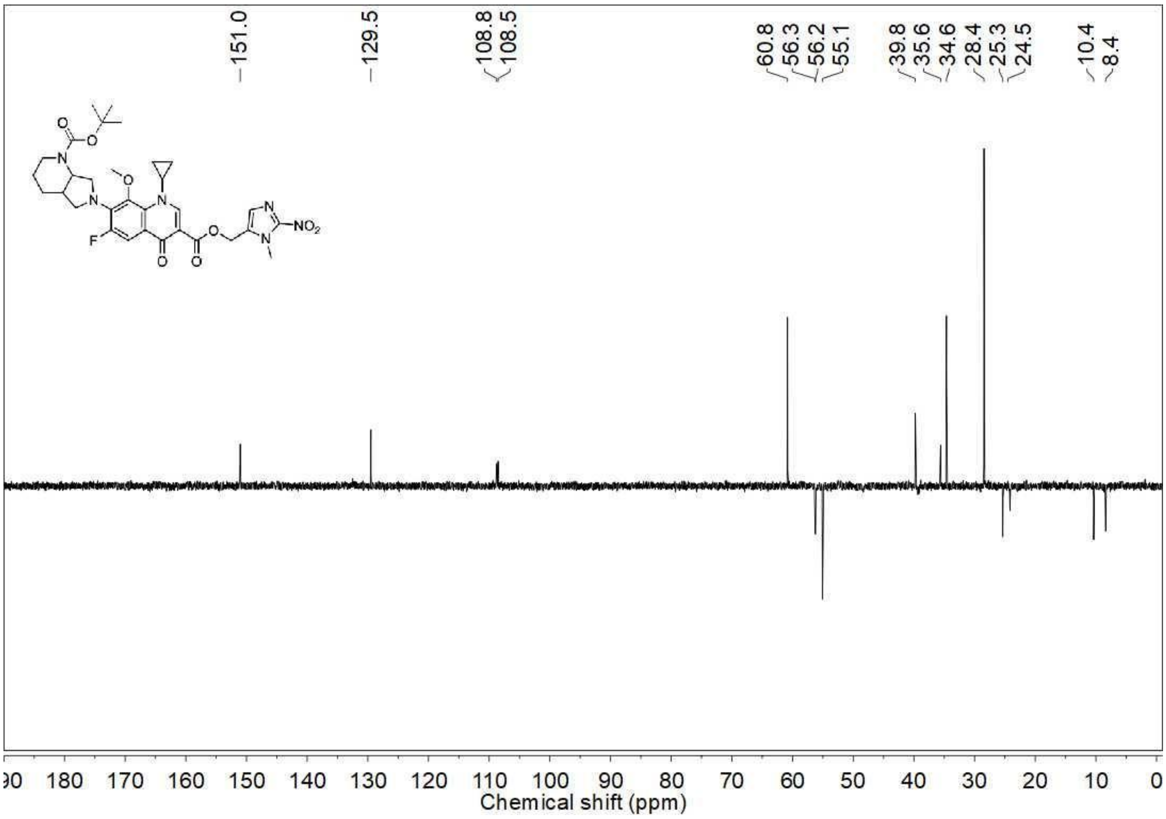


$^{19}$F NMR spectrum (376 MHz, $CDCl_3$) of **3e**

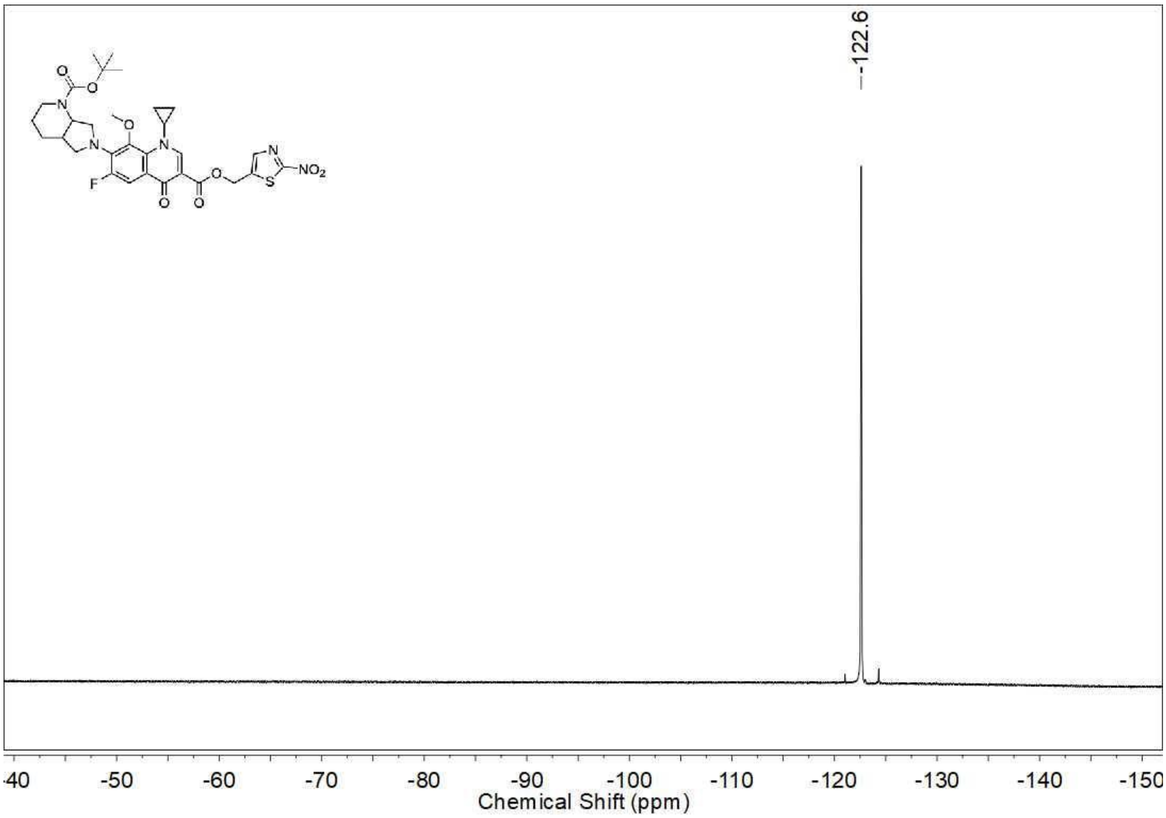

$^{1}H$ NMR spectrum (400 MHz, $CDCl_3$) of **3e**

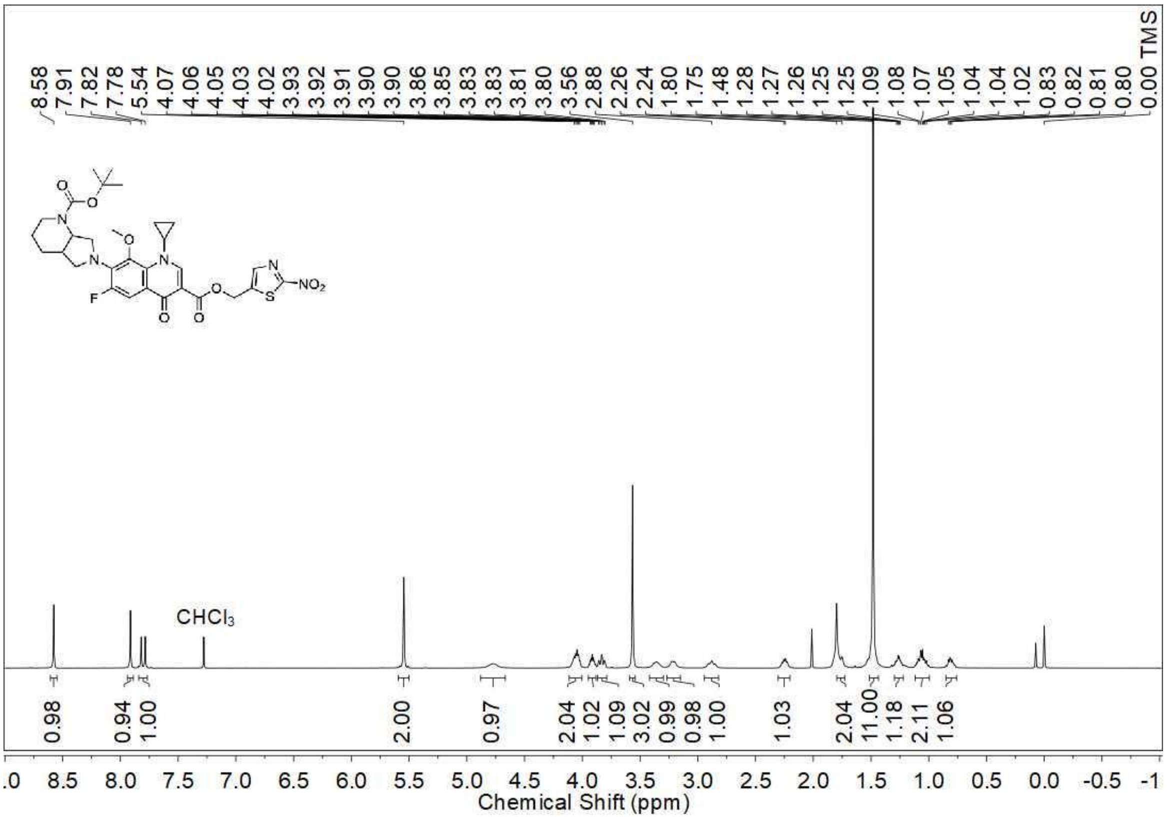


$^{13}C$ NMR spectrum (100 MHz, $CDCl_3$) of **3e**

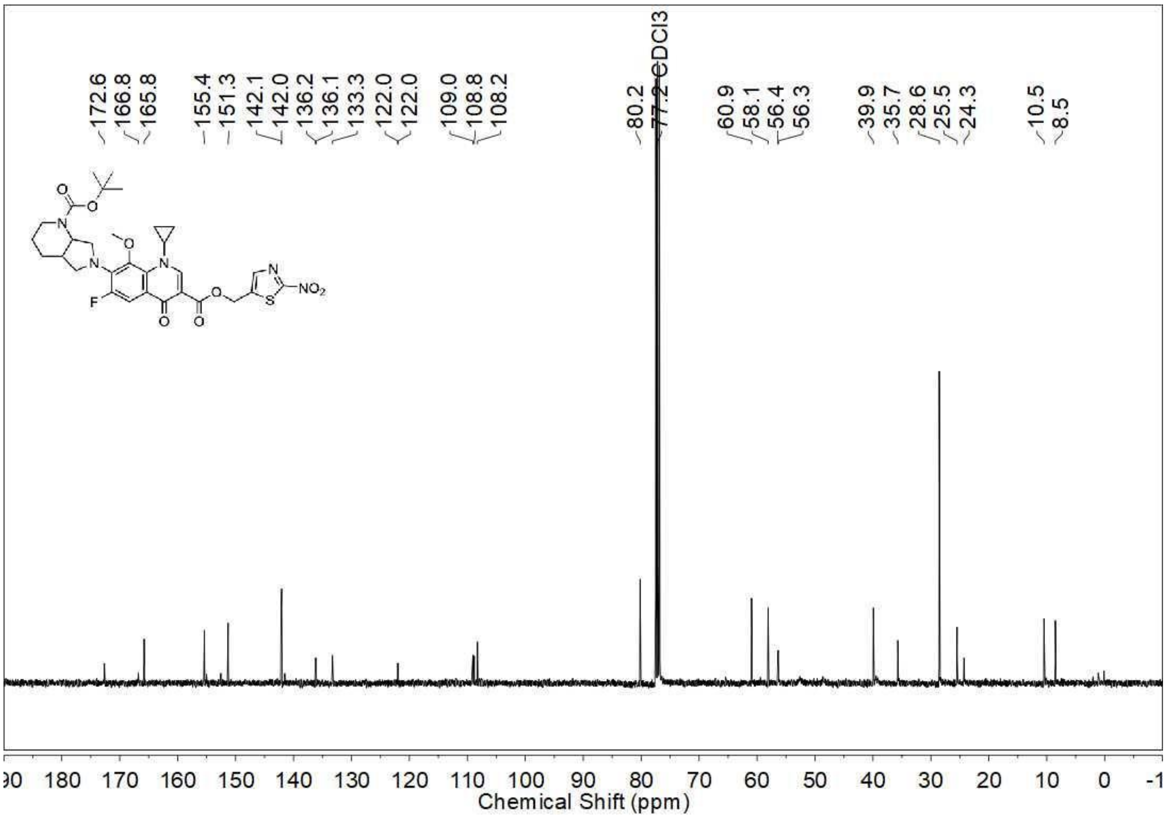

DEPT-135 NMR spectrum (100 MHz, $CDCl_3$) of **3e**

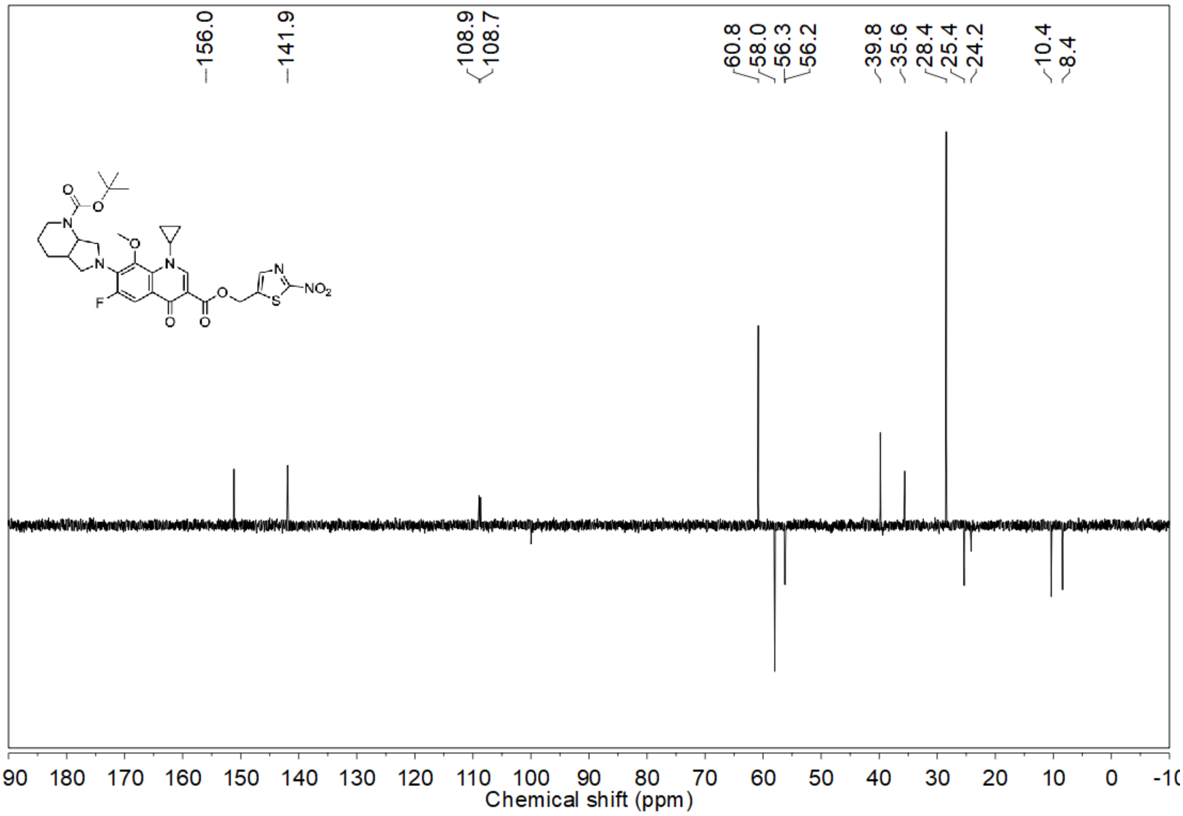


[19]F NMR spectrum (376 MHz, $CDCl_3$) of **3f**

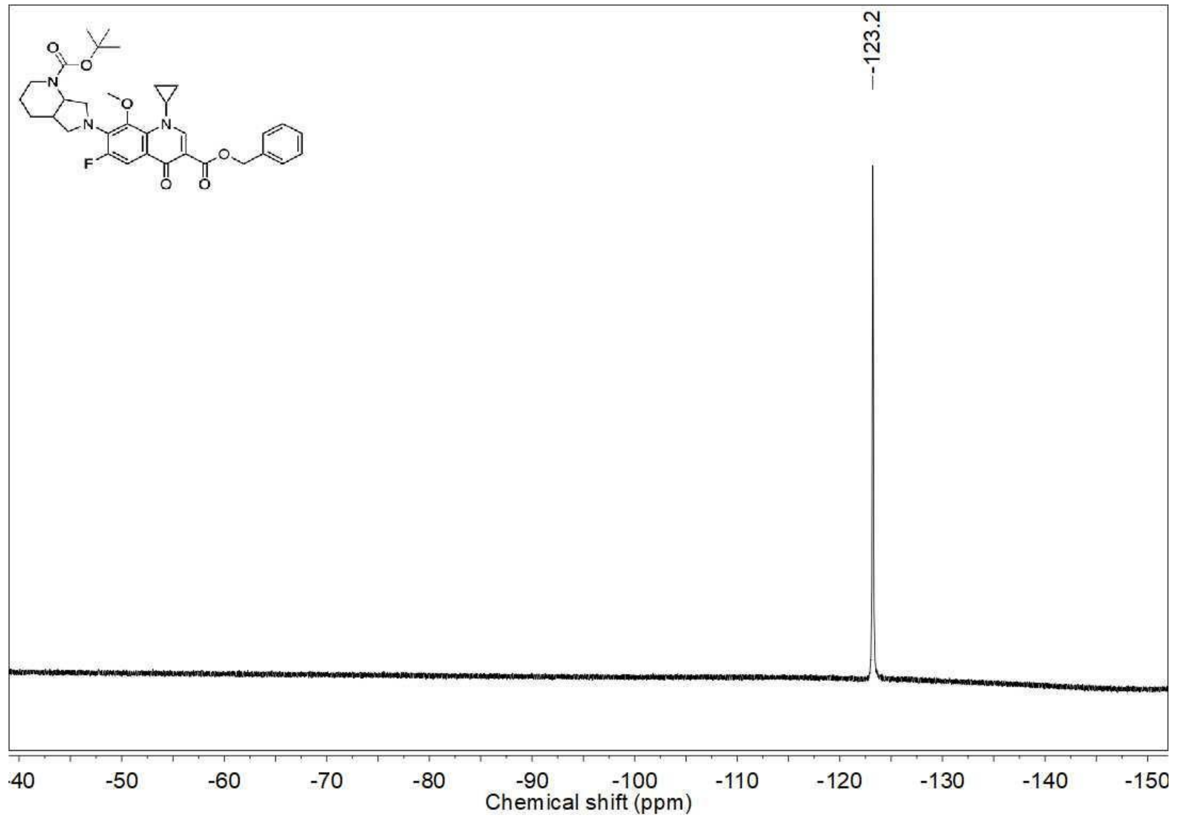

$^{1}$H NMR spectrum (400 MHz, $CDCl_3$) of **3f**

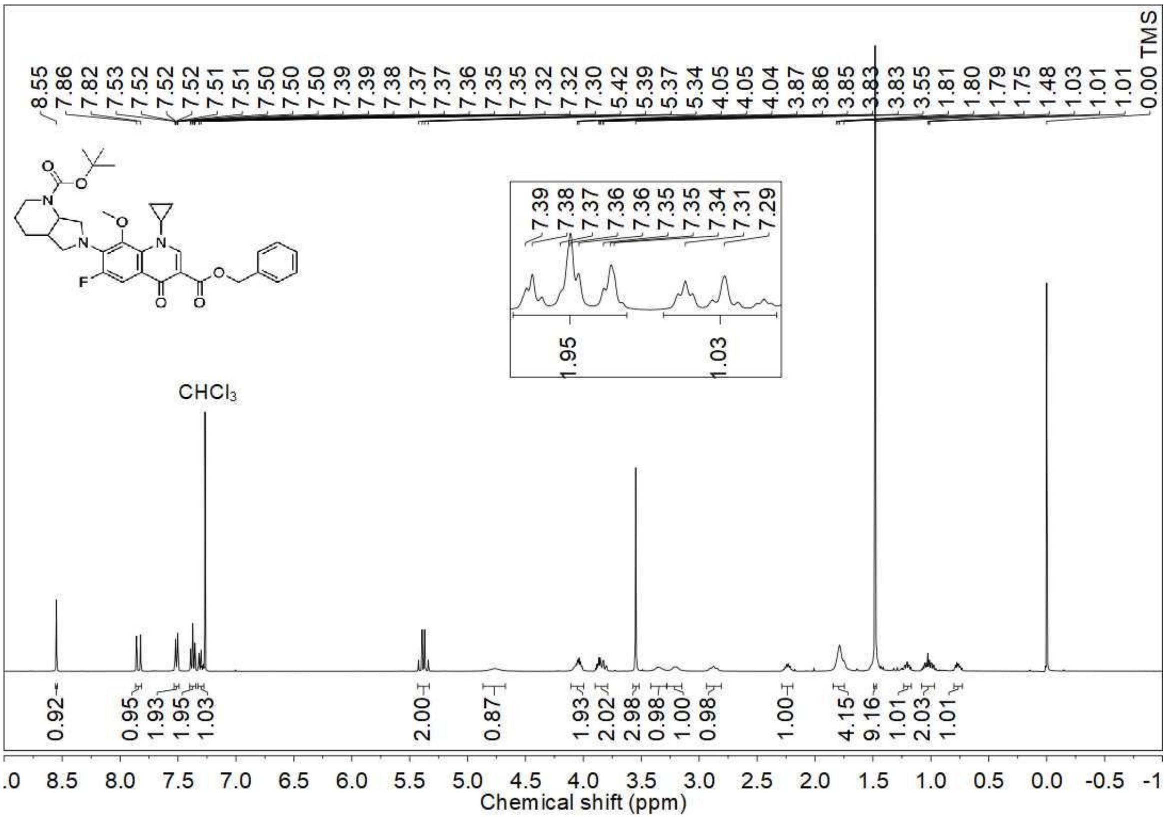


$^{13}$C NMR spectrum (100 MHz, $CDCl_3$) of **3f**

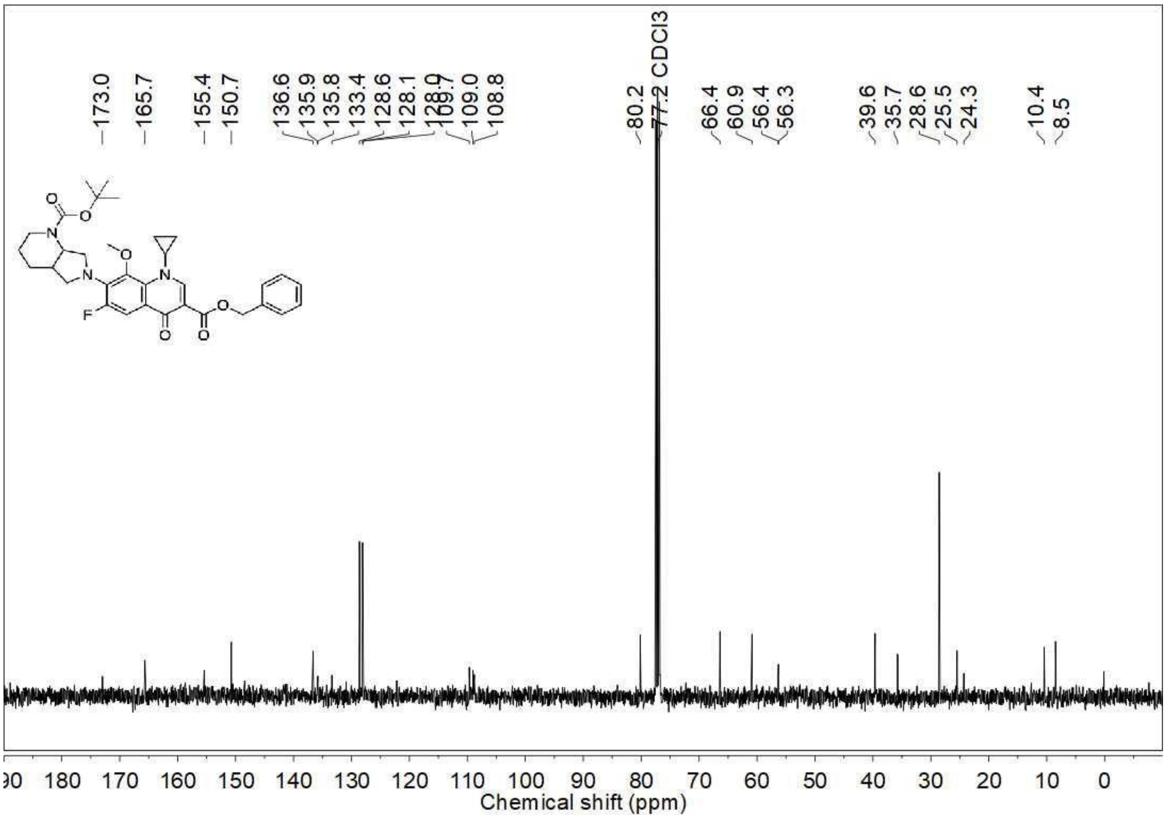

DEPT-135 NMR spectrum (100 MHz, $CDCl_3$) of **3f**

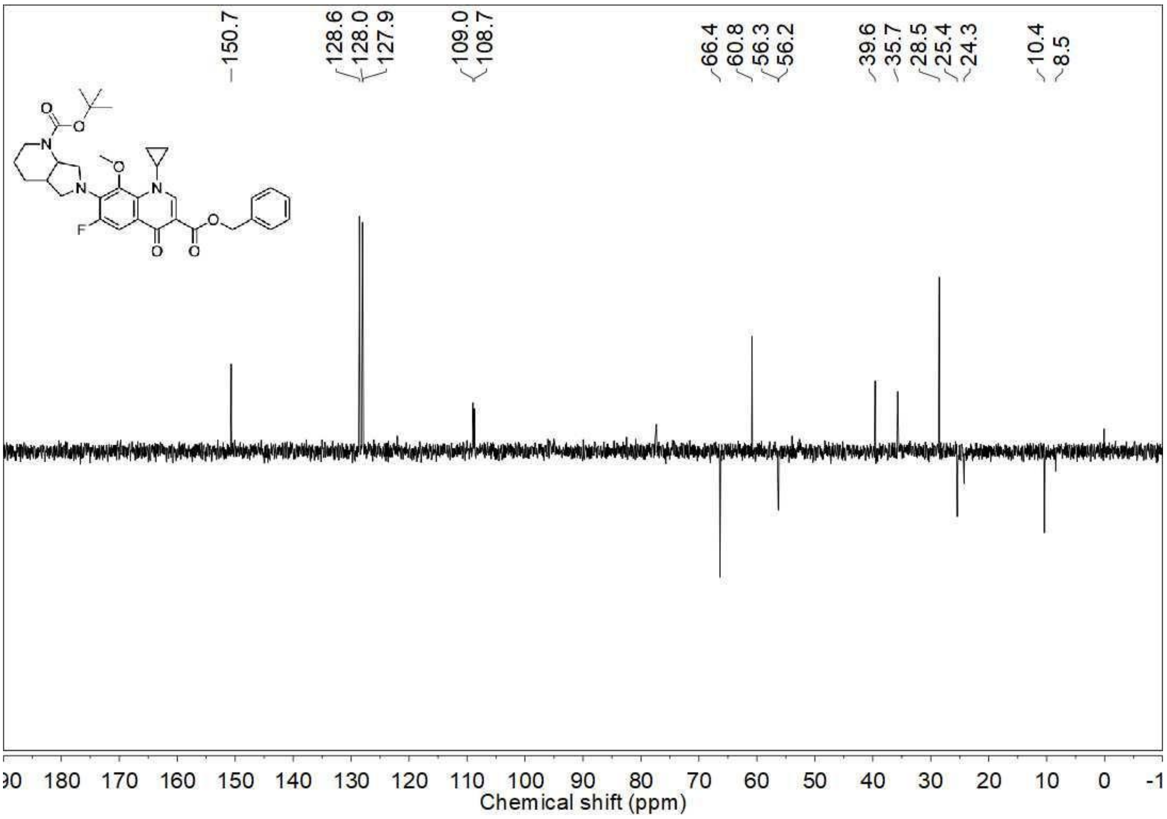


$^{19}F$ NMR spectrum (376 MHz, $CDCl_3$) of **3g**

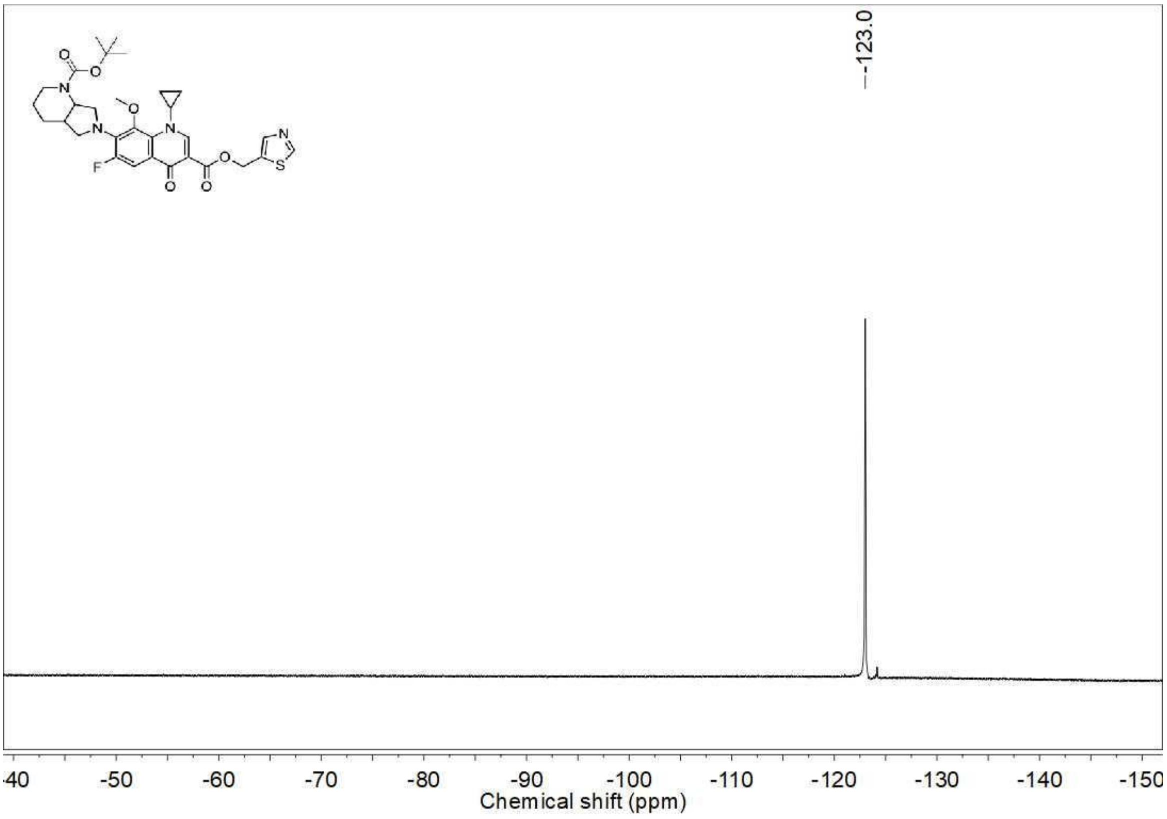

[1]H NMR spectrum (400 MHz, $CDCl_3$) of **3g**

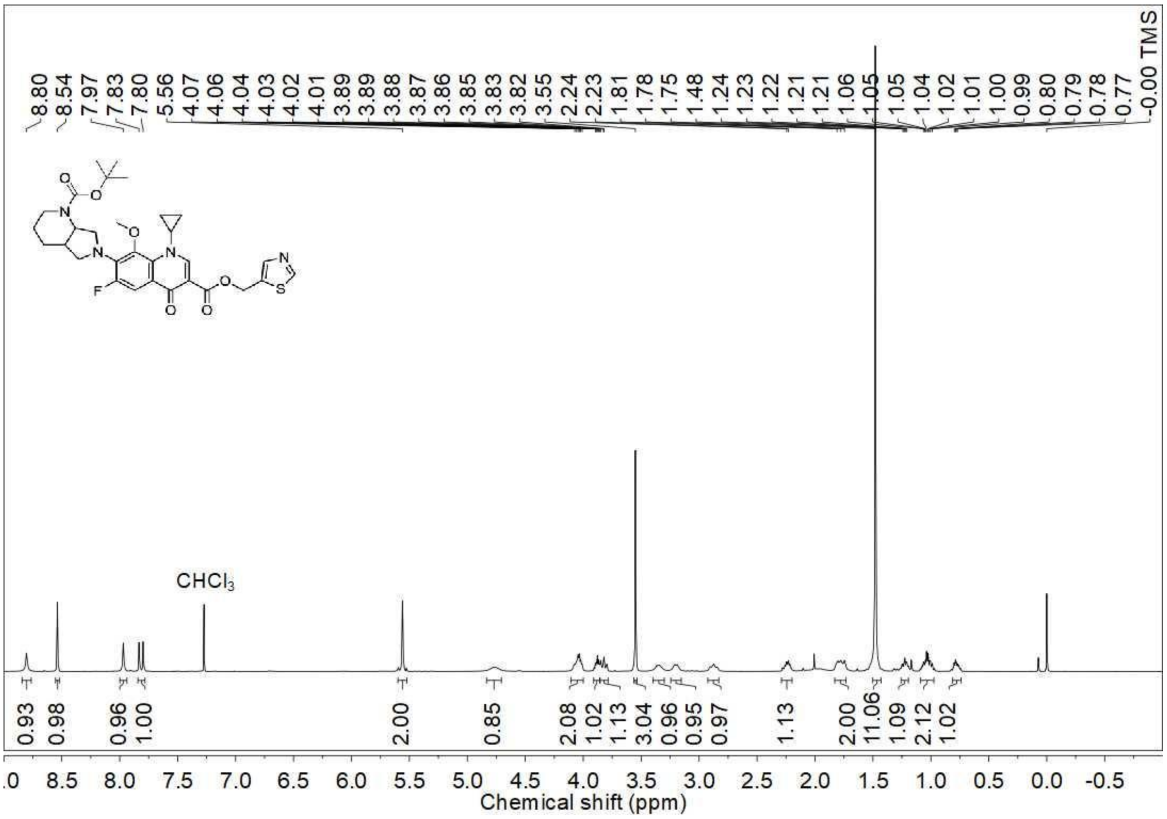

[13]C NMR spectrum (100 MHz, $CDCl_3$) of **3g**

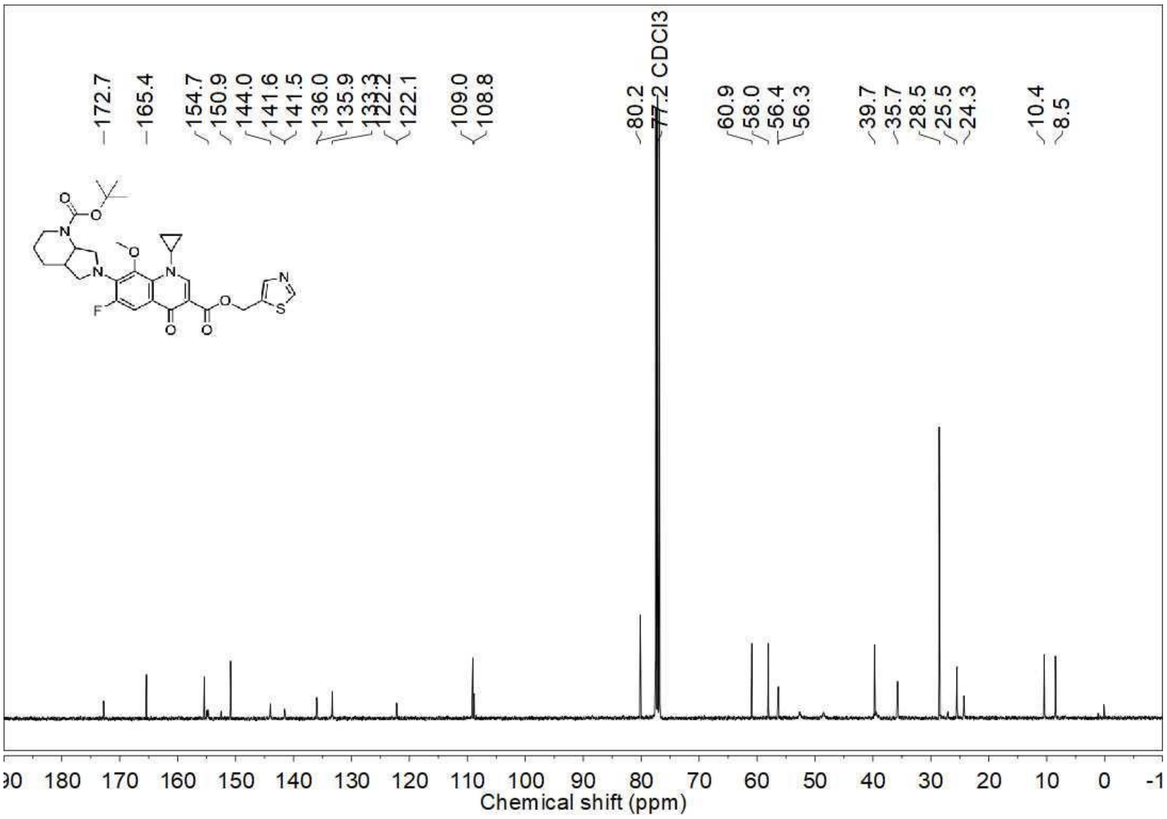

DEPT-135 NMR spectrum (100 MHz, $CDCl_3$) of **3g**

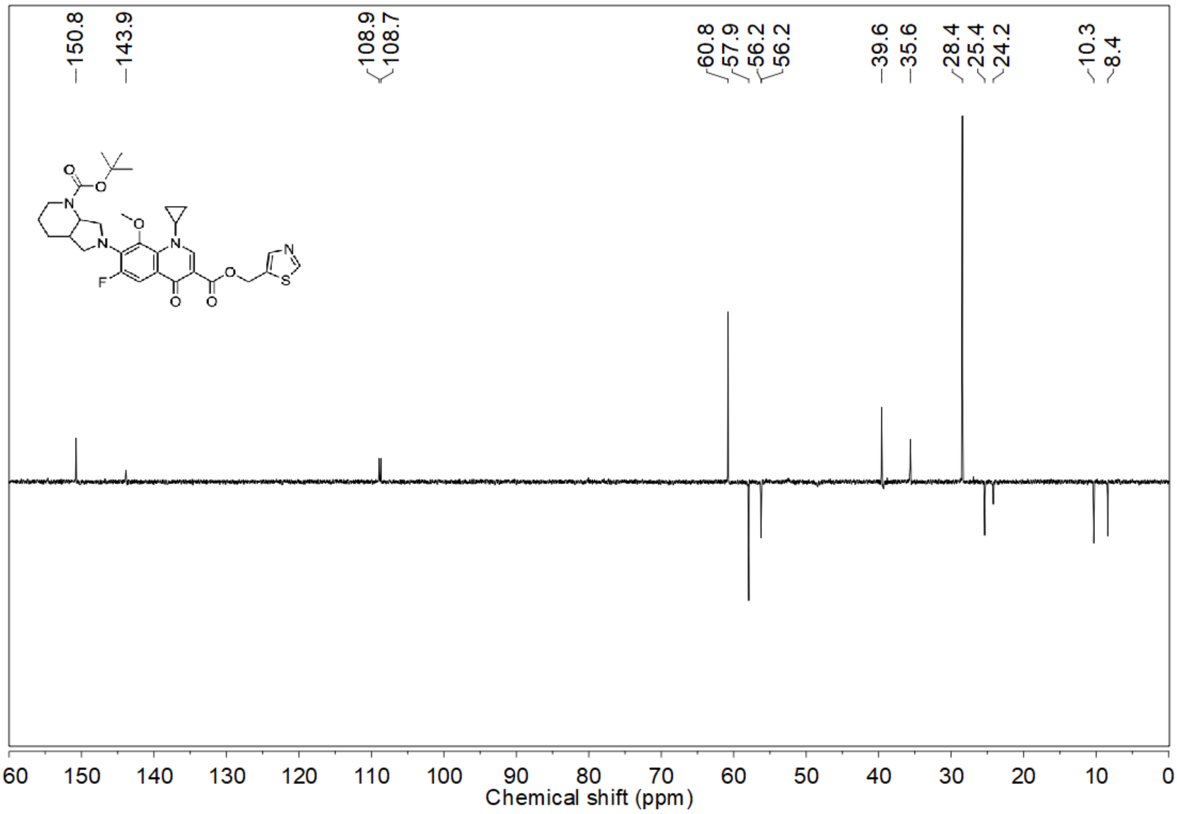


$^{19}F$ NMR spectrum (376 MHz, $CDCl_3$) of **4**

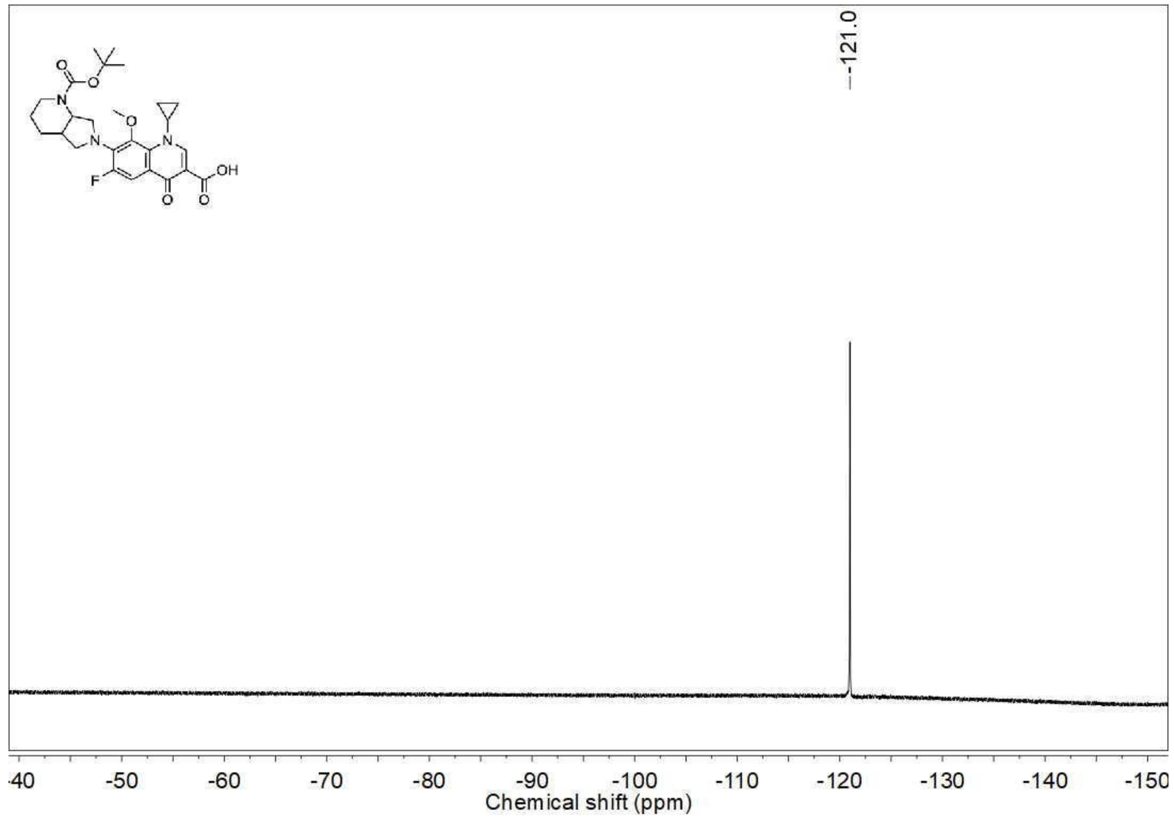

$^{1}$H NMR spectrum (400 MHz, $CDCl_3$) of **4**

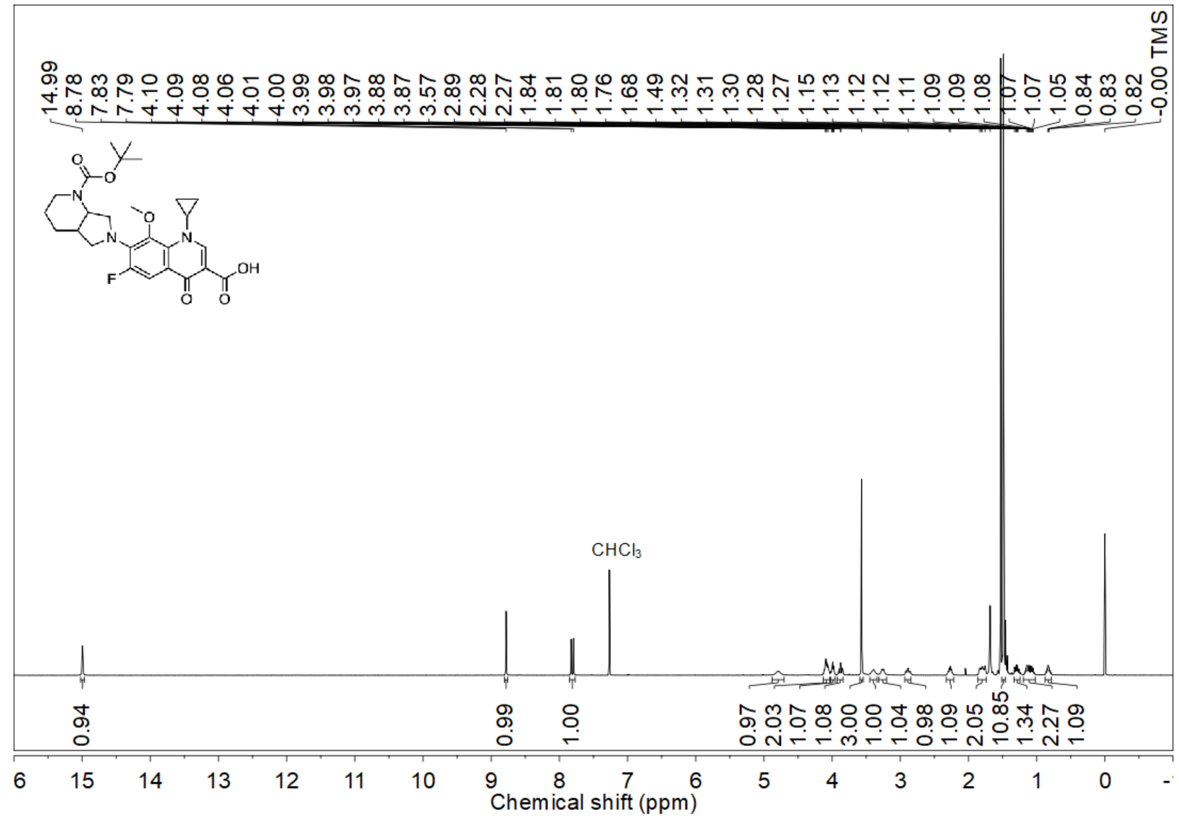


$^{1}$H NMR spectrum (400 MHz, $CDCl_3$) of **5a**

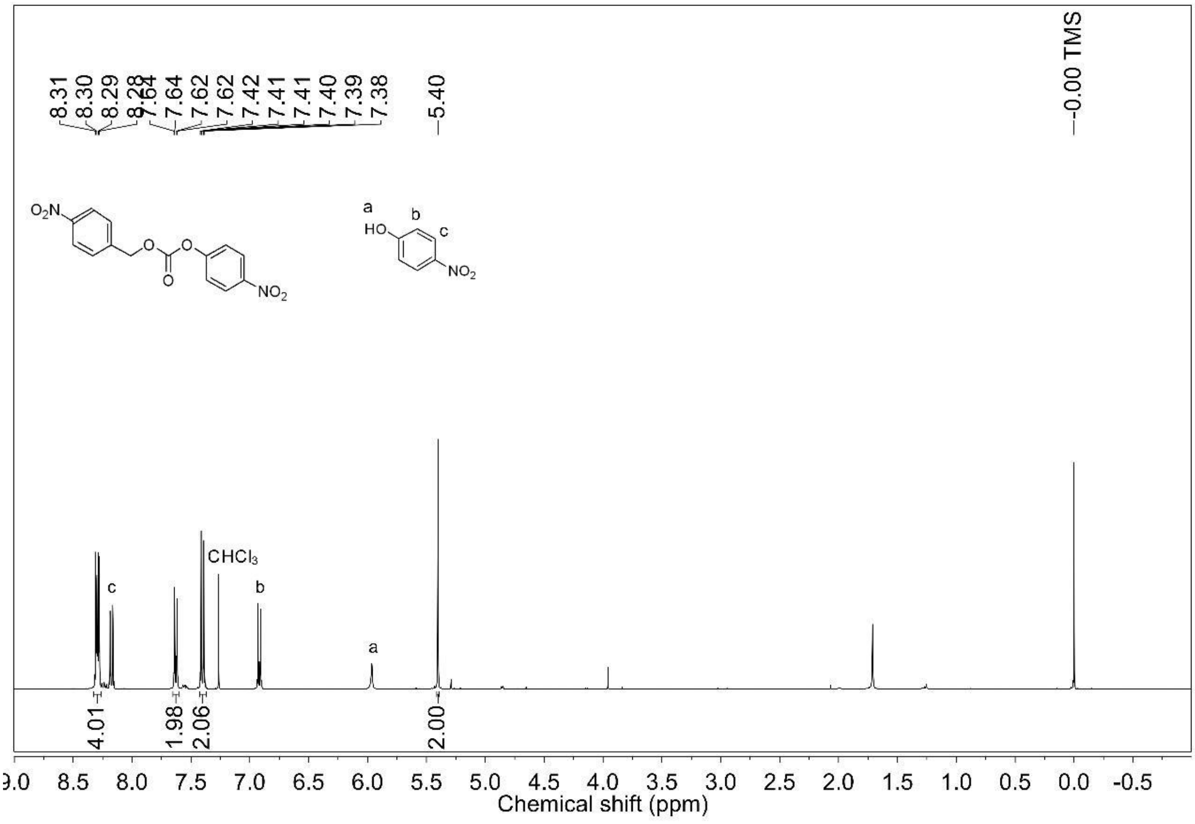

$^1$H NMR spectrum (400 MHz, $CDCl_3$) of **5b**

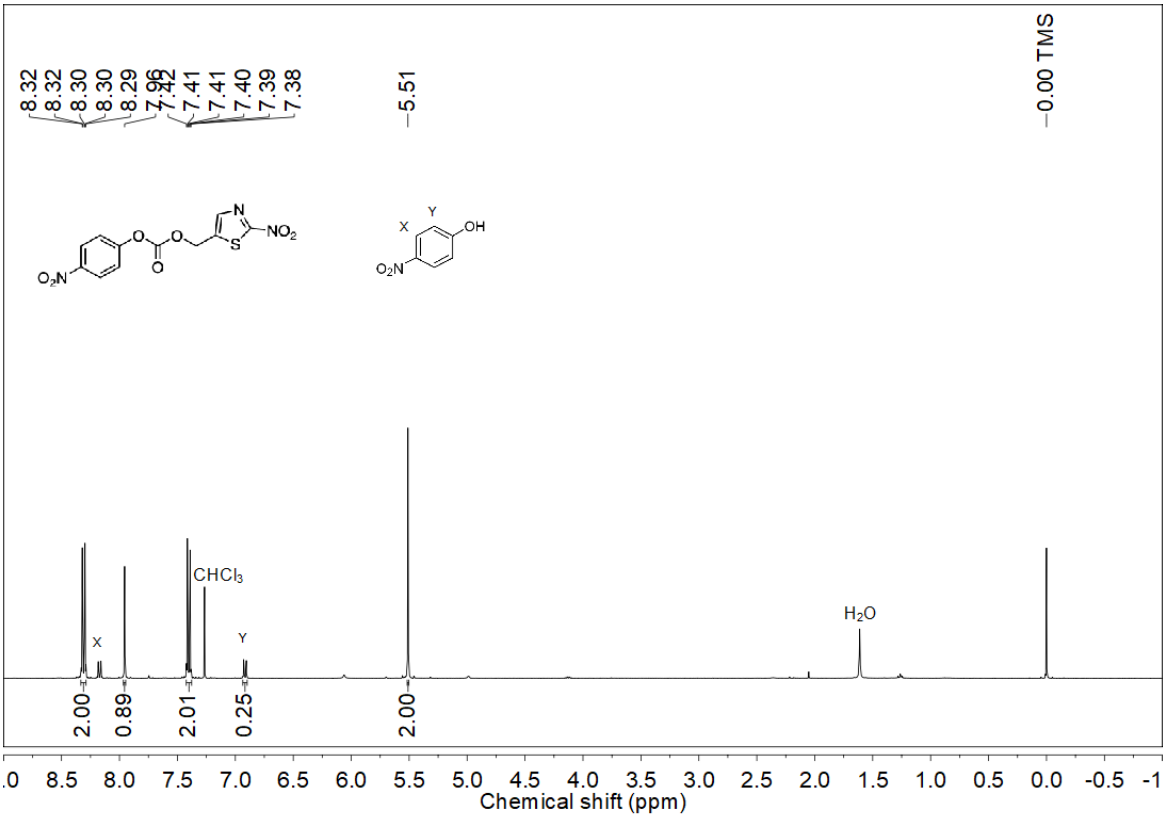


$^1$H NMR spectrum (400 MHz, $CDCl_3$) of **6b**

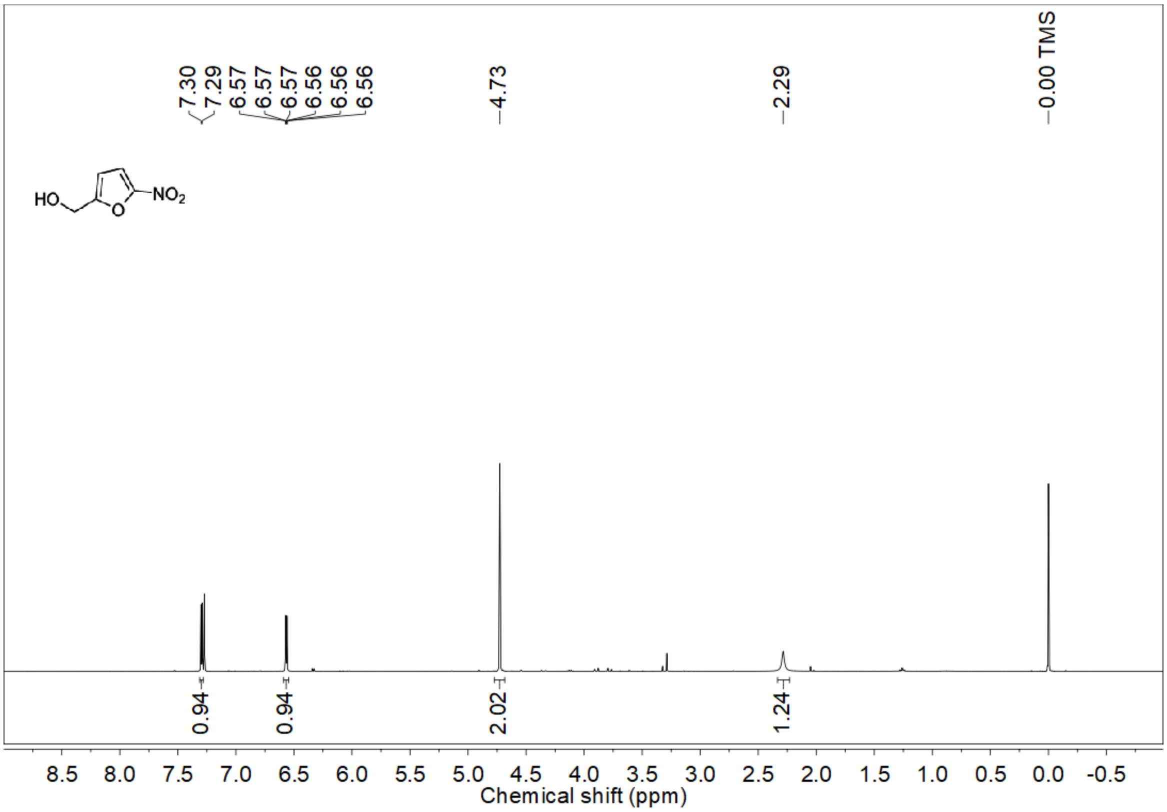

[1]H NMR spectrum (400 MHz, $CDCl_3$) of **6c**

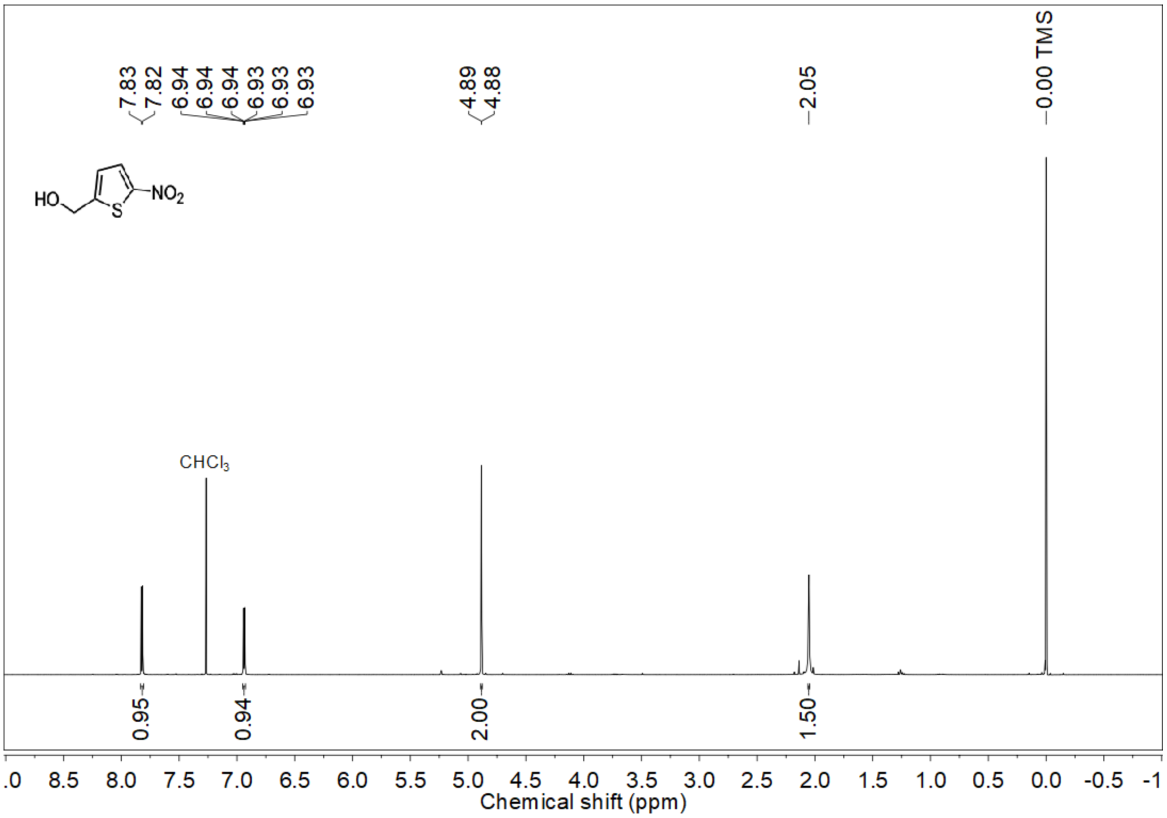


[1]H NMR spectrum (400 MHz, $CDCl_3$) of **6d**

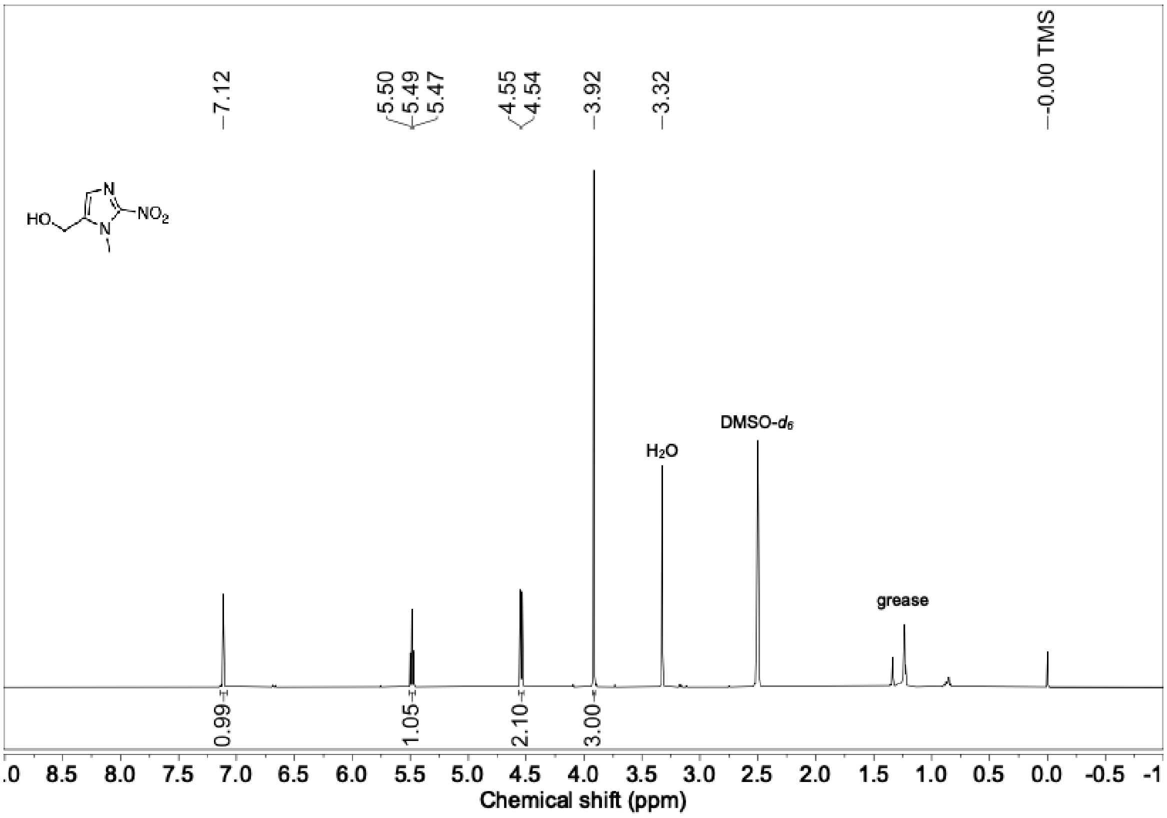

[1]H NMR spectrum (400 MHz, $CDCl_3$) of **6e**

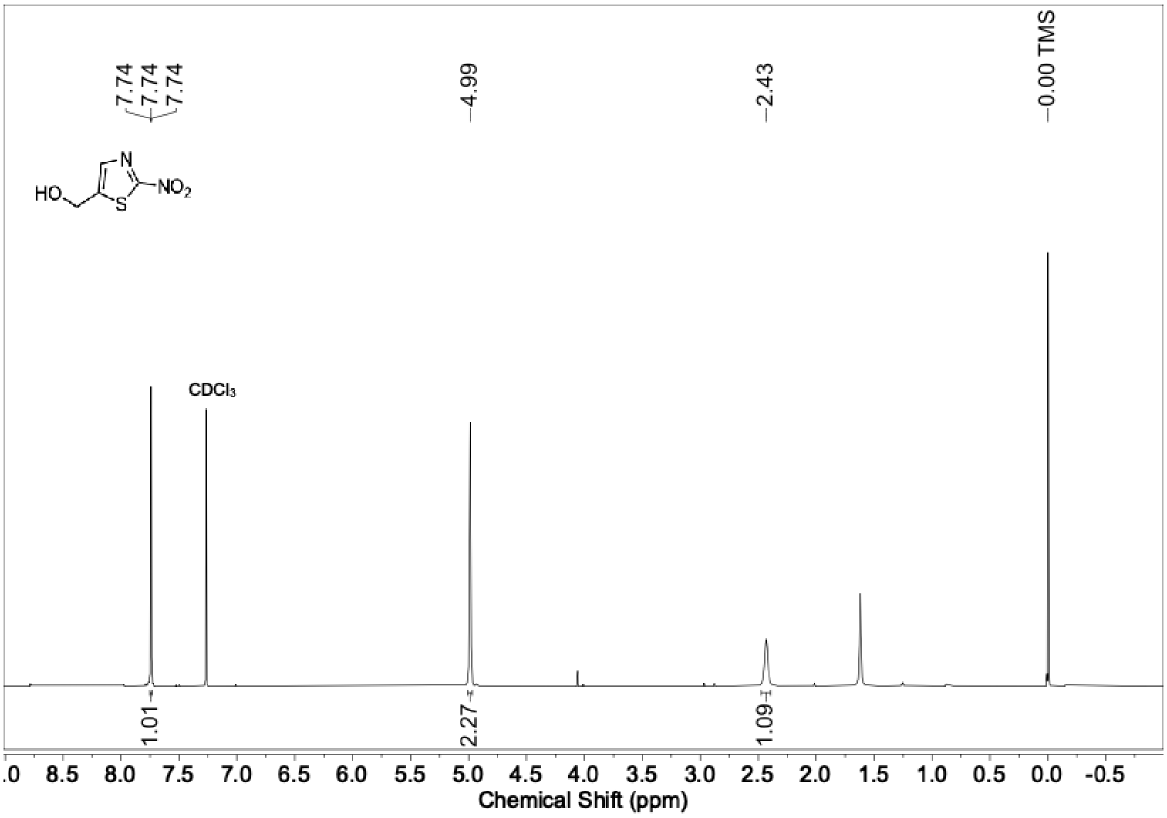


[1]H NMR spectrum (400 MHz, $CDCl_3$) of **7**

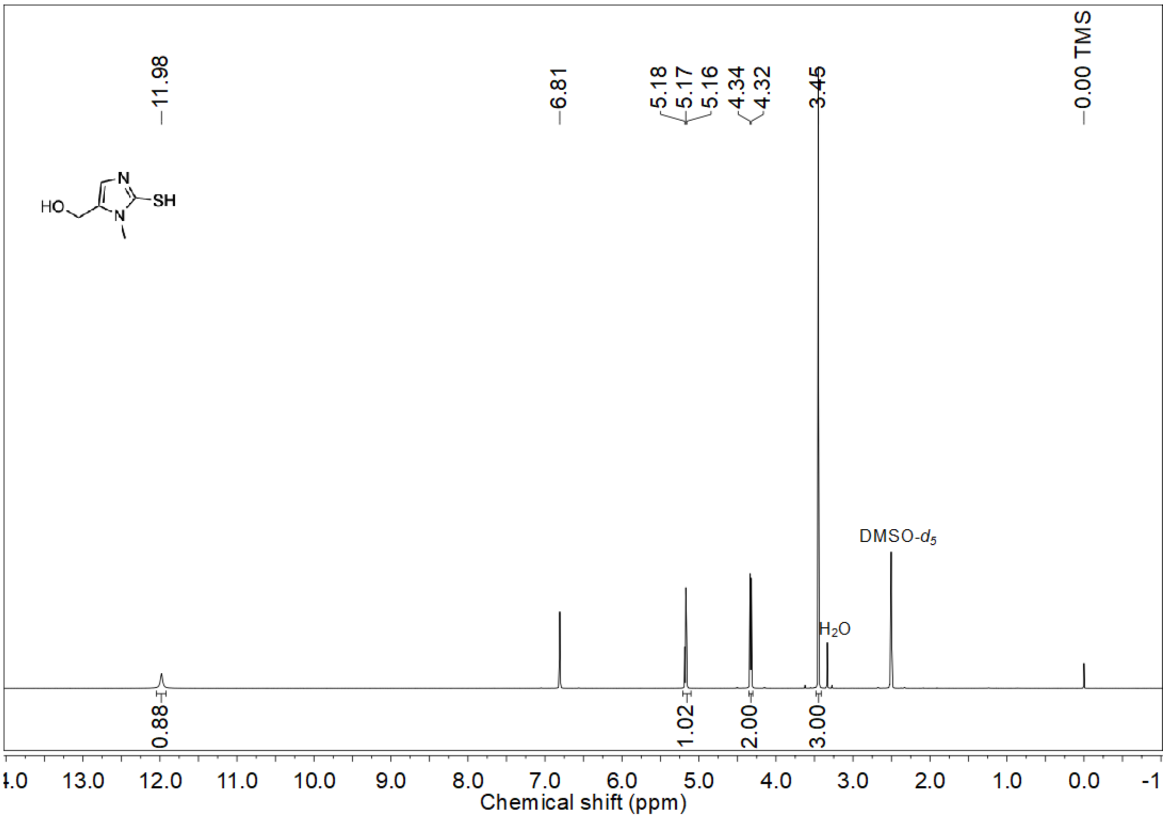

$^{1}H$ NMR spectrum (400 MHz, $CDCl_3$) of **8**

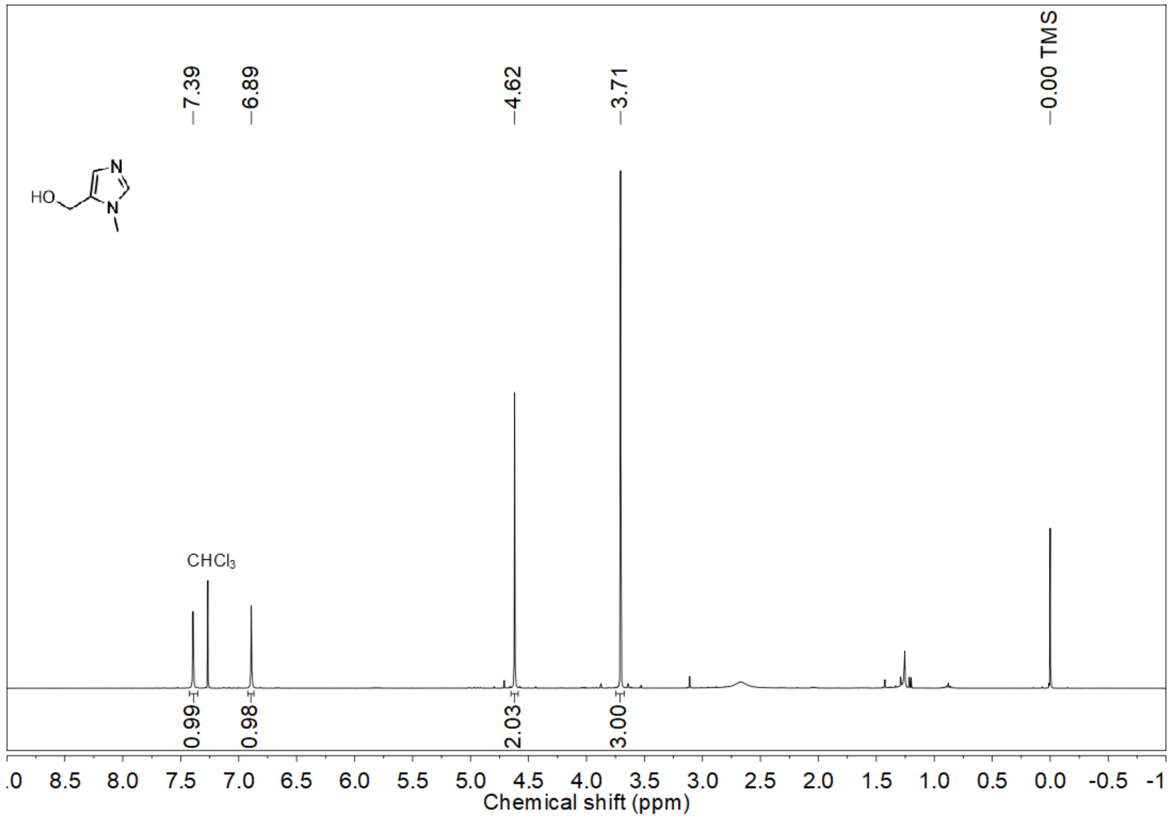


$^{1}H$ NMR spectrum (400 MHz, $CDCl_3$) of **9**

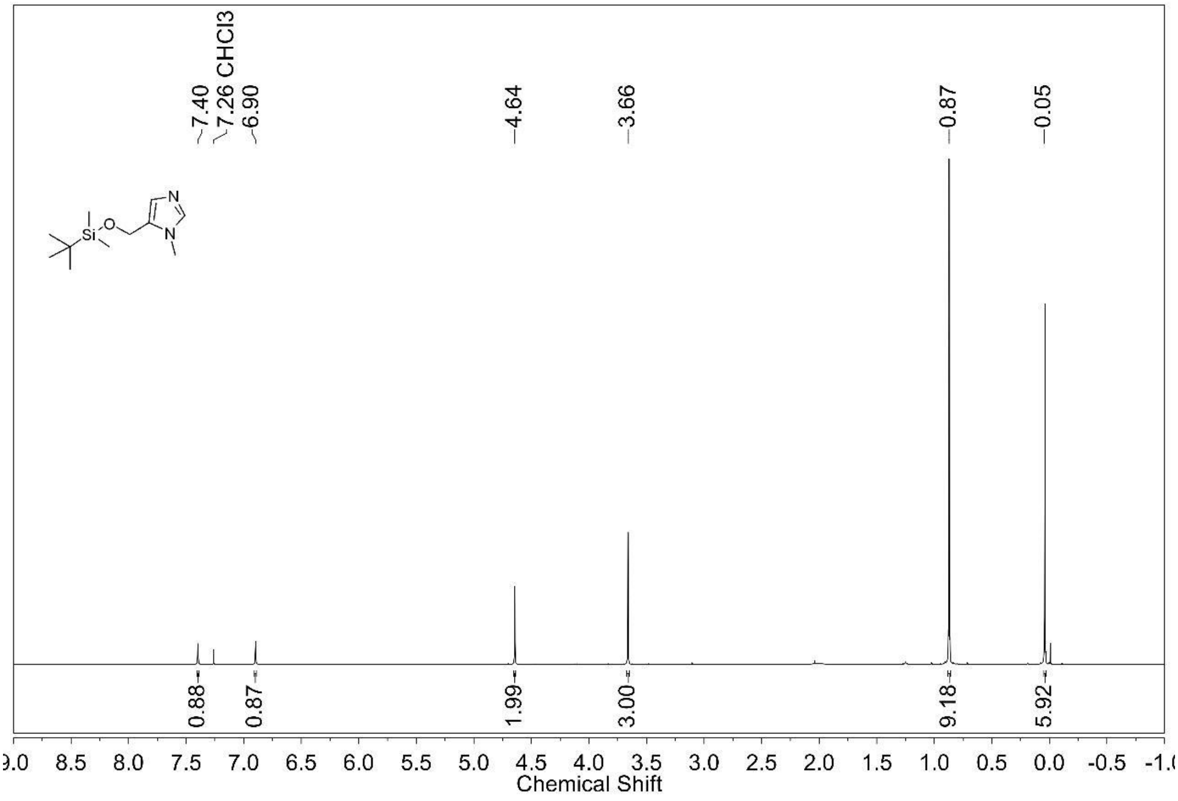

$^{1}H$ NMR spectrum (400 MHz, $CDCl_3$) of **10**

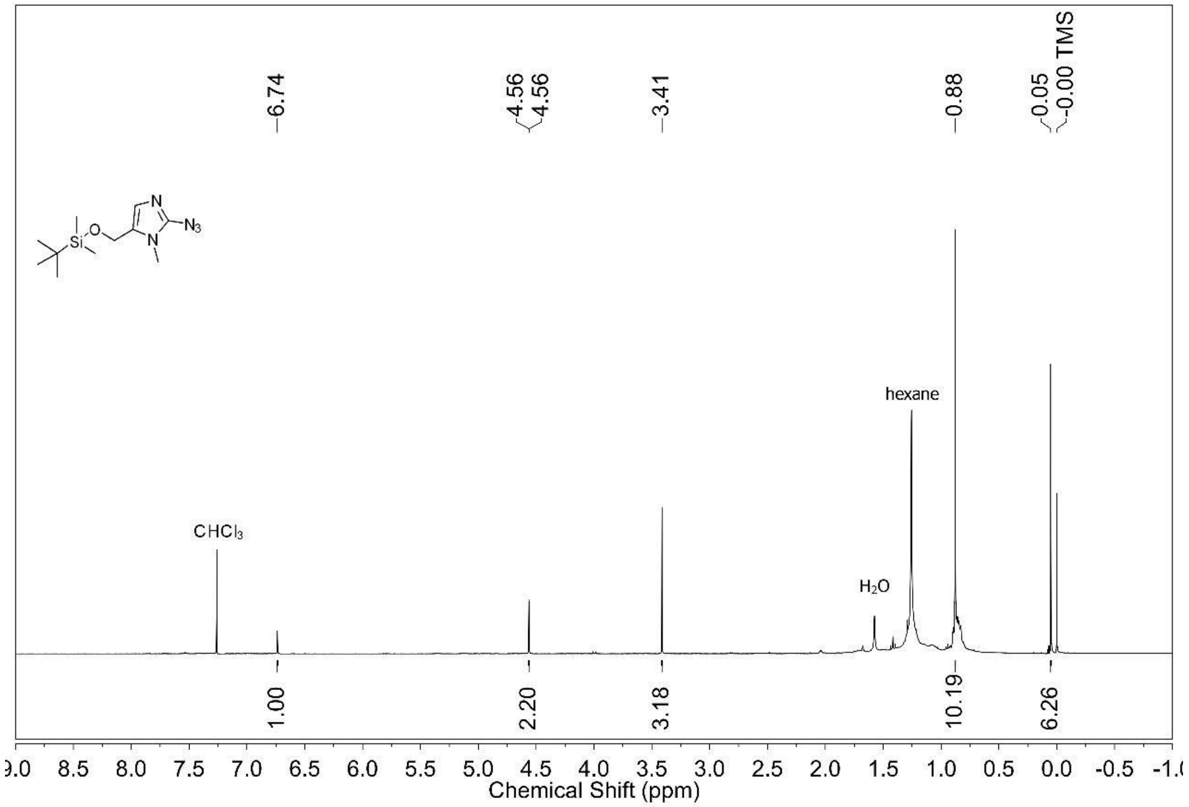


$^{1}H$ NMR spectrum (400 MHz, $CDCl_3$) of **11**

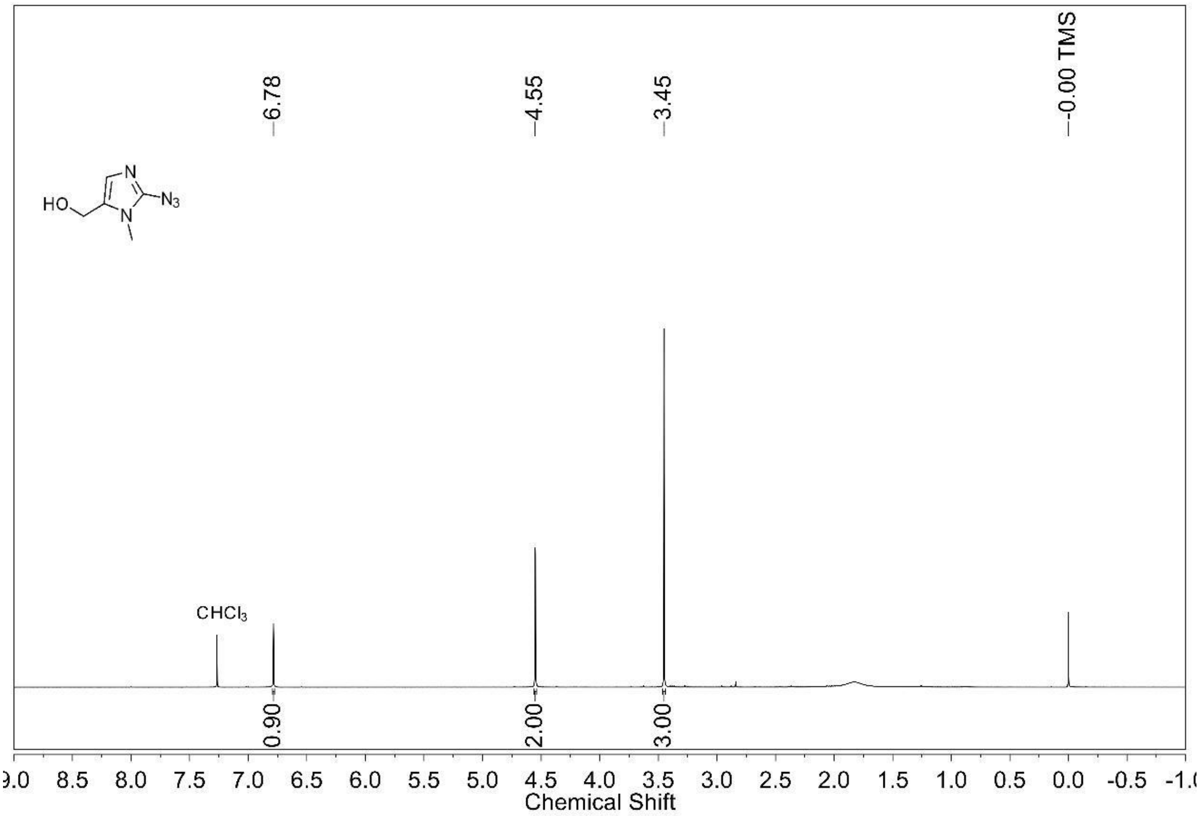

[1]H NMR spectrum (400 MHz, $CDCl_3$) of **12**

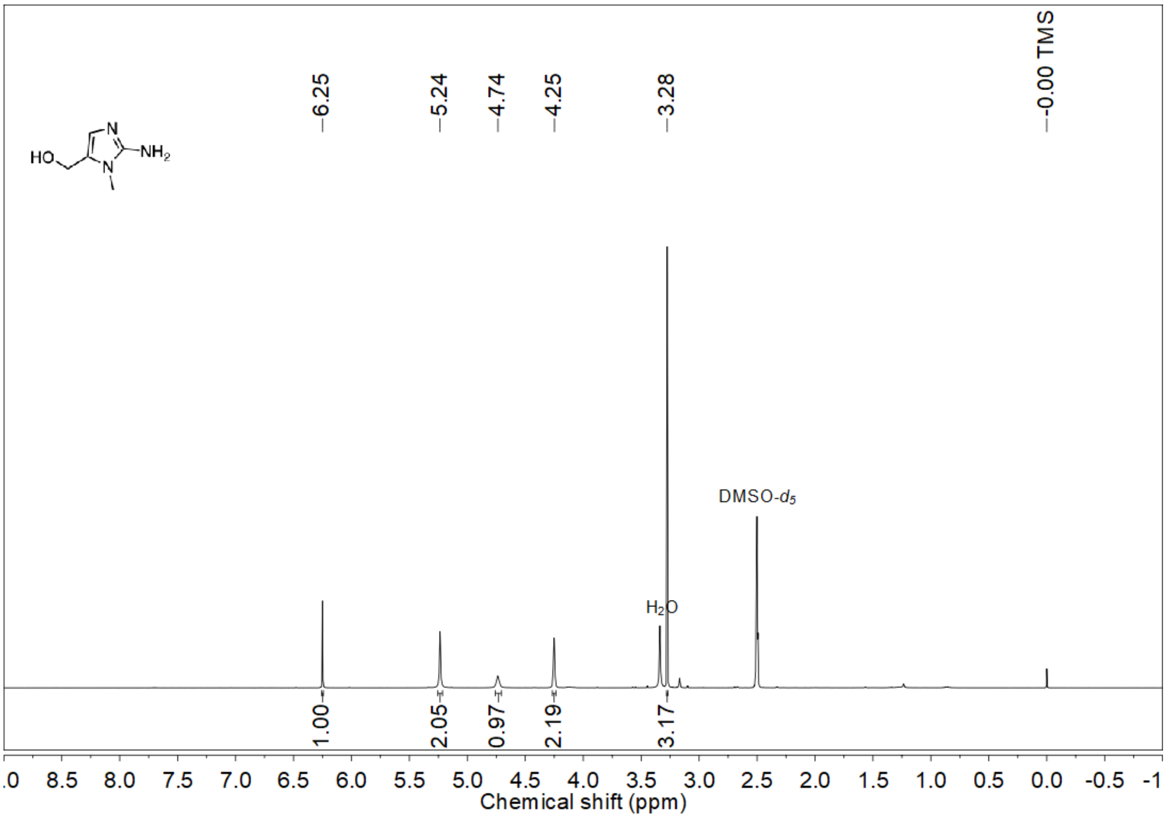


[1]H NMR spectrum (400 MHz, $CDCl_3$) of **13**

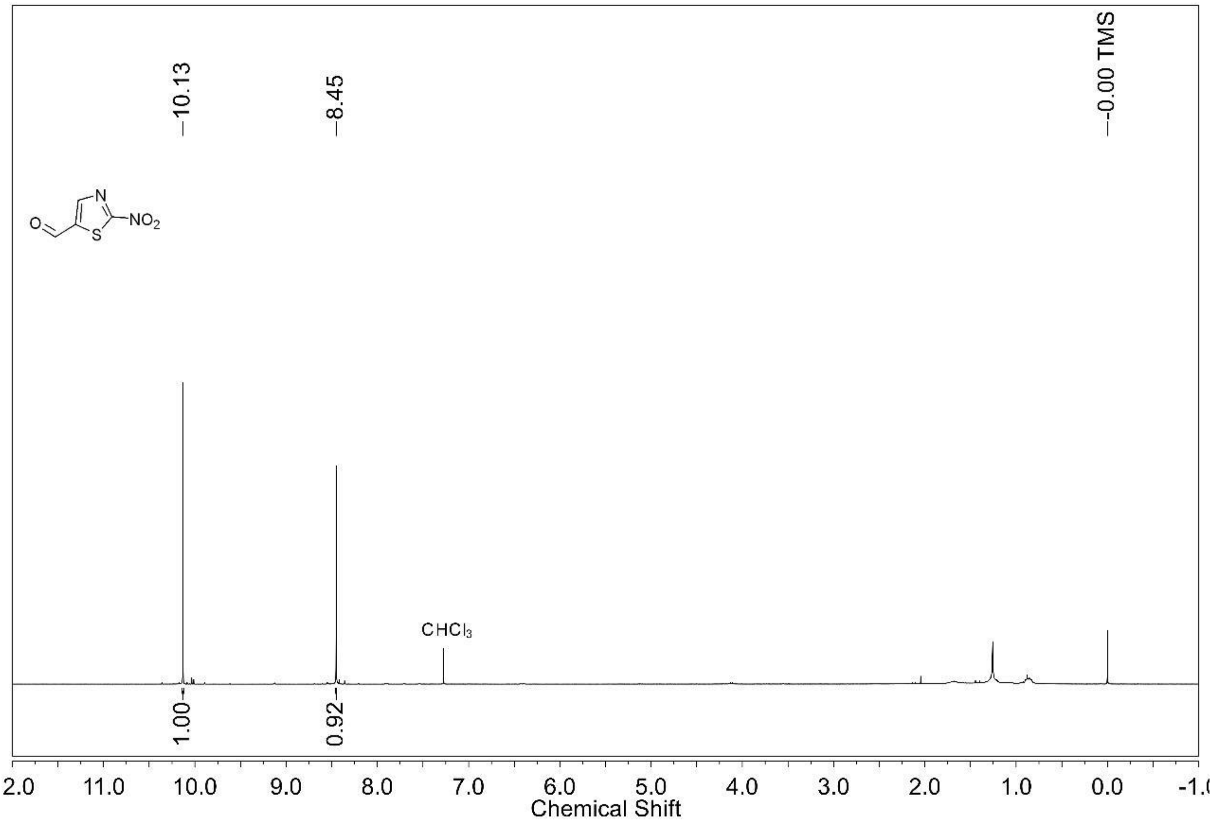

#### 8. HRMS spectra of compounds

HRMS spectrum of **1a**

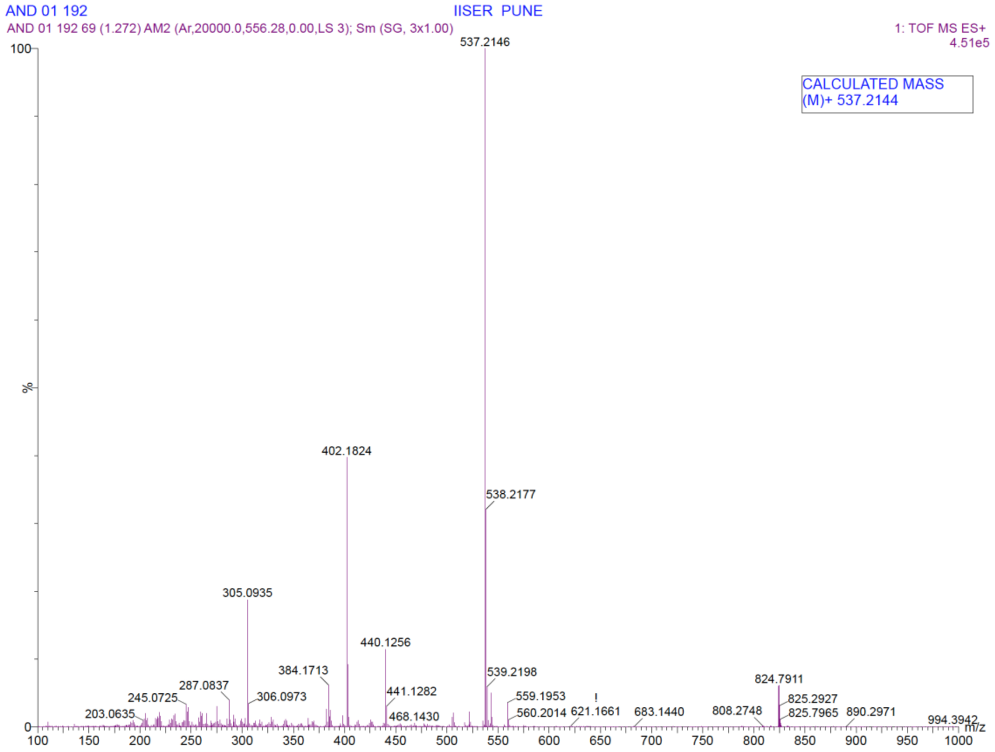


HRMS spectrum of **1b**

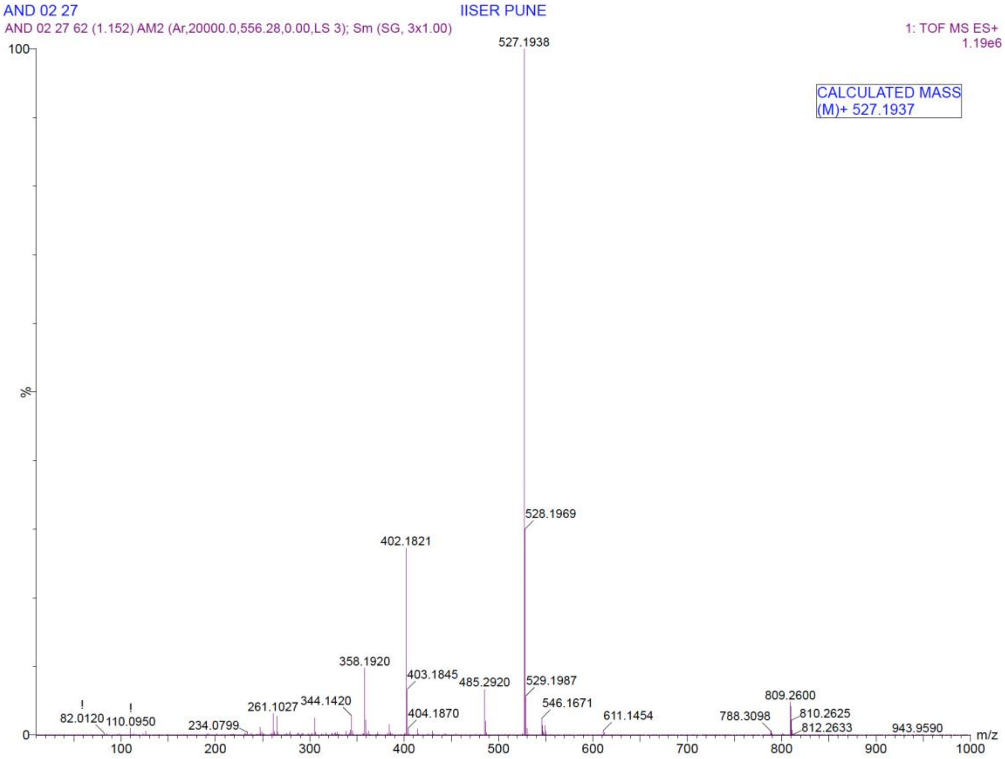

HRMS spectrum of **1c**

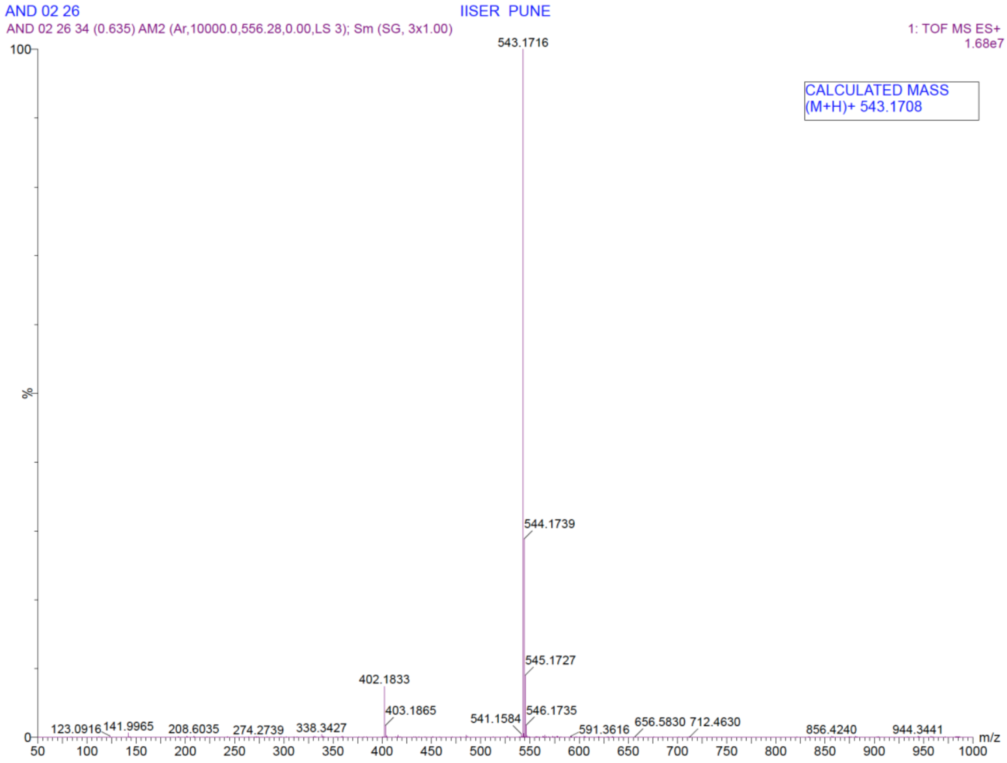


HRMS spectrum of **1d**

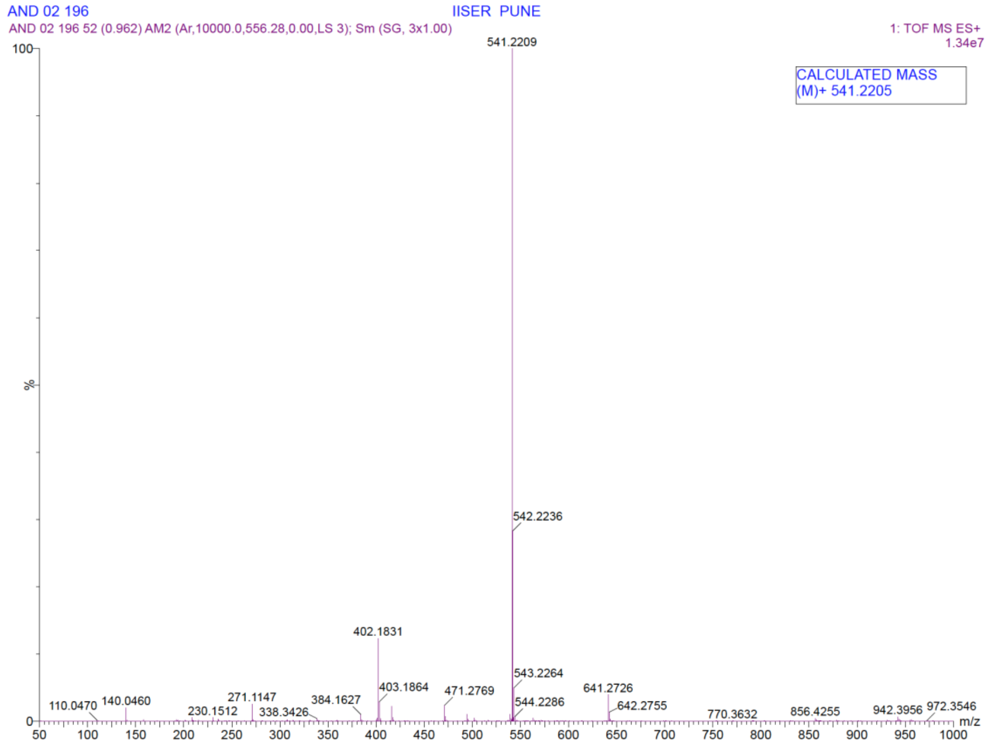

HRMS spectrum of **1e**

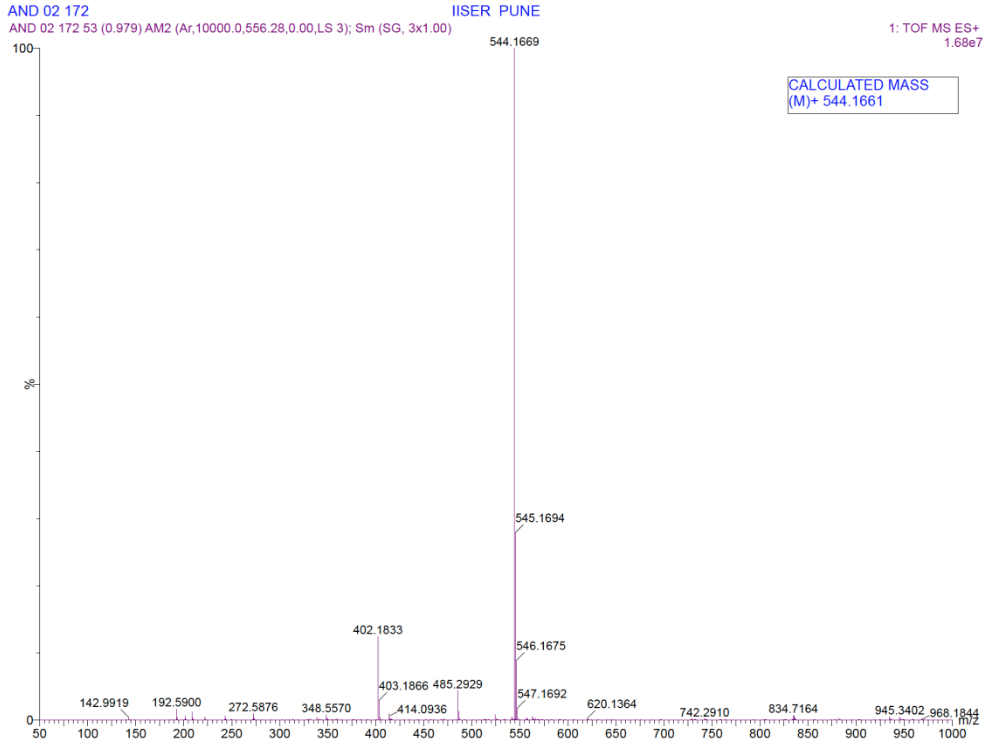


HRMS spectrum of **1f**

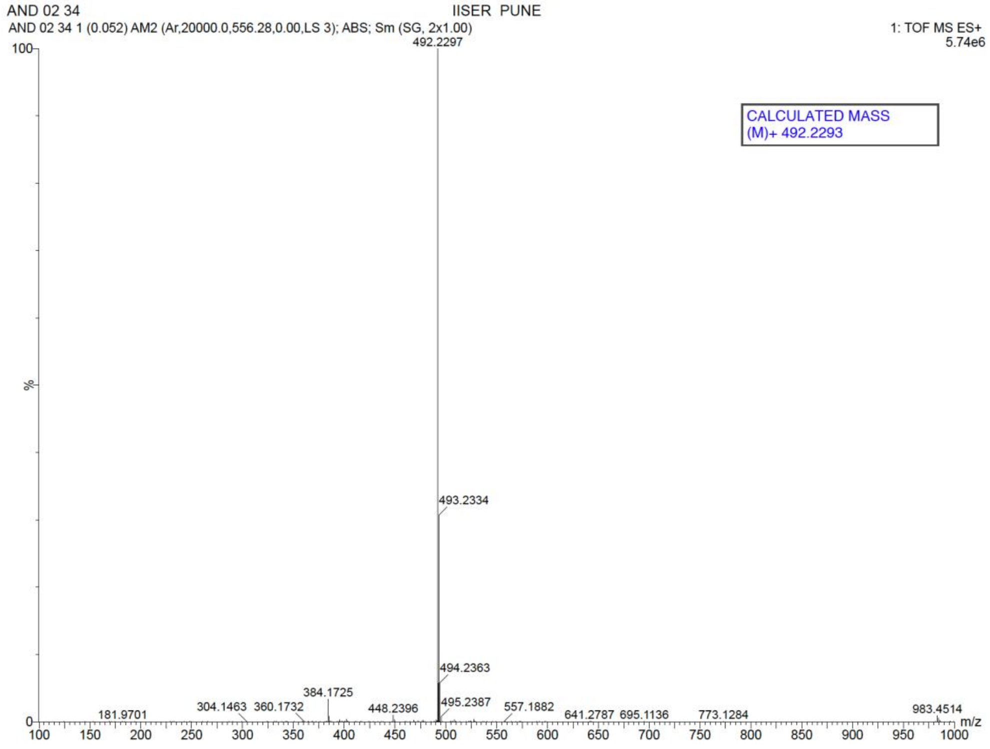

HRMS spectrum of **1g**

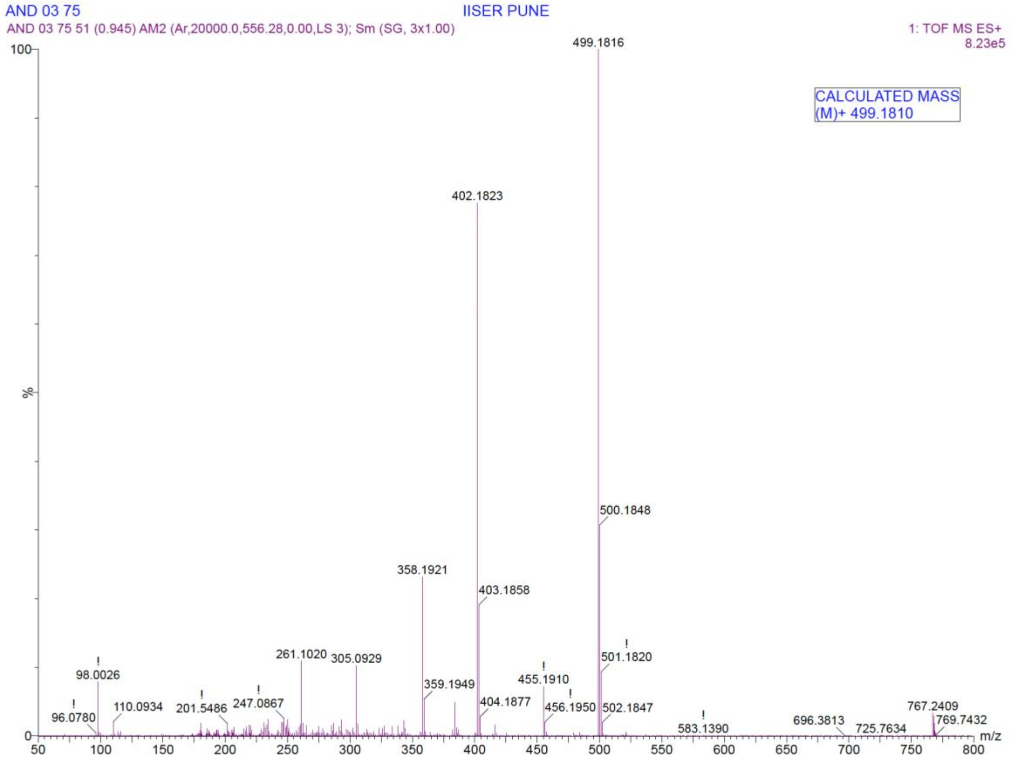


HRMS spectrum of **2a**

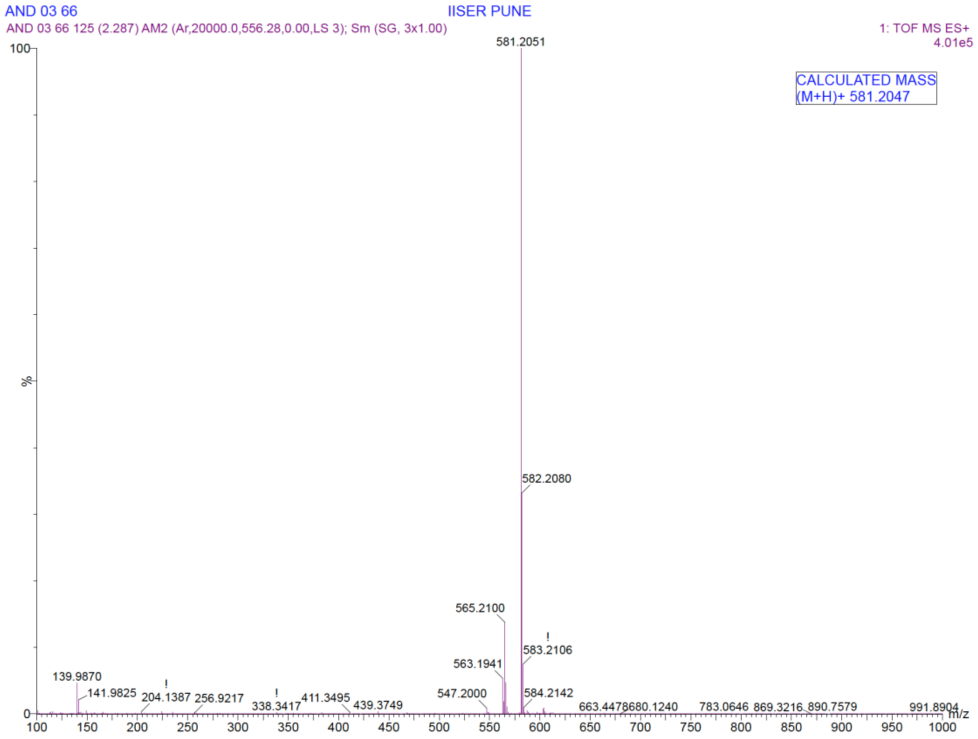

HRMS spectrum of **2b**

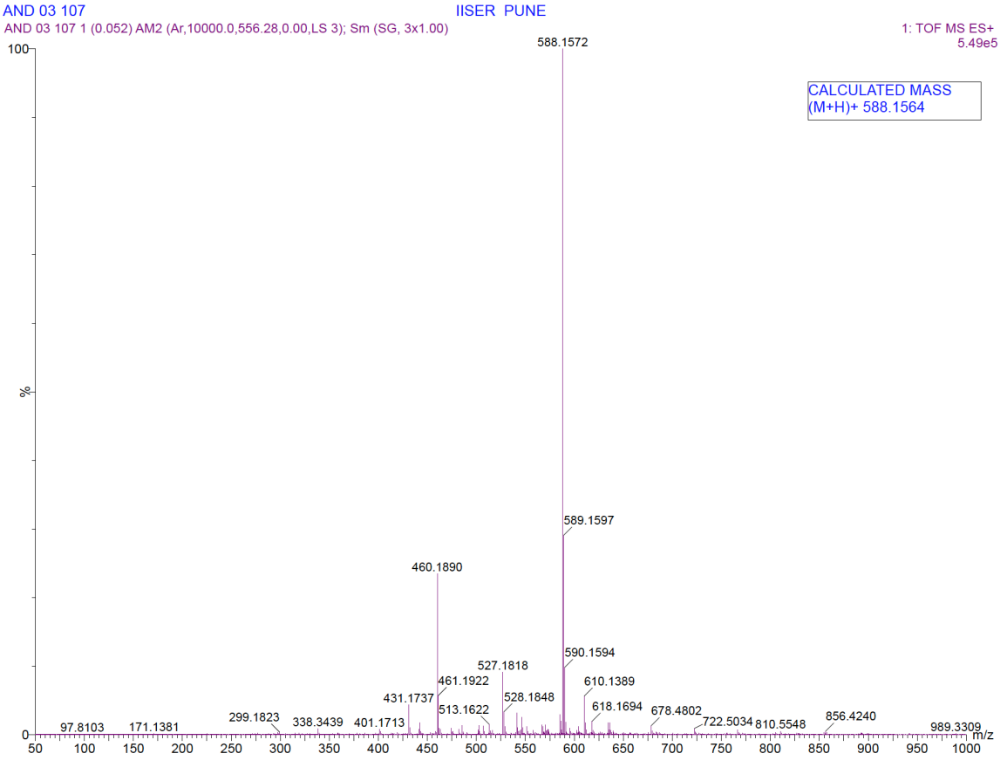
